# Discovery of prevalent, clinically actionable tumor neoepitopes via integrated biochemical and cell-based platforms

**DOI:** 10.1101/2022.10.27.513529

**Authors:** Hem Gurung, Amy Heidersbach, Martine Darwish, Pamela Chan, Jenny Li, Maureen Beresini, Oliver Zill, Andrew Wallace, Ann-Jay Tong, Dan Hascall, Eric Torres, Andy Chang, Kenny “Hei-Wai” Lou, Yassan Abdolazimi, Amanda Moore, Uzodinma Uche, Melanie Laur, Richard Notturno, Peter J.R. Ebert, Craig Blanchette, Benjamin Haley, Christopher M. Rose

**Author notes:** These authors contributed equally to this work (Hem Gurung, Amy Heidersbach, Martine Darwish).

## Abstract

Strategies for maximizing the potency and specificity of cancer immunotherapies have sparked efforts to identify recurrent epitopes presented in the context of defined tumor-associated neoantigens. Discovering these “neoepitopes” can be difficult owing to the limited number of peptides that arise from a single point mutation, a low number of copies presented on the cell surface, and variable binding specificity of the human leukocyte antigen (HLA) class I complex. Due to these limitations, many discovery efforts focus on identifying neoepitopes from a small number of cancer neoantigens in the context of few HLA alleles. Here we describe a systematic workflow to characterize binding and presentation of neoepitopes derived from 47 shared cancer neoantigens in the context of 15 HLA alleles. Through the development of a high-throughput neoepitope-HLA binding assay, we surveyed 24,149 candidate neoepitope-HLA combinations resulting in 587 stable complexes. These data were supplemented by computational prediction that identified an additional 257 neoepitope-HLA pairs, resulting in a total of 844 unique combinations. We used these results to build sensitive targeted mass spectrometry assays to validate neoepitope presentation on a panel of HLA-I monoallelic cell lines engineered to express neoantigens of interest as a single polypeptide. Altogether, our analyses detected 84 unique neoepitope-HLA pairs derived from 37 shared cancer neoantigens and presented across 12 HLA alleles. We subsequently identified multiple TCRs which specifically recognized two of these neoantigen-HLA combinations. Finally, these novel TCRs were utilized to elicit a T cell response suggesting that these neoepitopes are likely to be immunogenic. Together these data represent a validated, extensive resource of therapeutically relevant neoepitopes and the HLA context in which they can be targeted.

## Introduction

Neoantigen-specific T cells play a critical role in immune-mediated elimination of tumors and significant resources have been dedicated to developing clinically active drugs that amplify the cancer immunity cycle to improve the magnitude and breadth of elicited immune responses (Chen and Mellman, 2013; Rosenberg, 2014). Whether alone or paired with broad immune system activation, targeted immunotherapeutics may enable enhanced potency and safety profiles (Melero et al., 2015; Panchal et al., 2021). Further, T cells programmed to eradicate neoantigen expressing cells may facilitate design of next-generation cell therapies, particularly against solid tumors (Leidner et al., 2022; Yang and Rosenberg, 2016).

There are two broad categories of neoantigens, private neoantigens and shared neoantigens (Zhang et al., 2021). Private (a.k.a personalized) neoantigens represent the vast majority of mutations that arise during cancer progression and are somatic mutations unique to an individual’s tumor and not found across multiple patients or indications (Jhunjhunwala et al., 2021). Developing therapeutics against private neoantigens necessitates the creation of a personalized drug, a process that requires genomic analysis of a patient’s biopsy, HLA typing, and bioinformatic prediction of a small number of epitopes to target in the final therapeutic (Capietto et al., 2017; Lang et al., 2022). Within this workflow, optimization of the computational prediction algorithms remains an area of focus as epitopes predicted to bind may not be presented by a cell. An alternative approach, mass spectrometry based immunopeptidomics, can be used to confirm peptide presentation through direct peptide detection (Yadav et al., 2014), but has lacked adequate sensitivity to be used in a personalized approach. Despite these challenges, proof of concept studies have demonstrated the potential of such therapies (Ott et al., 2017; Sahin et al., 2017), but the road to broadly-available personalized neoantigen-targeting therapies has yet to be established.

In contrast to personalized neoantigens, shared neoantigens are recurrent mutations that are present across a wide scope of patients and indications (Zhang et al., 2021). The prevalence of shared neoantigens is related to their biological function and many putative shared neoantigens derive from oncogenic mutations within proteins such as KRAS, EGFR, TP53, and BRAF (Klebanoff and Wolchok, 2018). Prior knowledge of the specific mutation enables discovery and validation of target epitopes as well as a path towards “off the shelf” therapeutics that can be administered to any patient whose tumor bears the target mutation and appropriate HLA haplotype. Early examples of vaccines targeting shared neoantigens have shown promising preclinical efficacy, including vaccines targeting IDH1 (Schumacher et al., 2014), KRAS (Wang et al., 2016), and H3.3K27M (Chheda et al., 2018). Beyond vaccines, T cell therapies targeting shared neoantigens have shown clinical efficacy. For example, transfer of T cells specific to the KRAS G12D HLA-C*08:02 restricted neoepitopes were shown to produce an effective anti-tumor response in a human patient with lung metastatic tumors (Leidner et al., 2022; Tran et al., 2016). Efficacy has also been demonstrated in preclinical models for KRAS G12V/G12D HLA-A*11:01 restricted neoepitopes (Wang et al., 2016). Studies such as these have prompted development of vaccines and T cell therapies to additional frequent mutations found in cancer patients, however in these cases identifying shared neoantigen neoepitope and the HLA contexts in which they are presented remains a primary challenge.

Neoantigen specific T cell responses require the presentation of neoepitopes, peptides derived from mutated proteins, via cell surface-associated class I HLA molecules (HLA-I). T cell receptors (TCRs) interact with particular neoepitope-HLA complexes such that the therapeutic target definition comprises both the neoepitope sequence and HLA-I subtype upon which it is presented. Neoepitopes are generally 8-11 amino acids in length and as a result a single amino acid substitution may be presented within 38 possible neoepitopes (eight 8-mers, nine 9-mers, ten 10- mers, and eleven 11-mers). Additionally, HLA-I molecules are highly polymorphic with 1000s of documented variants that each have the capacity to bind a distinct subset of peptides. Consequently, the number of potential neoepitope-HLA targets increases quickly and even if development was focused on neoepitopes derived from the most common 50 cancer neoantigens across the most prevalent 15 HLA alleles, >28,000 neoepitope-HLA pairs could be formed. However, not all of these combinations are therapeutically relevant because even if a neoepitope can bind an HLA molecule, the specific neoepitope may not be processed and presented by HLA in the context of a tumor cell.

Generation of neoepitopes is dependent on the antigen processing pathway (APP) as cancer neoantigens are degraded by the proteasome and the resulting peptides are imported into the ER where they are further processed by ER resident aminopeptidases before finally being loaded into an HLA molecule for presentation (Pishesha et al., 2022). As a result, it is possible that a synthetic neoepitope peptide could bind an HLA molecule *in vitro*, but not be observed as a presented peptide within a cellular or *in vivo* context. A prime example is the description of a bi-specific antibody that targeted an A*02:01 restricted neoepitope of KRAS G12V (KLVVVGAVGV) (Douglass et al., 2021). This molecule demonstrated binding *in vitro* when HLA molecules were loaded with synthetic peptide, but failed to induce cell killing when tested in cell lines harboring the KRAS mutation. For this reason, direct identification of presented peptides through mass spectrometry (MS)-based immunopeptidomics approaches is a key aspect of neoantigen target validation. For example, a targeted MS approach was used to provide further evidence that the aforementioned KRAS A*02:01 neoepitope is not presented in a cellular context (Choi et al., 2021). While extremely sensitive, such targeted MS assays require heavy isotope-labeled peptides for each potential neoepitope as well as cell lines expressing the cancer neoantigen of interest. Due to these limitations, targeted MS assays are typically employed to evaluate a small number of cancer neoantigens within a given study.

Here, we present a pipeline for discovery and validation of neoepitope-HLA pairs presented on the surface of cells. Using a clinico-genomics approach we selected 47 common cancer point mutations and 15 prevalent HLA-I alleles to enable characterization of the neoepitope landscape for clinically-actionable neoantigen targets. We then employed a novel high throughput HLA binding assay to experimentally screen *in vitro* stabilization for 24,149 neoepitope-HLA combinations to identify 587 stable complexes (Darwish et al., 2021; Rodenko et al., 2006). We then compared these results to neoepitope-HLA pairs predicted to bind by NetMHCpan4.0 (Jurtz et al., 2017) and found that the two methods identified complementary sets of binding events. Consequently, neoepitope-HLA pairs considered for further analysis included those identified by the high throughput HLA binding screen as well as a subset of neoepitope-HLA pairs that were only predicted to bind by NetMHCpan4.0. The resulting neoepitope-HLA pairs were assayed for presentation using both untargeted and targeted mass spectrometry analysis of HLA-I monoallelic cell lines that simultaneously expressed ∼25 amino acid segments corresponding to each of the 47 cancer neoantigens. This analysis produced a list of 84 neoepitope-HLA pairs representing, to the best of our knowledge, the broadest list of experimentally validated presented neoepitopes. Lastly, to characterize the therapeutic potential for these targets we utilized TCRs discovered using the well-established Multiplex Identification of T cell Receptor Antigen (MIRA; Adaptive Biotechnologies) (Klinger et al., 2015) assay in a parallel therapeutic discovery effort to demonstrate mutant-selective T Cell activation and killing of cells expressing either an A*02:01 FLT3 D835Y or A*11:01 PIK3CA E454K neoepitope.

## Results

### Clinico-Genomics Analysis of Shared Cancer Neoantigens

The schematic highlighted in Figure 1 provides an overview of the workflow developed to enable neoepitope discovery in this study. We first identified the most common recurrent point mutations across cancer types from a large compendium of tumor and normal sequencing data (Chang et al., 2017), filtering at a per-indication case prevalence of 2%. For this analysis, gene fusions were excluded due to the high diversity of their breakpoints and resulting coding sequences. This led to a list of 36 shared cancer neoantigens (**Table S1**). Separately, we identified the most common HLA-I alleles across human populations, filtering at a carrier frequency of 10%. We also verified that these alleles were present at equivalent frequencies in patients from The Cancer Genome Atlas (TCGA). This additional filtering led to a list of 16 HLA-I alleles (**Table S2**). Co-prevalence of these shared cancer neoantigens and HLA-I alleles in the TCGA data was analyzed to ensure against biased co-expression of recurrent mutated genes and common HLA-I variants. We found that co-prevalence of each shared neoantigen matched expected values that were calculated as the product of the individual neoantigen and HLA-I allele prevalence. Together, these 36 neoantigens and 16 HLA-I alleles provided the foundation for development of our platform.

**Figure 1.**
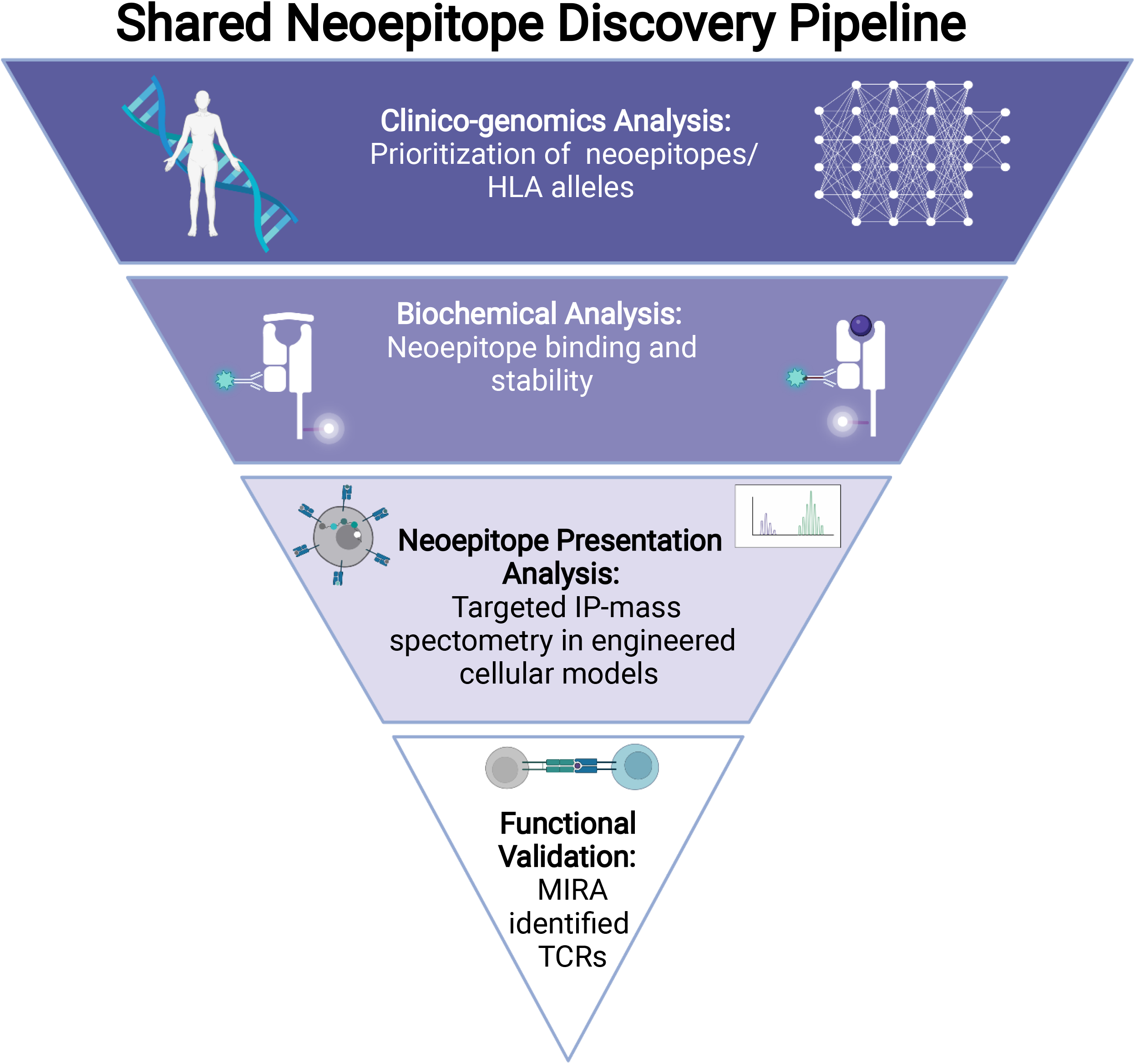
Overview of shared neoepitope discovery pipeline. The pipeline utilizes clinico- genomic analysis to select shared cancer neoantigens and HLA alleles to survey. Neoepitope-HLA pairs were then tested within a high-throughput binding assay. Binders were further interrogated within engineered cell lines expressing a 47-mer neoantigen cassette by untargeted and targeted immunopeptidomics to confirm presentation of neoepitope-HLA pairs. Select neoepitopes were then validated for immunogenic potential through discovery of specific TCRs that enable T cell activation and target cell killing.

### High throughput TR-FRET analysis of neoepitope exchange and neoepitope-HLA stability

T cell-mediated neoantigen-specific therapies require a neoepitope and HLA molecule to form a stable complex that can be presented on the surface of tumor cells. One strategy to determine if a peptide binds with a particular HLA allotype would be to perform an *in vitro* binding assay, either through peptide-HLA complex refolding (Harndahl et al., 2012) or peptide exchange into conditional HLA complexes containing a UV cleavable peptide (Darwish et al., 2021) While these assays cannot completely replicate peptide-HLA binding *in vivo, in vitro* binding affinity measurements have proven to be valuable data for training computational algorithms that predict peptide-MHC binding and presentation.

Given that our goal was to survey all potential neoepitopes from candidate cancer neoantigens across multiple HLA alleles, we developed a custom high-throughput (HTP) TR-FRET assay based on peptide-mediated stabilization of conditional HLA complexes to provide evidence of neoepitope-HLA complex formation (**Figure 2A**). We focused on the 36 prevalent shared cancer neoantigens identified by the clinico-genomic analysis described above, as well as a subset of 11 additional antigens used to further characterize the platform. Regarding HLA allele coverage, of the 16 HLA alleles identified in the in-silico screen, only 15 were available in the appropriate conditional HLA complex format needed for the TR-FRET assay (**Figure 2A)**. Consequently, the final TR-FRET assay was used to probe stable binding of neoepitopes from 47 shared cancer neoantigens across the 15 most prevalent HLA alleles, resulting in the characterization of 24,149 neoepitopes-HLA complexes (**Table S3**). Note, that the number neoepitope-HLA combinations is fewer than the 26,790 expected (15 alleles x 47 mutations x 38 mutation containing neoepitopes) due to overlapping neoepitopes sequences of some cancer neoantigens and technical issues with generation of a subset of reagents.

**Figure 2.**
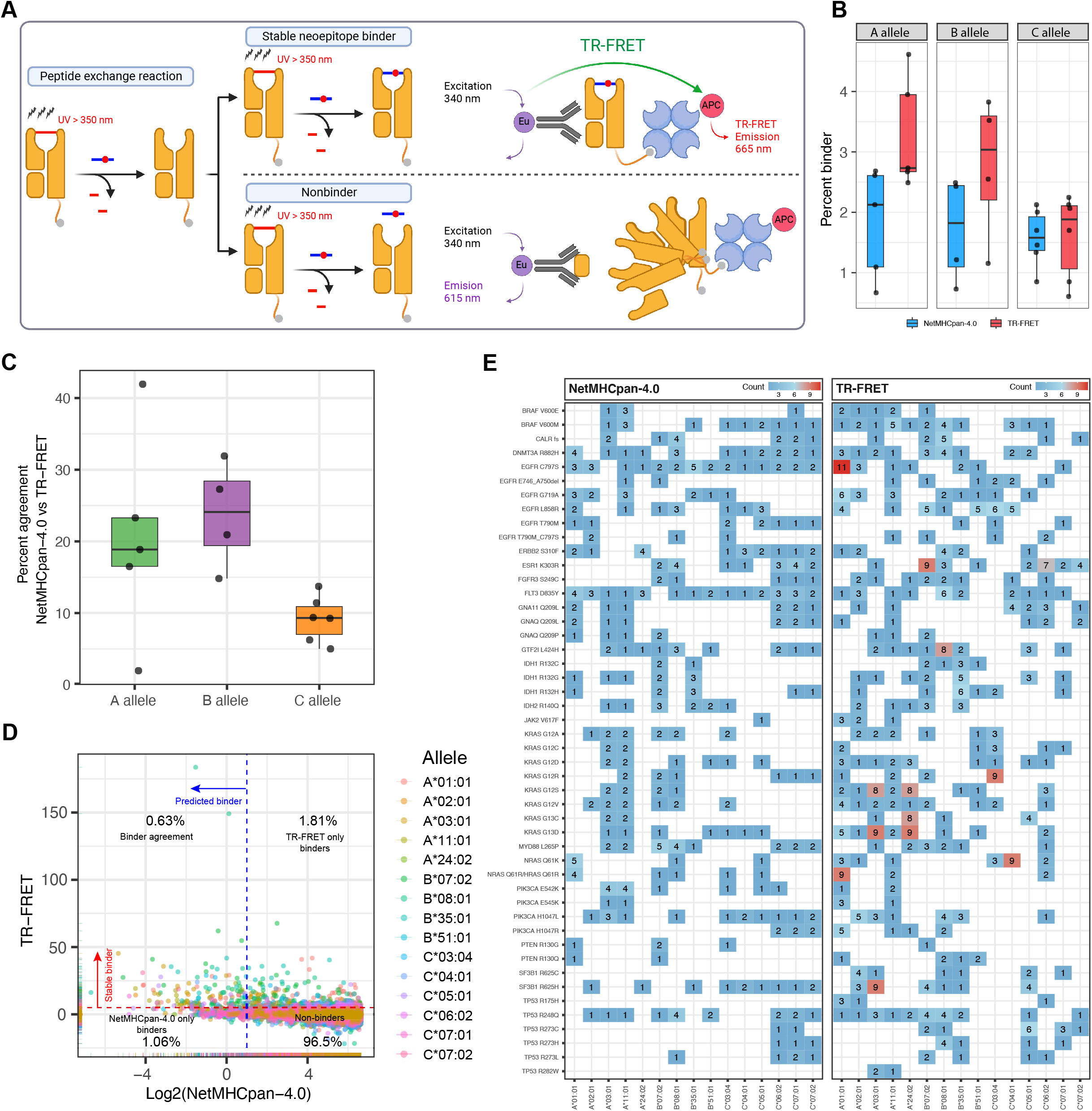
Characterization of neoepitope-HLA binding using a high-throughput TR-FRET assay and NetMHCpan4.0 prediction. **(A)** Schematic diagram of the TR-FRET assay used to measure stable neoepitope-HLA binding. **(B)** Percent of neoepitope-HLA combinations that were determined to be stable binders by TR-FRET (red bars) and NetMHC (blue bars) analysis across the HLA A, B and C alleles. **(C)** Percent of neoepitope-HLA pairs found to be binders by both NetMHC and TR-FRET across the A (green bar), B (purple bar) and C (orange) alleles. **(D)** Scatter plot of TR-FRET robust Z-score and Log2 NetMHC percentile rank. The dashed red line represents the cutoff for stable binders as measured by TR-FRET, where values higher than the red line are considered a stable binder. The dashed blue line represents the cutoff for binders based on NetMHC analysis, where values lower than the blue line are considered binders. **(E)** The number of neoepitope-HLA combinations found to be stable binders for each neoantigen (y- axis) across all 15 alleles (x-axis) for the TR-FRET and NetMHC analysis.

The TR-FRET assay utilized previously described conditional HLA ligands, peptides containing UV cleavable non-natural amino acids, to create conditional HLA complexes (HLA alpha chain, and Beta-2-microglobulin [β2M]) for the 15 HLA alleles (Darwish et al., 2021). Conditional HLA complexes were then incubated with a neoepitope of interest at 100-fold molar excess and exposed to UV light for 25 minutes. This reaction was expected to cleave the conditional ligand and convert the peptide from a stable high affinity “binder” to an unstable binder that dissociates from the HLA groove. In the presence of a binding neoepitope, peptide exchange occurred and stabilized the HLA complex (**Figure 2A, top**). In the presence of a non-binding neoepitope, peptide exchange did not occur, and the HLA complex dissociated (**Figure 2A, bottom**). To ensure the neoepitope-HLA complex remained stable at physiological temperatures, the samples were heated at 37°C for 24 hours prior to analysis. Complex stability was monitored using fluorescence of a TR-FRET donor (europium) conjugated to an anti-β2M antibody and a TR-FRET acceptor conjugated to streptavidin, which bound to a biotinylation of the HLA alpha chain. In these assays, a TR-FRET signal will only occur if the HLA complex remains intact due to the presence of a binding neoepitope. TR-FRET signals were quantified based on the ratio of relative fluorescent units and signals were subjected to a double normalization to generate a robust Z-score (RZ-score) for neoepitope comparison and ranking as described in the material and methods. For our analysis we considered any neoepitope-HLA combination with a RZ-score ≥ 5 to be a “stable binder”. This highly conservative cutoff was selected because all known binders fell within this range and to increase confidence that the combinations identified using this cutoff value were stable binders. When using this cutoff, 587 unique neoepitope-HLA pairs were classified as stable binders.

Computational algorithms that can predict neoepitope-HLA complex formation and presentation have become an integral part of personalized cancer therapies (Ott et al., 2020; Sahin et al., 2017). As mentioned above, these algorithms are trained with *in vitro* binding affinity measurements as well as mass spectrometry data to predict either binding or presentation of neoepitopes in the context of a particular HLA. To understand how TR-FRET results compared to computational prediction methods, we employed NetMHCpan-4.0 (hereafter NetMHC) to predict neoepitope presentation of the 24,149 neoepitope-HLA pairs assayed by TR-FRET. For this analysis the “eluted ligand” percentile rank (%Rank) values were used to determine if a neoepitope was a “binder” (%Rank ≤ 2), resulting in 408 unique predicted neoepitope-HLA pairs.

When measured as a percentage of all potential neoepitope-HLA complexes, TR-FRET generally identified more stable binders as compared to NetMHC, particularly for HLA-A and HLA-B alleles (**Figure 2B, Supplemental Figure S1C**). Examination of specific neoantigen HLA combinations revealed instances where concordance of TR-FRET and NetMHC was high, for example both approaches called the same two KRAS G12R neoepitopes as binders to B*07:02 (**Supplemental Figure S1A**). However, because TR-FRET generally called more neoepitope-HLA pairs, there were several cases where NetMHC did not predict the same binding events identified by TR-FRET. This was exemplified by ESR1 K303R where TR-FRET identified 9 stable binding neoepitopes and NetMHC only predicted 2 of those instances (**Supplemental Figure S1B**).

To better understand an overall correlation between NetMHC prediction and TR-FRET measurements, we compared the percent agreement when classifying binders and non-binders across the two methods (**Figure 2C, Supplemental Figure S1D and E**). When considering all potential neoepitope-HLA pairs, agreement of both methods for classification of binders and non-binders was strong, ranging between 95.1 - 98.6% depending on the allele (**Supplemental Figure S1D**). However, if only the identification of binders was considered, agreement between the two methods dropped significantly to 2.04 - 40% (**Figure 2C**). Agreement was generally higher for HLA-A and HLA-B alleles as compared to HLA-C alleles, however A*24:02 exhibited the lowest agreement at 2.04% (**Figure 2C and Supplemental Figure S1E**). When this same analysis was performed only considering non-binders, the agreement was once again very strong, ranging from 94.1 - 98.4% (**Supplemental Figure S1F**). These results demonstrate that both methods generally agree on non-binding events, whereas positive interactions are found with minimal overlap. This substantiates the power of combining both approaches to identify and prioritize unique neoepitope-HLA pairs for further characterization.

To better visualize the complementarity of binder identification by TR-FRET and NetMHC, the TR-FRET RZ-score and NetMHC %Rank were plotted for all candidate neoepitope-HLA pairs (**Figure 2D**). From this analysis we found that 0.63% of all candidate neoepitope-HLA pairs were found to be binders by both methods and that agreement varied from 0.06% to 1.49% of all potential neoepitope-HLA pairs when data were viewed at the allele level (**Figure 2D and Supplemental Figure S2**). Interestingly, each method identified nearly the same percentage of additional binding events for neoepitope-HLA pairs, 1.06% for NetMHC and 1.81% for TR-FRET, demonstrating that each method has the potential to identify unique binding combinations (**Figure 2D**). To more clearly visualize discrepant binders between TR-FRET and NetMHC, a heatmap was generated that displays the number of binders for each HLA allele and neoantigen combination (**Figure 2E**). Based on this analysis there are clear areas of overlap as well as significant gaps between the different approaches. For example, TR-FRET identified 11 epitopes as binders for EGFR C797S and A*01:01 whereas NetMHC only predicted 3 binding epitopes (**Figure 2E**). These findings highlight the power of our high-throughput biochemical assay to identify a potentially complementary set of neoepitope-HLA pairs and suggest using both methods together could lead to more comprehensive neoepitope discovery.

### Generation of HLA-I monoallelic cell lines co-expressing 47 shared cancer neoantigens

Despite observed peptide-HLA stabilization *in vitro*, expression and processing of a mutated protein may not result in presentation of a neoepitope in a cellular context (Garstka et al., 2015; Jappe et al., 2018; Skora et al., 2015). For this reason, validation of candidate neoepitopes typically requires genetic expression of target neoantigens followed by a readout of association with surface-bound HLA. The process of neoantigen-HLA discovery has been enhanced through the use of engineered “HLA monoallelic” cell lines, although these have relied to a large degree on endogenous mutant protein expression or expression of relatively few mutant transgenes, thus limiting throughput (Abelin et al., 2017; Bear et al., 2021; Wang et al., 2019).

We anticipated that simultaneous encoding of the mutated sequences (∼25 amino acids) for all 47 candidate neoantigens within a single HLA-null cell line would dramatically improve throughput of cell line generation and subsequent validation of TR-FRET identified neoepitope-HLA pairs by targeted mass spectrometry (**Figure 3A**). The HMy2.C1R lymphoblast cell line (hereafter C1R) was chosen as the model system due its lack of endogenous HLA-A and HLA-B protein as well as the ease of handling and expansion of suspension cells in culture. To generate the C1R^HLAnull^ cell line, the remaining HLA-C allele (HLA-C*04:01) was disrupted using CRISPheR/Cas9 and the HLA-null population was enriched by cell surface stain using a pan-HLA-I antibody and fluorescence activated cell sorting (FACS) (**Supplemental Fig 3 A,B**). There is precedence in the literature suggesting expression of B*35:03 in the C1R cell line (Schittenhelm et al., 2014), but we did not observe evidence of the B*35:03 motif in our immunopeptidomic analysis of resulting C1R^HLAnull^ cells.

**Figure 3.**
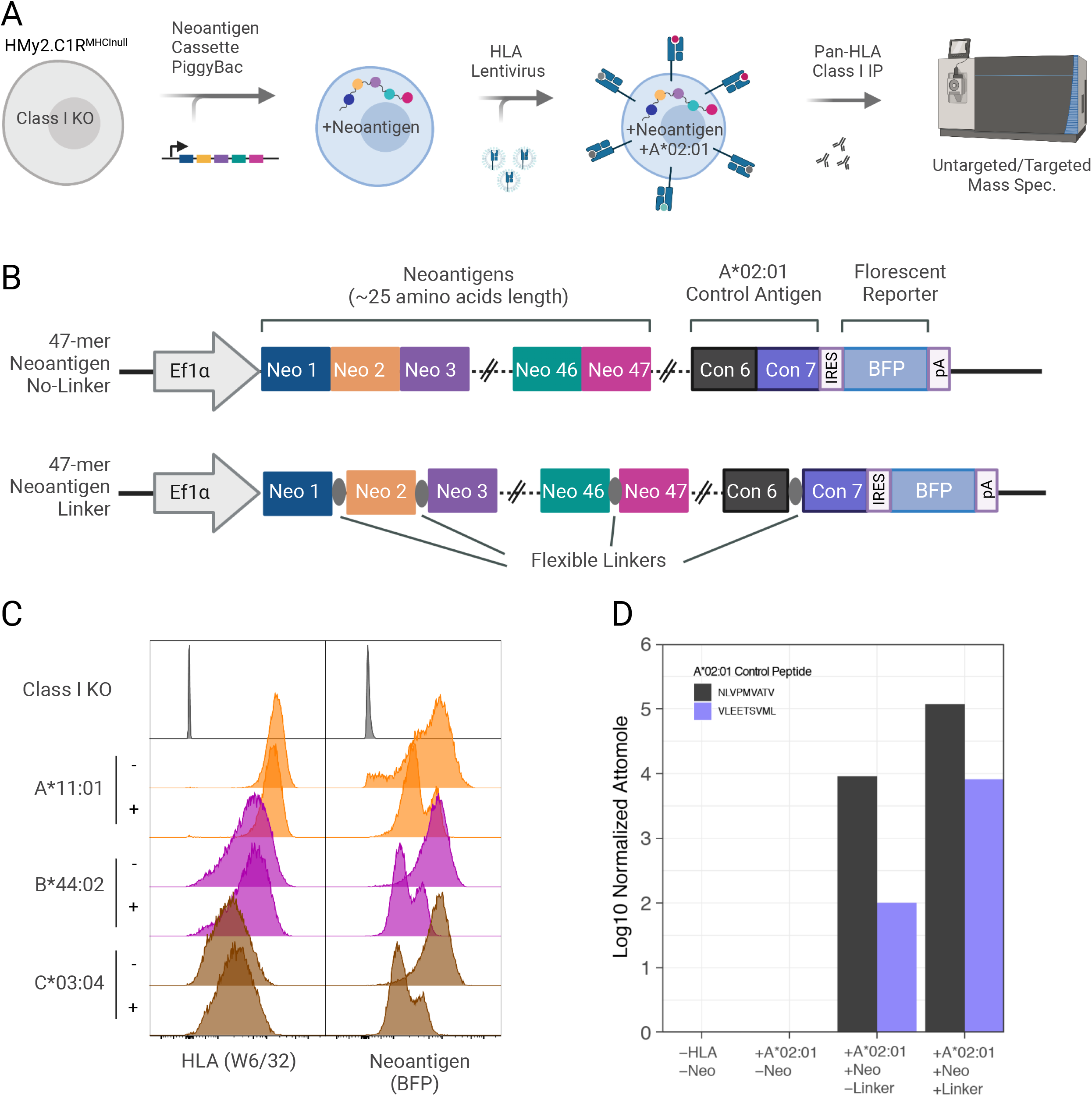
Generation of HLA class I monoallelic cell lines that stabley express a polyantigen cassette containing 47 shared cancer neoantigens. **(A)** Process overview for the cell engineering and mass spectrometry analysis of peptides presented by polyantigen expressing HLA-I monoallelic cell lines. **(B)** Vector map of the piggyBac polyantigen expression constructs utilized in this study. A single transcript containing 47 tandem neoantigens followed by 7 control peptides and an IRES linked mTagBFP2 (BFP) reporter is driven by a pol II EF1 alpha promoter. Neoantigens were either directly concatenated (No-Linker) or interspersed by short flexible linker sequences (Linker). **(C)** Flow cytometric detection of HLA-I expression (W6/32 antibody) and polyantigen cassette reporter (BFP) in selected cell lines or class I knockout parental line. Here, - indicates the absence of linkers and + indicates presence of linkers in the neoantigen construct. **(D)** Targeted immunopeptidomic detection of two neoantigen control peptides from pp65 (NLVPMVATV) and IE-1 (VLEETSVML) known to be presented by the A*02:01 allele.

Following development of a C1R^HLAnull^ cell population, synthesis and delivery of piggyBac expression vectors enabled stable transgene integration. Local sequence context has the potential to affect antigen processing (Gomez-Perosanz et al., 2020), and to address this possibility we generated two foundational cell lines that co-expressed mutated sequences (∼25 amino acids) for all 47 prioritized neoantigens in distinct configurations (**Figure 3B**). These two cell lines differed by the presence or absence of short amino acid linker sequences between most neoantigen segments within the polyantigen cassette; hereafter referred to as linker and no-linker cell lines, respectively (**Figure 3B**). Stable linker and no-linker cell populations were enriched using a separate TagBFP2 (BFP) marker to select for transgene-positive C1R^HLAnull^ cells. HLA-I monoallelic cells were then created by the introduction of the 15 HLA alleles as individual transgenes through stable lentiviral transduction of the linker and no-linker neoantigen-expressing C1R^HLAnull^ cell lines, resulting in 30 total cell populations for further analysis. HLA expression was confirmed by cell surface stain using a pan-HLA antibody (**Figure 3C and Supplemental Fig 3C,D**). To validate the functionality of our polyantigen cassettes, the linker and no-linker neoantigen constructs contained a set of control antigens with epitopes known to be presented by of A*02:01. HLA immunopeptidomics confirmed the presence of these peptides in both the linker and no-linker HLA-A*02:01-engineered cells (**Fig 3D**).

### Engineered polyantigen cassettes augment neoepitope presentation

One potential concern with this approach was that epitopes derived from a construct containing 47 concatenated neoantigen sequences of ∼25 amino acids may not reflect epitopes derived from a full-length mutant protein. We chose to investigate this further in the context of KRAS due to the recent description of A*11:01 restricted 9-mer and 10-mer neoepitopes detected by a targeted proteomic assay (Bear et al., 2021). To this end, we developed three C1R^HLAnull^ cell lines expressing HLA-A*11:01 as well as a doxycycline (dox)-inducible, full-length, wildtype, G12C, G12D, or G12V mutant KRAS protein, and compared the neoepitope presentation from these cell lines with a cell line expressing the no-linker variant of the polyantigen cassette.

We first confirmed mutant protein expression using a whole-cell targeted proteomic assay comprising a peptide that can detect total KRAS as well as three unique peptides that measured individual KRAS mutants (**Figure 4A**). This analysis validated dox-induced over-expression of the KRAS alleles by demonstrating an increase in total KRAS detected when dox was added to the culture medium (**Figure 4A**). Note, little to no signal was found at the steady-state protein level for these mutant peptides within the cell line containing a polyantigen cassette.

**Figure 4.**
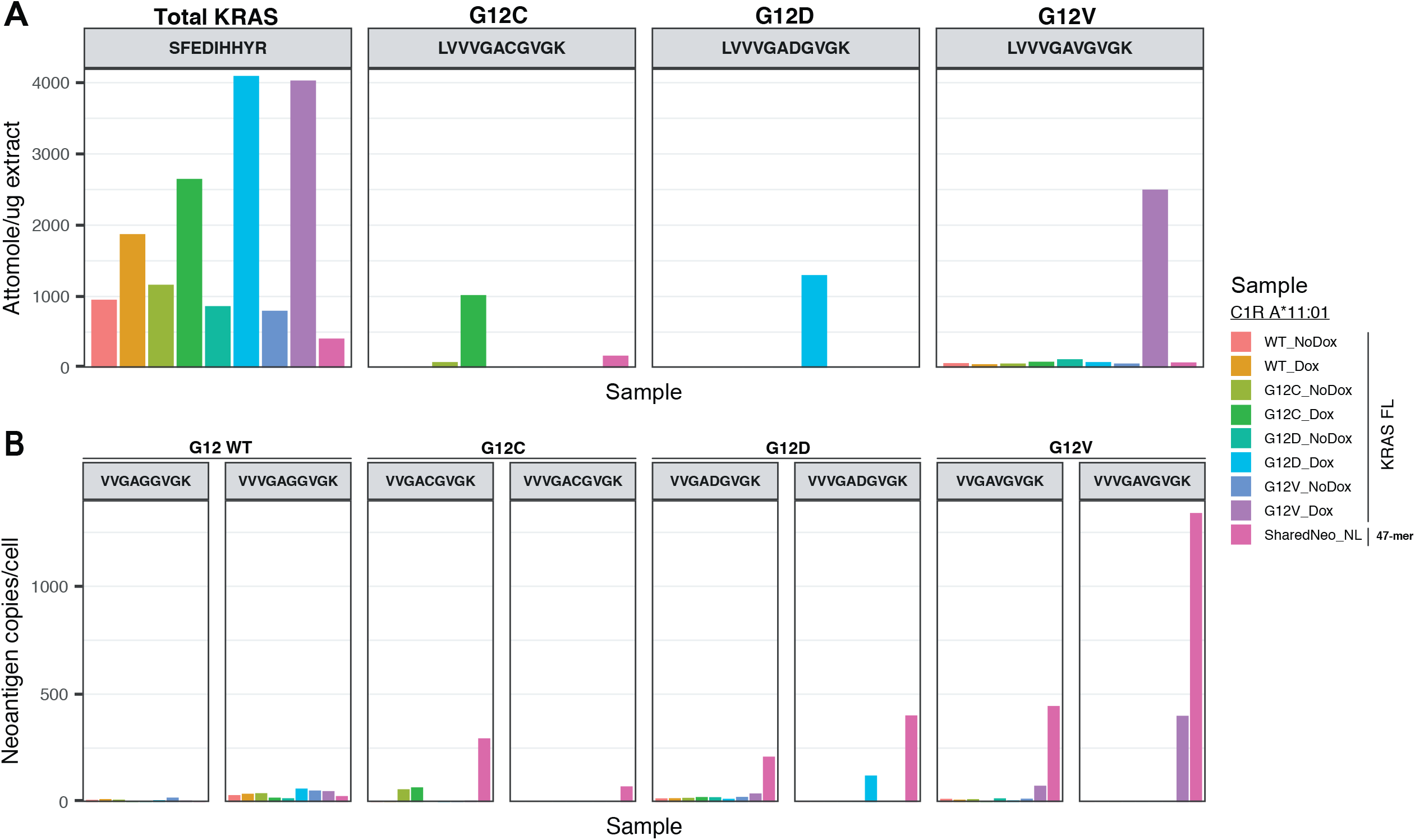
Characterization of protein and neoepitope abundance for C1R A*11:01 monoallelic cells expressing full length KRAS or the polyantigen cassette. For these analyses A*11:01 monoallelic cells were engineered to express doxycycline (dox)-inducible full length KRAS mutant proteins (G12C, G12D, G12V). These were compared against an A*11:01 monoallelic cell line containing the no-linker polyantigen cassette was used. (A) Absolute amount of KRAS WT and mutant proteins in the cell lysate by targeted mass spectrometry. (B) Copies per cell of presented KRAS 9-mer (VVGAXGVGK) and 10-mer (VVVGAXGVGK) neoepitopes as measured by A*11:01 monomers containing heavy synthetic neoepitope peptides spiked in prior to affinity purification and targeted mass spectrometry.

We then employed a targeted immunopeptidomic assay to quantify the level of presentation of previously identified 9-mer and 10-mer KRAS epitopes within the same cell lines described above (**Figure 4B**). For the cell lines containing a full-length mutant protein, induction of neoepitope presentation was observed for both G12V epitopes as well as the 10-mer epitope of G12D (**Figure 4B**). A weak signal was detected for the 9-mer epitope of G12C in both the control and dox-treated cell line. This could have been due to leaky/basal promoter activity as a weak signal for G12C was also detected at the protein level in both conditions (**Figure 4A-B**). Interestingly, the 9-mer and 10-mer epitopes of all KRAS mutants were detected in the cell line expressing the polyantigen cassette. Further, the polyantigen-modified cell line also presented higher absolute copies per cell of KRAS mutant epitopes as compared to cells expressing full length protein (**Figure 4B**). When combined with the lack of detection of mutant KRAS peptides at the protein level (**Figure 4A**), these results suggested the protein product of the polyantigen cassette is likely unstable and efficiently degraded such that epitope presentation is enhanced. This effect has been demonstrated previously in systems where induced protein degradation was used to increase presentation of epitopes derived from the degraded protein (Jensen et al., 2018; Moser et al., 2018). Therefore, monoallelic cells containing the polyantigen cassette provided both a higher-throughput and more sensitive system for discovery of neoepitopes from shared cancer neoantigens.

### Detection of neoepitope presentation on cell lines co-expressing 47 shared cancer neoantigens

After the generation of monoallelic cell lines containing either a linker and no-linker polyantigen cassette, peptide presentation was validated by HLA immunoprecipitation followed by both untargeted and targeted mass spectrometry (MS) analysis. Untargeted MS analysis enabled unbiased identification of peptides from the entire immunopeptidome, including peptides derived from the neoantigen constructs, but was limited in the sensitivity of neoepitope detection. Targeted analysis enabled sensitive detection of peptides presented at low copies per cell, but was constrained to peptides identified as binders within the TR-FRET assay as well as select peptides predicted to bind by NetMHC.

For untargeted MS analysis of each monoallelic cell line, the resulting data were searched using PEAKS (Zhang et al., 2012) against a human proteome database appended with the linker and no-linker 47-mer neoantigen construct and BFP sequences (**Figure 5A**). From this analysis we identified 218 to 6,663 unique 8-11 mer peptides within each monoallelic cell line, with the largest number of peptides identified in HLA-A alleles and the smallest numbers of peptides identified in HLA-C alleles (**Figure 5B**). Furthermore, the number of 8-11 mer peptides and general sequence features for each allele overlapped regardless of the polyantigen linker status and confirmed that presented peptides fit expected motifs, where known (**Supplemental Figures S4-S5**).

**Figure 5.**
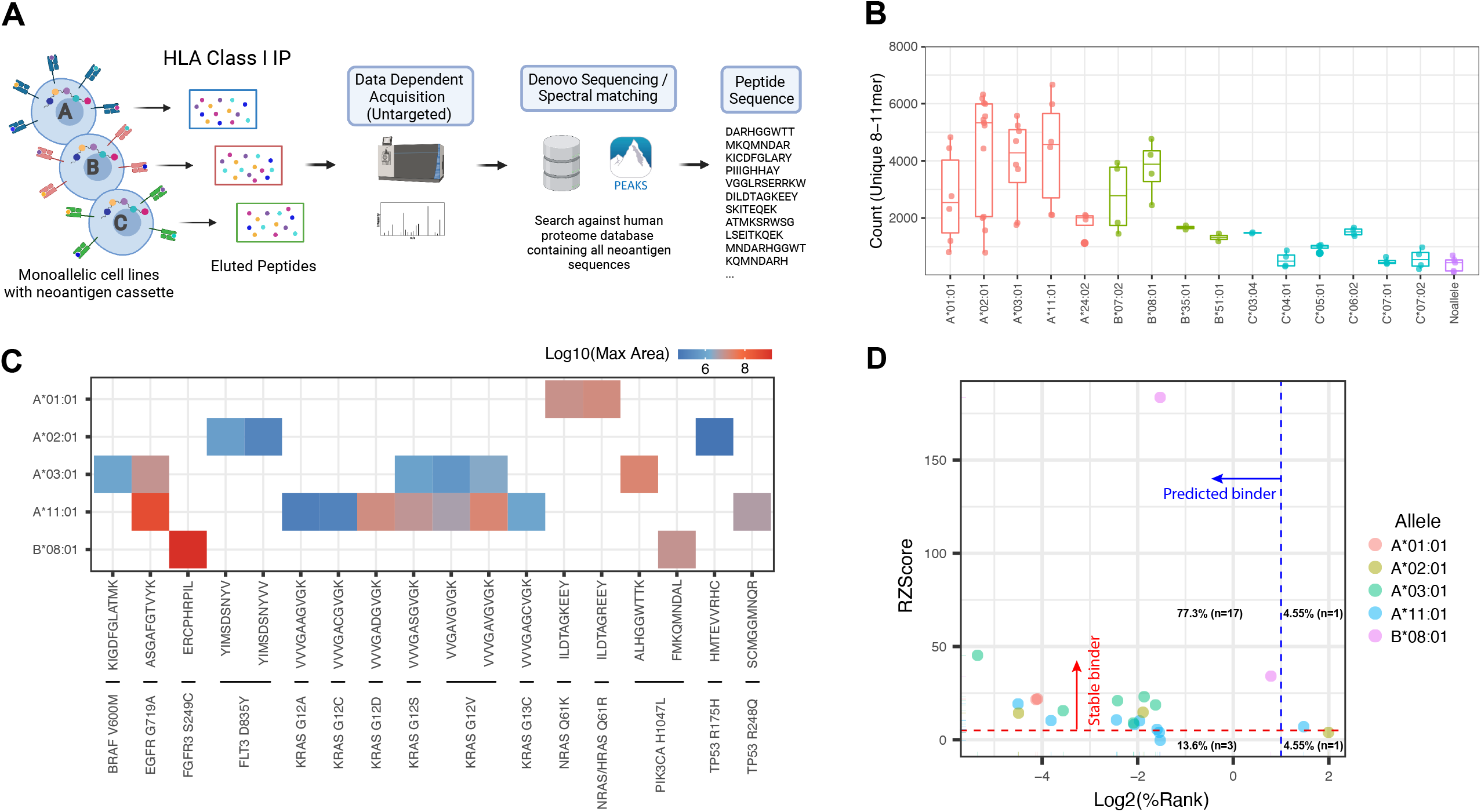
Untargeted immunopeptidomic analysis of monoallelic cell lines expressing the polyantigen cassette. **(A)** Workflow for untargeted immunopeptidomic analysis of monoallelic cell lines containing the polyantigen cassette. **(B)** Number of unique 8-11 mer peptides identified in untargeted immunopeptidomic analysis. **(C)** Identified shared cancer neoantigen epitopes. The color scale represents the log10 largest area across multiple analyses. **(D)** Comparison of TR-FRET Robust ZScore (RZScore) and NetMHC percentile rank (%Rank) score for each epitope identified through untargeted immunopeptidome analysis.

Through untargeted analysis we found 22 neoepitope-HLA pairs from 15 shared neoantigens across 5 HLA alleles representing ∼5.4% of neoepitope-HLA pairs predicted by NetMHC and ∼3.7% of neoepitope-HLA pairs identified within the TR-FRET assay (**Figure 5C, Table S4**). For untargeted analysis intensity measurements cannot be directly converted to absolute abundance, but neoepitopes from EGFR G719A (ASGAFGTVYK) and FGFR3 S249C (ERCPHRPIL) demonstrate much larger intensities as compared to other detected neoepitopes. For these 22 neoepitope-HLA pairs TR-FRET and NetMHC showed excellent concordance as 17 were identified as binders by both approaches (**Figure 5D**). However, there were also 1 and 3 neoepitope-HLA pairs uniquely identified as hits by TR-FRET and NetMHC, respectively, demonstrating each approach has the ability to predict distinct subsets of neoepitopes (**Figure 5D**). The final neoepitope-HLA pair identified by untargeted MS was derived from TP53 R175H (HMTEVVRHC) and was detected on A*02:01. This neoepitope-HLA pair had a RZ-score of 3.9 and a NetMHC %Rank of 3.98 and was not considered a hit by either approach. The detection of this neoepitope-HLA pair demonstrated that even for abundant epitopes, the chosen cutoffs for the TR-FRET assay and presentation prediction algorithms will produce some level of false-negative results. Lastly, of the 22 neoepitope-HLA pairs detected by untargeted analysis 12 have been previously described in the literature and the remaining 10 neoepitope-HLA pairs were presumed to be novel based on a search of both Tantigen (Olsen et al., 2017), CAatlas (Yi et al., 2021), NEPdb (Xia et l., 2021), and a cursory search of the literature (**Table S4**).

In addition to neoepitope peptides, untargeted analysis enabled the detection of peptides originating from the non-mutation bearing portions of each 25mer neoantigen sequence, as well as junction peptides created by sequential 25mers connected directly or separated by GS linkers within the polyantigen cassette. We detected 27 presented epitopes corresponding to amino acids that flanked but did not contain the mutated residue of the cancer neoantigens. Because these peptides have the same exact sequence as the endogenous version of the protein, we could not determine if these epitopes were derived from the neoantigen construct or the endogenous genome. However, these peptides still provide information about protein processing, including potential proteasome cleavage sites for these common cancer neoantigens. In addition to these epitopes we found 17 and 2 junction peptides from the non-linker and linker constructs, respectively (**Table S4**). We reasoned that the lower number of peptides from the linker construct was due to the fact the linker comprised G and S, amino acids which are not typically anchor residues. Lastly, we identified 5 epitopes from antigen sequences included as controls, as well as 13 epitopes derived from BFP **(Table S4)**, giving us confidence that the neoantigen construct was expressed in each of our monoallelic cell lines.

While untargeted analysis enables identification of thousands of peptides, it is challenged by stochastic sampling and limited detection of peptides presented at low copies per cell. Conversely, targeted MS analysis dedicates the entirety of the instrument duty cycle to the analysis of a small number of peptides, enabling detection of peptides presented at low copies per cell and improving data reproducibility. However, targeted approaches require heavy isotope labeled standard peptides and targeted analysis of all potential neoepitopes for the 47 cancer neoantigens within our polyantigen cassette would require 1,748 peptides to be synthesized. Instead, we used the TR-FRET assay as a preliminary screen and synthesized the 397 peptides with a RZ-score ≥ 5 within the TR-FRET. Due to the complementarity of TR-FRET and NetMHC described above, an additional 81 peptides were also synthesized that had a RZ-score < 5 but a NetMHC %Rank ≤ 2. Including these peptides allowed us to determine if there are neoepitopes predicted by NetMHC that would not have been found within the TR-FRET assay. In total, 479 peptides were divided into allele-specific peptide assays comprising 21 to 88 peptides which were used to quantify neoepitopes across 15 monoallelic cell lines (**Figure 6A-B**).

**Figure 6.**
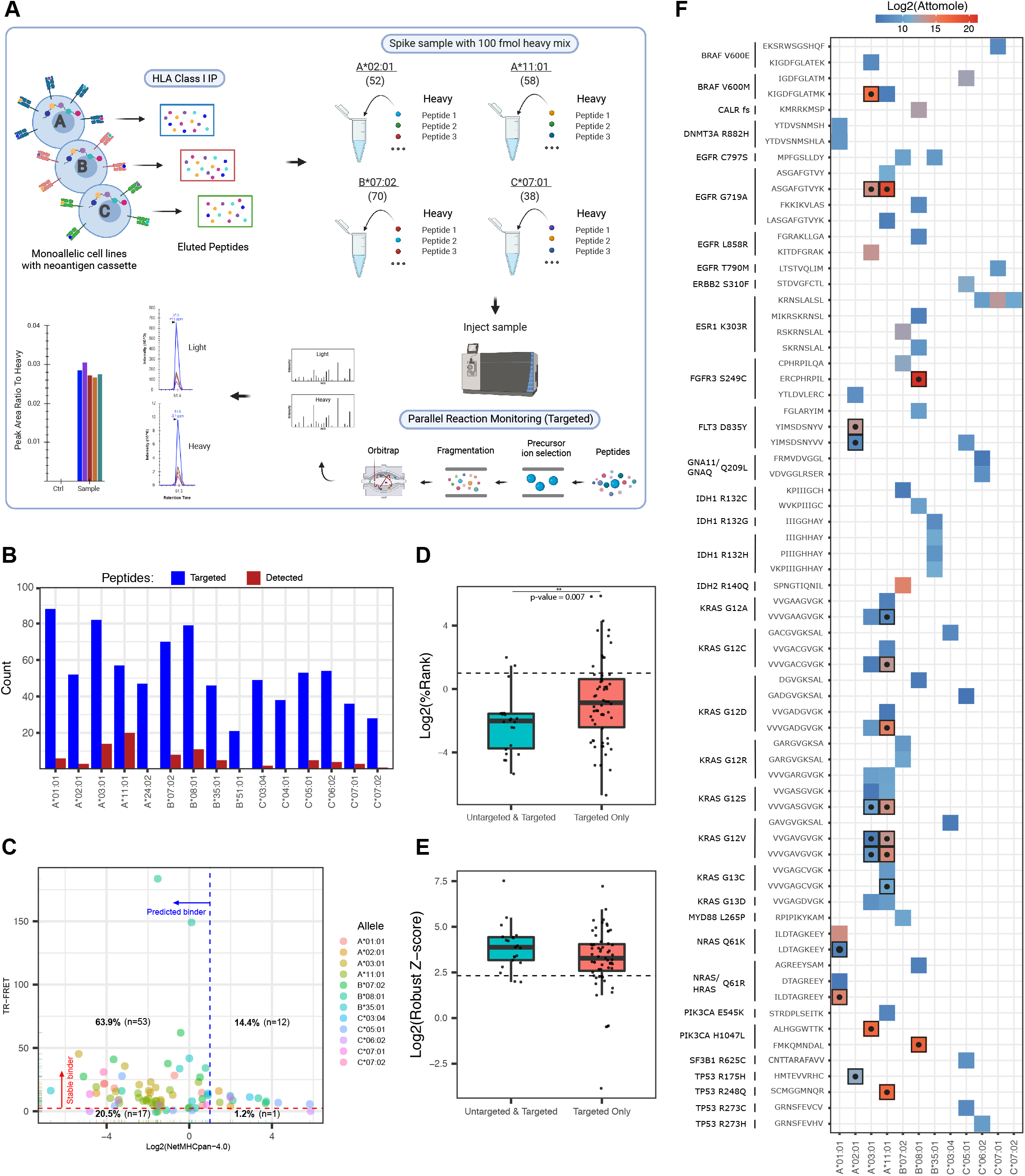
Targeted immunopeptidomic analysis of monoallelic cell lines expressing a polyantigen cassette. **(A)** Targeted immunopeptidomic workflow for the analysis of candidate neoepitopes within monoallelic cell lines expressing a polyantigen cassette. **(B)** The number of targeted (blue) and detected (red) shared cancer neoantigen epitopes within each targeted assay. **(C)** Comparison of TR-FRET Robust ZScore (RZ-Score) and NetMHC percentile rank (%Rank) score for each epitope identified through targeted immunopeptidome analysis. **(D)** NetMHC %Rank scores for neoeptiopes detected in both untargeted and targeted (teal) or targeted analysis alone (red). **(E)** TR-FRET RZ-Scores for neoepitopes detected in both untargeted and targeted (teal) or targeted analysis alone (red). **(F)** Summary of neoepitopes-HLA pairs detected from shared cancer neoantigens. Color represented attomol of neoepitopes detected on column during analysis. Bolded squares with centered dots represent neoepitopes also detected in untargeted analysis.

Targeted MS analysis identified 84 neoepitope-HLA pairs across 12 different alleles and 37 mutations, representing a fourfold improvement when compared to untargeted MS analysis of the same samples (**Figure 6B, Table S4**). Interestingly, 20 of 84 (∼24%) of all neoepitope-HLA pairs were identified within A*11:01. This was likely due to the presence of eight distinct KRAS neoantigen sequences within the polyantigen cassette as 14 of 20 A*11:01 specific neoepitopes mapped to KRAS G12X or G13X neoantigens. A similar pattern was observed in A*03:01 where 9 of 14 neoepitopes belonged to KRAS neoantigens. Following a search of the literature and relevant databases, 24 of the neoepitope-HLA pairs had been described previously, and 60 are novel (**Table S4**).

To understand the relative value of using the TR-FRET assay and NetMHC as a method to select peptides for targeted MS analysis we plotted RZ-score vs. NetMHC %Rank for each of the 83 neoepitope-HLA pairs detected by targeted MS (**Figure 6C**). This analysis revealed that 53 neoepitopes were stable binders by TR-FRET and predicted to be presented by NetMHC. Interestingly, 12 neoepitope-HLA pairs were found as hits in TR-FRET only while 17 neoepitope-HLA pairs were hits in NetMHC alone (**Figure 6C**). These data demonstrated again that both TR-FRET and NetMHC generally agreed on peptides that would be presented - but each method also identified a unique set of potential neoepitope-HLA combinations.

To better understand the binding characteristics of the 62 additional neoepitope-HLA pairs identified by targeted analysis we plotted RZ-scores and NetMHC %Rank scores for peptides observed in both untargeted and targeted analysis and compared them to peptides found only in targeted analysis (**Figure 6D-E**). We found that neoepitope-HLA pairs identified only by targeted analysis alone had a broader range of NetMHC %Rank scores as compared to neoepitopes also detected in untargeted analysis (**Figure 6D**). Furthermore, neoepitope-HLA pairs identified only by targeted analysis had a broader spread of RZ-scores within the TR-FRET assay (**Figure 6E**). These results suggest that targeted analysis can identify neoepitopes that are weaker binders as compared to those identified through untargeted analysis.

Unlike untargeted approaches, targeted MS permits absolute quantification of peptide presentation enabling comparison of presentation across neoepitopes. Here, the measured amount of neoepitope presentation spanned from 60 amol to 2.5 pmol (**Figure 6F**). Unsurprisingly, the peptides detected by untargeted MS were generally detected at higher absolute amounts. For example, both EGFR G719A (ASGAFGTVYK) and FGFR3 S249C (ERCPHRPIL) exhibited the highest absolute abundance - matching the results from the measured intensities from untargeted analysis (**Figure 6F**). However, some epitopes such as PIK3CA H1047L (ALHGGWTTK) exhibited high absolute abundance in targeted analysis, but were not detected in untargeted analysis (**Figure 6F**). When the absolute amounts of neoepitopes detected were compared to either RZ-score or NetMHC %Rank score for each allele no clear correlation could be found (**Supplemental Figure S6**). This suggests that each score could be predictive of whether or not a potential neoepitope was presented, but not the absolute amount presented.

Measurement of absolute levels of neoepitopes also enabled a comparison of presentation from cells containing the linker or no-linker constructs as well as characterization of the reproducibility of measurements across different analysis batches. To understand the impact of linkers within the neoantigen construct we plotted the highest absolute amount of peptide detected for neoepitopes from each cancer neoantigen within monoallelic cell lines containing the linker or no-linker neoantigen constructs (**Supplemental Figure S7-S8**). From these data we concluded that the presence of linkers did not impact presentation in a consistent manner, as presentation of some neoepitopes increased in the presence of linkers while presentation of other neoepitopes decreased (**Supplemental Figure S7-S8**). Lastly, the absolute quantification measurements were collected in 2-3 independent replicates of cell line growth and sample preparation (i.e., HLA-IP and MS analysis) and the absolute measurement of peptide presentation matched well between these replicates (**Supplemental Figure S9**)

### Functional validation of novel tumor associated antigen-HLA-I pairs

A large number of the 84 neoepitope-HLA pairs identified above represent novel candidate neoepitopes. However, presentation of a neoepitope alone does not ensure that it is capable of eliciting a T cell response. To determine whether the identified neoepitopes could be recognized by human T cells, we utilized a modified multiplexed TCR discovery method described by Klinger, et al. (Klinger et al., 2015). Focusing on 2 identified neoeptiope-HLA pairs (Flt3-p.D835Y/HLA- A*02:01,PIK3CA-p.E545K/HLA-A*11:01), briefly, neoepitopes were first allocated to peptide pools in unique combinations before healthy human donor CD8^+^ T cells were isolated and expanded using autologous monocyte-derived dendritic cells, restimulated with the neoepitope peptide pools, sorted for activation marker upregulation, and subjected to TCRβ sequencing. This method was utilized for donors spanning a range of HLA genotypes, enabling the association of TCRs with a variety of peptide-HLA pairs. However, due to the multiallelic nature of donor cells the HLA restriction of identified neoepitopes was not initially disambiguated among the 3-6 donor HLA alleles.

For neoepitopes that elicited a T cell response, associated TCRβ and TCRα sequences were determined using a parallel multiplexed assay (Howie et al., 2015) that enabled construction of paired TCR expression vectors and the selection of candidate neoepitope-specific TCRs. The specificity and potential efficacy of each TCR was then assessed through cellular assays. TCR-encoding *in vitro* transcribed RNA (ivtRNA) was introduced via electroporation into primary human T cells which were then incubated with either an increasing concentration of the candidate neoepitope in the presence of target cells expressing the predicted HLA allele or monoallelic K562 cells expressing both the predicted HLA allele and neoantigen of interest. These two approaches enabled characterization of TCR potency through activity of an exogenously loaded target cell and the potential for the neoepitope to elicit a T cell response when it is expressed, processed, and presented in a cellular context.

We found dose-dependent upregulation of CD137 after 12 hour co-culture of primary human CD8+ T cells transfected with predicted Flt3-p.D835Y/HLA-A*02:01-specific TCRs, in response to T2 cells (which express low levels of HLA-A*02:01) incubated with the indicated concentrations of exogenously delivered YIMSDSNYV peptide (**Figure 7A**). Furthermore, these T cells were activated by and specifically killed monoallelic A*02:01 K562 cells expressing a mutant FLT3- p.D835Y transgene, but were not activated by and did not kill monoallelic A*02:01 K562 cells expressing a wild type FLT3 transgene (**Figure 7B-C**). Interestingly, these TCRs appear to be exquisitely specific for the mutant neoepitope, an important characteristic because a similar non-mutant epitope IMSDSNYVV was identified by untargeted analysis in HLA-A*02:01 monoallelic cells. These data suggest a potential utility for these TCRs as a modality to address Flt3-p.D835Y expressing malignancies.

**Figure 7.**
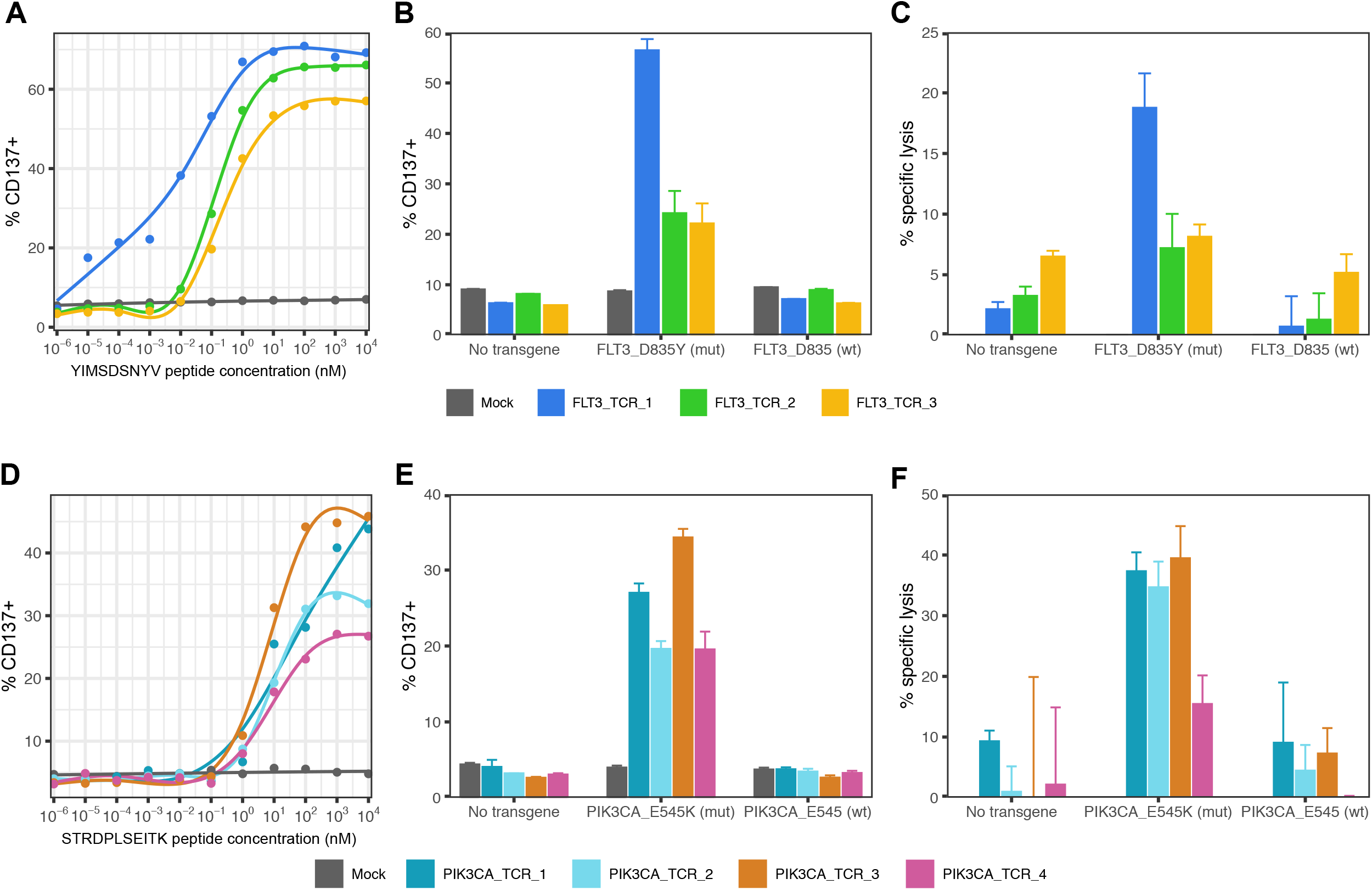
Discovery of neoepitope specific TCRs demonstrates immunogenic potential of discovered neoepitope-HLA pairs. Human CD8+ T cells were transfected with either FLT3- p.D835Y-specific or PIK3CA-p.E545K-specific TCR RNA. **(A)** CD137 expression was assessed after FLT3-p.D835Y-specific TCRs were co-cultured overnight with YIMSDSNYV peptide-pulsed- HLA-A*02:01+ T2 cells. CD137 expression and specific cell lysis were determined following co- culture with HLA-A02:01+ K562s transfected with no RNA, transgene containing mutant FLT3- p.D835Y or transgene containing FLT3-D835 wildtype sequence. **(B)** CD137 expression was assessed after PIK3CA-p.E454K-specific TCRs were co-cultured overnight with STRDPLSEITK peptide-pulsed HLA-A*11:01+ K562 cells. CD137 expression and specific cell lysis were determined following co-culture with HLA-A11:01+ K562 cells transfected with no RNA, transgene containing mutant PIK3CA-E545K or transgene containing PIK3CA-E545 wildtype sequence.

As a second proof of concept, T cells were transfected with predicted PIK3CA-p.E545K/HLA- A*11:01 TCRs and mixed with monoallelic HLA-A*11:01 expressing K562 cells incubated with an increasing concentration of the predicted neoepitope, STRDPLSEITK (**Figure 7D**). Here, TCR transfected T cells demonstrated dose-dependent activation as measured by CD137 expression. Furthermore, these T cells demonstrated higher levels of activation and cell killing when mixed with monoallelic A*11:01 K562 cells expressing a PIK3CA-p.E545K transgene as compared to cells that expressed a wild-type PIK3CA transgene (**Figure 7E-F**). Mutations that introduce anchor residues are thought to have a high immunogenic potential because the immune system has not built tolerance to a similar WT epitope. For PIK3CA-p.E545K/HLA-A*11:01 the E→K mutation introduces an anchor residue within the context of HLA-A*11:01 and the wild type STRDPLSEITE epitope was not detected in untargeted MS analyses of A*11:01 monoallelic cells. While lack of detection in MS analysis does not demonstrate absence, the WT epitope was also not predicted to bind HLA-A*11:01 by NetMHC (12.8). Taken together these data provide a clear mechanism for specificity of PIK3CA-p.E545K TCRs for recognition of mutant PIK3CA as compared to wild-type and support these TCRs as potential therapeutic candidates.

## Discussion

To date, most neoepitope discovery efforts have focused on a limited number of neoantigens, HLA alleles, or both in the search for immunogenic tumor-associated peptides. While a recent report expanded the number of neoantigens and HLA alleles studied at once, these neoepitopes were derived from mutations of the same gene - KRAS (Choi et al., 2021). Here, we developed a multiplexed platform that integrated a high throughput binding assay, computational neoepitope binding prediction, complex cellular engineering of monoallelic cell lines, and targeted mass spectrometry to identify unique tumor-associated neoepitopes that can be presented in the context of specific HLA-I alleles and function as potential targets for neoantigen based cancer immunotherapy. As demonstrated, our workflow leverages the combination of a high throughput biochemical neoepitope-HLA binding assay together with a computational neoepitope-HLA binding algorithm to enable systematic screening of all potential neoepitopes across 47 shared cancer neoantigens and 15 common HLA alleles. The combined analyses yielded a short list of 783 neoepitope-HLA combinations (out of 24,149 total combinations surveyed) that were identified as stable binders and potential candidates for cell surface HLA-I presentation. Separately, a custom engineered monoallelic cell line panel containing a polyantigen cassette encoding all 47 neoantigens was created to increase the breadth of putative neoepitope-HLA combinations presented on the cell surface. Using a combination of untargeted and targeted mass spectrometry analysis, we identified 80 unique neoepitope-HLA pairs derived from 36 shared cancer neoantigens across 12 of 15 surveyed HLA alleles. To validate the immunogenicity of these unique neoepitope-HLA pairs, and their potential as therapeutic targets, we selected two example combinations (Flt3-p.D835Y/HLA-A*02:01 and PIK3CA-p.E545K/HLA-A*11:01) and used cell-based assays to evaluate a cohort of neoantigen specific TCRs identified in a separate MIRA workflow. Not only were T cells activated in the presence of corresponding TCR-antigen/HLA-I combinations, but several TCRs exhibited mutant peptide-selectivity.

Beyond the analysis we provide here, the TR-FRET and MS data represent a valuable resource for future studies of neoepitope presentation. For example, the TR-FRET data could potentially be used as training or benchmarking data for more advanced computational algorithms that predict neoepitope-HLA complex formation. Additionally, we provide raw data for untargeted and targeted MS analysis - enabling future re-analysis with more advanced search algorithms (Vizcaíno et al., 2020), peptide false discovery rate determination (Wilhelm et al., 2021), or specific workflows that detect rare events within the antigen presentation pathway (e.g., spliced peptides (Faridi et al., 2021). In addition to the data generated in this study, the monoallelic cell lines expressing the polyantigen cassette also represent an ideal system for future studies characterizing the processing and presentation of clinically actionable shared cancer neoantigens.

Collectively, the workflow we describe herein provides the most comprehensive analyses of the neoepitope landscape performed to date and offers critical insight into therapeutic targets for neoepitope based cancer immunotherapies targeting shared neoantigens. One striking finding from this analysis was the relatively few (84 total out of 24,149 initially-screened neoepitope-HLA combinations, or 0.35%) shared neoepitope-HLA pairs presented and, consequently, made available for therapeutic development. Limited presentation of shared cancer neoepitopes could be driven by the diversity of HLA-I peptide binding as neoepitopes for 18 of 37 cancer neoantigens were detected in the context of only one HLA-I allele. For the 19 cancer neoantigens that presented epitopes across multiple HLA-I alleles, 7 were KRAS G12X or G13X mutations. Due to the low incidence of presentation across multiple prevalent HLA-alleles, a broadened use of this platform and additional neoepitope-HLA discovery efforts will be needed to identify the patient populations most likely to benefit from shared neoantigen-specific immunotherapies. For example, the KRAS G12D mutation was reported to be the most frequent in CRC with a frequency of 14.9% (Araujo et al., 2021). Based on our analysis, the most prevalent HLA allele presenting a KRAS G12D neoepitope was A*03:01, which on average constitutes an allele frequency of about 14% across patients of Caucasian and European descent. Therefore, for this highly common shared neoantigen within this indication, the maximum patient coverage for European and Caucasion demographic is only ∼2% (mutation frequency X allele frequency). This number decreases further when considering other demographics where the prevalence of HLA-A*03:01 is even lower, such as African American (1.1%), Chinese (0.2%), Hispanic (1.0%), and Southeast Asia (0.75%). Although there is growing evidence that neoantigen-based therapeutics are highly effective for the treatment of cancer, these collective findings suggest that targeting shared neoantigens will remain challenging.

## Limitations of the study

Although this analysis was broad, it is possible the neoepitope-HLA combinations were missed from either the biochemical assay and/or the prediction algorithms and were not included in the targeted mass spectrometry analysis. Also, presentation was measured in an engineered cell line overexpressing a polyantigen cassette encoding ∼25mer amino acid fragments spanning the mutated amino acid and it is possible that processing and presentation in this format could be altered compared to full length antigen in a tumor cell, resulting in missed neoepitopes. To address this potential limitation we evaluated neoepitope presentation from cell lines engineered with the full length neoantigen for KRAS G12C, G12D and G12V and observed the same neoepitopes presented (**Figure 4**). However, we did not validate this across all 47 neoantigens described in this manuscript. Despite these caveats, this is one of the most systematic analyses of the neoepitope landscape across the most relevant shared neoantigens and the collective findings yield valuable insight into druggable neoepitope targets for cancer immunotherapy.

## Supporting information

Supplemental Sequences

## Figure Legends

**Supplemental Figure 1.**
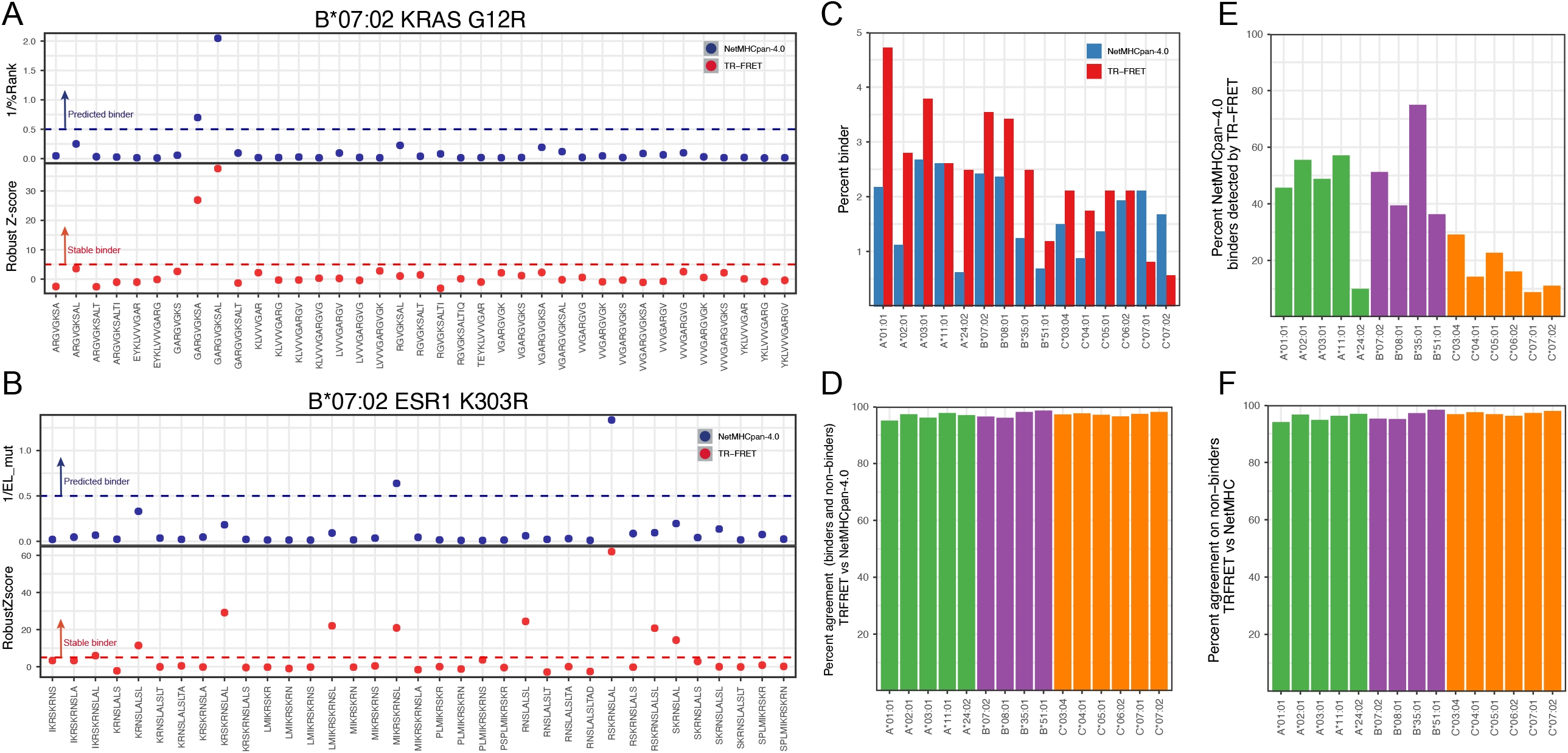
Comparison of TR-FRET and NetMHC analysis. **(A)** TR-FRET Robust Z-score (red dots) and 1/NetMHC Percentile Rank (blue dots) of all neoepitope-HLA combinations for the B*07:02 allele and KRAS G12R neoantigen. The blue and red lines represent the cutoff of stable binders for NetMHC and TR-FRET analysis, respectively. **(B)** TR-FRET Robust Z-score (red dots) and 1/NetMHC Percentile Rank (blue dots) of all neoepitope-MHCI combinations for the B*07:02 allele and ESR1 K303R neoantigen. The blue and red lines represent the cutoff of stable binders for NetMHC and TR-FRET analysis, respectively. **(C)** Percent of neoepitope-MHCI combinations that were determined to be stable binders by TR-FRET (red bars) and NetMHC (blue bars) analysis for each individual allele. **(D)** Percent neoepitopes-HLA pairs found to be binders and non-binders by both NetMHC and TR-FRET across the individual alleles. **(E)** Percent neoepitopes-HLA pairs found to be binders by both NetMHC and TR-FRET across the individual alleles. **(F)** Percent neoepitopes-HLA pairs found to be non-binders by both NetMHC and TR-FRET across the individual alleles

**Supplementary Figure 2.**
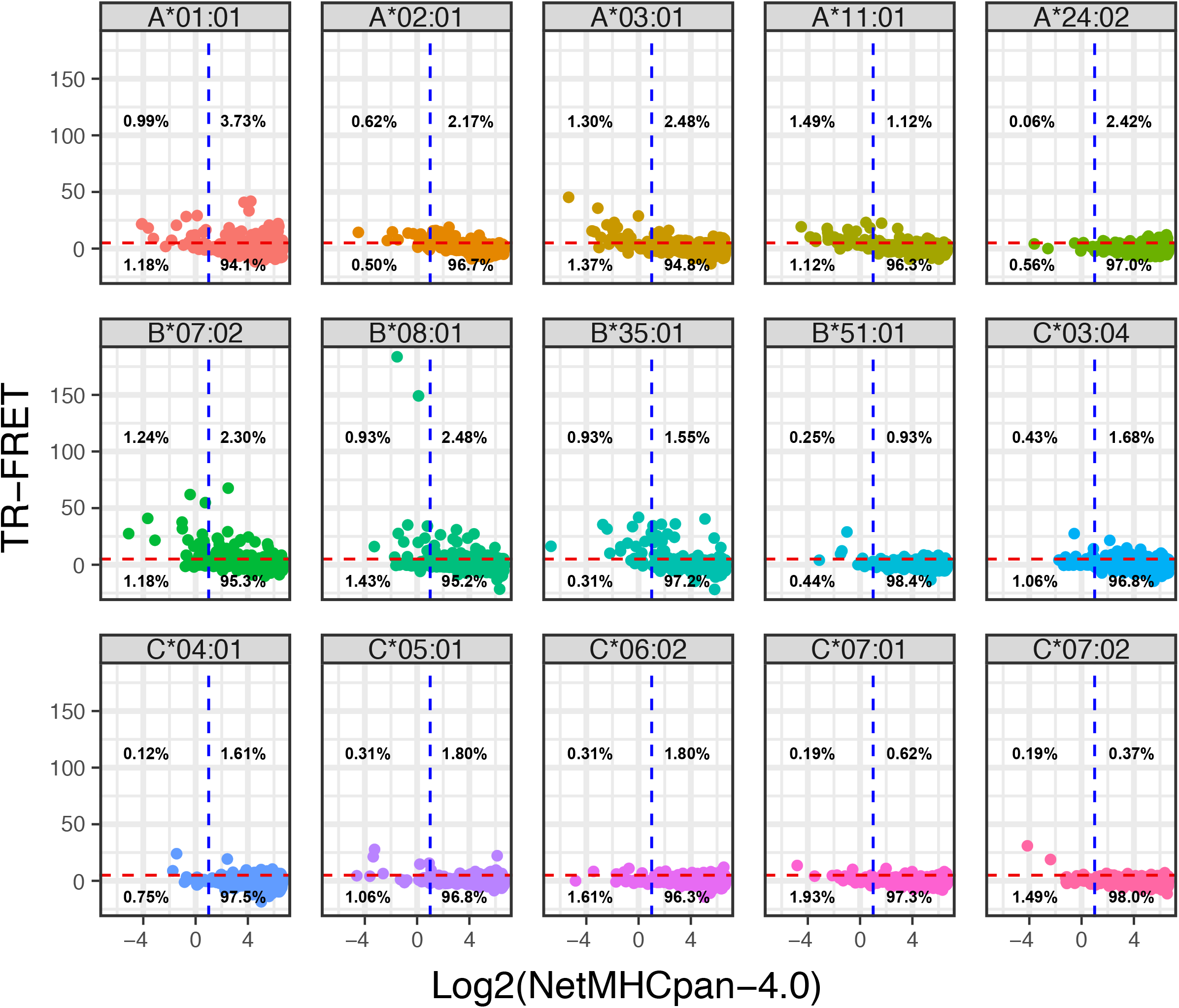
Scatter plot of TR-FRET Robust Z-score and Log2 NetMHC percentile rank for all 15 alleles across all neoepitopes evaluated. The dashed red line represents the cutoff for stable binders as measured by TR-FRET, where values higher than the red line are considered stable binder. The dashed blue line represents the cutoff for binders based on NetMHC analysis, where values lower than the blue line are considered binders.

**Supplementary Figure 3.**
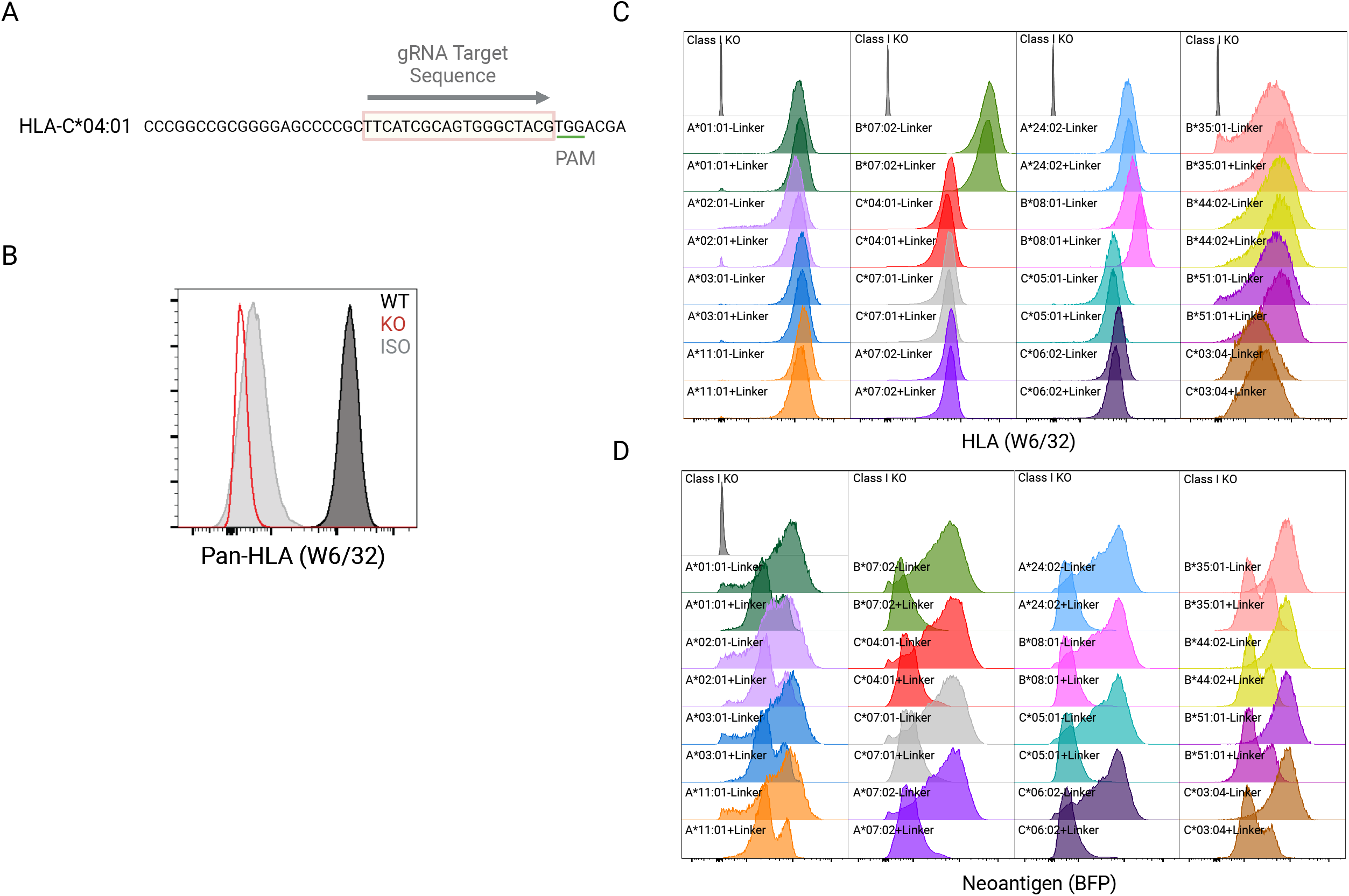
Further analysis of polyantigen expressing HLA class I monoallelic cell lines. **(A)** guide RNA sequence and genomic context for CRISPR/Cas9 mediated gene disruption of the HLA-C locus in HMy2.C1R cells. **(B)** Flow cytometric detection of pan-HLA-I expression in wild-type (WT) or HMy2.C1R HLA I-knockout (KO) cells using a pan-HLA-I detection antibody W6/32 or isotype (ISO) control. **(C)** Flow cytometric detection of pan-HLA-I expression of indicated monoallelic cell lines. **(D)** Flow cytometric detection of a transcriptionally linked TagBFP2 reporter gene in indicated polyantigen expressing monoallelic cell lines.

**Supplementary Figure 4.**
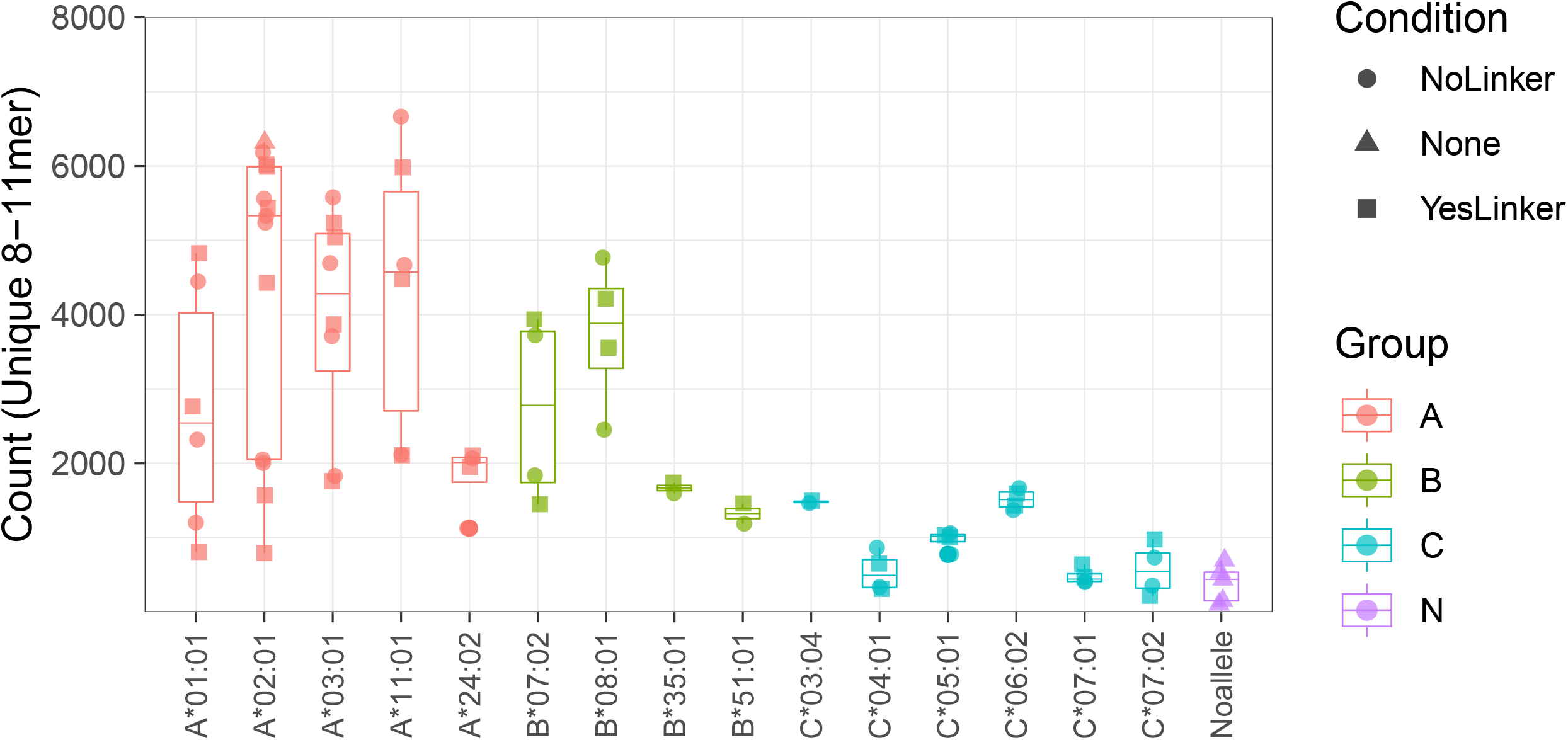
Summary of peptides identified in untargeted immunopeptidomics. Number of unique peptides (8-11 mer) identified in untargeted proteomics analysis stratified by allele and linker status of neoantigen construct. Each dot represents a measurement of a cell pellet. Boxes represent the interquartile range and line represents the median.

**Supplementary Figure 5.**
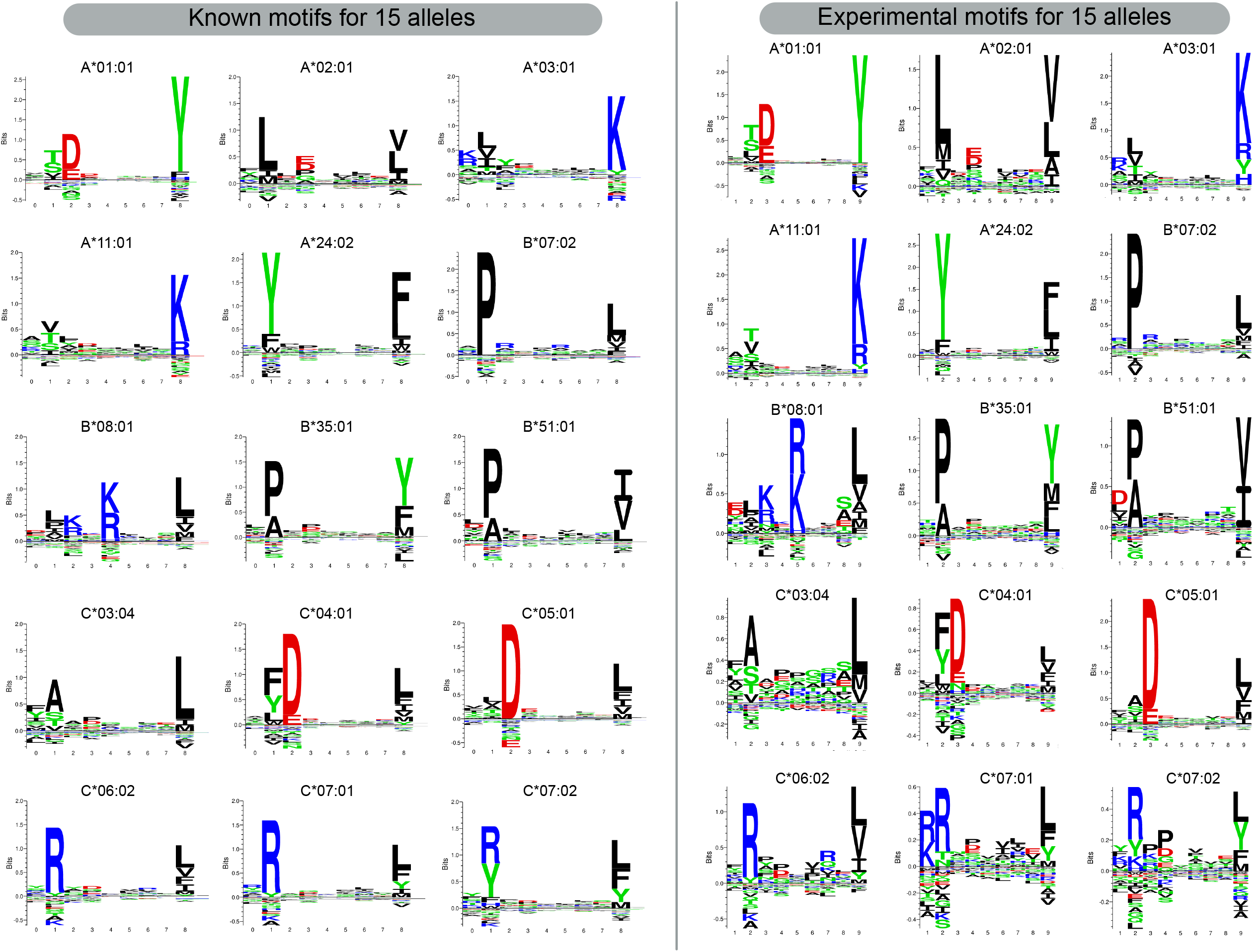
Motif analysis of untargeted proteomic analysis. Motifs were generated with GibbsCluster 2.0 with two bins allowing for one bin of the dominant motif and a second bin for non-specific peptides.

**Supplementary Figure 6.**
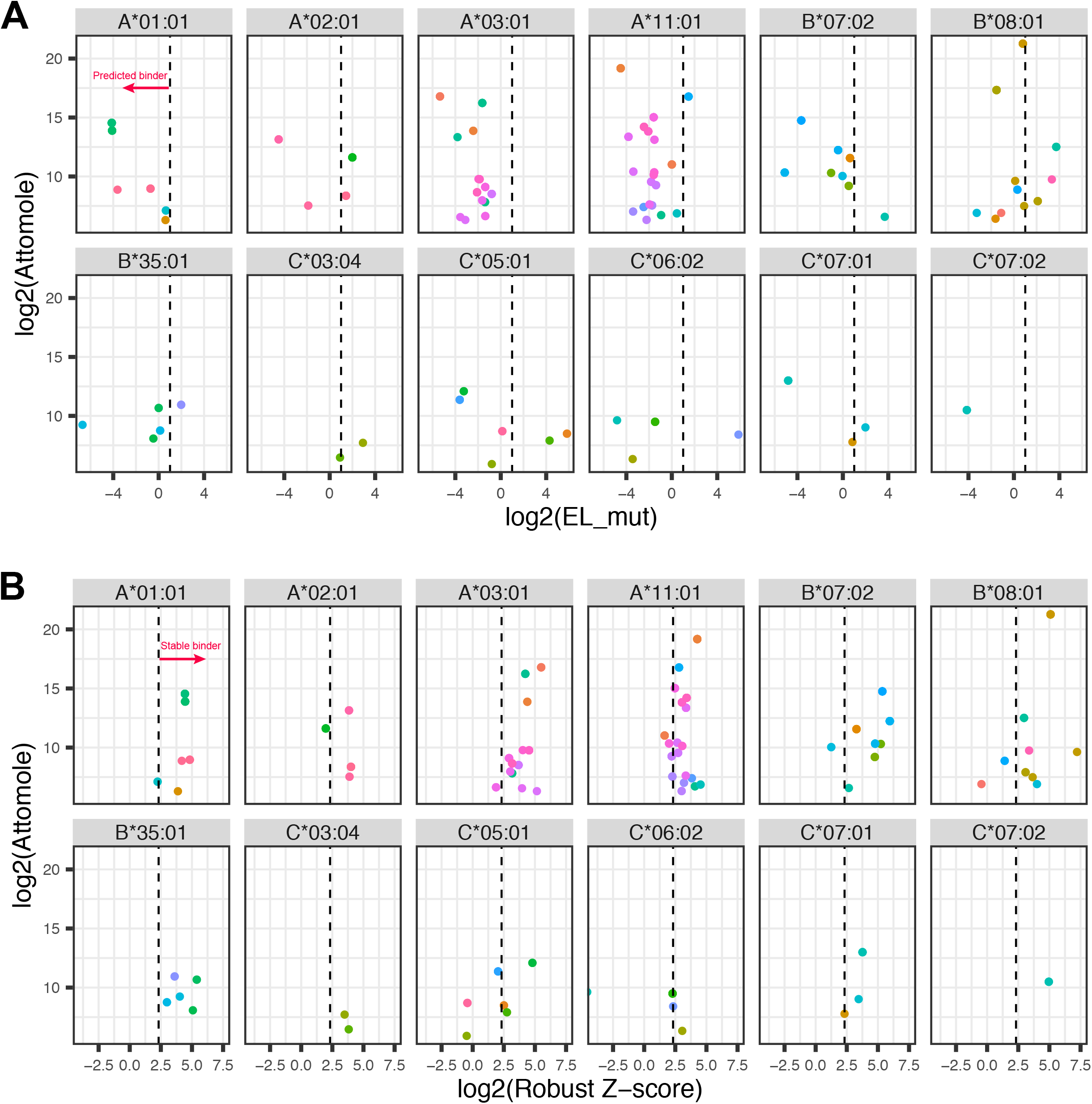
Comparison of absolute peptide presentation and NetMHC predicted binding score for TR-FRET Robust Z-score. Each dot represents a neoeptiope-HLA pair detected within the targeted proteomic analysis. If multiple neoepitope-HLA pairs were detected, the attomole value is the maximum value for that peptide-MHC pair across all analyses.

**Supplementary Figure 7.**
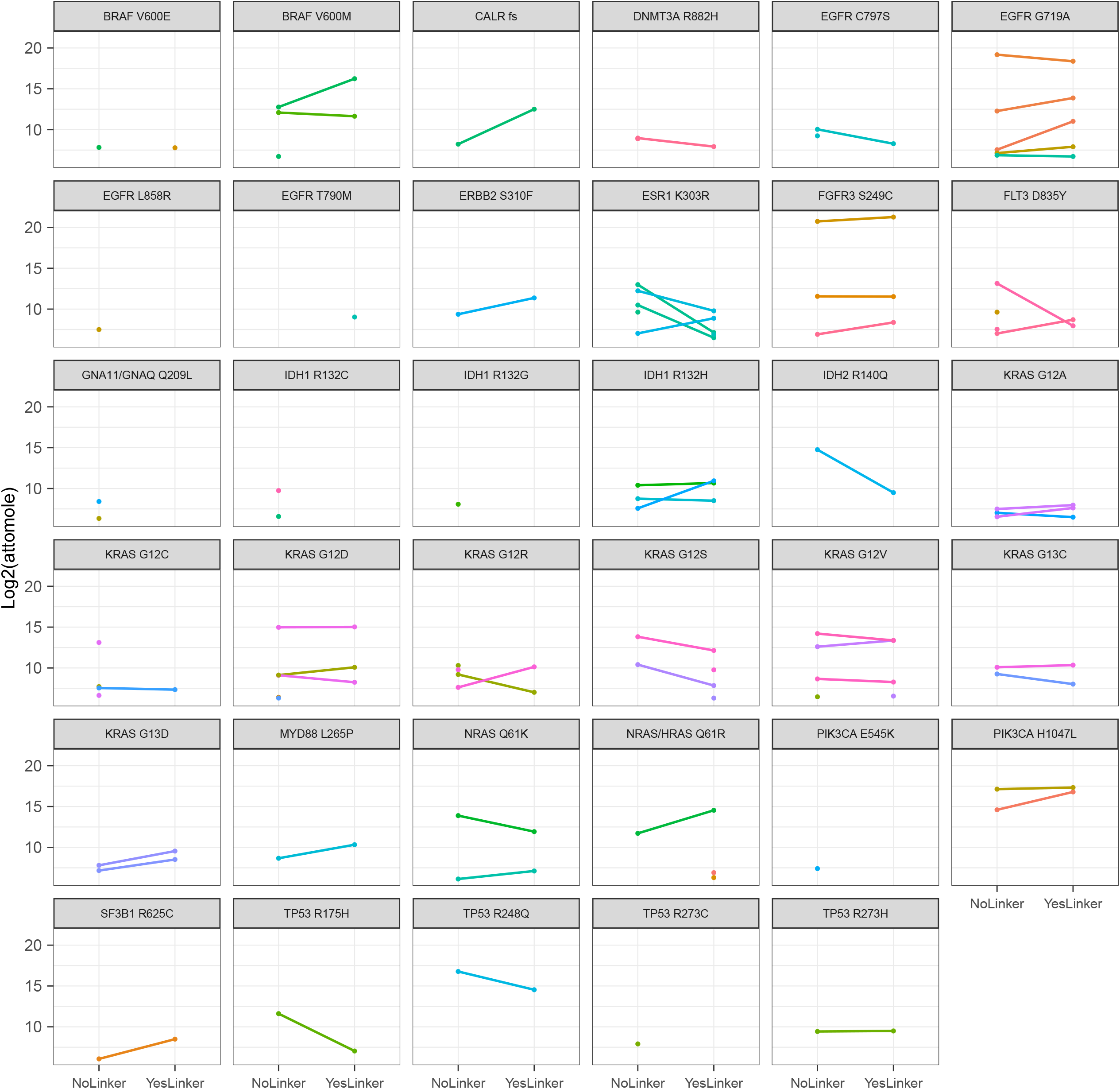
Comparison of epitope presentation for cell lines containing linker and no linker construct. Each line represents a specific epitope and the slope of the line demonstrates if expression was lower, higher, or similar between monoallelic cell lines containing linker and no-linker polyantigen cassettes.

**Supplementary Figure 8.**
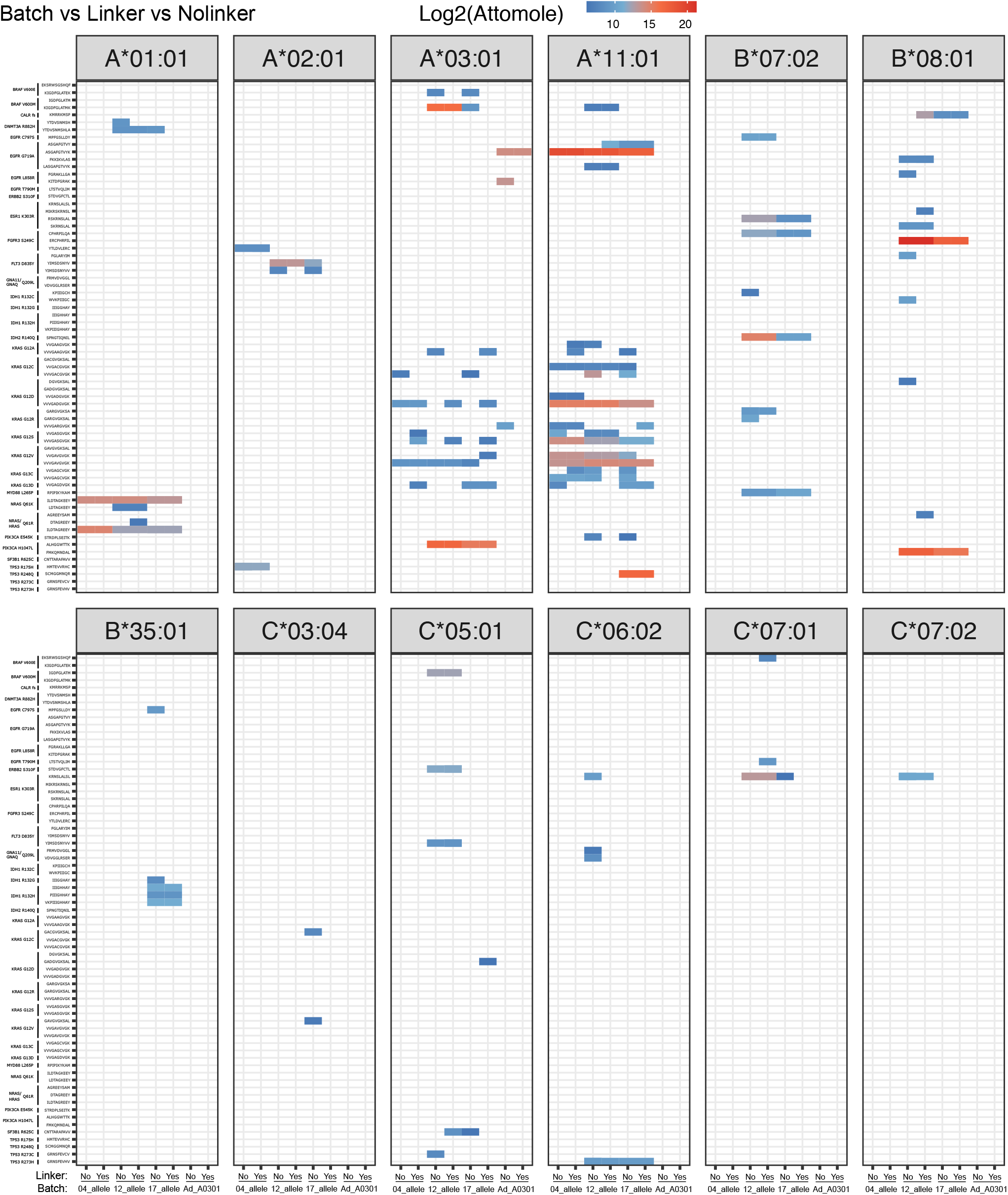
Analysis of epitope presentation across analysis batches with consideration of presence of linker in construct. Each column represents a specific analysis batch for a cell line expressing a linker or no-linker polyantigen cassettes. Color represents the absolute abundance measured by the attomole amount detected on column for each neoepitope.

**Supplementary Figure 9.**
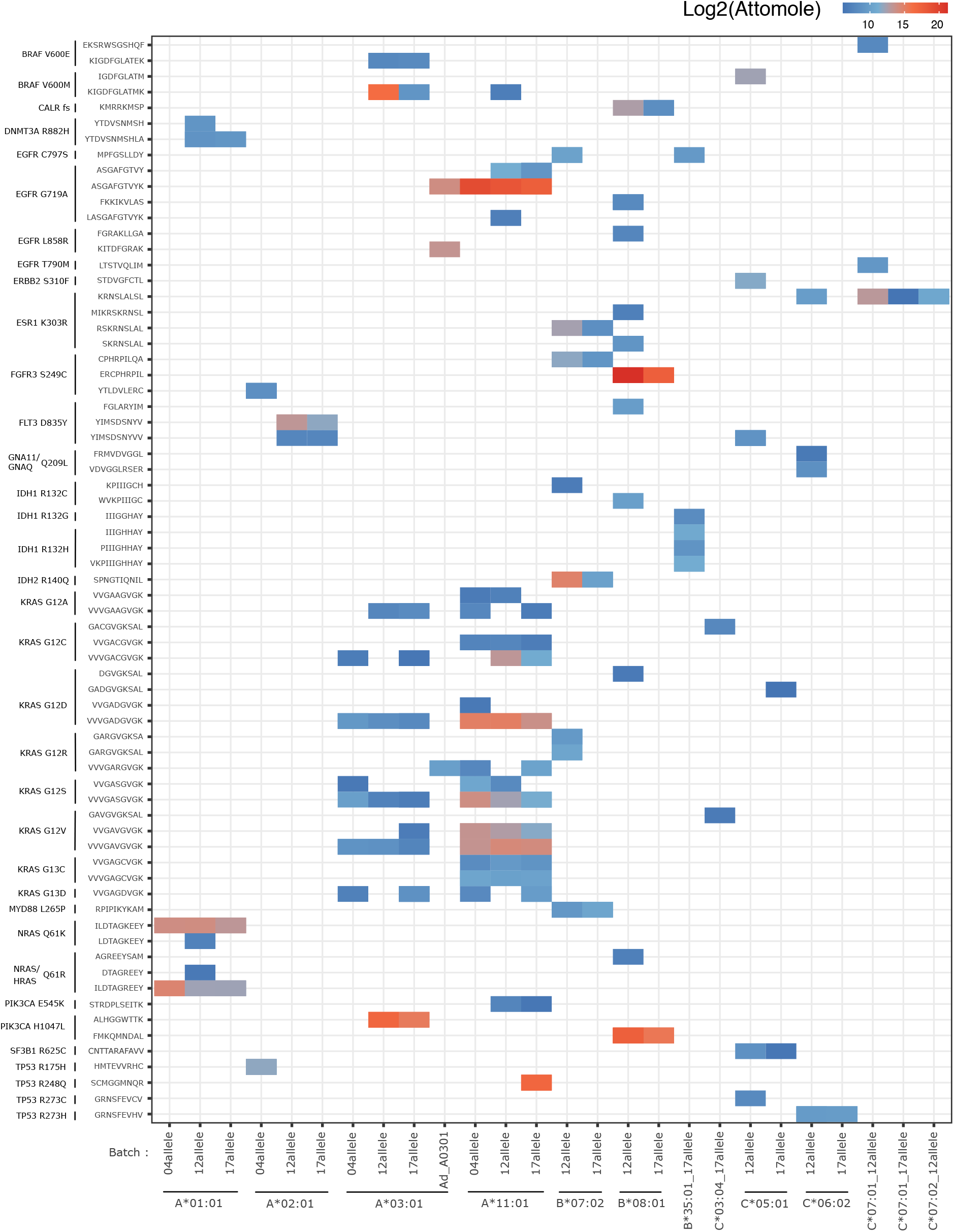
Comparison of epitope presentation across analysis batches. For each allele, detection of epitopes displayed for each batch that contained analysis of monoallelic cell lines expressing that particular allele. Color represents the absolute abundance measured by the attomole amount detected on column for each neoepitope.

## STAR★ Methods Key resources table

## Resource availability

### Lead contact

Further information and requests for resources and reagents should be directed to and will be fulfilled by the lead contact, Craig Blanchette

### Materials availability

- Plasmids generated in this study are property of Genentech but can be made available under an MTA
- Cell lines generated in this study are property of Genentech but can be made available under an MTA
- MHCI complexes generated in this study are property of Genentech but can be made available under an MTA

### Data and code availability

- All mass spec data has been deposited in the MASSIVE repository (Wang et al., 2018) and are publicly available as of the date of publication. Accession numbers are listed in the key resources table.
- Any additional information required to re analyze the data reported in this paper is available for the lead contact upon request.

## Experimental model and subject details

### Engineering of monoallelic polyantigen cassette expressing HMy2.CR1 cell lines

An HLA class-I (HLA-I) null cell population was generated by CRISPR/Cas9 mediated gene disruption of the endogenous HLA-C locus in HMy2.C1R cells (Cell engineering Class I KO fig A). Wildtype HMy2.C1R cells were electroporated with Cas9/RNP (Invitrogen) containing a HLA-C-specific sgRNA (Synthego), sequence:TTCATCGCAGTGGGCTACG (S3A) using an Amaxa V system (program D-023). Following an expansion period, cells were stained with anti-pan-HLA (W6/32) and antigen-negative cells were enriched via FACS (**Supplemental Figure S3B**).

HLA-I null HMy2.C1R cells were stably engineered with a piggyBac neoantigen expression plasmid system designed to co-express 47 shared cancer neoantigens and 7 HLA-A*02:01 control antigens **(Supplemental Sequences)**. Briefly, neoantigen segments (∼25 amino acids each) were concatenated and converted to codon-optimized DNA segments (IDT), with or without a flexible linker separating most neoantigen sequences. The polyantigen cassettes were synthesized and cloned into a piggyBac transposon plasmid downstream of a constitutive human EF1a and transcriptionally linked to an IRES-TagBFP2 reporter element. A separate hPGK promoter-driven puromycin resistance gene was included on the same vector for selection purposes. The polyantigen expression plasmid was co-electroporated with pBO (piggyBac transposase, Hera Biolabs) using a NEON system (Invitrogen) and 100 ul kit (buffer R, 1230 V, 20 ms, 3 pulses). Polyantigen expressing cells were selected by culture in 1ug/ml puromycin (Gibco) and further purified by FACS enrichment of the TagBFP2-positive population.

Unique HLA-I allele ORFs, each with a distinct 19 bp DNA barcode, were cloned downstream of the human EF1a promoter (Genscript) in a custom-modified pLenti6.3 backbone (ThermoFisher). Lentivirus was generated by lipofectamine 2000 (Invitrogen)-mediated co-transfection of HEK293T cells with individual lenti HLA expression constructs and packaging plasmids. 72 hours post-transfection, viral supernatant was harvested, filtered through a 0.45 uM filter, and concentrated via LentiX concentrator reagent (Takara) following the manufacturers recommended protocol. Linker or no-linker polyantigen expressing HLA-I-null HMy2.C1R cells were transduced with HLA expression vectors via spin infection (800xg for 30 min at room temperature with 8 ug/ml polybrene). Transgenic HLA-expressing cells subsequently were purified via magnetic bead-based enrichment (biotin-W6/32 Biolegend /SA-MACS). HLA allele identification was confirmed via barcode sequencing (Amplicon Primers: Fwd- CTCCCAGAGCCACCGTTACAC, Rev- GACTTAACGCGTCCTGGTTGC; sequencing primer: CTGGTTGCAGGCGTTTAGCGT) and uniform expression of both the HLA allele and polyantigen cassette were confirmed via flow cytometry (**Figure 3C**; **Supplemental Figure S3 B-C**) prior to analysis via mass spectrometry.

For studies evaluating neoantigen presentation in the context of full-length neoantigen containing proteins, A*11:01 monoallelic cell lines were stably engineered with an all-in-one doxycycline inducible piggyBac vector expressing wild-type or mutant alleles (G12D, G12C, G12V) of human KRAS **(Supplemental Sequences)**. The KRAS allele of interest and an IRES linked mCherry reporter was driven by a dox responsive TRE3G promoter. A puromycin resistance gene and the Tet-on3G element was encoded on the same vector downstream of a constitutive hPGK promoter. Post-puro selection and expansion, KRAS expression was induced by treating cells with 1ug/ml dox for 5 days prior to subsequent analysis.

## Methods

### Clinico-Genomics Analysis

Prevalence data for common cancer mutations (SNVs and indels) were obtained from the Cancer Hotspots database (http://cancerhotspots.org, (Chang et al., 2017)) and cross-referenced with TCGA data obtained from the cBioPortal for Cancer Genomics (http://cbioportal.org). Prevalence data for common HLA alleles from the general population were obtained from the Allele Frequency Net Database (http://allelefrequencies.net), as well as from HLA typing of >8,000 TCGA cases (A. Horowitz, personal communication). From these data sets, the 47 most common cancer mutations were determined based on prevalence per cancer type, and the 47 most common HLA-I alleles were determined. Additional ranking of these mutations was performed that considered the overall prevalence of each cancer type, and whether a neoantigen-specific therapy could be readily developed in a clinical setting.

### Predicted neoepitope landscape analysis

After translating mutations to peptide sequences, neoepitope-HLA binding predictions were generated using NetMHCpan-4.0 (Jurtz et al., 2017) on all combinations of 8, 9, 10, and 11-mer peptides derived from the 47 cancer neoantigens combined with 15 prevalent HLA alleles. Both “binding affinity” (BA) and “eluted ligand” EL predictions were obtained, which were then used for downstream analysis. Predicted neoepitopes were defined as neoepitope-HLA combinations with mutant EL percentile rank < 2.

### Protein expression and purification

Recombinant HLA and β2M were over expressed in E. coli, purified from inclusion bodies, and stored in denaturing buffer (6M Guanidine HCl, 25 mM Tris pH 8) at -80°C as described previously (Darwish et al., 2021). Briefly, β2M and HLA biomass pellets were resuspended in lysis buffer (PBS+1% Triton X-114) at 5 mL/g and homogenized twice in a microfluidizer at 1000 bar. The homogenized suspension was spun at 30000 g for 20 min in an ultracentrifuge. The pellets were collected and washed with 500 ml of 0.5% Triton X-114 in PBS. Collected samples were then centrifuged at 30000 g for 20 min. The pellet was collected again and washed as described above. The purified inclusion bodies were dissolved in denaturing buffer (20 mM MES, pH 6.0, 6 M Guanidine) at a concentration of 10 ml/g and stirred at 4°C overnight. The dissolved pellet was centrifuged at 40000 g for 60 min and the supernatant was collected and filtered through a 0.22 mm filter. The concentration was determined by UV-vis at 280 nm using the protein’s extinction coefficient. Samples were then snap-frozen and stored at -80°C prior to generation of complexes.

### HLA-I peptide refold, biotinylation, and purification

Conditional HLA-I complexes were generated in a 5L refold reactions in refold buffer (100 mM Tris, pH 8.0, 400 mM L-Arginine, 2 mM EDTA) as described previously (Darwish et al., 2021). Briefly, the refold reaction consisted of the conditional HLA-I ligand peptide containing a nonnatural UV cleavable amino acid (0.01mM), oxidized and reduced glutathione (0.5mM and 4.0mM, respectively), recombinant HLA (0.03mg/ml) and β2M (0.01mg/ml). The refold mixture was stirred for 3-5 days at 4°C, filtered through a 0.22 μm filter, and concentrated and buffer exchanged by tangential flow filtration (TFF) (Millipore P2C010C01) into 25 mM Tris pH 7.5. The concentrated and refolded HLA-I complex was then biotinylated through the addition of BirA (1:50 (wt:wt) enzyme:HLA-I), 100 mM ATP and 10X reaction buffer (100 mM MgOAc, 0.5 mM biotin) for an incubation period of 2 hr at room temp. The sample was dialyzed and analyzed by LC/MS to quantify biotinylation. The biotinylated HLA-I complex was purified by anion exchange chromatography using a 1ml HiTrap Q HP column on an AKTA Avant FPLC. The column was equilibrated with 10 column volumes (CV) of 25 mM Tris-HCl pH 7.5 at a flow rate of 5 ml/min. The refolded peptide-HLA-I sample was loaded on the column at a 5 ml/min flow rate and eluted using 0-60% 2.5 mM TrisHCl, pH 7.5, 1 M NaCl gradient over 30 CV. Fractions across the eluted peak were run on SDS-PAGE, and fractions containing both β2M and HLA bands were pooled. Pooled fractions were buffer-exchanged into storage buffer (25 mM Tris HCl, pH 8.0, 150 mM NaCl). Protein concentration was determined by UV absorbance at 280 nm, and samples were snap-frozen and stored at -80°C.

### Peptides synthesis for *in vitro* binding assay

Peptides for the binding screen were synthesized by JPT Peptide Technologies GmbH (Germany) and purified to >70% purity by HPLC. Peptides were dissolved in ethylene glycol (Sigma) at 1 mg/mL and stored at -80°C in Matrix 1.0 mL 2D screw cap tubes (Thermo Scientific). UV-cleavable peptides were synthesized with 3-amino-3-(2-nitrophenyl)propionic acid by Elim Biopharm and purified by HPLC to >70% purity.

### Automated high throughput neoepitope exchange

Peptides were diluted to 10 uM in 25 mM TRIS pH 8.0, 150 mM NaCl, 4 mM EDTA, 4.35% ethylene glycol, in 96 deep well plates (VWR) using a Biomek i5 automated liquid handler (Beckman Coulter). The peptide-buffer mixtures were dispensed and reformatted into 384 well plates (Labcyte) at a volume of 47.5 μl per well, resulting in identical plates of up to 352 unique Neoepitopes for screening against each of the 15 HLA alleles. The first two columns of the plate were reserved for controls. HLA A*02:01 with and without exchange peptide were included on each plate as positive and negative controls for exchange, respectively. The well characterized HLA A*02:01 specific viral epitope, CMV pp65 peptide (NLVPMVATV, Elim Biopharm), was plated in quadruplicate, as a positive control for peptide exchange. Negative controls for exchange included wells to which no peptide was added, and instead received ethylene glycol only during the peptide dilution step. Negative control wells for the HLA allele being screened were plated in octuplicate.

Using a Mantis liquid handler (Formulatrix), 2.5 μl of 0.1 mg/ml UV peptide-HLA complexes were added to each well with one HLA allele screened for binding per plate. Positive control wells received HLA A*02:01, and negative control wells received either HLA A*02:01 or the HLA allele specific to the plate. The resultant peptide exchange reaction mixtures contained 10 uM peptide, 0.1 uM UV-HLA complex, and 5% ethylene glycol v/v.

The peptide exchange protocol was adapted from a method previously described (17406393) by decreasing the UV exposure time and adding an incubation step after UV exposure. Plates containing the peptide exchange reaction mixtures were incubated under UV lamps (UVP 3UV Lamp, Analytik Jena) for 25 min using one lamp per plate. Plates were then sealed and incubated for 18 hours at room temperature.

### TR-FRET assay

The homogenous TR-FRET assay was carried out in a MAKO 1536-well white solid bottom plates(Aurora Microplates, Whitefish, MT). The total assay volume was 4 μL per well, including 2 μL diluted samples and 2 μL of reagent mix. In brief, 1.8 μL per well of assay diluent (PBS, 0.5% BSA + 0.05% Tween20 +10PPM Proclin, Genentech, Inc) was added to the 1536-well destination plate by a Multidrop™ Combi nL Dispenser (Thermo Fisher Scientific, Waltham, MA). Then 200nL of 5ug/mL of MHCI complex sample were dispensed from the Echo qualified 384- well source plate (Beckman Coulter Life Sciences, Indianapolis, IN)) into the destination plate by an Echo 550 acoustic liquid dispenser (Beckman Coulter Life Sciences, Indianapolis, IN). After centrifugation for three minutes, two μL of master mix donor at 2nM (Europium mouse anti-human β2-microglobulin (β2M), Biolegend, San Diego, CA, custom labeled by Perkin Elmer, Waltham, MA) and acceptor at 40nM (SureLight Allophycocyanin conjugated Streptavidin (SA-APC), PerkinElmer, Waltham, MA) in assay diluent were dispensed into each well of the destination plate with the Multidrop™ Combi nL dispenser. Following incubation at room temperature (RT) for one hour, the destination plates were read on the PHERAstar FS plate reader (BMG Labtech, Cary, NC) with donor excitation at 337nm, donor emission at 615nm and acceptor emission at 665nm.

The signal was expressed as the ratio of relative fluorescent units (RFU) in each well (RFU ratio = (665nm/615nm) X 10^4)^. For ranking the binders, a double normalization was applied to obtain % DeltaF. DeltaF(%) = {(RFU [Sample] – mean RFU [negative])/mean RFU[negative]} * 100. The Robust Z score was calculated on the sample plate basis. For screening quality control, large-scale prepared positive control (A*02:01 with pp65) and negative control (A*02:01 only) were added to designed wells in each sample plate. The acceptance of the screen was determined by Z-factor calculated from the assay controls (Z-factor = 1-{(3SD [positive] – 3SD [negative])/(mean [positive] – mean [negative]}. Sample plates had Z-factor > 0.4 were qualified for data process.

### Antibody coupling and crosslinking

Pan-HLA Class I-specific antibody (clone W6/32) was coupled to Protein-A resin packed into AssayMAP Bravo compatible large capacity cartridges (PA-W 25 μL) (Agilent, Part number G5496-60018). The coupled antibodies were then crosslinked with 20 mM Dimethyl pimelimidate dihydrochloride (DMP) (Sigma-Aldrich, cat. D8388-250 MG) in 100 mM sodium borate crosslinking buffer at pH 9.0 immediately after the end of the coupling step. The impurities within the cartridges were washed away with simultaneous dispensing of 200 mM ethanolamine (Sigma-Aldrich, cat. E9508-100ML) pH 8.0 and deionized H_2_O. The flow rates and other parameters of the affinity purification application within the AssayMAP Bravo software (VWorks) were used at the default settings. The antibody crosslinked Protein-A cartridges were stored in rack filled with TBS/0.025% sodium azide, sealed with parafilm, and kept at 4°C.

### Affinity purification of HLA-peptide complexes

Engineered monoallelic cell pellets (500 million cells/sample) were lysed at 4°C in 1% CHAPS (Roche Diagnostics, cat no. 10810126001) lysis buffer (pH 8.0) containing 20 mM TRIS, 150 mM NaCl, one tablet of cOmplete Protease Inhibitor Cocktail (Roche, cat. 4693159001) per 10 mL of the lysis buffer, and 0.2 mM phenylmethylsulfonyl fluoride (PMSF) (Sigma-Aldrich). The cell pellets were lysed with 2 mL of the lysis buffer by vortexing every 5 minutes for a total of 20 minutes at 4°C. The lysates were then transferred to LoBind tubes and centrifuged at 20,000 g at 4°C for 20 minutes. The supernatants were then carefully transferred to 0.45 μm polyethersulfone filter (Pall, cat. MCPM45C68). The samples were then centrifuged at 7000 g at 4°C for 30 minutes. The filtrate for each sample was carefully transferred to an AssayMAP Bravo compatible 96-well deep well plate making sure not to disturb any particulates that might have settled at the bottom of the conical tube. The deep well plate containing HLA-peptide complexes was transferred to the AssayMAP Bravo sample loading platform for automated dispensing of the samples through the W6/32 crosslinked Protein-A cartridges. The cartridges were primed and equilibrated with 20 mM Tris pH 8.0 and 150 mM NaCl in water. The sample impurities within the cartridges were washed away with automated dispensing of 20 mM Tris pH 8.0 and 400 mM NaCl in water followed by final wash with 20 mM Tris pH 8.0 in water. The antibody-bound HLA-peptide complexes were eluted with 0.1 M acetic acid in 0.1% trifluoroacetic acid (TFA). The flow rates and wash cycles were used at the default settings.

The eluates were transferred to ultra low adsorption *ProteoSave* autosampler vials (AMR Incorporated cat. PSVial100) and dried in a speed vacuum. The dried samples were then reconstituted in 100 uL 20 mM HEPES pH 8.0, reduced with 5 mM Dithiothreitol (DTT) (Thermofisher, cat. A39255) in the dark at 65°C for 30 minutes, and alkylated with 15 mM Iodoacetamide (IAA) (Sigma-Aldrich, cat. I1149-5G) in the dark at RT for 30 minutes. The samples were then acidified with TFA to pH ∼3.0, vortexed, and centrifuged at 14,000 g at RT for 5 minutes to pellet any debris. The samples were then carefully transferred to 96 well PCR, Full Skirt, PolyPro plate (Eppendorf, Part number 30129300) and loaded on the AssayMAP Bravo platform for final clean up before injection into the mass spectrometer. Four C18 cartridges (Agilent, Part number 5190-6532) were used per sample. The cartridges were primed with 80% acetonitrile (ACN) 0.1% TFA and equilibrated with 0.1% TFA. The samples were then loaded through the cartridges, washed with 0.1% TFA, and eluted with 30% ACN 0.1% TFA. After drying the samples in a speed vacuum, the samples were reconstituted in 6 uL 0.1% formic acid (FA) 0.05% heptafluorobutyric acid (HFBA) (Thermo Fisher Scientific, cat. 25003).

### Untargeted Mass Spectrometry and Database Search

One-third of each sample was loaded into a 25 cm x 75 μm ID, 1.6 μm C18 IonOpticks Aurora Series column (IonOpticks, Part Number AUR2-25075C18A) on a Thermo UltiMate 3000 high performance liquid chromatography (HPLC) system (Thermo Fisher Scientific) at a flow rate of 400 nL/min. Peptides were separated with a 90 minute gradient of 2% to 35% or 40% buffer B (98% ACN, 2% H_2_O, and 0.1% FA) at a flow rate of 300 nL/min. The gradient was further raised to 75% buffer B for 5 minutes and to 90% buffer B for 4 minutes at the same flow rate before final equilibration with 98% buffer A (98% H_2_O, 2% ACN, and 0.1% FA) and 2% buffer B for 10 minutes at a flow rate of 400 nL/min.

Peptide mass spectra were acquired using either Orbitrap Fusion Lumos or Orbitrap Eclipse Tribrid mass spectrometer (Thermo Fisher Scientific) with MS^1^ Orbitrap resolution of 240000 and MS/MS fragmentation of the precursor ions by collision-induced dissociation (CID) followed by spectra acquisition at MS^2^ Orbitrap resolution of 15000. All data-dependent acquisition (DDA) spectral raw files were searched in PEAKSOnline (Bioinformatics Solutions Inc.) against a Uniprot-derived *Homo sapiens* human proteome (downloaded October 03, 2019) that contained appended concatenated sequences of 47 most common mutations flanked by ∼13-mer sequences on either end of each mutation with or without stretches of Glycine and Serine residue (GS) linkers along with sequences of blue fluorescence protein (BFP). Within PEAKSOnline, since HLA-peptides are non-tryptic the enzyme specificity was set as none, CID was selected as an activation method, and Orbitrap (Orbi-Orbi) was chosen as an instrument parameter. In-depth de novo assisted database search and quantification were performed with precursor mass error tolerance of 15 parts per million (ppm), fragment mass error tolerance of 0.02 Da, and missed cleavage allowance of 3. Carbamidomethylation (Cys+57.02) was set as a fixed modification whereas deamidation (Asn+0.98, Gln+0.98) and oxidation (Met+15.99) were set as variable post translational modification (PTM) allowing a maximum of 3 variable PTMs per peptide. Additional report filters included peptide spectral match (PSM) false discovery rate (FDR) of 1%, Proteins - 10LgP ≥ 20, and denovo only amino acid residue average local confidence (ALC) of 50%. For label free analysis, a new group was created for each sample and match between runs was performed with default parameters except retention time (RT) shift tolerance was set to 4 minutes and base sample was selected as “Average”. Output csv files were exported and further analyzed in R.

### Targeted Mass Spectrometry

Absolute quantification (AQUA) synthetic heavy peptides (8-11mer) (Elim Biopharm) for all 47 mutation-derived neoantigens with TR-FRET RZ-score ≥ 5 (i.e. RobustzScore ≥ 5) or predicted NetMHC %Rank ≤ 2 (for a subset of mutations) were reconstituted in 30% ACN 0.1% FA. Dimethyl sulfoxide (DMSO) was added for peptides that were not readily soluble in 30 % ACN 0.1% FA. A working solution of 25 μM was made for each AQUA peptide from which allele-specific mastermix was made at 25 pmol/peptide. The peptides were reduced/alkylated and cleaned up with C18 cartridges on AssayMAP Bravo. After drying, the peptides were reconstituted in 0.1% FA 0.05% HFBA at 100 fmol/peptide. For each allele-specific assay the intact modified mass was calculated for each peptide in that assay using TomahaqCompanion software (Rose et al., 2019) which was then used to build an inclusion list mass spectrometry method for a scouting run to get the RT and mass-to-charge (*m/z*) of each target peptide. 1 μL of each assay was injected into the IonOpticks C18 column and sprayed into the mass spectrometer for a 125 minutes run as described above and the raw files were imported and analyzed in Skyline (64-bit, 19.1.0.193) to select appropriate charge for each peptide. A mass list table was built for each assay where 4 minutes RT window was created on both sides of the RT for each target peptide which was then imported into Xcalibur instrument method application and saved as an allele-specific parallel reaction monitoring (PRM) method. For both Fusion Lumos and Eclipse instruments MS^1^ was acquired at Orbitrap resolution of 240000 with a maximum injection time of 50 ms followed by quadrupole isolation window of 1.2 *m/z*, CID fragmentation of parent ions, maximum injection time of 300 ms, and MS^2^ acquisition at Orbitrap resolution of 60000. For Eclipse acquisition MS^1^ and MS^2^ AGC targets were set at 250% and 400% respectively. One-third of each monoallelic sample was spiked with 100 fmol of corresponding AQUA mastermix and injected into the mass spectrometer using the same HPLC set up as described above. Raw PRM data were imported and analyzed in Skyline in allele-specific manner. The ratios of the light peptides to their heavy counterparts across samples were exported as csv files and further analyzed in R. For each neoepitope, background signal detected in the synthetic peptide only analysis was subtracted from endogenous peptide signal before calculation of a final attomole amount.

For KRAS wild type (WT) and G12C/D/V mutant copy number presentation quantification in dox-inducible C1R A*11:01 KRAS full length (FL) cell lines and C1R A*11:01 47-neo sample recombinant heavy isotope-coded peptide MHC (hipMHC) (PMCID PMC7265461) monomers were made in-house for A*11:01 allele and KRAS WT/G12C/D/V (9 and 10 mer per target). These monomers were spiked at 1 pmol per 500 million cell lysate (KRAS FL samples) or 4.7 pmol per 500 million cell lysate (47-neo sample) immediately before the pan HLA Class I affinity purification step. Similar to shared neoantigen samples, an inclusion method and a 125 minutes PRM method were developed for A*11:01 KRAS FL samples where only 8 AQUA peptides were in the hipMHC assay mix. The raw data were analyzed as described above with additional steps where on-column AQUA peptide concentration and input cell count were taken into consideration to calculate antigen copies per cell.

For absolute quantification of total KRAS WT and G12C/D/V proteins in dox-inducible C1R A*11:01 KRAS FL cell lines 20 million cells per sample were lysed in 1 mL 8M Urea lysis buffer 20 mM HEPES pH 8.0. 25 μg of yeast digest and 50 μg from each sample were spiked with 2.5 pmol KRAS quantification concatemer (QconCAT) polypeptide (Polyquant) generated by concatenation of heavy WT and select mutant RAS tryptic proteotypic peptides. Samples were reduced with 5 mM DTT in the dark at 56°C for 10 minutes with shaking and alkylated with 15 mM IAA in the dark at RT for 15 minutes. Urea concentration across control and sample tubes were dropped to ∼2M with 20 mM HEPES pH 8.0 and were digested with 1 μg sequencing grade trypsin (Promega) overnight at 37°C in a nutator. Next day, trypsinization was quenched with 50% TFA and samples were cleaned up on C18 cartridges, dried, and reconstituted at 100 fmol/μL (yeast digest control) or 50 fmol/μL (samples) in 0.1% FA. The digested samples were run on Fusion Lumos mass spectrometer with a 65 minutes PRM method specific for RAS tryptic peptides present on QconCAT polypeptide. Data were analyzed on skyline and absolute quantification of each of the target KRAS peptides was calculated.

### TCR discovery

A total of 376 predicted and mass-spec identified neoantigen-derived peptides were synthesized (GenScript) and each was added to 6 of 11 peptide pools such that each neoepitope (or group of similar neoepitopes) occupied a unique combination of 6 pools (Klinger et al., 2015). CD8+ T cells were isolated (StemCell Technologies) from healthy human donor Leukopaks and expanded either on anti-CD3 coated plates (+anti-CD28/IL-2, BioLegend), or in the presence of matched donor-derived monocyte-derived dendritic cells (Wölfl and Greenberg, 2014) and a pool of all 376 neoepitopes. At day 10-15, T cells were recovered, supplemented with 1 of the 11 neoepitope pools, incubated 8-14 hours, enriched (Miltenyi Biotec) and then sorted using an anti-CD137 antibody (BioLegend). Sorted cells were then subjected either to immunoSEQ or pairSEQ (Adaptive Biotechnologies) to identify TCRB sequences displaying neoepitope-specific responsiveness and to associate TCRB with TCRA sequences in parallel, respectively. TCR sequences were encoded in pcDNA vectors as a single open reading frame, in the form of the full TCRB sequence followed by an RAKR motif and porcine teschovirus 2a cleavage peptide with the full TCRA sequence following in frame. TCR-encoding pcDNA vectors were then used as templates to generate TCR-encoding *in vitro* transcribed RNA (ivtRNA; mMessage mMachine, ThermoFisher) for electroporation of primary human T cells.

### TCR reactivity assays

CD8+ cells were enriched from human PBMCs with EasySep Human CD8+ T Cell Isolation Kit (Stem Cell Technologies) and stimulated with 5 ug/mL Ultra-LEAF anti-human CD3 (Biolegend) and 2.5 ug/mL Ultra-LEAF anti-human CD28 (Biolegend). Cells were cultured in the presence of 20 ng/mL recombinant human IL-2 for 6 days. Human expanded CD8^+^ T cells were transfected with FLT3-p.D835Y- or PIK3CA-p.E545K-specific TCR RNA using a Lonza 4D-Nucleofector, P3 primary cell 4D-nucleofector kit, program EO-115 (Lonza). RNA was purchased from Trilink or *in vitro* transcribed. FLT3-p.D835Y-specific TCRs were co-cultured overnight with HLA-A*02:01- expressing T2 cells pulsed with YIMSDSNYV or HLA-A*02:01-expressing K562 cells transfected with a construct encoding the mutant or wildtype sequence. K562 cells were transfected using a Lonza 4D-Nucleofector, SF cell line 4D-nucleofector kit, program FF-120 (Lonza). To determine specific cell lysis, an equal mixture of transfected HLA-A*02:01+ K562 cells and untransfected cellTrace FarRed (Thermofisher)-labeled HLA-A*02:01+ K562 cells were co-cultured overnight with T cells at a 2:1 E:T ratio. Percent (%) Specific Cell Lysis = (P_mock-transfected T-cells_ – P_TCR-transfected T-cells_)/(P_mock-transfected T-cells_)) x 100, where P is the proportion of transfected K562 targets relative to an untransfected K562 cells, as measured by flow cytometry. CD137 expression on CD8+ T cells was assessed after an overnight co-culture with an anti-CD137 PE antibody (BD Biosciences).

PIK3CA-p.E545K-specific TCRs were co-cultured overnight with HLA-A*11:01-expressing K562s pulsed with STRDPLSEITK or transfected with a construct encoding the mutant or wildtype sequence. Equal mixtures of cellTrace Far Red-labeled HLA-A*11:01+ K562 cells were added to each well. T cell response to PIK3CA-presenting K562 cells was assessed as above.

## Key Resources Table

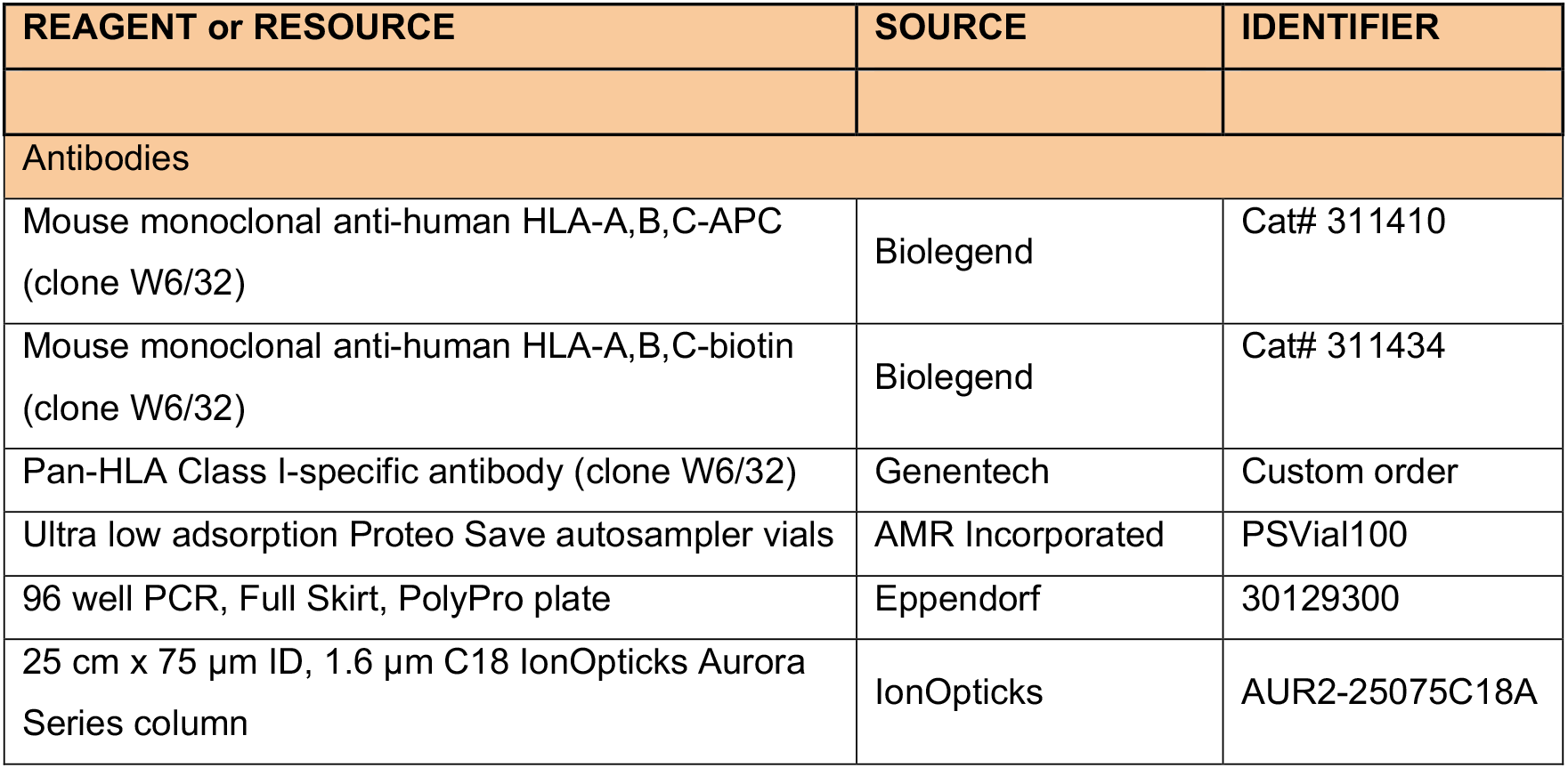
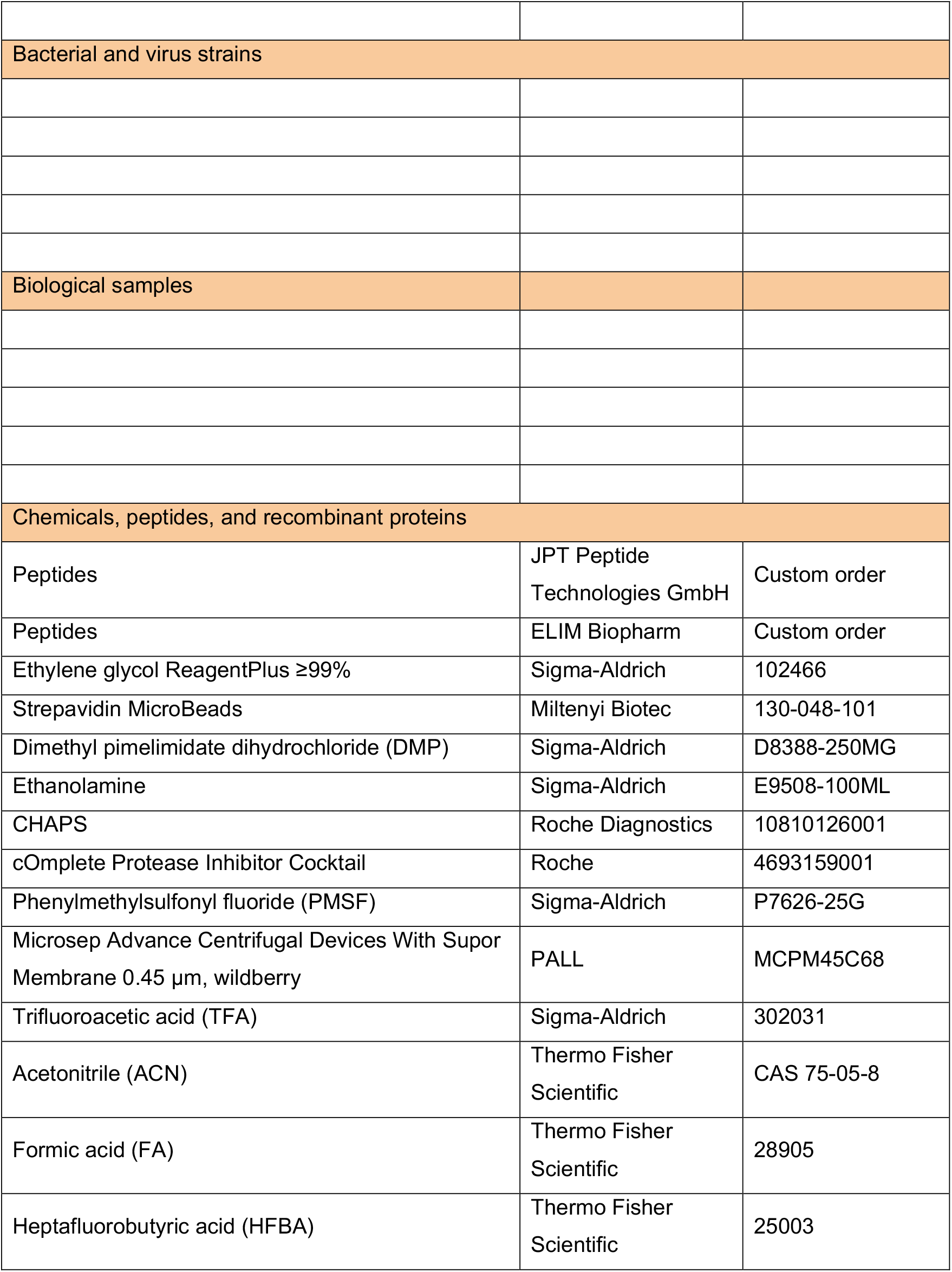
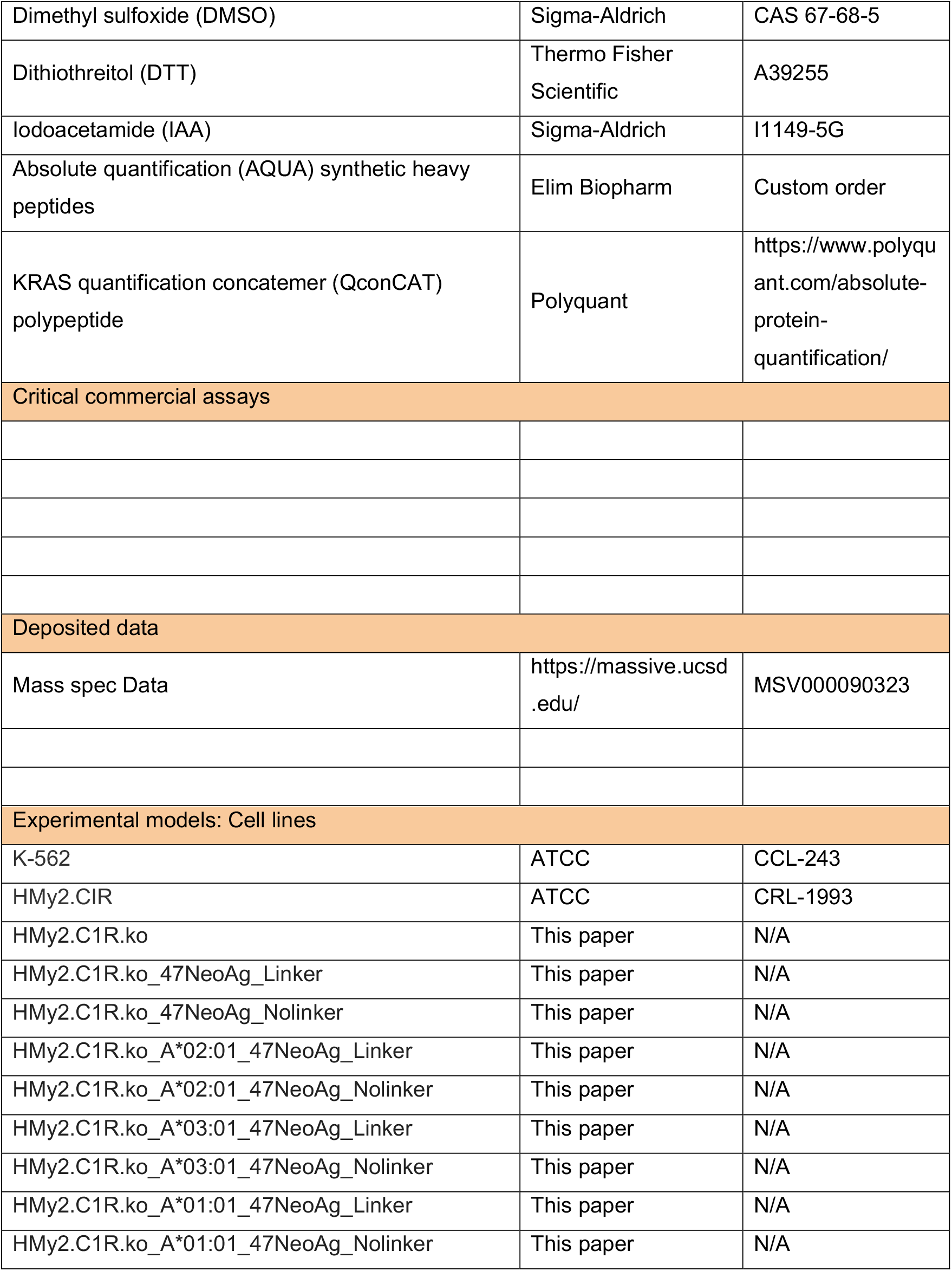
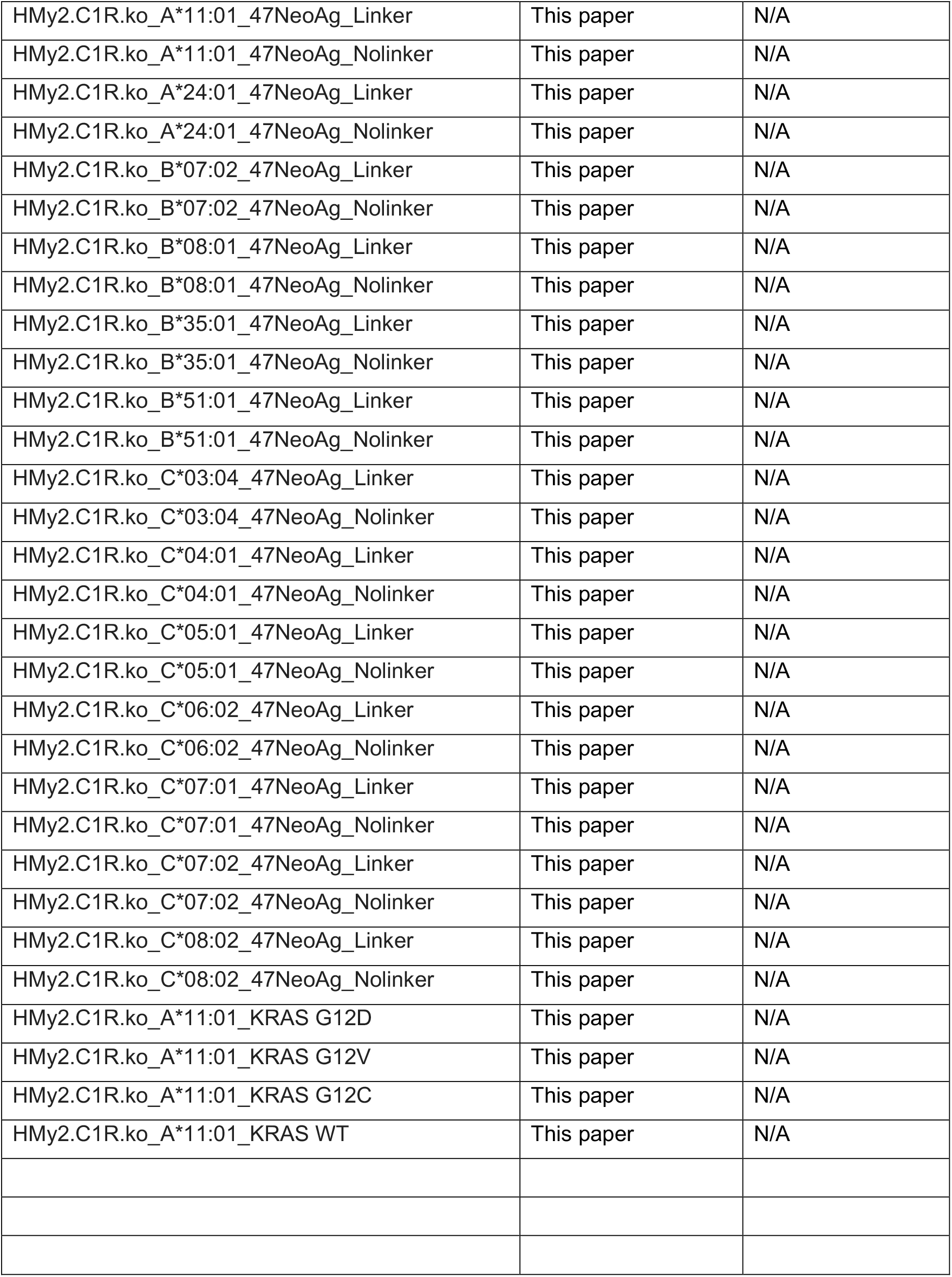
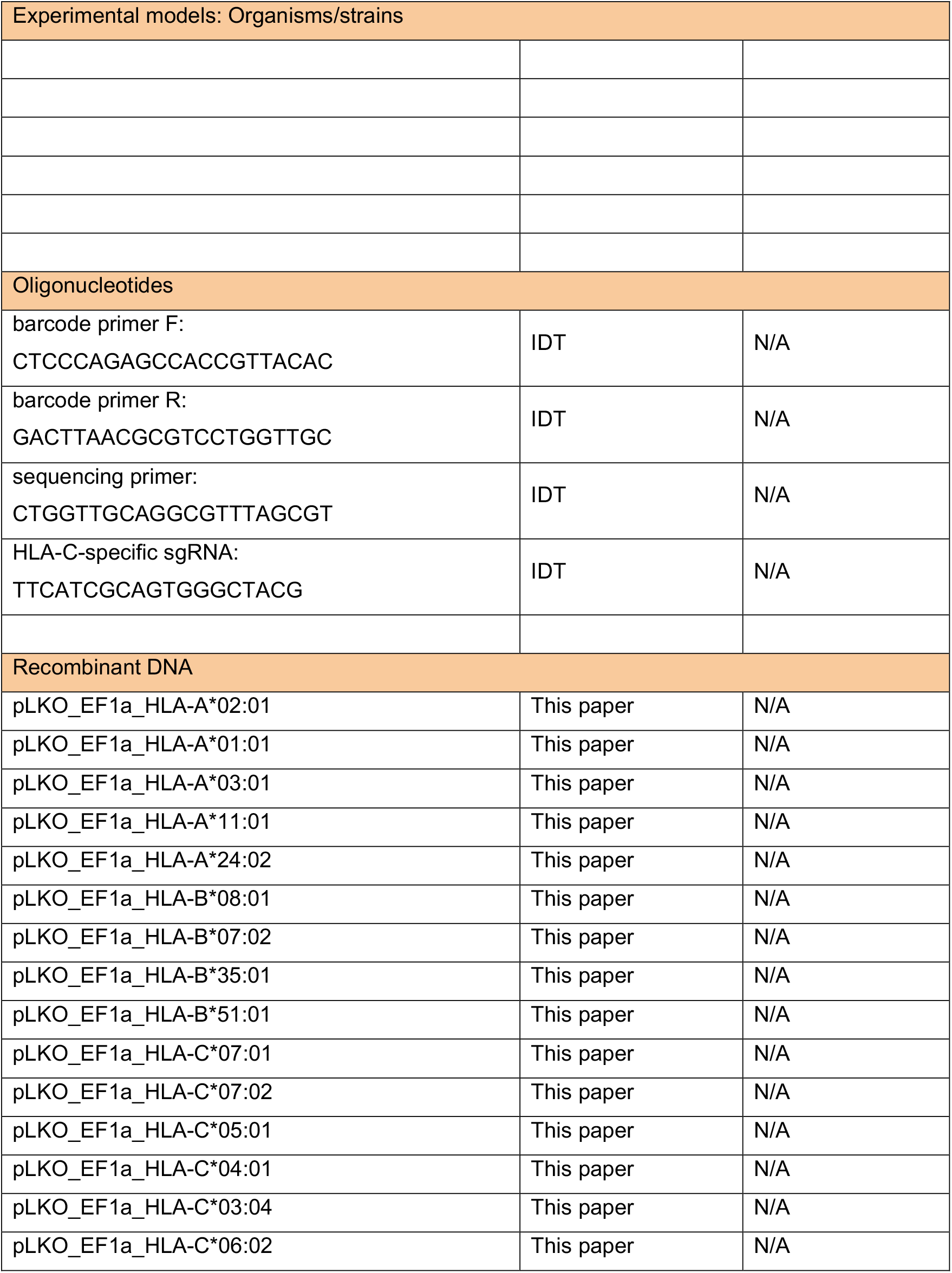
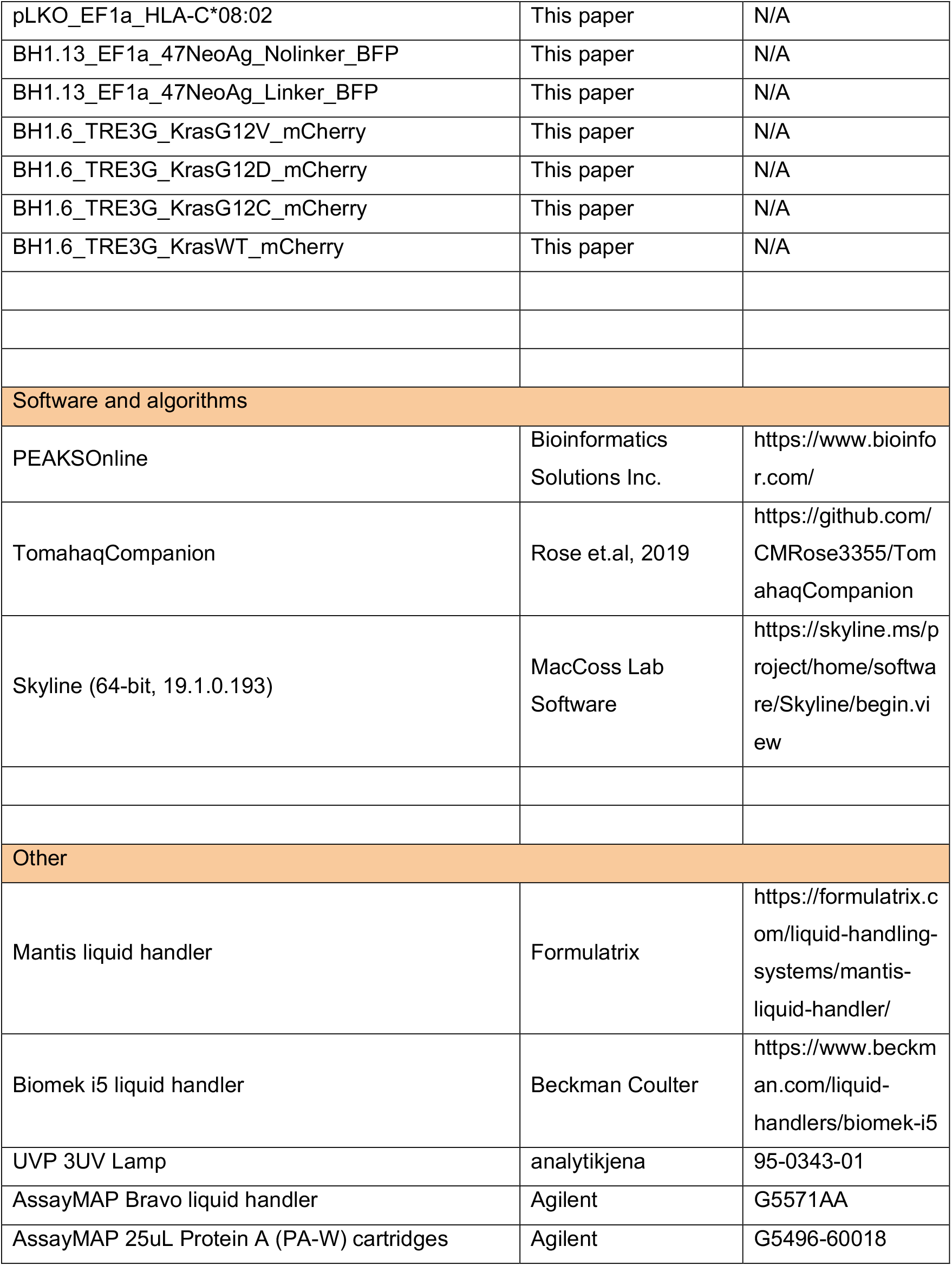
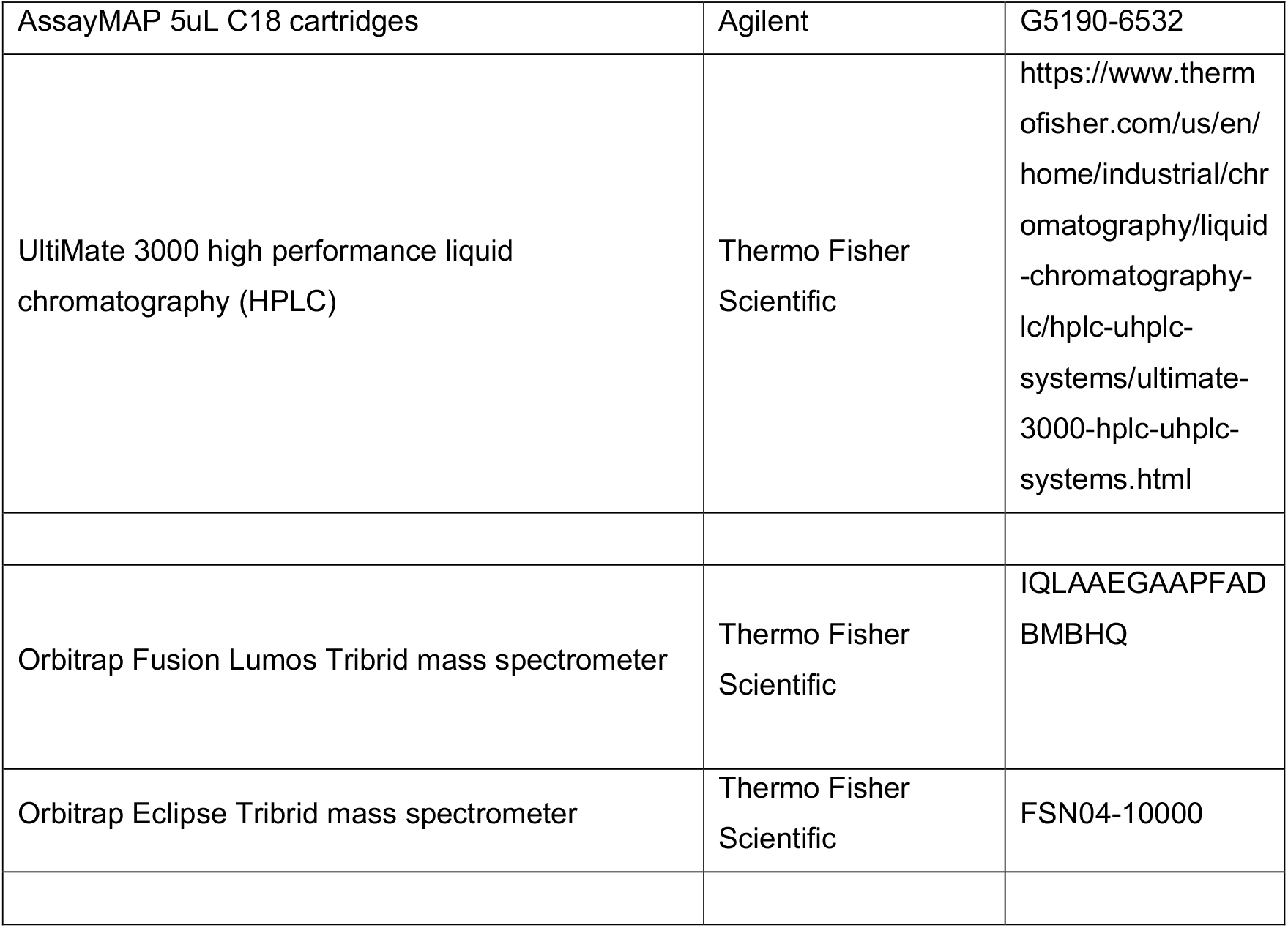

