## Supplemental Sequences for "Discovery of prevalent, clinically actionable tumor neoepitopes via integrated biochemical and cell-based platforms"

```
PiggyBac No Linker Neoantigens BFP

LOCUS       BH1.13_EF1a_47NeoAg_Nolinker_BFP               12954 bp DNA     circular SYN 24-OCT-2022
DEFINITION  synthetic circular DNA
ACCESSION   .
VERSION     .
KEYWORDS    .
SOURCE      synthetic DNA construct
  ORGANISM  synthetic DNA construct
REFERENCE   1  (bases 1 to 12954)
  AUTHORS   .
  TITLE     Direct Submission
  JOURNAL   Exported Oct 25, 2022 from SnapGene 6.0.5
            https://www.snapgene.com
FEATURES             Location/Qualifiers
     source          1..12954
                     /mol_type="other DNA"
                     /organism="synthetic DNA construct"
     CDS             complement(239..307)
                     /label=LacZ alpha
     primer_bind     378..395
                     /label=M13-fwd
     primer_bind     379..395
                     /label=M13 fwd
     protein_bind    complement(445..682)
                     /label=PB-TRE 874..1111
     enhancer        719..950
                     /label=Insulator 1148..1379
     enhancer        720..950
                     /label=Insulator 1701..1931
     misc_feature    1084..1085
                     /label=ClaI_SpeI fragment PCR clone
     promoter        1243..2424
                     /label=EF-1-alpha promoter
     intron          1473..2415
                     /label=EF-1-alpha intron A
     regulatory      2444..2449
                     /label=Kozak sequence
     CDS             2450..6856
                     /label=47nAg_NOLinker
                     /label=48nAg_NOLinker_all_controls_mmCO_ORF
     CDS             2450..2524
                     /codon_start=1
                     /label=KRAS G12D
                     /translation="MTEYKLVVVGADGVGKSALTIQLIQ"
     CDS             2525..2605
                     /codon_start=1
                     /label=JAK2 V617F
                     /translation="LSHKHLVLNYGVCFCGDENILVQEFVK"
     misc_feature    2606..2683
                     /label=KRAS G13D
     misc_feature    2684..2764
                     /label=NRAS Q61R/HRAS Q61R
     misc_feature    2765..2845
                     /label=EGFR G719A
     misc_feature    2846..2926
                     /label=EGFR T790M
     misc_feature    2927..3028
                     /label=EGFR T790M_C797S
     misc_feature    3029..3103
                     /label=KRAS G12S
     misc_feature    3104..3184
                     /label=PIK3CA E542K
     misc_feature    3185..3265
                     /label=PIK3CA H1047L
     misc_feature    3266..3346
                     /label=ERBB2 S310F
     misc_feature    3347..3427
                     /label=BRAF V600M
     misc_feature    3428..3508
                     /label=FGFR3 S249C
     misc_feature    3509..3589
                     /label=MYD88 L265P
     misc_feature    3590..3670
                     /label=PTEN R130Q
     misc_feature    3671..3751
                     /label=TP53 R248Q
     misc_feature    3752..3832
                     /label=TP53 R273H
     misc_feature    3833..3913
                     /label=TP53 R282W
     misc_feature    3914..3994
                     /label=ESR1 K303R
     misc_feature    3995..4075
                     /label=GNA11 Q209L
     misc_feature    4076..4156
                     /label=GNAQ Q209P
     misc_feature    4157..4237
                     /label=IDH1 R132C
     misc_feature    4238..4318
                     /label=IDH1 R132H
     misc_feature    4319..4399
                     /label=SF3B1 R625C
     misc_feature    4400..4474
                     /label=KRAS G12C
     misc_feature    4475..4549
                     /label=KRAS G12V
     misc_feature    4550..4630
                     /label=NRAS Q61K
     misc_feature    4631..4699
                     /label=EGFR E746_A750del
     misc_feature    4700..4780
                     /label=EGFR L858R
     misc_feature    4781..4861
                     /label=EGFR C797S
     misc_feature    4862..4936
                     /label=KRAS G12A
     misc_feature    4937..5014
                     /label=KRAS G13C
     misc_feature    5015..5095
                     /label=PIK3CA E545K
     misc_feature    5096..5176
                     /label=PIK3CA H1047R
     misc_feature    5177..5257
                     /label=BRAF V600E
     misc_feature    5258..5338
                     /label=DNMT3A R882H
     misc_feature    5339..5413
                     /label=KRAS G12R
     misc_feature    5414..5494
                     /label=PTEN R130G
     misc_feature    5495..5575
                     /label=TP53 R175H
     misc_feature    5576..5656
                     /label=TP53 R273C
     misc_feature    5657..5737
                     /label=TP53 R273L
     misc_feature    5738..5800
                     /label=CALR fs
     misc_feature    5801..5881
                     /label=FLT3 D835Y
     misc_feature    5882..5962
                     /label=GNAQ Q209L
     misc_feature    5963..6043
                     /label=BRAF V600M
     misc_feature    6044..6124
                     /label=IDH1 R132G
     misc_feature    6125..6205
                     /label=IDH2 R140Q
     misc_feature    6206..6286
                     /label=SF3B1 R625H
     CDS             6287..6367
                     /codon_start=1
                     /label=pp65 495-503
                     /translation="PWQAGILARNLVPMVATVQGQNLKYQE"
     CDS             6368..6448
                     /codon_start=1
                     /label=BMLF1
                     /translation="ARMQAIQNAGLCTLVAMLEETIFWLQE"
     CDS             6449..6529
                     /codon_start=1
                     /label=Influenza M
                     /translation="RPILSPLTKGILGFVFTLTVPSERGLQ"
     CDS             6530..6610
                     /codon_start=1
                     /label=IE-1
                     /translation="EFCRVLCCYVLEETSVMLAKRPLITKP"
     CDS             6611..6691
                     /codon_start=1
                     /label=Mart1
                     /translation="GIGILTVADELAGIGILTVADILTVIL"
     CDS             6692..6772
                     /codon_start=1
                     /label=MAGE A3
                     /translation="PGSDPACYEFLWGPRALVETSYVKVLH"
     CDS             6773..6853
                     /codon_start=1
                     /label=NY-ESO1
                     /translation="SISSCLQQLSLLMWITQVFLPVFLAQP"
     misc_feature    6945..7496
                     /label=IRES
     CDS             7508..8218
                     /label=mTagBFP2
     misc_feature    8223..8227
                     /label=via pLENTI6.3_ClaI_NheI
     polyA_signal    8234..8282
                     /label=poly(A) signal
     misc_feature    8284..8313
                     /label=buffer sequence
     polyA_signal    8314..8521
                     /label=bGH poly(A) signal
     misc_feature    8527..8644
                     /label=cPPT/CTS
     promoter        8669..9169
                     /label=hPGK promoter
     CDS             9183..9779
                     /label=PuroR
     polyA_signal    9789..9910
                     /label=SV40 poly(A) signal
     enhancer        10007..10238
                     /label=Insulator 1148..1379
     enhancer        10007..10237
                     /label=Insulator 1701..1931
     primer_bind     complement(10277..10296)
                     /label=T3
     promoter        complement(10279..10297)
                     /label=T3 promoter
     primer_bind     10304..10323
                     /label=T7
     promoter        10304..10322
                     /label=T7 promoter
     protein_bind    10373..10681
                     /label=PB-TRE 2067..2375
     misc_feature    10715
                     /label=HindIII killer
     primer_bind     complement(10729..10749)
                     /label=M13-rev
     primer_bind     complement(10733..10749)
                     /label=M13 rev
     misc_binding    complement(10755..10777)
                     /label=LacO
     protein_bind    10757..10773
                     /label=lac operator
     promoter        complement(10781..10811)
                     /label=lac promoter
     promoter        complement(10781..10787)
                     /label=lac promoter
     promoter        complement(10788..10805)
                     /label=lac promoter
     promoter        complement(10806..10811)
                     /label=lac promoter
     protein_bind    10826..10847
                     /label=CAP binding site
     rep_origin      complement(11117..11799)
                     /direction=LEFT
                     /label=ColE1 origin
     rep_origin      complement(11135..11723)
                     /direction=LEFT
                     /label=ori
     CDS             complement(11894..12685)
                     /label=AmpR
     CDS             complement(11897..12556)
                     /label=AmpR
     CDS             complement(12686..12754)
                     /label=AmpR
     promoter        complement(12755..12859)
                     /label=AmpR promoter
ORIGIN
        1 tcgcgcgttt cggtgatgac ggtgaaaacc tctgacacat gcagctcccg gagacggtca
       61 cagcttgtct gtaagcggat gccgggagca gacaagcccg tcagggcgcg tcagcgggtg
      121 ttggcgggtg tcggggctgg cttaactatg cggcatcaga gcagattgta ctgagagtgc
      181 accatatgcg gtgtgaaata ccgcacagat gcgtaaggag aaaataccgc atcaggcgcc
      241 attcgccatt caggctgcgc aactgttggg aagggcgatc ggtgcgggcc tcttcgctat
      301 tacgccagct ggcgaaaggg ggatgtgctg caaggcgatt aagttgggta acgccagggt
      361 tttcccagtc acgacgttgt aaaacgacgg ccagtgaatt aggcgcgcct gctcgacacg
      421 ctgcagaaca cgcagctaga ttaaccctag aaagataatc atattgtgac gtacgttaaa
      481 gataatcatg cgtaaaattg acgcatgtgt tttatcggtc tgtatatcga ggtttattta
      541 ttaatttgaa tagatattaa gttttattat atttacactt acatactaat aataaattca
      601 acaaacaatt tatttatgtt tatttattta ttaaaaaaaa acaaaaactc aaaatttctt
      661 ctataaagta acaaaacttt tatcgaattg ctgcagcccg ggcgatccat tagtactaga
      721 gggacagccc ccccccaaag cccccaggga tgtaattacg tccctccccc gctagggggc
      781 agcagcgagc cgcccggggc tccgctccgg tccggcgctc cccccgcatc cccgagccgg
      841 cagcgtgcgg ggacagcccg ggcacgggga aggtggcacg ggatcgcttt cctctgaacg
      901 cttctcgctg ctctttgagc ctgcagacac ctggggggat acggggaaaa ggcctccaag
      961 gccagcttcc cacaataagt tgggtgaatt ttggctcatt cctcctttct ataggattga
     1021 ggtcagagct ttgtgatggc aattctgtgg aatgtgtgtc agttagggtg tggaaagtca
     1081 agcatcgatg agtaattcat acaaaaggac tcgcccctgc cttggggaat cccagggacc
     1141 gtcgttaaac tcccactaac gtagaaccca gagatcgctg cgttcccgcc ccctcacccg
     1201 cccgctctcg tcatcactga ggtggagaag agcatgcgtg aggctccggt gcccgtcagt
     1261 gggcagagcg cacatcgccc acagtccccg agaagttggg gggaggggtc ggcaattgaa
     1321 ccggtgccta gagaaggtgg cgcggggtaa actgggaaag tgatgtcgtg tactggctcc
     1381 gcctttttcc cgagggtggg ggagaaccgt atataagtgc agtagtcgcc gtgaacgttc
     1441 tttttcgcaa cgggtttgcc gccagaacac aggtaagtgc cgtgtgtggt tcccgcgggc
     1501 ctggcctctt tacgggttat ggcccttgcg tgccttgaat tacttccacg cccctggctg
     1561 cagtacgtga ttcttgatcc cgagcttcgg gttggaagtg ggtgggagag ttcgaggcct
     1621 tgcgcttaag gagccccttc gcctcgtgct tgagttgagg cctggcttgg gcgctggggc
     1681 cgccgcgtgc gaatctggtg gcaccttcgc gcctgtctcg ctgctttcga taagtctcta
     1741 gccatttaaa atttttgatg acctgctgcg acgctttttt tctggcaaga tagtcttgta
     1801 aatgcgggcc aagatctgca cactggtatt tcggtttttg gggccgcggg cggcgacggg
     1861 gcccgtgcgt cccagcgcac atgttcggcg aggcggggcc tgcgagcgcg gccaccgaga
     1921 atcggacggg ggtagtctca agctggccgg cctgctctgg tgcctggcct cgcgccgccg
     1981 tgtatcgccc cgccctgggc ggcaaggctg gcccggtcgg caccagttgc gtgagcggaa
     2041 agatggccgc ttcccggccc tgctgcaggg agctcaaaat ggaggacgcg gcgctcggga
     2101 gagcgggcgg gtgagtcacc cacacaaagg aaaagggcct ttccgtcctc agccgtcgct
     2161 tcatgtgact ccacggagta ccgggcgccg tccaggcacc tcgattagtt ctcaagcttt
     2221 tggagtacgt cgtctttagg ttggggggag gggttttatg cgatggagtt tccccacact
     2281 gagtgggtgg agactgaagt taggccagct tggcacttga tgtaattctc cttggaattt
     2341 gccctttttg agtttggatc ttggttcatt ctcaagcctc agacagtggt tcaaagtttt
     2401 tttcttccat ttcaggtgtc gtgaccctag cgctacctct agagccacca tgactgagta
     2461 caagttggtc gtggtcggag ccgacggcgt gggaaagtca gcactgacta ttcagctcat
     2521 tcagttgagc cacaagcatt tggtgttgaa ctacggcgtg tgcttttgcg gtgacgagaa
     2581 tattttggtg caggagtttg ttaagatgac cgagtataaa ttggtcgtcg tcggtgcagg
     2641 tgatgtaggg aaatctgccc tgacaatcca acttattcag aatggcgaaa catgtctcct
     2701 ggacatcctt gataccgcag ggcgggagga atatagcgcc atgagagatc aatacatgcg
     2761 aactatcctc aaggagactg agtttaaaaa aatcaaggtt cttgcatctg gtgcctttgg
     2821 gacagtttac aaggggcttt ggatattgct cgggatttgc ctcacatcaa ctgttcagct
     2881 gattatgcaa ctgatgccct tcggatgttt gcttgattat gtaaggctgt tgggtatatg
     2941 cttgacctcc acagtgcagc tgatcatgca attgatgcca tttggcagtc ttctggatta
     3001 cgtccgagag cataaagata atatcggtat gaccgagtat aagttggtgg tcgtcggcgc
     3061 ttccggagtg gggaagtctg cactgactat tcagctcatc caggaacaac ttaaagcaat
     3121 ctctactaga gatcccctct ctaaaataac tgagcaggag aaggatttcc tttggagtca
     3181 ccgagaagct cttgagtatt ttatgaagca gatgaatgat gcacttcacg gcggctggac
     3241 taccaagatg gattggatat tccatacagc atgtccttat aattacttgt ccactgatgt
     3301 tgggttctgt acattggtct gtcccttgca taatcaagaa gtaaccgatt tgactgtgaa
     3361 gattggcgat ttcgggctcg ccactatgaa gagcaggtgg tctggcagcc atcaattcga
     3421 gcaacttagc atacgacaaa catatacatt ggacgttctt gagagatgcc ctcacagacc
     3481 tatcctgcag gctgggttgc ccgccaattt cgccctttct ctgagtccag gcgcacacca
     3541 aaaaagacct ataccaatta agtataaagc aatgaagaaa gagtttccaa atcatgtcgc
     3601 cgccattcac tgtaaggccg gcaaagggca gaccggcgta atgatttgcg catacctcct
     3661 gcaccgggga aattacatgt gcaactccag ctgcatggga ggcatgaacc agcggcccat
     3721 actgacaatt attacattgg aggacagttc ttcatctgga aacctgttgg gcagaaactc
     3781 ttttgaagtt catgtgtgtg catgtcccgg cagagaccga cgcactgaag agagcttcga
     3841 ggttcgagtt tgtgcctgtc ctggccgaga ctggcgaacc gaagaagaga accttagaaa
     3901 aaaaggagaa cccaatttgt ggccctcccc cctcatgata aaacgaagca agcgaaactc
     3961 tcttgctctg tctttgaccg ccgaccaaat ggtcctcgaa aatattatct ttagaatggt
     4021 tgacgtcggt ggattgcgga gcgagcgacg aaagtggatt cattgtttcg agaaccttca
     4081 atccgttata tttaggatgg tagacgtcgg gggcccacgc agtgaacggc gcaaatggat
     4141 ccattgtttc gagaacagac tggtcagtgg atgggtcaag cccatcatta ttggctgcca
     4201 tgcttatggt gaccaatacc gggctaccga cttcgttcgc ttggtttccg ggtgggtcaa
     4261 gccaattatt ataggtcatc atgcctacgg cgatcaatat cgagcaaccg actttgtcac
     4321 tatgcggcct gacatagaca atatggatga atatgtatgt aatactactg cccgagcttt
     4381 tgctgtcgtg gcctctgcaa tgactgaata taagttggtc gttgttggag cttgcggtgt
     4441 tggcaagtct gctctgacta tacaactgat acaaatgacc gagtacaagc tcgtggtagt
     4501 gggcgcagtt ggagttggca agagcgcttt gacaatccaa cttatacagg gtgaaacctg
     4561 tctgctcgat atacttgata cagccggaaa ggaggagtac tcagccatgc gggaccaata
     4621 tatgcgcacc cccgaggggg agaaggtcaa aattcccgtg gccataaaaa cctcacccaa
     4681 agcaaacaaa gagattctgg tgaaaacacc acaacatgta aagatcaccg acttcggaag
     4741 ggctaagttg ttgggtgctg aggaaaaaga atatcatgca agcaccgtac agcttatcac
     4801 acaactcatg ccctttgggt cacttctgga ttacgtaagg gagcacaagg ataatattgg
     4861 catgaccgag tacaaactcg tggtagtagg agccgctggc gtggggaaga gtgcccttac
     4921 cattcagctt atccagatga cagaatataa actggtcgtc gtgggagctg gatgtgttgg
     4981 gaaaagcgca cttacaatac aactgattca gaacaaagcc atttctacaa gggatccact
     5041 gtcagagatc accaagcaag agaaggattt cttgtggtct cacaggcatt actgtgaagc
     5101 ccttgagtac tttatgaagc agatgaatga tgcccgccat ggggggtgga ctactaaaat
     5161 ggattggatt ttccatgatc tcacagtgaa aatcggagac tttggtcttg ctacagagaa
     5221 atctaggtgg agtggaagcc accaatttga gcaattgggc tttcccgtac attatactga
     5281 cgtgagtaat atgagtcatc tcgcccgaca acgcctcctc ggtcggagtt ggtcagtaat
     5341 gaccgagtat aagctcgttg tggtcggcgc aaggggggtt ggcaaaagtg ctcttaccat
     5401 acaactcatc cagaaccacg tcgccgccat acattgtaaa gctggcaaag gtgggacagg
     5461 tgtcatgata tgtgcttacc ttctgcaccg cggcatatat aagcaaagtc agcatatgac
     5521 cgaggtagtg cggcactgcc cacaccacga gcgatgttct gactccgacg ggctgagttc
     5581 cggaaacctg ttggggcgaa acagttttga ggtctgcgta tgcgcttgtc caggacgaga
     5641 caggcgcacc gaggaatcct ctgggaacct cctgggcaga aattcattcg aagttcttgt
     5701 gtgcgcctgt ccagggcgcg acagacggac tgaagagcgg atgcgacgga tgcgacgcac
     5761 aaggaggaaa atgagaagga agatgtcccc tgcaagaccc gggaaagtgg tgaagatttg
     5821 cgacttcggc ctcgcacgat atattatgtc tgactcaaac tatgttgtgc gggggaatgc
     5881 tttgcaaagt gtcatcttcc gaatggtaga tgtcgggggg ctcaggtcag agaggagaaa
     5941 gtggattcac tgttttgaaa acgatcttac agttaaaata ggtgatttcg ggcttgccac
     6001 aatgaaatct cgatggtcag gaagtcacca atttgagcag ctccgacttg tcagtggatg
     6061 ggtcaagcca attatcatag gtgggcatgc atacggagac caataccggg ccacagattt
     6121 tgttaagctc aagaaaatgt ggaagtcacc aaatggcaca atccagaata tccttggtgg
     6181 caccgttttc cgggaaccca tcataaccat gcgaccagat attgataata tggatgaata
     6241 tgttcataat actaccgccc gagcatttgc tgtcgtggct agtgcaccat ggcaggctgg
     6301 tatcctcgca cggaaccttg tgccaatggt cgccactgta cagggacaaa acctgaagta
     6361 ccaagaggcc cgaatgcaag ccatccaaaa tgctgggctg tgcacactcg ttgccatgct
     6421 ggaagagacc atcttctggc tgcaagagag gccaatcctt agccctctta caaaaggcat
     6481 cttgggcttc gtatttacat tgactgtccc ttcagaaagg ggcttgcaag aattttgtcg
     6541 cgttttgtgt tgttatgtgc ttgaagaaac ttcagttatg ttggctaaac gacccctcat
     6601 aactaagcct ggcatcggta tattgacagt tgcagatgag ttggctggta ttgggatact
     6661 tacagttgcc gatattctga ctgtaattct tcccggatct gatccagctt gctatgagtt
     6721 tctgtgggga ccacgagccc tcgtcgaaac atcctacgtc aaggtcctgc attctataag
     6781 cagctgcctc caacagctca gcctccttat gtggattacc caggttttcc tccctgtatt
     6841 ccttgctcaa ccatgactcg aggttaaccg cccgccccac gacccgcagc gcccgaccga
     6901 aaggagcgca cgaccccatc atccaattcc gccccccccc cctaacgtta ctggccgaag
     6961 ccgcttggaa taaggccggt gtgcgtttgt ctatatgtta ttttccacca tattgccgtc
     7021 ttttggcaat gtgagggccc ggaaacctgg ccctgtcttc ttgacgagca ttcctagggg
     7081 tctttcccct ctcgccaaag gaatgcaagg tctgttgaat gtcgtgaagg aagcagttcc
     7141 tctggaagct tcttgaagac aaacaacgtc tgtagcgacc ctttgcaggc agcggaaccc
     7201 cccacctggc gacaggtgcc tctgcggcca aaagccacgt gtataagata cacctgcaaa
     7261 ggcggcacaa ccccagtgcc acgttgtgag ttggatagtt gtggaaagag tcaaatggct
     7321 ctcctcaagc gtattcaaca aggggctgaa ggatgcccag aaggtacccc attgtatggg
     7381 atctgatctg gggcctcggt gcacatgctt tacatgtgtt tagtcgaggt taaaaaacgt
     7441 ctaggccccc cgaaccacgg ggacgtggtt ttcctttgaa aaacacgatg ataatatggc
     7501 cacaaccatg gtgtctaagg gcgaagagct gattaaggag aacatgcaca tgaagctgta
     7561 catggagggc accgtggaca accatcactt caagtgcaca tccgagggcg aaggcaagcc
     7621 ctacgagggc acccagacca tgagaatcaa ggtggtcgag ggcggccctc tccccttcgc
     7681 cttcgacatc ctggctacta gcttcctcta cggcagcaag accttcatca accacaccca
     7741 gggcatcccc gacttcttca agcagtcctt ccctgagggc ttcacatggg agagagtcac
     7801 cacatacgaa gacgggggcg tgctgaccgc tacccaggac accagcctcc aggacggctg
     7861 cctcatctac aacgtcaaga tcagaggggt gaacttcaca tccaacggcc ctgtgatgca
     7921 gaagaaaaca ctcggctggg aggccttcac cgagacgctg taccccgctg acggcggcct
     7981 ggaaggcaga aacgacatgg ccctgaagct cgtgggcggg agccatctga tcgcaaacgc
     8041 caagaccaca tatagatcca agaaacccgc taagaacctc aagatgcctg gcgtctacta
     8101 tgtggactac agactggaaa gaatcaagga ggccaacaac gagacctacg tcgagcagca
     8161 cgaggtggca gtggccagat actgcgacct ccctagcaaa ctggggcaca agcttaattg
     8221 agctagcgtc gacaataaaa tatctttatt ttcattacat ctgtgtgttg gttttttgtg
     8281 tgtggcacct cctgccatct gtctttccgt ttcctgtgcc ttctagttgc cagccatctg
     8341 ttgtttgccc ctcccccgtg ccttccttga ccctggaagg tgccactccc actgtccttt
     8401 cctaataaaa tgaggaaatt gcatcgcatt gtctgagtag gtgtcattct attctggggg
     8461 gtggggtggg gcaggacagc aagggggagg attgggaaga gaatagcagg catgctgggg
     8521 aaattgtttt aaaagaaaag gggggattgg ggggtacagt gcaggggaaa gaatagtaga
     8581 cataatagca acagacatac aaactaaaga attacaaaaa caaattacaa aaattcaaaa
     8641 ttttacccca gaacgcgtgg ggcccgggtt gcgccttttc caaggcagcc ctgggtttgc
     8701 gcagggacgc ggctgctctg ggcgtggttc cgggaaacgc agcggcgccg accctgggtc
     8761 tcgcacattc ttcacgtccg ttcgcagcgt cacccggatc ttcgccgcta cccttgtggg
     8821 ccccccggcg acgcttcctg ctccgcccct aagtcgggaa ggttccttgc ggttcgcggc
     8881 gtgccggacg tgacaaacgg aagccgcacg tctcactagt accctcgcag acggacagcg
     8941 ccagggagca atggcagcgc gccgaccgcg atgggctgtg gccaatagcg gctgctcagc
     9001 agggcgcgcc gagagcagcg gccgggaagg ggcggtgcgg gaggcggggt gtggggcggt
     9061 agtgtgggcc ctgttcctgc ccgcgcggtg ttccgcattc tgcaagcctc cggagcgcac
     9121 gtcggcagtc ggctccctcg ttgaccgaat caccgacctc tctccccagg gagatctcaa
     9181 ccatgaccga gtacaagccc acggtgcgcc tcgccacccg cgacgacgtc ccccgggccg
     9241 tacgcaccct cgccgccgcg ttcgccgact accccgccac gcgccacacc gtggacccgg
     9301 accgccacat cgagcgggtc accgagctgc aagaactctt cctcacgcgc gtcgggctcg
     9361 acatcggcaa ggtgtgggtc gcggacgacg gcgccgcggt ggcggtctgg accacgccgg
     9421 agagcgtcga agcgggggcg gtgttcgccg agatcggccc gcgcatggcc gagttgagcg
     9481 gttcccggct ggccgcgcag caacagatgg aaggcctcct ggcgccgcac cggcccaagg
     9541 agcccgcgtg gttcctggcc accgtcggcg tctcgcccga ccaccagggc aagggtctgg
     9601 gcagcgccgt cgtgctcccc ggagtggagg cggccgagcg cgccggggtg cccgccttcc
     9661 tggagacctc cgcgccccgc aacctcccct tctacgagcg gctcggcttc accgtcaccg
     9721 ccgacgtcga ggtgcccgaa ggaccgcgca cctggtgcat gacccgcaag cccggtgcct
     9781 gagctagcaa cttgtttatt gcagcttata atggttacaa ataaagcaat agcatcacaa
     9841 atttcacaaa taaagcattt ttttcactgc attctagttg tggtttgtcc aaactcatca
     9901 atgtatctta tgtcctcaca ggaacgaagt ccctaaagaa acagtggcag ccaggtttag
     9961 ccccggaatt gactggattc cttttttagg gcccattggt atggcttttt ccccgtatcc
    10021 ccccaggtgt ctgcaggctc aaagagcagc gagaagcgtt cagaggaaag cgatcccgtg
    10081 ccaccttccc cgtgcccggg ctgtccccgc acgctgccgg ctcggggatg cggggggagc
    10141 gccggaccgg agcggagccc cgggcggctc gctgctgccc cctagcgggg gagggacgta
    10201 attacatccc tgggggcttt gggggggggc tgtccctctt gaacgaccgc caccgcggtg
    10261 gagctccagc ttttgttccc tttagtgagg gttaattaaa tcttaatacg actcactata
    10321 gggcgaattg ggaaccgggc cccccctgga gatcgacggt atccataagc ttgatatcta
    10381 taacaagaaa atatatatat aataagttat cacgtaagta gaacatgaaa taacaatata
    10441 attatcgtat gagttaaatc ttaaaagtca cgtaaaagat aatcatgcgt cattttgact
    10501 cacgcggtcg ttatagttca aaatcagtga cacttaccgc attgacaagc acgcctcacg
    10561 ggagctccaa gcggcgactg agatgtccta aatgcacagc gacggattcg cgctatttag
    10621 aaagagagag caatatttca agaatgcatg cgtcaatttt acgcagacta tctttctagg
    10681 gttaatctag ctgcatcagg atcatagcgg ccgccagctt ggcgtaatca tggtcatagc
    10741 tgtttcctgt gtgaaattgt tatccgctca caattccaca caacatacga gccggaagca
    10801 taaagtgtaa agcctggggt gcctaatgag tgagctaact cacattaatt gcgttgcgct
    10861 cactgcccgc tttccagtcg ggaaacctgt cgtgccagct gcattaatga atcggccaac
    10921 gcgcggggag aggcggtttg cgtattgggc gctcttccgc ttcctcgctc actgactcgc
    10981 tgcgctcggt cgttcggctg cggcgagcgg tatcagctca ctcaaaggcg gtaatacggt
    11041 tatccacaga atcaggggat aacgcaggaa agaacatgtg agcaaaaggc cagcaaaagg
    11101 ccaggaaccg taaaaaggcc gcgttgctgg cgtttttcca taggctccgc ccccctgacg
    11161 agcatcacaa aaatcgacgc tcaagtcaga ggtggcgaaa cccgacagga ctataaagat
    11221 accaggcgtt tccccctgga agctccctcg tgcgctctcc tgttccgacc ctgccgctta
    11281 ccggatacct gtccgccttt ctcccttcgg gaagcgtggc gctttctcat agctcacgct
    11341 gtaggtatct cagttcggtg taggtcgttc gctccaagct gggctgtgtg cacgaacccc
    11401 ccgttcagcc cgaccgctgc gccttatccg gtaactatcg tcttgagtcc aacccggtaa
    11461 gacacgactt atcgccactg gcagcagcca ctggtaacag gattagcaga gcgaggtatg
    11521 taggcggtgc tacagagttc ttgaagtggt ggcctaacta cggctacact agaagaacag
    11581 tatttggtat ctgcgctctg ctgaagccag ttaccttcgg aaaaagagtt ggtagctctt
    11641 gatccggcaa acaaaccacc gctggtagcg gtggtttttt tgtttgcaag cagcagatta
    11701 cgcgcagaaa aaaaggatct caagaagatc ctttgatctt ttctacgggg tctgacgctc
    11761 agtggaacga aaactcacgt taagggattt tggtcatgag attatcaaaa aggatcttca
    11821 cctagatcct tttaaattaa aaatgaagtt ttaaatcaat ctaaagtata tatgagtaaa
    11881 cttggtctga cagttaccaa tgcttaatca gtgaggcacc tatctcagcg atctgtctat
    11941 ttcgttcatc catagttgcc tgactccccg tcgtgtagat aactacgata cgggagggct
    12001 taccatctgg ccccagtgct gcaatgatac cgcgagaccc acgctcaccg gctccagatt
    12061 tatcagcaat aaaccagcca gccggaaggg ccgagcgcag aagtggtcct gcaactttat
    12121 ccgcctccat ccagtctatt aattgttgcc gggaagctag agtaagtagt tcgccagtta
    12181 atagtttgcg caacgttgtt gccattgcta caggcatcgt ggtgtcacgc tcgtcgtttg
    12241 gtatggcttc attcagctcc ggttcccaac gatcaaggcg agttacatga tcccccatgt
    12301 tgtgcaaaaa agcggttagc tccttcggtc ctccgatcgt tgtcagaagt aagttggccg
    12361 cagtgttatc actcatggtt atggcagcac tgcataattc tcttactgtc atgccatccg
    12421 taagatgctt ttctgtgact ggtgagtact caaccaagtc attctgagaa tagtgtatgc
    12481 ggcgaccgag ttgctcttgc ccggcgtcaa tacgggataa taccgcgcca catagcagaa
    12541 ctttaaaagt gctcatcatt ggaaaacgtt cttcggggcg aaaactctca aggatcttac
    12601 cgctgttgag atccagttcg atgtaaccca ctcgtgcacc caactgatct tcagcatctt
    12661 ttactttcac cagcgtttct gggtgagcaa aaacaggaag gcaaaatgcc gcaaaaaagg
    12721 gaataagggc gacacggaaa tgttgaatac tcatactctt cctttttcaa tattattgaa
    12781 gcatttatca gggttattgt ctcatgagcg gatacatatt tgaatgtatt tagaaaaata
    12841 aacaaatagg ggttccgcgc acatttcccc gaaaagtgcc acctgacgtc taagaaacca
    12901 ttattatcat gacattaacc tataaaaata ggcgtatcac gaggcccttt cgtc
//

PiggyBac Linker Neoantigens BFP

LOCUS BH1.13_EF1a_47NeoAg_Linker_BFP               14574 bp DNA     circular SYN 24-OCT-2022
DEFINITION  synthetic circular DNA
ACCESSION   .
VERSION     .
KEYWORDS    .
SOURCE      synthetic DNA construct
  ORGANISM  synthetic DNA construct
REFERENCE   1  (bases 1 to 14574)
  AUTHORS   .
  TITLE     Direct Submission
  JOURNAL   Exported Oct 25, 2022 from SnapGene 6.0.5
            https://www.snapgene.com
FEATURES             Location/Qualifiers
     source          1..14574
                     /mol_type="other DNA"
                     /organism="synthetic DNA construct"
     CDS             complement(239..307)
                     /label=LacZ alpha
     primer_bind     378..395
                     /label=M13-fwd
     primer_bind     379..395
                     /label=M13 fwd
     protein_bind    complement(445..682)
                     /label=PB-TRE 874..1111
     enhancer        719..950
                     /label=Insulator 1148..1379
     enhancer        720..950
                     /label=Insulator 1701..1931
     misc_feature    1084..1085
                     /label=ClaI_SpeI fragment PCR clone
     promoter        1243..2424
                     /label=EF-1-alpha promoter
     intron          1473..2415
                     /label=EF-1-alpha intron A
     regulatory      2444..2449
                     /label=Kozak sequence
     CDS             2450..8476
                     /label=47nAg_YESLinkers
                     /label=48nAg_YESLinkers_all_controls_mmCO_ORF
     misc_feature    2450..2524
                     /label=KRAS G12D
     misc_feature    2555..2635
                     /label=JAK2 V617F
     misc_feature    2666..2743
                     /label=KRAS G13D
     misc_feature    2774..2854
                     /label=NRAS Q61R/HRAS Q61R
     misc_feature    2885..2965
                     /label=EGFR G719A
     misc_feature    3002..3082
                     /label=EGFR T790M
     misc_feature    3113..3214
                     /label=EGFR T790M_C797S
     misc_feature    3245..3319
                     /label=KRAS G12S
     misc_feature    3350..3430
                     /label=PIK3CA E542K
     misc_feature    3461..3541
                     /label=PIK3CA H1047L
     misc_feature    3578..3658
                     /label=ERBB2 S310F
     misc_feature    3689..3769
                     /label=BRAF V600M
     misc_feature    3689..3769
                     /label=GTF2I L424H
     misc_feature    3800..3880
                     /label=FGFR3 S249C
     misc_feature    3911..3991
                     /label=MYD88 L265P
     misc_feature    4022..4102
                     /label=PTEN R130Q
     misc_feature    4139..4219
                     /label=TP53 R248Q
     misc_feature    4250..4330
                     /label=TP53 R273H
     misc_feature    4361..4441
                     /label=TP53 R282W
     misc_feature    4472..4552
                     /label=ESR1 K303R
     misc_feature    4583..4663
                     /label=GNA11 Q209L
     misc_feature    4700..4780
                     /label=GNAQ Q209P
     misc_feature    4811..4891
                     /label=IDH1 R132C
     misc_feature    4922..5002
                     /label=IDH1 R132H
     misc_feature    5033..5113
                     /label=SF3B1 R625C
     misc_feature    5144..5218
                     /label=KRAS G12C
     misc_feature    5255..5329
                     /label=KRAS G12V
     misc_feature    5360..5440
                     /label=NRAS Q61K
     misc_feature    5471..5539
                     /label=EGFR E746_A750del
     misc_feature    5570..5650
                     /label=EGFR L858R
     misc_feature    5681..5761
                     /label=EGFR C797S
     misc_feature    5798..5872
                     /label=KRAS G12A
     misc_feature    5903..5980
                     /label=KRAS G13C
     misc_feature    6011..6091
                     /label=PIK3CA E545K
     misc_feature    6092..6172
                     /label=PIK3CA H1047R
     misc_feature    6173..6253
                     /label=BRAF V600E
     misc_feature    6284..6364
                     /label=DNMT3A R882H
     misc_feature    6395..6469
                     /label=KRAS G12R
     misc_feature    6506..6586
                     /label=PTEN R130G
     misc_feature    6617..6697
                     /label=TP53 R175H
     misc_feature    6728..6808
                     /label=TP53 R273C
     misc_feature    6839..6919
                     /label=TP53 R273L
     misc_feature    6950..7012
                     /label=CALR fs
     misc_feature    7049..7129
                     /label=FLT3 D835Y
     misc_feature    7160..7240
                     /label=GNAQ Q209L
     misc_feature    7271..7351
                     /label=BRAF V600M
     misc_feature    7271..7351
                     /label=GTF2I L424H
     misc_feature    7382..7462
                     /label=IDH1 R132G
     misc_feature    7493..7573
                     /label=IDH2 R140Q
     misc_feature    7610..7690
                     /label=SF3B1 R625H
     CDS             7721..7801
                     /codon_start=1
                     /label=pp65 495-503
                     /translation="PWQAGILARNLVPMVATVQGQNLKYQE"
     CDS             7838..7918
                     /codon_start=1
                     /label=BMLF1
                     /translation="ARMQAIQNAGLCTLVAMLEETIFWLQE"
     CDS             7949..8029
                     /codon_start=1
                     /label=Influenza M
                     /translation="RPILSPLTKGILGFVFTLTVPSERGLQ"
     CDS             8060..8140
                     /codon_start=1
                     /label=IE-1
                     /translation="EFCRVLCCYVLEETSVMLAKRPLITKP"
     CDS             8171..8251
                     /codon_start=1
                     /label=Mart1
                     /translation="GIGILTVADELAGIGILTVADILTVIL"
     CDS             8282..8362
                     /codon_start=1
                     /label=MAGE A3
                     /translation="PGSDPACYEFLWGPRALVETSYVKVLH"
     CDS             8393..8473
                     /codon_start=1
                     /label=NY-ESO1
                     /translation="SISSCLQQLSLLMWITQVFLPVFLAQP"
     misc_feature    8565..9116
                     /label=IRES
     CDS             9128..9838
                     /label=mTagBFP2
     misc_feature    9843..9847
                     /label=via pLENTI6.3_ClaI_NheI
     polyA_signal    9854..9902
                     /label=poly(A) signal
     misc_feature    9904..9933
                     /label=buffer sequence
     polyA_signal    9934..10141
                     /label=bGH poly(A) signal
     misc_feature    10147..10264
                     /label=cPPT/CTS
     promoter        10289..10789
                     /label=hPGK promoter
     CDS             10803..11399
                     /label=PuroR
     polyA_signal    11409..11530
                     /label=SV40 poly(A) signal
     enhancer        11627..11858
                     /label=Insulator 1148..1379
     enhancer        11627..11857
                     /label=Insulator 1701..1931
     primer_bind     complement(11897..11916)
                     /label=T3
     promoter        complement(11899..11917)
                     /label=T3 promoter
     primer_bind     11924..11943
                     /label=T7
     promoter        11924..11942
                     /label=T7 promoter
     protein_bind    11993..12301
                     /label=PB-TRE 2067..2375
     misc_feature    12335
                     /label=HindIII killer
     primer_bind     complement(12349..12369)
                     /label=M13-rev
     primer_bind     complement(12353..12369)
                     /label=M13 rev
     misc_binding    complement(12375..12397)
                     /label=LacO
     protein_bind    12377..12393
                     /label=lac operator
     promoter        complement(12401..12431)
                     /label=lac promoter
     promoter        complement(12401..12407)
                     /label=lac promoter
     promoter        complement(12408..12425)
                     /label=lac promoter
     promoter        complement(12426..12431)
                     /label=lac promoter
     protein_bind    12446..12467
                     /label=CAP binding site
     rep_origin      complement(12737..13419)
                     /direction=LEFT
                     /label=ColE1 origin
     rep_origin      complement(12755..13343)
                     /direction=LEFT
                     /label=ori
     CDS             complement(13514..14305)
                     /label=AmpR
     CDS             complement(13517..14176)
                     /label=AmpR
     CDS             complement(14306..14374)
                     /label=AmpR
     promoter        complement(14375..14479)
                     /label=AmpR promoter
ORIGIN
        1 tcgcgcgttt cggtgatgac ggtgaaaacc tctgacacat gcagctcccg gagacggtca
       61 cagcttgtct gtaagcggat gccgggagca gacaagcccg tcagggcgcg tcagcgggtg
      121 ttggcgggtg tcggggctgg cttaactatg cggcatcaga gcagattgta ctgagagtgc
      181 accatatgcg gtgtgaaata ccgcacagat gcgtaaggag aaaataccgc atcaggcgcc
      241 attcgccatt caggctgcgc aactgttggg aagggcgatc ggtgcgggcc tcttcgctat
      301 tacgccagct ggcgaaaggg ggatgtgctg caaggcgatt aagttgggta acgccagggt
      361 tttcccagtc acgacgttgt aaaacgacgg ccagtgaatt aggcgcgcct gctcgacacg
      421 ctgcagaaca cgcagctaga ttaaccctag aaagataatc atattgtgac gtacgttaaa
      481 gataatcatg cgtaaaattg acgcatgtgt tttatcggtc tgtatatcga ggtttattta
      541 ttaatttgaa tagatattaa gttttattat atttacactt acatactaat aataaattca
      601 acaaacaatt tatttatgtt tatttattta ttaaaaaaaa acaaaaactc aaaatttctt
      661 ctataaagta acaaaacttt tatcgaattg ctgcagcccg ggcgatccat tagtactaga
      721 gggacagccc ccccccaaag cccccaggga tgtaattacg tccctccccc gctagggggc
      781 agcagcgagc cgcccggggc tccgctccgg tccggcgctc cccccgcatc cccgagccgg
      841 cagcgtgcgg ggacagcccg ggcacgggga aggtggcacg ggatcgcttt cctctgaacg
      901 cttctcgctg ctctttgagc ctgcagacac ctggggggat acggggaaaa ggcctccaag
      961 gccagcttcc cacaataagt tgggtgaatt ttggctcatt cctcctttct ataggattga
     1021 ggtcagagct ttgtgatggc aattctgtgg aatgtgtgtc agttagggtg tggaaagtca
     1081 agcatcgatg agtaattcat acaaaaggac tcgcccctgc cttggggaat cccagggacc
     1141 gtcgttaaac tcccactaac gtagaaccca gagatcgctg cgttcccgcc ccctcacccg
     1201 cccgctctcg tcatcactga ggtggagaag agcatgcgtg aggctccggt gcccgtcagt
     1261 gggcagagcg cacatcgccc acagtccccg agaagttggg gggaggggtc ggcaattgaa
     1321 ccggtgccta gagaaggtgg cgcggggtaa actgggaaag tgatgtcgtg tactggctcc
     1381 gcctttttcc cgagggtggg ggagaaccgt atataagtgc agtagtcgcc gtgaacgttc
     1441 tttttcgcaa cgggtttgcc gccagaacac aggtaagtgc cgtgtgtggt tcccgcgggc
     1501 ctggcctctt tacgggttat ggcccttgcg tgccttgaat tacttccacg cccctggctg
     1561 cagtacgtga ttcttgatcc cgagcttcgg gttggaagtg ggtgggagag ttcgaggcct
     1621 tgcgcttaag gagccccttc gcctcgtgct tgagttgagg cctggcttgg gcgctggggc
     1681 cgccgcgtgc gaatctggtg gcaccttcgc gcctgtctcg ctgctttcga taagtctcta
     1741 gccatttaaa atttttgatg acctgctgcg acgctttttt tctggcaaga tagtcttgta
     1801 aatgcgggcc aagatctgca cactggtatt tcggtttttg gggccgcggg cggcgacggg
     1861 gcccgtgcgt cccagcgcac atgttcggcg aggcggggcc tgcgagcgcg gccaccgaga
     1921 atcggacggg ggtagtctca agctggccgg cctgctctgg tgcctggcct cgcgccgccg
     1981 tgtatcgccc cgccctgggc ggcaaggctg gcccggtcgg caccagttgc gtgagcggaa
     2041 agatggccgc ttcccggccc tgctgcaggg agctcaaaat ggaggacgcg gcgctcggga
     2101 gagcgggcgg gtgagtcacc cacacaaagg aaaagggcct ttccgtcctc agccgtcgct
     2161 tcatgtgact ccacggagta ccgggcgccg tccaggcacc tcgattagtt ctcaagcttt
     2221 tggagtacgt cgtctttagg ttggggggag gggttttatg cgatggagtt tccccacact
     2281 gagtgggtgg agactgaagt taggccagct tggcacttga tgtaattctc cttggaattt
     2341 gccctttttg agtttggatc ttggttcatt ctcaagcctc agacagtggt tcaaagtttt
     2401 tttcttccat ttcaggtgtc gtgaccctag cgctacctct agagccacca tgaccgagta
     2461 taaactggtg gtggttggtg cagatggggt ggggaaatca gctctcacta tccagcttat
     2521 acaaggttcc ggaagcggtt ctggaagtgg gtcccttagc cacaaacacc ttgttctgaa
     2581 ctatggcgtt tgtttttgtg gtgacgagaa cattctggtc caggaattcg taaaaggtgg
     2641 gtccagtggc ggtggtagtt caggaatgac cgagtacaag ctcgtcgttg tgggggctgg
     2701 agatgtcggc aagagcgcct tgactattca actcatccaa aacggctcag ccggtagtgc
     2761 cgcagggtcc ggtggcgaga cctgtctgct cgatatcctg gatactgcag gacgggaaga
     2821 atacagcgca atgcgagacc aatatatgcg gacaggaggc tctggaggag gcggatctgg
     2881 cggcattttg aaggaaaccg aattcaaaaa gatcaaagta cttgcaagtg gggcctttgg
     2941 cacagtctat aaaggtttgt ggattggctc tgcaggcagc gcagcaggga gtggtgagtt
     3001 tctgctcggc atctgtctta caagcacagt tcaacttatt atgcaactta tgccattcgg
     3061 ctgtctcctg gactacgtcc ggggctccgg aagcgggagc ggaagcggca gcttgcttgg
     3121 tatctgcctg acaagtactg ttcaactgat aatgcaactc atgccatttg gcagcttgtt
     3181 ggattacgta agagaacaca aagacaacat cgggggaggt tctagcggtg gagggtcctc
     3241 tgggatgact gagtataagc tcgtggttgt cggtgcatca ggcgtgggta agtcagcact
     3301 taccatacag cttatccagg gttcagccgg ttccgctgca ggatcaggtg agcaactcaa
     3361 ggcaatatct actagggatc ccctttccaa gataacagaa caggagaaag acttcctgtg
     3421 gtcccaccga ggtggctctg gcgggggcgg aagtggggga gaggcccttg agtattttat
     3481 gaaacagatg aatgatgcac ttcacggtgg ctggaccaca aagatggact ggatttttca
     3541 tgggtcagca ggctccgccg ccggatctgg tgaatttaca gcttgccctt ataactatct
     3601 cagtactgat gtgggatttt gcaccctggt ttgtccactt cataaccagg aggtcacagg
     3661 gagtggatca ggttccggaa gtggctcaga tctcactgtt aagatcggtg actttgggtt
     3721 ggccactatg aaatccagat ggagtggaag ccatcagttc gagcaactcg gggggtcaag
     3781 cggtggtgga tcctccggta gcattcggca gacttatact cttgacgtgc ttgagcggtg
     3841 cccacatcgg cccatccttc aggctgggct tcctgctaat gggtcagccg gctcagctgc
     3901 tggcagcggc ttcgccctga gtttgtcacc cggcgctcac caaaagagac ccatacctat
     3961 caaatataaa gctatgaaga aggagttccc tggtggatca ggcggcgggg ggtccggagg
     4021 caaccatgtt gccgcaatac actgcaaggc aggtaagggt caaactggag tgatgatttg
     4081 tgcctatctc ttgcatcgcg ggggctctgc aggaagcgct gccggcagtg gcgaatttaa
     4141 ctacatgtgt aactcctctt gtatgggcgg gatgaaccaa cggcctattc tcactattat
     4201 caccctggaa gattctagtg gtagtggttc cggatctggg agcggtagca gttccggaaa
     4261 cctgttgggc cgcaacagct tcgaagtgca tgtctgcgct tgtcctggtc gggaccgccg
     4321 gacagaagag ggtggatcat ctggtggtgg ctcaagcgga agcttcgaag tgcgagtctg
     4381 cgcttgccca ggacgagact ggagaaccga agaagaaaac cttcgaaaga aaggcgagcc
     4441 tggttctgcc ggcagtgctg ctggaagtgg gaatttgtgg cccagccccc tgatgataaa
     4501 acgatccaaa cgaaactcct tggctctgag tcttactgct gaccaaatgg taggggggtc
     4561 tgggggtgga gggtccggcg ggcttgaaaa tatcatattc cggatggtcg atgtcggggg
     4621 cttgagaagc gaacgccgca agtggataca ctgcttcgaa aacggctcag caggctctgc
     4681 tgcagggtca ggggagttct tgcaaagtgt gatctttaga atggtagatg taggaggtcc
     4741 ccgatcagaa cgcagaaaat ggatacactg ttttgaaaat ggctctggta gcggaagtgg
     4801 gagcggaagc cgacttgttt ccggatgggt taagcccatc atcattggtt gtcacgctta
     4861 cggcgatcaa taccgggcaa ccgattttgt agggggatca tccggaggag gtagctcagg
     4921 gcggctcgtg agtggatggg tgaaacccat aatcatcggt caccacgcct atggtgacca
     4981 atatcgggct actgactttg ttggttctgc cggtagtgcc gcaggttcag ggactatgag
     5041 gcccgacatc gacaatatgg atgaatacgt gtgtaataca accgcaagag cattcgcagt
     5101 agttgccagc gctggtggat caggtggagg ggggtcagga ggcatgaccg agtacaagtt
     5161 ggtagtagta ggagcttgcg gcgtgggtaa aagcgccctg actattcagc tcatacaggg
     5221 atctgcaggc tctgctgctg ggagtgggga atttatgacc gagtataaac tcgtagtggt
     5281 aggagcagta ggggttggaa aatctgctct gactattcag cttatccagg ggtctgggtc
     5341 agggtctggg agcggtagtg gtgaaacttg tcttctcgat attcttgata ctgctggtaa
     5401 ggaagagtac tccgcaatga gggaccaata tatgcgaacc ggcggctcat ccggtggtgg
     5461 ttcttcaggg cccgagggag aaaaagtaaa aattcctgtc gctatcaaaa cctcccctaa
     5521 ggctaacaaa gagattcttg gatctgctgg ctccgcagca ggctcaggtg tcaaaactcc
     5581 ccaacatgtt aagatcaccg actttggccg ggctaagttg ttgggggcag aagagaagga
     5641 ataccacgcc gggggctctg gaggaggggg cagcggggga agtacagttc aattgatcac
     5701 acagttgatg ccatttggct ccctgctcga ttatgtgcgg gagcacaaag acaacatcgg
     5761 gggtagcgcc ggttctgcag caggttcagg tgagttcatg acagagtaca aattggtagt
     5821 ggtgggagcc gctggagtag gcaaatccgc actcactatc cagttgattc agggatccgg
     5881 ttccggaagc gggtcaggat caatgaccga atataaactg gtggtcgtcg gtgctgggtg
     5941 tgttggtaaa agcgctctga ccatacaact tatacagaac ggcgggagta gcgggggtgg
     6001 ttcaagcggg aaggccatta gtacccgaga ccccctttca gaaatcacaa aacaggaaaa
     6061 agactttctg tggagccatc gccattattg tgaggctttg gaatatttta tgaagcagat
     6121 gaacgacgcc cgccacggtg gatggactac caaaatggat tggatattcc atgacctcac
     6181 cgtaaaaatt ggcgattttg gcctcgcaac tgagaaaagc cgatggagcg gttctcatca
     6241 gttcgagcag ctcggttccg cagggtcagc tgctggctca gggggatttc ccgtgcatta
     6301 tactgatgtt agcaacatgt cacatcttgc taggcagaga ctccttggtc gcagctggtc
     6361 tgtgggggga tctggcggtg gtggatctgg ggggatgact gagtacaagc tcgtggtggt
     6421 aggagcccga ggagtcggca agtctgcact cactattcaa ctgattcagg ggtccgcagg
     6481 atccgctgcc ggcagcgggg aatttaatca cgtcgctgcc attcactgta aagcaggcaa
     6541 aggtggaaca ggcgttatga tatgtgcata ccttctgcac cgcgggggga gcggttctgg
     6601 ttctggaagt ggaagcattt acaaacaaag tcaacatatg actgaagtag tgcgacactg
     6661 tccacatcat gaacgctgct ctgatagtga tgggcttggg ggatcttctg ggggcggctc
     6721 ctcaggaagt tccgggaacc tgctgggtcg caatagcttt gaagtgtgtg tttgtgcatg
     6781 tccagggcgc gaccgacgca ccgaagaagg ctccgccggt tcagccgctg gatctggatc
     6841 tagcggaaat ctgcttggga ggaacagctt tgaggtactc gtctgcgcct gtcctggacg
     6901 cgatagacgc actgaggagg gcgggtctgg tggaggcgga tccgggggga gaatgcggag
     6961 gatgagacgc actagaagga aaatgcggcg aaaaatgagc ccagctcgac caggtagtgc
     7021 tggaagtgct gctgggtcag gggagtttgg taaagtagtt aagatctgtg actttgggct
     7081 ggcacgctat attatgagcg attctaatta tgtagtcaga ggcaatgcag ggtccggcag
     7141 cgggagtggt agtggttcat tgcaatctgt aatttttaga atggtagatg taggcggatt
     7201 gagaagcgag agacgaaaat ggattcactg tttcgagaat ggggggtcaa gtggaggggg
     7261 ctcttcaggc gatctgaccg tcaagattgg cgattttggt ttggccacaa tgaaaagtcg
     7321 gtggagcggg tcacatcagt ttgagcagct tggttctgct ggaagcgccg ccggttcagg
     7381 tagactcgtg agcgggtggg tcaagcccat cataataggg ggccatgctt acggcgatca
     7441 atacagagct acagacttcg tgggggggtc aggaggaggt ggttccggtg gtaaactcaa
     7501 gaagatgtgg aaaagcccaa acggaacaat tcaaaatata ctggggggca ctgttttcag
     7561 ggagccaatt attggttccg ccgggtcagc cgctgggtca ggggaattca ccatgaggcc
     7621 tgacatagac aatatggacg agtatgtcca caataccaca gcacgagctt tcgccgtagt
     7681 agcatctgct ggcggctccg gtggaggagg atcaggaggc ccatggcaag ctgggattct
     7741 cgcaagaaac cttgtcccta tggtagcaac agtgcagggg cagaacctta agtaccaaga
     7801 aggaagcgct ggtagcgctg cagggagcgg cgaattcgct cggatgcaag ccatacaaaa
     7861 tgccggcttg tgtacccttg ttgcaatgct ggaagagact atattctggt tgcaggaagg
     7921 gtccggatca gggtctgggt ctggctccag gcccattctg agcccactca caaagggtat
     7981 tctgggcttt gtatttacat tgactgtccc cagcgaaaga ggcctccaag gcggttcttc
     8041 tggaggaggg tcctctggcg agttttgccg ggtactgtgt tgctatgtgc tcgaagagac
     8101 cagcgtcatg ttggctaaaa ggcctttgat aaccaaacca ggaggttcta gtgggggagg
     8161 gagctcaggt ggtataggta tccttacagt cgccgacgaa ctcgctggta taggtatatt
     8221 gactgtggcc gacatcctga ccgtgatcct gggatcagct gggtcagccg ctggttcagg
     8281 gcccggaagt gaccccgcct gctatgaatt cctttggggg ccacgcgccc ttgtagaaac
     8341 atcttacgtt aaggtactcc atggcgggtc cggaggtggc ggaagcggag gatcaattag
     8401 cagctgtctc cagcagttgt ccttgcttat gtggataacc caggtatttt tgcccgtctt
     8461 cttggcccag ccctgactcg aggttaaccg cccgccccac gacccgcagc gcccgaccga
     8521 aaggagcgca cgaccccatc atccaattcc gccccccccc cctaacgtta ctggccgaag
     8581 ccgcttggaa taaggccggt gtgcgtttgt ctatatgtta ttttccacca tattgccgtc
     8641 ttttggcaat gtgagggccc ggaaacctgg ccctgtcttc ttgacgagca ttcctagggg
     8701 tctttcccct ctcgccaaag gaatgcaagg tctgttgaat gtcgtgaagg aagcagttcc
     8761 tctggaagct tcttgaagac aaacaacgtc tgtagcgacc ctttgcaggc agcggaaccc
     8821 cccacctggc gacaggtgcc tctgcggcca aaagccacgt gtataagata cacctgcaaa
     8881 ggcggcacaa ccccagtgcc acgttgtgag ttggatagtt gtggaaagag tcaaatggct
     8941 ctcctcaagc gtattcaaca aggggctgaa ggatgcccag aaggtacccc attgtatggg
     9001 atctgatctg gggcctcggt gcacatgctt tacatgtgtt tagtcgaggt taaaaaacgt
     9061 ctaggccccc cgaaccacgg ggacgtggtt ttcctttgaa aaacacgatg ataatatggc
     9121 cacaaccatg gtgtctaagg gcgaagagct gattaaggag aacatgcaca tgaagctgta
     9181 catggagggc accgtggaca accatcactt caagtgcaca tccgagggcg aaggcaagcc
     9241 ctacgagggc acccagacca tgagaatcaa ggtggtcgag ggcggccctc tccccttcgc
     9301 cttcgacatc ctggctacta gcttcctcta cggcagcaag accttcatca accacaccca
     9361 gggcatcccc gacttcttca agcagtcctt ccctgagggc ttcacatggg agagagtcac
     9421 cacatacgaa gacgggggcg tgctgaccgc tacccaggac accagcctcc aggacggctg
     9481 cctcatctac aacgtcaaga tcagaggggt gaacttcaca tccaacggcc ctgtgatgca
     9541 gaagaaaaca ctcggctggg aggccttcac cgagacgctg taccccgctg acggcggcct
     9601 ggaaggcaga aacgacatgg ccctgaagct cgtgggcggg agccatctga tcgcaaacgc
     9661 caagaccaca tatagatcca agaaacccgc taagaacctc aagatgcctg gcgtctacta
     9721 tgtggactac agactggaaa gaatcaagga ggccaacaac gagacctacg tcgagcagca
     9781 cgaggtggca gtggccagat actgcgacct ccctagcaaa ctggggcaca agcttaattg
     9841 agctagcgtc gacaataaaa tatctttatt ttcattacat ctgtgtgttg gttttttgtg
     9901 tgtggcacct cctgccatct gtctttccgt ttcctgtgcc ttctagttgc cagccatctg
     9961 ttgtttgccc ctcccccgtg ccttccttga ccctggaagg tgccactccc actgtccttt
    10021 cctaataaaa tgaggaaatt gcatcgcatt gtctgagtag gtgtcattct attctggggg
    10081 gtggggtggg gcaggacagc aagggggagg attgggaaga gaatagcagg catgctgggg
    10141 aaattgtttt aaaagaaaag gggggattgg ggggtacagt gcaggggaaa gaatagtaga
    10201 cataatagca acagacatac aaactaaaga attacaaaaa caaattacaa aaattcaaaa
    10261 ttttacccca gaacgcgtgg ggcccgggtt gcgccttttc caaggcagcc ctgggtttgc
    10321 gcagggacgc ggctgctctg ggcgtggttc cgggaaacgc agcggcgccg accctgggtc
    10381 tcgcacattc ttcacgtccg ttcgcagcgt cacccggatc ttcgccgcta cccttgtggg
    10441 ccccccggcg acgcttcctg ctccgcccct aagtcgggaa ggttccttgc ggttcgcggc
    10501 gtgccggacg tgacaaacgg aagccgcacg tctcactagt accctcgcag acggacagcg
    10561 ccagggagca atggcagcgc gccgaccgcg atgggctgtg gccaatagcg gctgctcagc
    10621 agggcgcgcc gagagcagcg gccgggaagg ggcggtgcgg gaggcggggt gtggggcggt
    10681 agtgtgggcc ctgttcctgc ccgcgcggtg ttccgcattc tgcaagcctc cggagcgcac
    10741 gtcggcagtc ggctccctcg ttgaccgaat caccgacctc tctccccagg gagatctcaa
    10801 ccatgaccga gtacaagccc acggtgcgcc tcgccacccg cgacgacgtc ccccgggccg
    10861 tacgcaccct cgccgccgcg ttcgccgact accccgccac gcgccacacc gtggacccgg
    10921 accgccacat cgagcgggtc accgagctgc aagaactctt cctcacgcgc gtcgggctcg
    10981 acatcggcaa ggtgtgggtc gcggacgacg gcgccgcggt ggcggtctgg accacgccgg
    11041 agagcgtcga agcgggggcg gtgttcgccg agatcggccc gcgcatggcc gagttgagcg
    11101 gttcccggct ggccgcgcag caacagatgg aaggcctcct ggcgccgcac cggcccaagg
    11161 agcccgcgtg gttcctggcc accgtcggcg tctcgcccga ccaccagggc aagggtctgg
    11221 gcagcgccgt cgtgctcccc ggagtggagg cggccgagcg cgccggggtg cccgccttcc
    11281 tggagacctc cgcgccccgc aacctcccct tctacgagcg gctcggcttc accgtcaccg
    11341 ccgacgtcga ggtgcccgaa ggaccgcgca cctggtgcat gacccgcaag cccggtgcct
    11401 gagctagcaa cttgtttatt gcagcttata atggttacaa ataaagcaat agcatcacaa
    11461 atttcacaaa taaagcattt ttttcactgc attctagttg tggtttgtcc aaactcatca
    11521 atgtatctta tgtcctcaca ggaacgaagt ccctaaagaa acagtggcag ccaggtttag
    11581 ccccggaatt gactggattc cttttttagg gcccattggt atggcttttt ccccgtatcc
    11641 ccccaggtgt ctgcaggctc aaagagcagc gagaagcgtt cagaggaaag cgatcccgtg
    11701 ccaccttccc cgtgcccggg ctgtccccgc acgctgccgg ctcggggatg cggggggagc
    11761 gccggaccgg agcggagccc cgggcggctc gctgctgccc cctagcgggg gagggacgta
    11821 attacatccc tgggggcttt gggggggggc tgtccctctt gaacgaccgc caccgcggtg
    11881 gagctccagc ttttgttccc tttagtgagg gttaattaaa tcttaatacg actcactata
    11941 gggcgaattg ggaaccgggc cccccctgga gatcgacggt atccataagc ttgatatcta
    12001 taacaagaaa atatatatat aataagttat cacgtaagta gaacatgaaa taacaatata
    12061 attatcgtat gagttaaatc ttaaaagtca cgtaaaagat aatcatgcgt cattttgact
    12121 cacgcggtcg ttatagttca aaatcagtga cacttaccgc attgacaagc acgcctcacg
    12181 ggagctccaa gcggcgactg agatgtccta aatgcacagc gacggattcg cgctatttag
    12241 aaagagagag caatatttca agaatgcatg cgtcaatttt acgcagacta tctttctagg
    12301 gttaatctag ctgcatcagg atcatagcgg ccgccagctt ggcgtaatca tggtcatagc
    12361 tgtttcctgt gtgaaattgt tatccgctca caattccaca caacatacga gccggaagca
    12421 taaagtgtaa agcctggggt gcctaatgag tgagctaact cacattaatt gcgttgcgct
    12481 cactgcccgc tttccagtcg ggaaacctgt cgtgccagct gcattaatga atcggccaac
    12541 gcgcggggag aggcggtttg cgtattgggc gctcttccgc ttcctcgctc actgactcgc
    12601 tgcgctcggt cgttcggctg cggcgagcgg tatcagctca ctcaaaggcg gtaatacggt
    12661 tatccacaga atcaggggat aacgcaggaa agaacatgtg agcaaaaggc cagcaaaagg
    12721 ccaggaaccg taaaaaggcc gcgttgctgg cgtttttcca taggctccgc ccccctgacg
    12781 agcatcacaa aaatcgacgc tcaagtcaga ggtggcgaaa cccgacagga ctataaagat
    12841 accaggcgtt tccccctgga agctccctcg tgcgctctcc tgttccgacc ctgccgctta
    12901 ccggatacct gtccgccttt ctcccttcgg gaagcgtggc gctttctcat agctcacgct
    12961 gtaggtatct cagttcggtg taggtcgttc gctccaagct gggctgtgtg cacgaacccc
    13021 ccgttcagcc cgaccgctgc gccttatccg gtaactatcg tcttgagtcc aacccggtaa
    13081 gacacgactt atcgccactg gcagcagcca ctggtaacag gattagcaga gcgaggtatg
    13141 taggcggtgc tacagagttc ttgaagtggt ggcctaacta cggctacact agaagaacag
    13201 tatttggtat ctgcgctctg ctgaagccag ttaccttcgg aaaaagagtt ggtagctctt
    13261 gatccggcaa acaaaccacc gctggtagcg gtggtttttt tgtttgcaag cagcagatta
    13321 cgcgcagaaa aaaaggatct caagaagatc ctttgatctt ttctacgggg tctgacgctc
    13381 agtggaacga aaactcacgt taagggattt tggtcatgag attatcaaaa aggatcttca
    13441 cctagatcct tttaaattaa aaatgaagtt ttaaatcaat ctaaagtata tatgagtaaa
    13501 cttggtctga cagttaccaa tgcttaatca gtgaggcacc tatctcagcg atctgtctat
    13561 ttcgttcatc catagttgcc tgactccccg tcgtgtagat aactacgata cgggagggct
    13621 taccatctgg ccccagtgct gcaatgatac cgcgagaccc acgctcaccg gctccagatt
    13681 tatcagcaat aaaccagcca gccggaaggg ccgagcgcag aagtggtcct gcaactttat
    13741 ccgcctccat ccagtctatt aattgttgcc gggaagctag agtaagtagt tcgccagtta
    13801 atagtttgcg caacgttgtt gccattgcta caggcatcgt ggtgtcacgc tcgtcgtttg
    13861 gtatggcttc attcagctcc ggttcccaac gatcaaggcg agttacatga tcccccatgt
    13921 tgtgcaaaaa agcggttagc tccttcggtc ctccgatcgt tgtcagaagt aagttggccg
    13981 cagtgttatc actcatggtt atggcagcac tgcataattc tcttactgtc atgccatccg
    14041 taagatgctt ttctgtgact ggtgagtact caaccaagtc attctgagaa tagtgtatgc
    14101 ggcgaccgag ttgctcttgc ccggcgtcaa tacgggataa taccgcgcca catagcagaa
    14161 ctttaaaagt gctcatcatt ggaaaacgtt cttcggggcg aaaactctca aggatcttac
    14221 cgctgttgag atccagttcg atgtaaccca ctcgtgcacc caactgatct tcagcatctt
    14281 ttactttcac cagcgtttct gggtgagcaa aaacaggaag gcaaaatgcc gcaaaaaagg
    14341 gaataagggc gacacggaaa tgttgaatac tcatactctt cctttttcaa tattattgaa
    14401 gcatttatca gggttattgt ctcatgagcg gatacatatt tgaatgtatt tagaaaaata
    14461 aacaaatagg ggttccgcgc acatttcccc gaaaagtgcc acctgacgtc taagaaacca
    14521 ttattatcat gacattaacc tataaaaata ggcgtatcac gaggcccttt cgtc
//

PiggyBac TRE3G_KrasG12C_mCherry

LOCUS BH1.6_TRE3G_KrasG12C_mCherry                8564 bp DNA     circular SYN 24-OCT-2022
DEFINITION  synthetic circular DNA
ACCESSION   .
VERSION     .
KEYWORDS    .
SOURCE      synthetic DNA construct
  ORGANISM  synthetic DNA construct
REFERENCE   1  (bases 1 to 8564)
  AUTHORS   .
  TITLE     Direct Submission
  JOURNAL   Exported Oct 25, 2022 from SnapGene 6.0.5
            https://www.snapgene.com
COMMENT     U179MFC040-45
FEATURES             Location/Qualifiers
     source          1..8564
                     /mol_type="other DNA"
                     /organism="synthetic DNA construct"
     primer_bind     379..395
                     /label=M13 fwd
     protein_bind    complement(445..682)
                     /label=PB-TRE 874..1111
     enhancer        719..950
                     /label=Insulator 1148..1379
     promoter        1090..1468
                     /label=TRE3G promoter
     protein_bind    1098..1116
                     /label=tet operator
     protein_bind    1134..1152
                     /label=tet operator
     protein_bind    1170..1188
                     /label=tet operator
     protein_bind    1206..1224
                     /label=tet operator
     protein_bind    1242..1260
                     /label=tet operator
     protein_bind    1278..1296
                     /label=tet operator
     protein_bind    1314..1332
                     /label=tet operator
     CDS             1475..2044
                     /codon_start=1
                     /label=KRAS G12C
                     /translation="MTEYKLVVVGACGVGKSALTIQLIQNHFVDEYDPTIEDSYRKQVV
                     IDGETCLLDILDTAGQEEYSAMRDQYMRTGEGFLCVFAINNTKSFEDIHHYREQIKRVK
                     DSEDVPMVLVGNKCDLPSRTVDTKQAQDLARSYGIPFIETSAKTRQRVEDAFYTLVREI
                     RQYRLKKISKEEKTPGCVKIKKCIIM"
     misc_feature    2127..2549
                     /label=IRES
     CDS             2690..3400
                     /label=mCherry
     polyA_signal    3407..3455
                     /label=poly(A) signal
     promoter        3472..3972
                     /label=hPGK promoter
     CDS             3986..4582
                     /label=PuroR
     CDS             4592..4645
                     /label=T2A
     CDS             4646..5392
                     /label=Tet-On(R) 3G
     polyA_signal    5399..5520
                     /label=SV40 poly(A) signal
     enhancer        5617..5848
                     /label=Insulator 1148..1379
     promoter        complement(5889..5907)
                     /label=T3 promoter
     promoter        5914..5932
                     /label=T7 promoter
     protein_bind    5983..6291
                     /label=PB-TRE 2067..2375
     primer_bind     complement(6343..6359)
                     /label=M13 rev
     protein_bind    6367..6383
                     /label=lac operator
     promoter        complement(6398..6415)
                     /label=lac promoter
     protein_bind    6436..6457
                     /label=CAP binding site
     rep_origin      complement(6745..7333)
                     /direction=LEFT
                     /label=ori
     CDS             complement(7504..8295)
                     /label=AmpR
     CDS             complement(8296..8364)
                     /label=AmpR
     promoter        complement(8365..8469)
                     /label=AmpR promoter
ORIGIN
        1 tcgcgcgttt cggtgatgac ggtgaaaacc tctgacacat gcagctcccg gagacggtca
       61 cagcttgtct gtaagcggat gccgggagca gacaagcccg tcagggcgcg tcagcgggtg
      121 ttggcgggtg tcggggctgg cttaactatg cggcatcaga gcagattgta ctgagagtgc
      181 accatatgcg gtgtgaaata ccgcacagat gcgtaaggag aaaataccgc atcaggcgcc
      241 attcgccatt caggctgcgc aactgttggg aagggcgatc ggtgcgggcc tcttcgctat
      301 tacgccagct ggcgaaaggg ggatgtgctg caaggcgatt aagttgggta acgccagggt
      361 tttcccagtc acgacgttgt aaaacgacgg ccagtgaatt aggcgcgcct gctcgacacg
      421 ctgcagaaca cgcagctaga ttaaccctag aaagataatc atattgtgac gtacgttaaa
      481 gataatcatg cgtaaaattg acgcatgtgt tttatcggtc tgtatatcga ggtttattta
      541 ttaatttgaa tagatattaa gttttattat atttacactt acatactaat aataaattca
      601 acaaacaatt tatttatgtt tatttattta ttaaaaaaaa acaaaaactc aaaatttctt
      661 ctataaagta acaaaacttt tatcgaattg ctgcagcccg ggcgatccat tagtactaga
      721 gggacagccc ccccccaaag cccccaggga tgtaattacg tccctccccc gctagggggc
      781 agcagcgagc cgcccggggc tccgctccgg tccggcgctc cccccgcatc cccgagccgg
      841 cagcgtgcgg ggacagcccg ggcacgggga aggtggcacg ggatcgcttt cctctgaacg
      901 cttctcgctg ctctttgagc ctgcagacac ctggggggat acggggaaaa ggcctccaag
      961 gccagcttcc cacaataagt tgggtgaatt ttggctcatt cctcctttct ataggattga
     1021 ggtcagagct ttgtgatggc aattctgtgg aatgtgtgtc agttagggtg tggaaagtca
     1081 agcatcgatg agtttactcc ctatcagtga tagagaacgt atgaagagtt tactccctat
     1141 cagtgataga gaacgtatgc agactttact ccctatcagt gatagagaac gtataaggag
     1201 tttactccct atcagtgata gagaacgtat gaccagttta ctccctatca gtgatagaga
     1261 acgtatctac agtttactcc ctatcagtga tagagaacgt atatccagtt tactccctat
     1321 cagtgataga gaacgtataa gctttaggcg tgtacggtgg gcgcctataa aagcagagct
     1381 cgtttagtga accgtcagat cgcctggagc aattccacaa cacttttgtc ttataccaac
     1441 tttccgtacc acttcctacc ctcgtaaaac cggtatgact gaatataaac ttgtggtagt
     1501 tggagcttgt ggcgtaggca agagtgcctt gacgatacag ctaattcaga atcattttgt
     1561 ggacgaatat gatccaacaa tagaggattc ctacaggaag caagtagtaa ttgatggaga
     1621 aacctgtctc ttggatattc tcgacacagc aggtcaagag gagtacagtg caatgaggga
     1681 ccagtacatg aggactgggg agggctttct ttgtgtattt gccataaata atactaaatc
     1741 atttgaagat attcaccatt atagagaaca aattaaaaga gttaaggact ctgaagatgt
     1801 acctatggtc ctagtaggaa ataaatgtga tttgccttct agaacagtag acacaaaaca
     1861 ggctcaggac ttagcaagaa gttatggaat tccttttatt gaaacatcag caaagacaag
     1921 acagagagtg gaggatgctt tttatacatt ggtgagagag atccgacaat acagattgaa
     1981 aaaaatcagc aaagaagaaa agactcctgg ctgtgtgaaa attaaaaaat gcattataat
     2041 gtaagttaac cgcccgcccc acgacccgca gcgcccgacc gaaaggagcg cacgacccca
     2101 tcatccaatt ccgccccccc cccctaacgt tactggccga agccgcttgg aataaggccg
     2161 gtgtgcgttt gtctatatgt tattttccac catattgccg tcttttggca atgtgagggc
     2221 ccggaaacct ggccctgtct tcttgacgag cattcctagg ggtctttccc ctctcgccaa
     2281 aggaatgcaa ggtctgttga atgtcgtgaa ggaagcagtt cctctggaag cttcttgaag
     2341 acaaacaacg tctgtagcga ccctttgcag gcagcggaac cccccacctg gcgacaggtg
     2401 cctctgcggc caaaagccac gtgtataaga tacacctgca aaggcggcac aaccccagtg
     2461 ccacgttgtg agttggatag ttgtggaaag agtcaaatgg ctctcctcaa gcgtattcaa
     2521 caaggggctg aaggatgccc agaaggtacc ccattgtatg ggatctgatc tggggcctcg
     2581 gtgcacatgc tttacatgtg tttagtcgag gttaaaaaac gtctaggccc cccgaaccac
     2641 ggggacgtgg ttttcctttg aaaaacacga tgataatatg gccacaacca tggtgagcaa
     2701 gggcgaggag gataacatgg ccatcatcaa ggagttcatg cgcttcaagg tgcacatgga
     2761 gggctccgtg aacggccacg agttcgagat cgagggcgag ggcgagggcc gcccctacga
     2821 gggcacccag accgccaagc tgaaggtgac caagggtggc cccctgccct tcgcctggga
     2881 catcctgtcc cctcagttca tgtacggctc caaggcctac gtgaagcacc ccgccgacat
     2941 ccccgactac ttgaagctgt ccttccccga gggcttcaag tgggagcgcg tgatgaactt
     3001 cgaggacggc ggcgtggtga ccgtgaccca ggactcctcc ctgcaggacg gcgagttcat
     3061 ctacaaggtg aagctgcgcg gcaccaactt cccctccgac ggccccgtaa tgcagaagaa
     3121 gaccatgggc tgggaggcct cctccgagcg gatgtacccc gaggacggcg ccctgaaggg
     3181 cgagatcaag cagaggctga agctgaagga cggcggccac tacgacgctg aggtcaagac
     3241 cacctacaag gccaagaagc ccgtgcagct gcccggcgcc tacaacgtca acatcaagtt
     3301 ggacatcacc tcccacaacg aggactacac catcgtggaa cagtacgaac gcgccgaggg
     3361 ccgccactcc accggcggca tggacgagct gtacaagtag gtcgacaata aaatatcttt
     3421 attttcatta catctgtgtg ttggtttttt gtgtgacgcg tggggcccgg gttgcgcctt
     3481 ttccaaggca gccctgggtt tgcgcaggga cgcggctgct ctgggcgtgg ttccgggaaa
     3541 cgcagcggcg ccgaccctgg gtctcgcaca ttcttcacgt ccgttcgcag cgtcacccgg
     3601 atcttcgccg ctacccttgt gggccccccg gcgacgcttc ctgctccgcc cctaagtcgg
     3661 gaaggttcct tgcggttcgc ggcgtgccgg acgtgacaaa cggaagccgc acgtctcact
     3721 agtaccctcg cagacggaca gcgccaggga gcaatggcag cgcgccgacc gcgatgggct
     3781 gtggccaata gcggctgctc agcagggcgc gccgagagca gcggccggga aggggcggtg
     3841 cgggaggcgg ggtgtggggc ggtagtgtgg gccctgttcc tgcccgcgcg gtgttccgca
     3901 ttctgcaagc ctccggagcg cacgtcggca gtcggctccc tcgttgaccg aatcaccgac
     3961 ctctctcccc agggagatct caaccatgac cgagtacaag cccacggtgc gcctcgccac
     4021 ccgcgacgac gtccccaggg ccgtacgcac cctcgccgcc gcgttcgccg actaccccgc
     4081 cacgcgccac accgtcgatc cggaccgcca catcgagcgg gtcaccgagc tgcaagaact
     4141 cttcctcacg cgcgtcgggc tcgacatcgg caaggtgtgg gtcgcggacg acggcgccgc
     4201 cgtggcggtc tggaccacgc cggagagcgt cgaagcgggg gcggtgttcg ccgagatcgg
     4261 cccgcgcatg gccgagttga gcggttcccg gctggccgcg cagcaacaga tggaaggcct
     4321 cctggcgccg caccggccca aggagcccgc gtggttcctg gccaccgtcg gcgtctcgcc
     4381 cgaccaccag ggcaagggtc tgggcagcgc cgtcgtgctc cccggagtgg aggcggccga
     4441 gcgcgccggg gtgcccgcct tcctggagac ctccgcgccc cgcaacctcc ccttctacga
     4501 gcggctcggc ttcaccgtca ccgccgacgt cgaggtgccc gaaggaccgc gcacctggtg
     4561 catgacccgc aagcccggtg ccggaagcgg agaaggtaga ggttctctcc tcacttgtgg
     4621 tgatgttgaa gaaaaccctg gtccaatgtc tagactggac aagagcaaag tcataaactc
     4681 tgctctggaa ttactcaatg gagtcggtat cgaaggcctg acgacaagga aactcgctca
     4741 aaagctggga gttgagcagc ctaccctgta ctggcacgtg aagaacaagc gggccctgct
     4801 cgatgccctg ccaatcgaga tgctggacag gcatcatacc cactcctgcc ccctggaagg
     4861 cgagtcatgg caagactttc tgcggaacaa cgccaagtca taccgctgtg ctctcctctc
     4921 acatcgcgac ggggctaaag tgcatctcgg cacccgccca acagagaaac agtacgaaac
     4981 cctggaaaat cagctcgcgt tcctgtgtca gcaaggcttc tccctggaga acgcactgta
     5041 cgctctgtcc gccgtgggcc actttacact gggctgcgta ttggaggaac aggagcatca
     5101 agtagcaaaa gaggaaagag agacacctac caccgattct atgcccccac ttctgaaaca
     5161 agcaattgag ctgttcgacc ggcagggagc cgaacctgcc ttccttttcg gcctggaact
     5221 aatcatatgt ggcctggaga aacagctaaa gtgcgaaagc ggcgggccga ccgacgccct
     5281 tgacgatttt gacttagaca tgctcccagc cgatgccctt gacgactttg accttgatat
     5341 gctgcctgct gacgctcttg acgattttga ccttgacatg ctccccgggt aagctagcaa
     5401 cttgtttatt gcagcttata atggttacaa ataaagcaat agcatcacaa atttcacaaa
     5461 taaagcattt ttttcactgc attctagttg tggtttgtcc aaactcatca atgtatctta
     5521 tgtcctcaca ggaacgaagt ccctaaagaa acagtggcag ccaggtttag ccccggaatt
     5581 gactggattc cttttttagg gcccattggt atggcttttt ccccgtatcc ccccaggtgt
     5641 ctgcaggctc aaagagcagc gagaagcgtt cagaggaaag cgatcccgtg ccaccttccc
     5701 cgtgcccggg ctgtccccgc acgctgccgg ctcggggatg cggggggagc gccggaccgg
     5761 agcggagccc cgggcggctc gctgctgccc cctagcgggg gagggacgta attacatccc
     5821 tgggggcttt gggggggggc tgtccctctt gaacgaccgc caccgcggtg gagctccagc
     5881 ttttgttccc tttagtgagg gttaattaaa tcttaatacg actcactata gggcgaattg
     5941 ggaaccgggc cccccctgga gatcgacggt atccataagc ttgatatcta taacaagaaa
     6001 atatatatat aataagttat cacgtaagta gaacatgaaa taacaatata attatcgtat
     6061 gagttaaatc ttaaaagtca cgtaaaagat aatcatgcgt cattttgact cacgcggtcg
     6121 ttatagttca aaatcagtga cacttaccgc attgacaagc acgcctcacg ggagctccaa
     6181 gcggcgactg agatgtccta aatgcacagc gacggattcg cgctatttag aaagagagag
     6241 caatatttca agaatgcatg cgtcaatttt acgcagacta tctttctagg gttaatctag
     6301 ctgcatcagg atcatagcgg ccgccagctt ggcgtaatca tggtcatagc tgtttcctgt
     6361 gtgaaattgt tatccgctca caattccaca caacatacga gccggaagca taaagtgtaa
     6421 agcctggggt gcctaatgag tgagctaact cacattaatt gcgttgcgct cactgcccgc
     6481 tttccagtcg ggaaacctgt cgtgccagct gcattaatga atcggccaac gcgcggggag
     6541 aggcggtttg cgtattgggc gctcttccgc ttcctcgctc actgactcgc tgcgctcggt
     6601 cgttcggctg cggcgagcgg tatcagctca ctcaaaggcg gtaatacggt tatccacaga
     6661 atcaggggat aacgcaggaa agaacatgtg agcaaaaggc cagcaaaagg ccaggaaccg
     6721 taaaaaggcc gcgttgctgg cgtttttcca taggctccgc ccccctgacg agcatcacaa
     6781 aaatcgacgc tcaagtcaga ggtggcgaaa cccgacagga ctataaagat accaggcgtt
     6841 tccccctgga agctccctcg tgcgctctcc tgttccgacc ctgccgctta ccggatacct
     6901 gtccgccttt ctcccttcgg gaagcgtggc gctttctcat agctcacgct gtaggtatct
     6961 cagttcggtg taggtcgttc gctccaagct gggctgtgtg cacgaacccc ccgttcagcc
     7021 cgaccgctgc gccttatccg gtaactatcg tcttgagtcc aacccggtaa gacacgactt
     7081 atcgccactg gcagcagcca ctggtaacag gattagcaga gcgaggtatg taggcggtgc
     7141 tacagagttc ttgaagtggt ggcctaacta cggctacact agaagaacag tatttggtat
     7201 ctgcgctctg ctgaagccag ttaccttcgg aaaaagagtt ggtagctctt gatccggcaa
     7261 acaaaccacc gctggtagcg gtggtttttt tgtttgcaag cagcagatta cgcgcagaaa
     7321 aaaaggatct caagaagatc ctttgatctt ttctacgggg tctgacgctc agtggaacga
     7381 aaactcacgt taagggattt tggtcatgag attatcaaaa aggatcttca cctagatcct
     7441 tttaaattaa aaatgaagtt ttaaatcaat ctaaagtata tatgagtaaa cttggtctga
     7501 cagttaccaa tgcttaatca gtgaggcacc tatctcagcg atctgtctat ttcgttcatc
     7561 catagttgcc tgactccccg tcgtgtagat aactacgata cgggagggct taccatctgg
     7621 ccccagtgct gcaatgatac cgcgagaccc acgctcaccg gctccagatt tatcagcaat
     7681 aaaccagcca gccggaaggg ccgagcgcag aagtggtcct gcaactttat ccgcctccat
     7741 ccagtctatt aattgttgcc gggaagctag agtaagtagt tcgccagtta atagtttgcg
     7801 caacgttgtt gccattgcta caggcatcgt ggtgtcacgc tcgtcgtttg gtatggcttc
     7861 attcagctcc ggttcccaac gatcaaggcg agttacatga tcccccatgt tgtgcaaaaa
     7921 agcggttagc tccttcggtc ctccgatcgt tgtcagaagt aagttggccg cagtgttatc
     7981 actcatggtt atggcagcac tgcataattc tcttactgtc atgccatccg taagatgctt
     8041 ttctgtgact ggtgagtact caaccaagtc attctgagaa tagtgtatgc ggcgaccgag
     8101 ttgctcttgc ccggcgtcaa tacgggataa taccgcgcca catagcagaa ctttaaaagt
     8161 gctcatcatt ggaaaacgtt cttcggggcg aaaactctca aggatcttac cgctgttgag
     8221 atccagttcg atgtaaccca ctcgtgcacc caactgatct tcagcatctt ttactttcac
     8281 cagcgtttct gggtgagcaa aaacaggaag gcaaaatgcc gcaaaaaagg gaataagggc
     8341 gacacggaaa tgttgaatac tcatactctt cctttttcaa tattattgaa gcatttatca
     8401 gggttattgt ctcatgagcg gatacatatt tgaatgtatt tagaaaaata aacaaatagg
     8461 ggttccgcgc acatttcccc gaaaagtgcc acctgacgtc taagaaacca ttattatcat
     8521 gacattaacc tataaaaata ggcgtatcac gaggcccttt cgtc
//

PiggyBac TRE3G_KrasG12V_mCherry

LOCUS BH1.6_TRE3G_KrasG12V_mCherry                8564 bp DNA     circular SYN 24-OCT-2022
DEFINITION  synthetic circular DNA
ACCESSION   .
VERSION     .
KEYWORDS    .
SOURCE      synthetic DNA construct
  ORGANISM  synthetic DNA construct
REFERENCE   1  (bases 1 to 8564)
  AUTHORS   .
  TITLE     Direct Submission
  JOURNAL   Exported Oct 25, 2022 from SnapGene 6.0.5
            https://www.snapgene.com
COMMENT     U179MFC040-33
FEATURES             Location/Qualifiers
     source          1..8564
                     /mol_type="other DNA"
                     /organism="synthetic DNA construct"
     primer_bind     379..395
                     /label=M13 fwd
     protein_bind    complement(445..682)
                     /label=PB-TRE 874..1111
     enhancer        719..950
                     /label=Insulator 1148..1379
     promoter        1090..1468
                     /label=TRE3G promoter
     protein_bind    1098..1116
                     /label=tet operator
     protein_bind    1134..1152
                     /label=tet operator
     protein_bind    1170..1188
                     /label=tet operator
     protein_bind    1206..1224
                     /label=tet operator
     protein_bind    1242..1260
                     /label=tet operator
     protein_bind    1278..1296
                     /label=tet operator
     protein_bind    1314..1332
                     /label=tet operator
     CDS             1475..2044
                     /codon_start=1
                     /label=KRAS G12V
                     /translation="MTEYKLVVVGAVGVGKSALTIQLIQNHFVDEYDPTIEDSYRKQVV
                     IDGETCLLDILDTAGQEEYSAMRDQYMRTGEGFLCVFAINNTKSFEDIHHYREQIKRVK
                     DSEDVPMVLVGNKCDLPSRTVDTKQAQDLARSYGIPFIETSAKTRQRVEDAFYTLVREI
                     RQYRLKKISKEEKTPGCVKIKKCIIM"
     misc_feature    2127..2549
                     /label=IRES
     CDS             2690..3400
                     /label=mCherry
     polyA_signal    3407..3455
                     /label=poly(A) signal
     promoter        3472..3972
                     /label=hPGK promoter
     CDS             3986..4582
                     /label=PuroR
     CDS             4592..4645
                     /label=T2A
     CDS             4646..5392
                     /label=Tet-On(R) 3G
     polyA_signal    5399..5520
                     /label=SV40 poly(A) signal
     enhancer        5617..5848
                     /label=Insulator 1148..1379
     promoter        complement(5889..5907)
                     /label=T3 promoter
     promoter        5914..5932
                     /label=T7 promoter
     protein_bind    5983..6291
                     /label=PB-TRE 2067..2375
     primer_bind     complement(6343..6359)
                     /label=M13 rev
     protein_bind    6367..6383
                     /label=lac operator
     promoter        complement(6398..6415)
                     /label=lac promoter
     protein_bind    6436..6457
                     /label=CAP binding site
     rep_origin      complement(6745..7333)
                     /direction=LEFT
                     /label=ori
     CDS             complement(7504..8295)
                     /label=AmpR
     CDS             complement(8296..8364)
                     /label=AmpR
     promoter        complement(8365..8469)
                     /label=AmpR promoter
ORIGIN
        1 tcgcgcgttt cggtgatgac ggtgaaaacc tctgacacat gcagctcccg gagacggtca
       61 cagcttgtct gtaagcggat gccgggagca gacaagcccg tcagggcgcg tcagcgggtg
      121 ttggcgggtg tcggggctgg cttaactatg cggcatcaga gcagattgta ctgagagtgc
      181 accatatgcg gtgtgaaata ccgcacagat gcgtaaggag aaaataccgc atcaggcgcc
      241 attcgccatt caggctgcgc aactgttggg aagggcgatc ggtgcgggcc tcttcgctat
      301 tacgccagct ggcgaaaggg ggatgtgctg caaggcgatt aagttgggta acgccagggt
      361 tttcccagtc acgacgttgt aaaacgacgg ccagtgaatt aggcgcgcct gctcgacacg
      421 ctgcagaaca cgcagctaga ttaaccctag aaagataatc atattgtgac gtacgttaaa
      481 gataatcatg cgtaaaattg acgcatgtgt tttatcggtc tgtatatcga ggtttattta
      541 ttaatttgaa tagatattaa gttttattat atttacactt acatactaat aataaattca
      601 acaaacaatt tatttatgtt tatttattta ttaaaaaaaa acaaaaactc aaaatttctt
      661 ctataaagta acaaaacttt tatcgaattg ctgcagcccg ggcgatccat tagtactaga
      721 gggacagccc ccccccaaag cccccaggga tgtaattacg tccctccccc gctagggggc
      781 agcagcgagc cgcccggggc tccgctccgg tccggcgctc cccccgcatc cccgagccgg
      841 cagcgtgcgg ggacagcccg ggcacgggga aggtggcacg ggatcgcttt cctctgaacg
      901 cttctcgctg ctctttgagc ctgcagacac ctggggggat acggggaaaa ggcctccaag
      961 gccagcttcc cacaataagt tgggtgaatt ttggctcatt cctcctttct ataggattga
     1021 ggtcagagct ttgtgatggc aattctgtgg aatgtgtgtc agttagggtg tggaaagtca
     1081 agcatcgatg agtttactcc ctatcagtga tagagaacgt atgaagagtt tactccctat
     1141 cagtgataga gaacgtatgc agactttact ccctatcagt gatagagaac gtataaggag
     1201 tttactccct atcagtgata gagaacgtat gaccagttta ctccctatca gtgatagaga
     1261 acgtatctac agtttactcc ctatcagtga tagagaacgt atatccagtt tactccctat
     1321 cagtgataga gaacgtataa gctttaggcg tgtacggtgg gcgcctataa aagcagagct
     1381 cgtttagtga accgtcagat cgcctggagc aattccacaa cacttttgtc ttataccaac
     1441 tttccgtacc acttcctacc ctcgtaaaac cggtatgact gaatataaac ttgtggtagt
     1501 tggagctgtt ggcgtaggca agagtgcctt gacgatacag ctaattcaga atcattttgt
     1561 ggacgaatat gatccaacaa tagaggattc ctacaggaag caagtagtaa ttgatggaga
     1621 aacctgtctc ttggatattc tcgacacagc aggtcaagag gagtacagtg caatgaggga
     1681 ccagtacatg aggactgggg agggctttct ttgtgtattt gccataaata atactaaatc
     1741 atttgaagat attcaccatt atagagaaca aattaaaaga gttaaggact ctgaagatgt
     1801 acctatggtc ctagtaggaa ataaatgtga tttgccttct agaacagtag acacaaaaca
     1861 ggctcaggac ttagcaagaa gttatggaat tccttttatt gaaacatcag caaagacaag
     1921 acagagagtg gaggatgctt tttatacatt ggtgagagag atccgacaat acagattgaa
     1981 aaaaatcagc aaagaagaaa agactcctgg ctgtgtgaaa attaaaaaat gcattataat
     2041 gtaagttaac cgcccgcccc acgacccgca gcgcccgacc gaaaggagcg cacgacccca
     2101 tcatccaatt ccgccccccc cccctaacgt tactggccga agccgcttgg aataaggccg
     2161 gtgtgcgttt gtctatatgt tattttccac catattgccg tcttttggca atgtgagggc
     2221 ccggaaacct ggccctgtct tcttgacgag cattcctagg ggtctttccc ctctcgccaa
     2281 aggaatgcaa ggtctgttga atgtcgtgaa ggaagcagtt cctctggaag cttcttgaag
     2341 acaaacaacg tctgtagcga ccctttgcag gcagcggaac cccccacctg gcgacaggtg
     2401 cctctgcggc caaaagccac gtgtataaga tacacctgca aaggcggcac aaccccagtg
     2461 ccacgttgtg agttggatag ttgtggaaag agtcaaatgg ctctcctcaa gcgtattcaa
     2521 caaggggctg aaggatgccc agaaggtacc ccattgtatg ggatctgatc tggggcctcg
     2581 gtgcacatgc tttacatgtg tttagtcgag gttaaaaaac gtctaggccc cccgaaccac
     2641 ggggacgtgg ttttcctttg aaaaacacga tgataatatg gccacaacca tggtgagcaa
     2701 gggcgaggag gataacatgg ccatcatcaa ggagttcatg cgcttcaagg tgcacatgga
     2761 gggctccgtg aacggccacg agttcgagat cgagggcgag ggcgagggcc gcccctacga
     2821 gggcacccag accgccaagc tgaaggtgac caagggtggc cccctgccct tcgcctggga
     2881 catcctgtcc cctcagttca tgtacggctc caaggcctac gtgaagcacc ccgccgacat
     2941 ccccgactac ttgaagctgt ccttccccga gggcttcaag tgggagcgcg tgatgaactt
     3001 cgaggacggc ggcgtggtga ccgtgaccca ggactcctcc ctgcaggacg gcgagttcat
     3061 ctacaaggtg aagctgcgcg gcaccaactt cccctccgac ggccccgtaa tgcagaagaa
     3121 gaccatgggc tgggaggcct cctccgagcg gatgtacccc gaggacggcg ccctgaaggg
     3181 cgagatcaag cagaggctga agctgaagga cggcggccac tacgacgctg aggtcaagac
     3241 cacctacaag gccaagaagc ccgtgcagct gcccggcgcc tacaacgtca acatcaagtt
     3301 ggacatcacc tcccacaacg aggactacac catcgtggaa cagtacgaac gcgccgaggg
     3361 ccgccactcc accggcggca tggacgagct gtacaagtag gtcgacaata aaatatcttt
     3421 attttcatta catctgtgtg ttggtttttt gtgtgacgcg tggggcccgg gttgcgcctt
     3481 ttccaaggca gccctgggtt tgcgcaggga cgcggctgct ctgggcgtgg ttccgggaaa
     3541 cgcagcggcg ccgaccctgg gtctcgcaca ttcttcacgt ccgttcgcag cgtcacccgg
     3601 atcttcgccg ctacccttgt gggccccccg gcgacgcttc ctgctccgcc cctaagtcgg
     3661 gaaggttcct tgcggttcgc ggcgtgccgg acgtgacaaa cggaagccgc acgtctcact
     3721 agtaccctcg cagacggaca gcgccaggga gcaatggcag cgcgccgacc gcgatgggct
     3781 gtggccaata gcggctgctc agcagggcgc gccgagagca gcggccggga aggggcggtg
     3841 cgggaggcgg ggtgtggggc ggtagtgtgg gccctgttcc tgcccgcgcg gtgttccgca
     3901 ttctgcaagc ctccggagcg cacgtcggca gtcggctccc tcgttgaccg aatcaccgac
     3961 ctctctcccc agggagatct caaccatgac cgagtacaag cccacggtgc gcctcgccac
     4021 ccgcgacgac gtccccaggg ccgtacgcac cctcgccgcc gcgttcgccg actaccccgc
     4081 cacgcgccac accgtcgatc cggaccgcca catcgagcgg gtcaccgagc tgcaagaact
     4141 cttcctcacg cgcgtcgggc tcgacatcgg caaggtgtgg gtcgcggacg acggcgccgc
     4201 cgtggcggtc tggaccacgc cggagagcgt cgaagcgggg gcggtgttcg ccgagatcgg
     4261 cccgcgcatg gccgagttga gcggttcccg gctggccgcg cagcaacaga tggaaggcct
     4321 cctggcgccg caccggccca aggagcccgc gtggttcctg gccaccgtcg gcgtctcgcc
     4381 cgaccaccag ggcaagggtc tgggcagcgc cgtcgtgctc cccggagtgg aggcggccga
     4441 gcgcgccggg gtgcccgcct tcctggagac ctccgcgccc cgcaacctcc ccttctacga
     4501 gcggctcggc ttcaccgtca ccgccgacgt cgaggtgccc gaaggaccgc gcacctggtg
     4561 catgacccgc aagcccggtg ccggaagcgg agaaggtaga ggttctctcc tcacttgtgg
     4621 tgatgttgaa gaaaaccctg gtccaatgtc tagactggac aagagcaaag tcataaactc
     4681 tgctctggaa ttactcaatg gagtcggtat cgaaggcctg acgacaagga aactcgctca
     4741 aaagctggga gttgagcagc ctaccctgta ctggcacgtg aagaacaagc gggccctgct
     4801 cgatgccctg ccaatcgaga tgctggacag gcatcatacc cactcctgcc ccctggaagg
     4861 cgagtcatgg caagactttc tgcggaacaa cgccaagtca taccgctgtg ctctcctctc
     4921 acatcgcgac ggggctaaag tgcatctcgg cacccgccca acagagaaac agtacgaaac
     4981 cctggaaaat cagctcgcgt tcctgtgtca gcaaggcttc tccctggaga acgcactgta
     5041 cgctctgtcc gccgtgggcc actttacact gggctgcgta ttggaggaac aggagcatca
     5101 agtagcaaaa gaggaaagag agacacctac caccgattct atgcccccac ttctgaaaca
     5161 agcaattgag ctgttcgacc ggcagggagc cgaacctgcc ttccttttcg gcctggaact
     5221 aatcatatgt ggcctggaga aacagctaaa gtgcgaaagc ggcgggccga ccgacgccct
     5281 tgacgatttt gacttagaca tgctcccagc cgatgccctt gacgactttg accttgatat
     5341 gctgcctgct gacgctcttg acgattttga ccttgacatg ctccccgggt aagctagcaa
     5401 cttgtttatt gcagcttata atggttacaa ataaagcaat agcatcacaa atttcacaaa
     5461 taaagcattt ttttcactgc attctagttg tggtttgtcc aaactcatca atgtatctta
     5521 tgtcctcaca ggaacgaagt ccctaaagaa acagtggcag ccaggtttag ccccggaatt
     5581 gactggattc cttttttagg gcccattggt atggcttttt ccccgtatcc ccccaggtgt
     5641 ctgcaggctc aaagagcagc gagaagcgtt cagaggaaag cgatcccgtg ccaccttccc
     5701 cgtgcccggg ctgtccccgc acgctgccgg ctcggggatg cggggggagc gccggaccgg
     5761 agcggagccc cgggcggctc gctgctgccc cctagcgggg gagggacgta attacatccc
     5821 tgggggcttt gggggggggc tgtccctctt gaacgaccgc caccgcggtg gagctccagc
     5881 ttttgttccc tttagtgagg gttaattaaa tcttaatacg actcactata gggcgaattg
     5941 ggaaccgggc cccccctgga gatcgacggt atccataagc ttgatatcta taacaagaaa
     6001 atatatatat aataagttat cacgtaagta gaacatgaaa taacaatata attatcgtat
     6061 gagttaaatc ttaaaagtca cgtaaaagat aatcatgcgt cattttgact cacgcggtcg
     6121 ttatagttca aaatcagtga cacttaccgc attgacaagc acgcctcacg ggagctccaa
     6181 gcggcgactg agatgtccta aatgcacagc gacggattcg cgctatttag aaagagagag
     6241 caatatttca agaatgcatg cgtcaatttt acgcagacta tctttctagg gttaatctag
     6301 ctgcatcagg atcatagcgg ccgccagctt ggcgtaatca tggtcatagc tgtttcctgt
     6361 gtgaaattgt tatccgctca caattccaca caacatacga gccggaagca taaagtgtaa
     6421 agcctggggt gcctaatgag tgagctaact cacattaatt gcgttgcgct cactgcccgc
     6481 tttccagtcg ggaaacctgt cgtgccagct gcattaatga atcggccaac gcgcggggag
     6541 aggcggtttg cgtattgggc gctcttccgc ttcctcgctc actgactcgc tgcgctcggt
     6601 cgttcggctg cggcgagcgg tatcagctca ctcaaaggcg gtaatacggt tatccacaga
     6661 atcaggggat aacgcaggaa agaacatgtg agcaaaaggc cagcaaaagg ccaggaaccg
     6721 taaaaaggcc gcgttgctgg cgtttttcca taggctccgc ccccctgacg agcatcacaa
     6781 aaatcgacgc tcaagtcaga ggtggcgaaa cccgacagga ctataaagat accaggcgtt
     6841 tccccctgga agctccctcg tgcgctctcc tgttccgacc ctgccgctta ccggatacct
     6901 gtccgccttt ctcccttcgg gaagcgtggc gctttctcat agctcacgct gtaggtatct
     6961 cagttcggtg taggtcgttc gctccaagct gggctgtgtg cacgaacccc ccgttcagcc
     7021 cgaccgctgc gccttatccg gtaactatcg tcttgagtcc aacccggtaa gacacgactt
     7081 atcgccactg gcagcagcca ctggtaacag gattagcaga gcgaggtatg taggcggtgc
     7141 tacagagttc ttgaagtggt ggcctaacta cggctacact agaagaacag tatttggtat
     7201 ctgcgctctg ctgaagccag ttaccttcgg aaaaagagtt ggtagctctt gatccggcaa
     7261 acaaaccacc gctggtagcg gtggtttttt tgtttgcaag cagcagatta cgcgcagaaa
     7321 aaaaggatct caagaagatc ctttgatctt ttctacgggg tctgacgctc agtggaacga
     7381 aaactcacgt taagggattt tggtcatgag attatcaaaa aggatcttca cctagatcct
     7441 tttaaattaa aaatgaagtt ttaaatcaat ctaaagtata tatgagtaaa cttggtctga
     7501 cagttaccaa tgcttaatca gtgaggcacc tatctcagcg atctgtctat ttcgttcatc
     7561 catagttgcc tgactccccg tcgtgtagat aactacgata cgggagggct taccatctgg
     7621 ccccagtgct gcaatgatac cgcgagaccc acgctcaccg gctccagatt tatcagcaat
     7681 aaaccagcca gccggaaggg ccgagcgcag aagtggtcct gcaactttat ccgcctccat
     7741 ccagtctatt aattgttgcc gggaagctag agtaagtagt tcgccagtta atagtttgcg
     7801 caacgttgtt gccattgcta caggcatcgt ggtgtcacgc tcgtcgtttg gtatggcttc
     7861 attcagctcc ggttcccaac gatcaaggcg agttacatga tcccccatgt tgtgcaaaaa
     7921 agcggttagc tccttcggtc ctccgatcgt tgtcagaagt aagttggccg cagtgttatc
     7981 actcatggtt atggcagcac tgcataattc tcttactgtc atgccatccg taagatgctt
     8041 ttctgtgact ggtgagtact caaccaagtc attctgagaa tagtgtatgc ggcgaccgag
     8101 ttgctcttgc ccggcgtcaa tacgggataa taccgcgcca catagcagaa ctttaaaagt
     8161 gctcatcatt ggaaaacgtt cttcggggcg aaaactctca aggatcttac cgctgttgag
     8221 atccagttcg atgtaaccca ctcgtgcacc caactgatct tcagcatctt ttactttcac
     8281 cagcgtttct gggtgagcaa aaacaggaag gcaaaatgcc gcaaaaaagg gaataagggc
     8341 gacacggaaa tgttgaatac tcatactctt cctttttcaa tattattgaa gcatttatca
     8401 gggttattgt ctcatgagcg gatacatatt tgaatgtatt tagaaaaata aacaaatagg
     8461 ggttccgcgc acatttcccc gaaaagtgcc acctgacgtc taagaaacca ttattatcat
     8521 gacattaacc tataaaaata ggcgtatcac gaggcccttt cgtc
//

PiggyBac TRE3G_KrasWT_mCherry

LOCUS BH1.6_TRE3G_KrasWT_mCherry                8564 bp DNA     circular SYN 24-OCT-2022
DEFINITION  synthetic circular DNA
ACCESSION   .
VERSION     .
KEYWORDS    .
SOURCE      synthetic DNA construct
  ORGANISM  synthetic DNA construct
REFERENCE   1  (bases 1 to 8564)
  AUTHORS   .
  TITLE     Direct Submission
  JOURNAL   Exported Oct 25, 2022 from SnapGene 6.0.5
            https://www.snapgene.com
COMMENT     U179MFC040-21
FEATURES             Location/Qualifiers
     source          1..8564
                     /mol_type="other DNA"
                     /organism="synthetic DNA construct"
     primer_bind     379..395
                     /label=M13 fwd
     protein_bind    complement(445..682)
                     /label=PB-TRE 874..1111
     enhancer        719..950
                     /label=Insulator 1148..1379
     promoter        1090..1468
                     /label=TRE3G promoter
     protein_bind    1098..1116
                     /label=tet operator
     protein_bind    1134..1152
                     /label=tet operator
     protein_bind    1170..1188
                     /label=tet operator
     protein_bind    1206..1224
                     /label=tet operator
     protein_bind    1242..1260
                     /label=tet operator
     protein_bind    1278..1296
                     /label=tet operator
     protein_bind    1314..1332
                     /label=tet operator
     CDS             1475..2044
                     /codon_start=1
                     /label=KRAS WT
                     /translation="MTEYKLVVVGAGGVGKSALTIQLIQNHFVDEYDPTIEDSYRKQVV
                     IDGETCLLDILDTAGQEEYSAMRDQYMRTGEGFLCVFAINNTKSFEDIHHYREQIKRVK
                     DSEDVPMVLVGNKCDLPSRTVDTKQAQDLARSYGIPFIETSAKTRQRVEDAFYTLVREI
                     RQYRLKKISKEEKTPGCVKIKKCIIM"
     misc_feature    2127..2549
                     /label=IRES
     CDS             2690..3400
                     /label=mCherry
     polyA_signal    3407..3455
                     /label=poly(A) signal
     promoter        3472..3972
                     /label=hPGK promoter
     CDS             3986..4582
                     /label=PuroR
     CDS             4592..4645
                     /label=T2A
     CDS             4646..5392
                     /label=Tet-On(R) 3G
     polyA_signal    5399..5520
                     /label=SV40 poly(A) signal
     enhancer        5617..5848
                     /label=Insulator 1148..1379
     promoter        complement(5889..5907)
                     /label=T3 promoter
     promoter        5914..5932
                     /label=T7 promoter
     protein_bind    5983..6291
                     /label=PB-TRE 2067..2375
     primer_bind     complement(6343..6359)
                     /label=M13 rev
     protein_bind    6367..6383
                     /label=lac operator
     promoter        complement(6398..6415)
                     /label=lac promoter
     protein_bind    6436..6457
                     /label=CAP binding site
     rep_origin      complement(6745..7333)
                     /direction=LEFT
                     /label=ori
     CDS             complement(7504..8295)
                     /label=AmpR
     CDS             complement(8296..8364)
                     /label=AmpR
     promoter        complement(8365..8469)
                     /label=AmpR promoter
ORIGIN
        1 tcgcgcgttt cggtgatgac ggtgaaaacc tctgacacat gcagctcccg gagacggtca
       61 cagcttgtct gtaagcggat gccgggagca gacaagcccg tcagggcgcg tcagcgggtg
      121 ttggcgggtg tcggggctgg cttaactatg cggcatcaga gcagattgta ctgagagtgc
      181 accatatgcg gtgtgaaata ccgcacagat gcgtaaggag aaaataccgc atcaggcgcc
      241 attcgccatt caggctgcgc aactgttggg aagggcgatc ggtgcgggcc tcttcgctat
      301 tacgccagct ggcgaaaggg ggatgtgctg caaggcgatt aagttgggta acgccagggt
      361 tttcccagtc acgacgttgt aaaacgacgg ccagtgaatt aggcgcgcct gctcgacacg
      421 ctgcagaaca cgcagctaga ttaaccctag aaagataatc atattgtgac gtacgttaaa
      481 gataatcatg cgtaaaattg acgcatgtgt tttatcggtc tgtatatcga ggtttattta
      541 ttaatttgaa tagatattaa gttttattat atttacactt acatactaat aataaattca
      601 acaaacaatt tatttatgtt tatttattta ttaaaaaaaa acaaaaactc aaaatttctt
      661 ctataaagta acaaaacttt tatcgaattg ctgcagcccg ggcgatccat tagtactaga
      721 gggacagccc ccccccaaag cccccaggga tgtaattacg tccctccccc gctagggggc
      781 agcagcgagc cgcccggggc tccgctccgg tccggcgctc cccccgcatc cccgagccgg
      841 cagcgtgcgg ggacagcccg ggcacgggga aggtggcacg ggatcgcttt cctctgaacg
      901 cttctcgctg ctctttgagc ctgcagacac ctggggggat acggggaaaa ggcctccaag
      961 gccagcttcc cacaataagt tgggtgaatt ttggctcatt cctcctttct ataggattga
     1021 ggtcagagct ttgtgatggc aattctgtgg aatgtgtgtc agttagggtg tggaaagtca
     1081 agcatcgatg agtttactcc ctatcagtga tagagaacgt atgaagagtt tactccctat
     1141 cagtgataga gaacgtatgc agactttact ccctatcagt gatagagaac gtataaggag
     1201 tttactccct atcagtgata gagaacgtat gaccagttta ctccctatca gtgatagaga
     1261 acgtatctac agtttactcc ctatcagtga tagagaacgt atatccagtt tactccctat
     1321 cagtgataga gaacgtataa gctttaggcg tgtacggtgg gcgcctataa aagcagagct
     1381 cgtttagtga accgtcagat cgcctggagc aattccacaa cacttttgtc ttataccaac
     1441 tttccgtacc acttcctacc ctcgtaaaac cggtatgact gaatataaac ttgtggtagt
     1501 tggagctggt ggcgtaggca agagtgcctt gacgatacag ctaattcaga atcattttgt
     1561 ggacgaatat gatccaacaa tagaggattc ctacaggaag caagtagtaa ttgatggaga
     1621 aacctgtctc ttggatattc tcgacacagc aggtcaagag gagtacagtg caatgaggga
     1681 ccagtacatg aggactgggg agggctttct ttgtgtattt gccataaata atactaaatc
     1741 atttgaagat attcaccatt atagagaaca aattaaaaga gttaaggact ctgaagatgt
     1801 acctatggtc ctagtaggaa ataaatgtga tttgccttct agaacagtag acacaaaaca
     1861 ggctcaggac ttagcaagaa gttatggaat tccttttatt gaaacatcag caaagacaag
     1921 acagagagtg gaggatgctt tttatacatt ggtgagagag atccgacaat acagattgaa
     1981 aaaaatcagc aaagaagaaa agactcctgg ctgtgtgaaa attaaaaaat gcattataat
     2041 gtaagttaac cgcccgcccc acgacccgca gcgcccgacc gaaaggagcg cacgacccca
     2101 tcatccaatt ccgccccccc cccctaacgt tactggccga agccgcttgg aataaggccg
     2161 gtgtgcgttt gtctatatgt tattttccac catattgccg tcttttggca atgtgagggc
     2221 ccggaaacct ggccctgtct tcttgacgag cattcctagg ggtctttccc ctctcgccaa
     2281 aggaatgcaa ggtctgttga atgtcgtgaa ggaagcagtt cctctggaag cttcttgaag
     2341 acaaacaacg tctgtagcga ccctttgcag gcagcggaac cccccacctg gcgacaggtg
     2401 cctctgcggc caaaagccac gtgtataaga tacacctgca aaggcggcac aaccccagtg
     2461 ccacgttgtg agttggatag ttgtggaaag agtcaaatgg ctctcctcaa gcgtattcaa
     2521 caaggggctg aaggatgccc agaaggtacc ccattgtatg ggatctgatc tggggcctcg
     2581 gtgcacatgc tttacatgtg tttagtcgag gttaaaaaac gtctaggccc cccgaaccac
     2641 ggggacgtgg ttttcctttg aaaaacacga tgataatatg gccacaacca tggtgagcaa
     2701 gggcgaggag gataacatgg ccatcatcaa ggagttcatg cgcttcaagg tgcacatgga
     2761 gggctccgtg aacggccacg agttcgagat cgagggcgag ggcgagggcc gcccctacga
     2821 gggcacccag accgccaagc tgaaggtgac caagggtggc cccctgccct tcgcctggga
     2881 catcctgtcc cctcagttca tgtacggctc caaggcctac gtgaagcacc ccgccgacat
     2941 ccccgactac ttgaagctgt ccttccccga gggcttcaag tgggagcgcg tgatgaactt
     3001 cgaggacggc ggcgtggtga ccgtgaccca ggactcctcc ctgcaggacg gcgagttcat
     3061 ctacaaggtg aagctgcgcg gcaccaactt cccctccgac ggccccgtaa tgcagaagaa
     3121 gaccatgggc tgggaggcct cctccgagcg gatgtacccc gaggacggcg ccctgaaggg
     3181 cgagatcaag cagaggctga agctgaagga cggcggccac tacgacgctg aggtcaagac
     3241 cacctacaag gccaagaagc ccgtgcagct gcccggcgcc tacaacgtca acatcaagtt
     3301 ggacatcacc tcccacaacg aggactacac catcgtggaa cagtacgaac gcgccgaggg
     3361 ccgccactcc accggcggca tggacgagct gtacaagtag gtcgacaata aaatatcttt
     3421 attttcatta catctgtgtg ttggtttttt gtgtgacgcg tggggcccgg gttgcgcctt
     3481 ttccaaggca gccctgggtt tgcgcaggga cgcggctgct ctgggcgtgg ttccgggaaa
     3541 cgcagcggcg ccgaccctgg gtctcgcaca ttcttcacgt ccgttcgcag cgtcacccgg
     3601 atcttcgccg ctacccttgt gggccccccg gcgacgcttc ctgctccgcc cctaagtcgg
     3661 gaaggttcct tgcggttcgc ggcgtgccgg acgtgacaaa cggaagccgc acgtctcact
     3721 agtaccctcg cagacggaca gcgccaggga gcaatggcag cgcgccgacc gcgatgggct
     3781 gtggccaata gcggctgctc agcagggcgc gccgagagca gcggccggga aggggcggtg
     3841 cgggaggcgg ggtgtggggc ggtagtgtgg gccctgttcc tgcccgcgcg gtgttccgca
     3901 ttctgcaagc ctccggagcg cacgtcggca gtcggctccc tcgttgaccg aatcaccgac
     3961 ctctctcccc agggagatct caaccatgac cgagtacaag cccacggtgc gcctcgccac
     4021 ccgcgacgac gtccccaggg ccgtacgcac cctcgccgcc gcgttcgccg actaccccgc
     4081 cacgcgccac accgtcgatc cggaccgcca catcgagcgg gtcaccgagc tgcaagaact
     4141 cttcctcacg cgcgtcgggc tcgacatcgg caaggtgtgg gtcgcggacg acggcgccgc
     4201 cgtggcggtc tggaccacgc cggagagcgt cgaagcgggg gcggtgttcg ccgagatcgg
     4261 cccgcgcatg gccgagttga gcggttcccg gctggccgcg cagcaacaga tggaaggcct
     4321 cctggcgccg caccggccca aggagcccgc gtggttcctg gccaccgtcg gcgtctcgcc
     4381 cgaccaccag ggcaagggtc tgggcagcgc cgtcgtgctc cccggagtgg aggcggccga
     4441 gcgcgccggg gtgcccgcct tcctggagac ctccgcgccc cgcaacctcc ccttctacga
     4501 gcggctcggc ttcaccgtca ccgccgacgt cgaggtgccc gaaggaccgc gcacctggtg
     4561 catgacccgc aagcccggtg ccggaagcgg agaaggtaga ggttctctcc tcacttgtgg
     4621 tgatgttgaa gaaaaccctg gtccaatgtc tagactggac aagagcaaag tcataaactc
     4681 tgctctggaa ttactcaatg gagtcggtat cgaaggcctg acgacaagga aactcgctca
     4741 aaagctggga gttgagcagc ctaccctgta ctggcacgtg aagaacaagc gggccctgct
     4801 cgatgccctg ccaatcgaga tgctggacag gcatcatacc cactcctgcc ccctggaagg
     4861 cgagtcatgg caagactttc tgcggaacaa cgccaagtca taccgctgtg ctctcctctc
     4921 acatcgcgac ggggctaaag tgcatctcgg cacccgccca acagagaaac agtacgaaac
     4981 cctggaaaat cagctcgcgt tcctgtgtca gcaaggcttc tccctggaga acgcactgta
     5041 cgctctgtcc gccgtgggcc actttacact gggctgcgta ttggaggaac aggagcatca
     5101 agtagcaaaa gaggaaagag agacacctac caccgattct atgcccccac ttctgaaaca
     5161 agcaattgag ctgttcgacc ggcagggagc cgaacctgcc ttccttttcg gcctggaact
     5221 aatcatatgt ggcctggaga aacagctaaa gtgcgaaagc ggcgggccga ccgacgccct
     5281 tgacgatttt gacttagaca tgctcccagc cgatgccctt gacgactttg accttgatat
     5341 gctgcctgct gacgctcttg acgattttga ccttgacatg ctccccgggt aagctagcaa
     5401 cttgtttatt gcagcttata atggttacaa ataaagcaat agcatcacaa atttcacaaa
     5461 taaagcattt ttttcactgc attctagttg tggtttgtcc aaactcatca atgtatctta
     5521 tgtcctcaca ggaacgaagt ccctaaagaa acagtggcag ccaggtttag ccccggaatt
     5581 gactggattc cttttttagg gcccattggt atggcttttt ccccgtatcc ccccaggtgt
     5641 ctgcaggctc aaagagcagc gagaagcgtt cagaggaaag cgatcccgtg ccaccttccc
     5701 cgtgcccggg ctgtccccgc acgctgccgg ctcggggatg cggggggagc gccggaccgg
     5761 agcggagccc cgggcggctc gctgctgccc cctagcgggg gagggacgta attacatccc
     5821 tgggggcttt gggggggggc tgtccctctt gaacgaccgc caccgcggtg gagctccagc
     5881 ttttgttccc tttagtgagg gttaattaaa tcttaatacg actcactata gggcgaattg
     5941 ggaaccgggc cccccctgga gatcgacggt atccataagc ttgatatcta taacaagaaa
     6001 atatatatat aataagttat cacgtaagta gaacatgaaa taacaatata attatcgtat
     6061 gagttaaatc ttaaaagtca cgtaaaagat aatcatgcgt cattttgact cacgcggtcg
     6121 ttatagttca aaatcagtga cacttaccgc attgacaagc acgcctcacg ggagctccaa
     6181 gcggcgactg agatgtccta aatgcacagc gacggattcg cgctatttag aaagagagag
     6241 caatatttca agaatgcatg cgtcaatttt acgcagacta tctttctagg gttaatctag
     6301 ctgcatcagg atcatagcgg ccgccagctt ggcgtaatca tggtcatagc tgtttcctgt
     6361 gtgaaattgt tatccgctca caattccaca caacatacga gccggaagca taaagtgtaa
     6421 agcctggggt gcctaatgag tgagctaact cacattaatt gcgttgcgct cactgcccgc
     6481 tttccagtcg ggaaacctgt cgtgccagct gcattaatga atcggccaac gcgcggggag
     6541 aggcggtttg cgtattgggc gctcttccgc ttcctcgctc actgactcgc tgcgctcggt
     6601 cgttcggctg cggcgagcgg tatcagctca ctcaaaggcg gtaatacggt tatccacaga
     6661 atcaggggat aacgcaggaa agaacatgtg agcaaaaggc cagcaaaagg ccaggaaccg
     6721 taaaaaggcc gcgttgctgg cgtttttcca taggctccgc ccccctgacg agcatcacaa
     6781 aaatcgacgc tcaagtcaga ggtggcgaaa cccgacagga ctataaagat accaggcgtt
     6841 tccccctgga agctccctcg tgcgctctcc tgttccgacc ctgccgctta ccggatacct
     6901 gtccgccttt ctcccttcgg gaagcgtggc gctttctcat agctcacgct gtaggtatct
     6961 cagttcggtg taggtcgttc gctccaagct gggctgtgtg cacgaacccc ccgttcagcc
     7021 cgaccgctgc gccttatccg gtaactatcg tcttgagtcc aacccggtaa gacacgactt
     7081 atcgccactg gcagcagcca ctggtaacag gattagcaga gcgaggtatg taggcggtgc
     7141 tacagagttc ttgaagtggt ggcctaacta cggctacact agaagaacag tatttggtat
     7201 ctgcgctctg ctgaagccag ttaccttcgg aaaaagagtt ggtagctctt gatccggcaa
     7261 acaaaccacc gctggtagcg gtggtttttt tgtttgcaag cagcagatta cgcgcagaaa
     7321 aaaaggatct caagaagatc ctttgatctt ttctacgggg tctgacgctc agtggaacga
     7381 aaactcacgt taagggattt tggtcatgag attatcaaaa aggatcttca cctagatcct
     7441 tttaaattaa aaatgaagtt ttaaatcaat ctaaagtata tatgagtaaa cttggtctga
     7501 cagttaccaa tgcttaatca gtgaggcacc tatctcagcg atctgtctat ttcgttcatc
     7561 catagttgcc tgactccccg tcgtgtagat aactacgata cgggagggct taccatctgg
     7621 ccccagtgct gcaatgatac cgcgagaccc acgctcaccg gctccagatt tatcagcaat
     7681 aaaccagcca gccggaaggg ccgagcgcag aagtggtcct gcaactttat ccgcctccat
     7741 ccagtctatt aattgttgcc gggaagctag agtaagtagt tcgccagtta atagtttgcg
     7801 caacgttgtt gccattgcta caggcatcgt ggtgtcacgc tcgtcgtttg gtatggcttc
     7861 attcagctcc ggttcccaac gatcaaggcg agttacatga tcccccatgt tgtgcaaaaa
     7921 agcggttagc tccttcggtc ctccgatcgt tgtcagaagt aagttggccg cagtgttatc
     7981 actcatggtt atggcagcac tgcataattc tcttactgtc atgccatccg taagatgctt
     8041 ttctgtgact ggtgagtact caaccaagtc attctgagaa tagtgtatgc ggcgaccgag
     8101 ttgctcttgc ccggcgtcaa tacgggataa taccgcgcca catagcagaa ctttaaaagt
     8161 gctcatcatt ggaaaacgtt cttcggggcg aaaactctca aggatcttac cgctgttgag
     8221 atccagttcg atgtaaccca ctcgtgcacc caactgatct tcagcatctt ttactttcac
     8281 cagcgtttct gggtgagcaa aaacaggaag gcaaaatgcc gcaaaaaagg gaataagggc
     8341 gacacggaaa tgttgaatac tcatactctt cctttttcaa tattattgaa gcatttatca
     8401 gggttattgt ctcatgagcg gatacatatt tgaatgtatt tagaaaaata aacaaatagg
     8461 ggttccgcgc acatttcccc gaaaagtgcc acctgacgtc taagaaacca ttattatcat
     8521 gacattaacc tataaaaata ggcgtatcac gaggcccttt cgtc
//

PiggyBac TRE3G_KrasG12D_mCherry

LOCUS BH1.6_TRE3G_KrasG12D_mCherry               9134 bp DNA     circular SYN 24-OCT-2022
DEFINITION  synthetic circular DNA
ACCESSION   .
VERSION     .
KEYWORDS    .
SOURCE      synthetic DNA construct
  ORGANISM  synthetic DNA construct
REFERENCE   1  (bases 1 to 9134)
  AUTHORS   .
  TITLE     Direct Submission
  JOURNAL   Exported Oct 25, 2022 from SnapGene 6.0.5
            https://www.snapgene.com
COMMENT     U179MFC040-24
FEATURES             Location/Qualifiers
     source          1..9134
                     /mol_type="other DNA"
                     /organism="synthetic DNA construct"
     primer_bind     379..395
                     /label=M13 fwd
     misc_feature    401
                     /label=EcoRI killer
     protein_bind    complement(445..682)
                     /label=PB-TRE 874..1111
     enhancer        719..950
                     /label=Insulator 1148..1379
     promoter        1090..1468
                     /label=TRE3G promoter
     protein_bind    1098..1116
                     /label=tet operator
     protein_bind    1134..1152
                     /label=tet operator
     protein_bind    1170..1188
                     /label=tet operator
     protein_bind    1206..1224
                     /label=tet operator
     protein_bind    1242..1260
                     /label=tet operator
     protein_bind    1278..1296
                     /label=tet operator
     protein_bind    1314..1332
                     /label=tet operator
     CDS             1475..2044
                     /codon_start=1
                     /label=KRAS G12D
                     /translation="MTEYKLVVVGADGVGKSALTIQLIQNHFVDEYDPTIEDSYRKQVV
                     IDGETCLLDILDTAGQEEYSAMRDQYMRTGEGFLCVFAINNTKSFEDIHHYREQIKRVK
                     DSEDVPMVLVGNKCDLPSRTVDTKQAQDLARSYGIPFIETSAKTRQRVEDAFYTLVREI
                     RQYRLKKISKEEKTPGCVKIKKCIIM"
     misc_feature    2127..2549
                     /label=IRES
     CDS             2690..3400
                     /label=mCherry
     polyA_signal    3407..3455
                     /label=poly(A) signal
     promoter        3472..3972
                     /label=hPGK promoter
     CDS             3986..4582
                     /label=PuroR
     CDS             4592..4645
                     /label=T2A
     CDS             4646..5392
                     /label=Tet-On(R) 3G
     polyA_signal    5399..5520
                     /label=SV40 poly(A) signal
     enhancer        5617..5847
                     /label=Insulator 1701..1931
     promoter        complement(5889..5907)
                     /label=T3 promoter
     promoter        5914..5932
                     /label=T7 promoter
     protein_bind    5983..6291
                     /label=PB-TRE 2067..2375
     misc_feature    6325
                     /label=HindIII killer
     primer_bind     complement(6343..6359)
                     /label=M13 rev
     protein_bind    6367..6383
                     /label=lac operator
     promoter        complement(6391..6397)
                     /label=lac promoter
     promoter        complement(6398..6415)
                     /label=lac promoter
     promoter        complement(6416..6421)
                     /label=lac promoter
     protein_bind    6436..6457
                     /label=CAP binding site
     rep_origin      complement(6745..7333)
                     /direction=LEFT
                     /label=ori
     CDS             complement(7504..8295)
                     /label=AmpR
     CDS             complement(8296..8364)
                     /label=AmpR
     promoter        complement(8365..8469)
                     /label=AmpR promoter
     CDS             complement(8565..8865)
                     /label=AmpR
     CDS             complement(8866..8934)
                     /label=AmpR
     promoter        complement(8935..9039)
                     /label=AmpR promoter
ORIGIN
        1 tcgcgcgttt cggtgatgac ggtgaaaacc tctgacacat gcagctcccg gagacggtca
       61 cagcttgtct gtaagcggat gccgggagca gacaagcccg tcagggcgcg tcagcgggtg
      121 ttggcgggtg tcggggctgg cttaactatg cggcatcaga gcagattgta ctgagagtgc
      181 accatatgcg gtgtgaaata ccgcacagat gcgtaaggag aaaataccgc atcaggcgcc
      241 attcgccatt caggctgcgc aactgttggg aagggcgatc ggtgcgggcc tcttcgctat
      301 tacgccagct ggcgaaaggg ggatgtgctg caaggcgatt aagttgggta acgccagggt
      361 tttcccagtc acgacgttgt aaaacgacgg ccagtgaatt aggcgcgcct gctcgacacg
      421 ctgcagaaca cgcagctaga ttaaccctag aaagataatc atattgtgac gtacgttaaa
      481 gataatcatg cgtaaaattg acgcatgtgt tttatcggtc tgtatatcga ggtttattta
      541 ttaatttgaa tagatattaa gttttattat atttacactt acatactaat aataaattca
      601 acaaacaatt tatttatgtt tatttattta ttaaaaaaaa acaaaaactc aaaatttctt
      661 ctataaagta acaaaacttt tatcgaattg ctgcagcccg ggcgatccat tagtactaga
      721 gggacagccc ccccccaaag cccccaggga tgtaattacg tccctccccc gctagggggc
      781 agcagcgagc cgcccggggc tccgctccgg tccggcgctc cccccgcatc cccgagccgg
      841 cagcgtgcgg ggacagcccg ggcacgggga aggtggcacg ggatcgcttt cctctgaacg
      901 cttctcgctg ctctttgagc ctgcagacac ctggggggat acggggaaaa ggcctccaag
      961 gccagcttcc cacaataagt tgggtgaatt ttggctcatt cctcctttct ataggattga
     1021 ggtcagagct ttgtgatggc aattctgtgg aatgtgtgtc agttagggtg tggaaagtca
     1081 agcatcgatg agtttactcc ctatcagtga tagagaacgt atgaagagtt tactccctat
     1141 cagtgataga gaacgtatgc agactttact ccctatcagt gatagagaac gtataaggag
     1201 tttactccct atcagtgata gagaacgtat gaccagttta ctccctatca gtgatagaga
     1261 acgtatctac agtttactcc ctatcagtga tagagaacgt atatccagtt tactccctat
     1321 cagtgataga gaacgtataa gctttaggcg tgtacggtgg gcgcctataa aagcagagct
     1381 cgtttagtga accgtcagat cgcctggagc aattccacaa cacttttgtc ttataccaac
     1441 tttccgtacc acttcctacc ctcgtaaaac cggtatgact gaatataaac ttgtggtagt
     1501 tggagctgat ggcgtaggca agagtgcctt gacgatacag ctaattcaga atcattttgt
     1561 ggacgaatat gatccaacaa tagaggattc ctacaggaag caagtagtaa ttgatggaga
     1621 aacctgtctc ttggatattc tcgacacagc aggtcaagag gagtacagtg caatgaggga
     1681 ccagtacatg aggactgggg agggctttct ttgtgtattt gccataaata atactaaatc
     1741 atttgaagat attcaccatt atagagaaca aattaaaaga gttaaggact ctgaagatgt
     1801 acctatggtc ctagtaggaa ataaatgtga tttgccttct agaacagtag acacaaaaca
     1861 ggctcaggac ttagcaagaa gttatggaat tccttttatt gaaacatcag caaagacaag
     1921 acagagagtg gaggatgctt tttatacatt ggtgagagag atccgacaat acagattgaa
     1981 aaaaatcagc aaagaagaaa agactcctgg ctgtgtgaaa attaaaaaat gcattataat
     2041 gtaagttaac cgcccgcccc acgacccgca gcgcccgacc gaaaggagcg cacgacccca
     2101 tcatccaatt ccgccccccc cccctaacgt tactggccga agccgcttgg aataaggccg
     2161 gtgtgcgttt gtctatatgt tattttccac catattgccg tcttttggca atgtgagggc
     2221 ccggaaacct ggccctgtct tcttgacgag cattcctagg ggtctttccc ctctcgccaa
     2281 aggaatgcaa ggtctgttga atgtcgtgaa ggaagcagtt cctctggaag cttcttgaag
     2341 acaaacaacg tctgtagcga ccctttgcag gcagcggaac cccccacctg gcgacaggtg
     2401 cctctgcggc caaaagccac gtgtataaga tacacctgca aaggcggcac aaccccagtg
     2461 ccacgttgtg agttggatag ttgtggaaag agtcaaatgg ctctcctcaa gcgtattcaa
     2521 caaggggctg aaggatgccc agaaggtacc ccattgtatg ggatctgatc tggggcctcg
     2581 gtgcacatgc tttacatgtg tttagtcgag gttaaaaaac gtctaggccc cccgaaccac
     2641 ggggacgtgg ttttcctttg aaaaacacga tgataatatg gccacaacca tggtgagcaa
     2701 gggcgaggag gataacatgg ccatcatcaa ggagttcatg cgcttcaagg tgcacatgga
     2761 gggctccgtg aacggccacg agttcgagat cgagggcgag ggcgagggcc gcccctacga
     2821 gggcacccag accgccaagc tgaaggtgac caagggtggc cccctgccct tcgcctggga
     2881 catcctgtcc cctcagttca tgtacggctc caaggcctac gtgaagcacc ccgccgacat
     2941 ccccgactac ttgaagctgt ccttccccga gggcttcaag tgggagcgcg tgatgaactt
     3001 cgaggacggc ggcgtggtga ccgtgaccca ggactcctcc ctgcaggacg gcgagttcat
     3061 ctacaaggtg aagctgcgcg gcaccaactt cccctccgac ggccccgtaa tgcagaagaa
     3121 gaccatgggc tgggaggcct cctccgagcg gatgtacccc gaggacggcg ccctgaaggg
     3181 cgagatcaag cagaggctga agctgaagga cggcggccac tacgacgctg aggtcaagac
     3241 cacctacaag gccaagaagc ccgtgcagct gcccggcgcc tacaacgtca acatcaagtt
     3301 ggacatcacc tcccacaacg aggactacac catcgtggaa cagtacgaac gcgccgaggg
     3361 ccgccactcc accggcggca tggacgagct gtacaagtag gtcgacaata aaatatcttt
     3421 attttcatta catctgtgtg ttggtttttt gtgtgacgcg tggggcccgg gttgcgcctt
     3481 ttccaaggca gccctgggtt tgcgcaggga cgcggctgct ctgggcgtgg ttccgggaaa
     3541 cgcagcggcg ccgaccctgg gtctcgcaca ttcttcacgt ccgttcgcag cgtcacccgg
     3601 atcttcgccg ctacccttgt gggccccccg gcgacgcttc ctgctccgcc cctaagtcgg
     3661 gaaggttcct tgcggttcgc ggcgtgccgg acgtgacaaa cggaagccgc acgtctcact
     3721 agtaccctcg cagacggaca gcgccaggga gcaatggcag cgcgccgacc gcgatgggct
     3781 gtggccaata gcggctgctc agcagggcgc gccgagagca gcggccggga aggggcggtg
     3841 cgggaggcgg ggtgtggggc ggtagtgtgg gccctgttcc tgcccgcgcg gtgttccgca
     3901 ttctgcaagc ctccggagcg cacgtcggca gtcggctccc tcgttgaccg aatcaccgac
     3961 ctctctcccc agggagatct caaccatgac cgagtacaag cccacggtgc gcctcgccac
     4021 ccgcgacgac gtccccaggg ccgtacgcac cctcgccgcc gcgttcgccg actaccccgc
     4081 cacgcgccac accgtcgatc cggaccgcca catcgagcgg gtcaccgagc tgcaagaact
     4141 cttcctcacg cgcgtcgggc tcgacatcgg caaggtgtgg gtcgcggacg acggcgccgc
     4201 cgtggcggtc tggaccacgc cggagagcgt cgaagcgggg gcggtgttcg ccgagatcgg
     4261 cccgcgcatg gccgagttga gcggttcccg gctggccgcg cagcaacaga tggaaggcct
     4321 cctggcgccg caccggccca aggagcccgc gtggttcctg gccaccgtcg gcgtctcgcc
     4381 cgaccaccag ggcaagggtc tgggcagcgc cgtcgtgctc cccggagtgg aggcggccga
     4441 gcgcgccggg gtgcccgcct tcctggagac ctccgcgccc cgcaacctcc ccttctacga
     4501 gcggctcggc ttcaccgtca ccgccgacgt cgaggtgccc gaaggaccgc gcacctggtg
     4561 catgacccgc aagcccggtg ccggaagcgg agaaggtaga ggttctctcc tcacttgtgg
     4621 tgatgttgaa gaaaaccctg gtccaatgtc tagactggac aagagcaaag tcataaactc
     4681 tgctctggaa ttactcaatg gagtcggtat cgaaggcctg acgacaagga aactcgctca
     4741 aaagctggga gttgagcagc ctaccctgta ctggcacgtg aagaacaagc gggccctgct
     4801 cgatgccctg ccaatcgaga tgctggacag gcatcatacc cactcctgcc ccctggaagg
     4861 cgagtcatgg caagactttc tgcggaacaa cgccaagtca taccgctgtg ctctcctctc
     4921 acatcgcgac ggggctaaag tgcatctcgg cacccgccca acagagaaac agtacgaaac
     4981 cctggaaaat cagctcgcgt tcctgtgtca gcaaggcttc tccctggaga acgcactgta
     5041 cgctctgtcc gccgtgggcc actttacact gggctgcgta ttggaggaac aggagcatca
     5101 agtagcaaaa gaggaaagag agacacctac caccgattct atgcccccac ttctgaaaca
     5161 agcaattgag ctgttcgacc ggcagggagc cgaacctgcc ttccttttcg gcctggaact
     5221 aatcatatgt ggcctggaga aacagctaaa gtgcgaaagc ggcgggccga ccgacgccct
     5281 tgacgatttt gacttagaca tgctcccagc cgatgccctt gacgactttg accttgatat
     5341 gctgcctgct gacgctcttg acgattttga ccttgacatg ctccccgggt aagctagcaa
     5401 cttgtttatt gcagcttata atggttacaa ataaagcaat agcatcacaa atttcacaaa
     5461 taaagcattt ttttcactgc attctagttg tggtttgtcc aaactcatca atgtatctta
     5521 tgtcctcaca ggaacgaagt ccctaaagaa acagtggcag ccaggtttag ccccggaatt
     5581 gactggattc cttttttagg gcccattggt atggcttttt ccccgtatcc ccccaggtgt
     5641 ctgcaggctc aaagagcagc gagaagcgtt cagaggaaag cgatcccgtg ccaccttccc
     5701 cgtgcccggg ctgtccccgc acgctgccgg ctcggggatg cggggggagc gccggaccgg
     5761 agcggagccc cgggcggctc gctgctgccc cctagcgggg gagggacgta attacatccc
     5821 tgggggcttt gggggggggc tgtccctctt gaacgaccgc caccgcggtg gagctccagc
     5881 ttttgttccc tttagtgagg gttaattaaa tcttaatacg actcactata gggcgaattg
     5941 ggaaccgggc cccccctgga gatcgacggt atccataagc ttgatatcta taacaagaaa
     6001 atatatatat aataagttat cacgtaagta gaacatgaaa taacaatata attatcgtat
     6061 gagttaaatc ttaaaagtca cgtaaaagat aatcatgcgt cattttgact cacgcggtcg
     6121 ttatagttca aaatcagtga cacttaccgc attgacaagc acgcctcacg ggagctccaa
     6181 gcggcgactg agatgtccta aatgcacagc gacggattcg cgctatttag aaagagagag
     6241 caatatttca agaatgcatg cgtcaatttt acgcagacta tctttctagg gttaatctag
     6301 ctgcatcagg atcatagcgg ccgccagctt ggcgtaatca tggtcatagc tgtttcctgt
     6361 gtgaaattgt tatccgctca caattccaca caacatacga gccggaagca taaagtgtaa
     6421 agcctggggt gcctaatgag tgagctaact cacattaatt gcgttgcgct cactgcccgc
     6481 tttccagtcg ggaaacctgt cgtgccagct gcattaatga atcggccaac gcgcggggag
     6541 aggcggtttg cgtattgggc gctcttccgc ttcctcgctc actgactcgc tgcgctcggt
     6601 cgttcggctg cggcgagcgg tatcagctca ctcaaaggcg gtaatacggt tatccacaga
     6661 atcaggggat aacgcaggaa agaacatgtg agcaaaaggc cagcaaaagg ccaggaaccg
     6721 taaaaaggcc gcgttgctgg cgtttttcca taggctccgc ccccctgacg agcatcacaa
     6781 aaatcgacgc tcaagtcaga ggtggcgaaa cccgacagga ctataaagat accaggcgtt
     6841 tccccctgga agctccctcg tgcgctctcc tgttccgacc ctgccgctta ccggatacct
     6901 gtccgccttt ctcccttcgg gaagcgtggc gctttctcat agctcacgct gtaggtatct
     6961 cagttcggtg taggtcgttc gctccaagct gggctgtgtg cacgaacccc ccgttcagcc
     7021 cgaccgctgc gccttatccg gtaactatcg tcttgagtcc aacccggtaa gacacgactt
     7081 atcgccactg gcagcagcca ctggtaacag gattagcaga gcgaggtatg taggcggtgc
     7141 tacagagttc ttgaagtggt ggcctaacta cggctacact agaagaacag tatttggtat
     7201 ctgcgctctg ctgaagccag ttaccttcgg aaaaagagtt ggtagctctt gatccggcaa
     7261 acaaaccacc gctggtagcg gtggtttttt tgtttgcaag cagcagatta cgcgcagaaa
     7321 aaaaggatct caagaagatc ctttgatctt ttctacgggg tctgacgctc agtggaacga
     7381 aaactcacgt taagggattt tggtcatgag attatcaaaa aggatcttca cctagatcct
     7441 tttaaattaa aaatgaagtt ttaaatcaat ctaaagtata tatgagtaaa cttggtctga
     7501 cagttaccaa tgcttaatca gtgaggcacc tatctcagcg atctgtctat ttcgttcatc
     7561 catagttgcc tgactccccg tcgtgtagat aactacgata cgggagggct taccatctgg
     7621 ccccagtgct gcaatgatac cgcgagaccc acgctcaccg gctccagatt tatcagcaat
     7681 aaaccagcca gccggaaggg ccgagcgcag aagtggtcct gcaactttat ccgcctccat
     7741 ccagtctatt aattgttgcc gggaagctag agtaagtagt tcgccagtta atagtttgcg
     7801 caacgttgtt gccattgcta caggcatcgt ggtgtcacgc tcgtcgtttg gtatggcttc
     7861 attcagctcc ggttcccaac gatcaaggcg agttacatga tcccccatgt tgtgcaaaaa
     7921 agcggttagc tccttcggtc ctccgatcgt tgtcagaagt aagttggccg cagtgttatc
     7981 actcatggtt atggcagcac tgcataattc tcttactgtc atgccatccg taagatgctt
     8041 ttctgtgact ggtgagtact caaccaagtc attctgagaa tagtgtatgc ggcgaccgag
     8101 ttgctcttgc ccggcgtcaa tacgggataa taccgcgcca catagcagaa ctttaaaagt
     8161 gctcatcatt ggaaaacgtt cttcggggcg aaaactctca aggatcttac cgctgttgag
     8221 atccagttcg atgtaaccca ctcgtgcacc caactgatct tcagcatctt ttactttcac
     8281 cagcgtttct gggtgagcaa aaacaggaag gcaaaatgcc gcaaaaaagg gaataagggc
     8341 gacacggaaa tgttgaatac tcatactctt cctttttcaa tattattgaa gcatttatca
     8401 gggttattgt ctcatgagcg gatacatatt tgaatgtatt tagaaaaata aacaaatagg
     8461 ggttccgcgc acatttcccc gaaaagtgcc acctgacgtc taagaaacca ttattatcat
     8521 gacattaacc tataaaaata ggcgtatcac gaggcccttt cgtccagcac tgcataattc
     8581 tcttactgtc atgccatccg taagatgctt ttctgtgact ggtgagtact caaccaagtc
     8641 attctgagaa tagtgtatgc ggcgaccgag ttgctcttgc ccggcgtcaa tacgggataa
     8701 taccgcgcca catagcagaa ctttaaaagt gctcatcatt ggaaaacgtt cttcggggcg
     8761 aaaactctca aggatcttac cgctgttgag atccagttcg atgtaaccca ctcgtgcacc
     8821 caactgatct tcagcatctt ttactttcac cagcgtttct gggtgagcaa aaacaggaag
     8881 gcaaaatgcc gcaaaaaagg gaataagggc gacacggaaa tgttgaatac tcatactctt
     8941 cctttttcaa tattattgaa gcatttatca gggttattgt ctcatgagcg gatacatatt
     9001 tgaatgtatt tagaaaaata aacaaatagg ggttccgcgc acatttcccc gaaaagtgcc
     9061 acctgacgtc taagaaacca ttattatcat gacattaacc tataaaaata ggcgtatcac
     9121 gaggcccttt cgtc
//

Lenti plKO_EF1A_ORF_HLA_A*01 01

LOCUS plKO_EF1A_ORF_HLA_A*01 01               8739 bp DNA     circular SYN 24-OCT-2022
DEFINITION  synthetic circular DNA
ACCESSION   .
VERSION     .
KEYWORDS    .
SOURCE      synthetic DNA construct
  ORGANISM  synthetic DNA construct
REFERENCE   1  (bases 1 to 8739)
  AUTHORS   .
  TITLE     Direct Submission
  JOURNAL   Exported Oct 25, 2022 from SnapGene 6.0.5
            https://www.snapgene.com
COMMENT     U515NEI230-112 DNA889095
FEATURES             Location/Qualifiers
     source          1..8739
                     /mol_type="other DNA"
                     /organism="synthetic DNA construct"
     misc_feature    10..598
                     /label=WPRE
     CDS             complement(481..492)
                     /label=Factor Xa site
     LTR             670..903
                     /label=3' LTR (Delta-U3)
     polyA_signal    981..1102
                     /label=SV40 poly(A) signal
     rep_origin      1142..1277
                     /label=SV40 ori
     promoter        complement(1298..1316)
                     /label=T7 promoter
     primer_bind     complement(1326..1342)
                     /label=M13 fwd
     rep_origin      1484..1939
                     /label=f1 ori
     promoter        1965..2069
                     /label=AmpR promoter
     CDS             2070..2138
                     /label=AmpR
     CDS             2139..2930
                     /label=AmpR
     rep_origin      3101..3689
                     /label=ori
     promoter        4019..4036
                     /label=lac promoter
     protein_bind    4051..4067
                     /label=lac operator
     primer_bind     4075..4091
                     /label=M13 rev
     promoter        4112..4130
                     /label=T3 promoter
     promoter        4158..4384
                     /label=RSV promoter
     LTR             4385..4565
                     /label=5' LTR (truncated)
     misc_feature    4612..4737
                     /label=HIV-1 Psi
     misc_feature    5230..5463
                     /label=RRE
     misc_feature    5959..6076
                     /label=cPPT/CTS
     promoter        6236..7417
                     /label=EF-1-alpha promoter
     intron          6466..7408
                     /label=EF-1-alpha intron A
     regulatory      7437..7446
                     /label=Kozak sequence
     CDS             7443..8540
                     /codon_start=1
                     /label=HLA-A*01:01
                     /label=Translation 3449-4546
                     /translation="MAVMAPRTLLLLLSGALALTQTWAGSHSMRYFFTSVSRPGRGEPR
                     FIAVGYVDDTQFVRFDSDAASQKMEPRAPWIEQEGPEYWDQETRNMKAHSQTDRANLGT
                     LRGYYNQSEDGSHTIQIMYGCDVGPDGRFLRGYRQDAYDGKDYIALNEDLRSWTAADMA
                     AQITKRKWEAVHAAEQRRVYLEGRCVDGLRRYLENGKETLQRTDPPKTHMTHHPISDHE
                     ATLRCWALGFYPAEITLTWQRDGEDQTQDTELVETRPAGDGTFQKWAAVVVPSGEEQRY
                     TCHVQHEGLPKPLTLRWELSSQPTIPIVGIIAGLVLLGAVITGAVVAAVMWRRKSSDRK
                     GGSYTQAASSDSAQGSDVSLTACKV"
     misc_RNA        complement(8588..8663)
                     /label=gRNA scaffold/Barcode
ORIGIN
        1 taagtcgaca atcaacctct ggattacaaa atttgtgaaa gattgactgg tattcttaac
       61 tatgttgctc cttttacgct atgtggatac gctgctttaa tgcctttgta tcatgctatt
      121 gcttcccgta tggctttcat tttctcctcc ttgtataaat cctggttgct gtctctttat
      181 gaggagttgt ggcccgttgt caggcaacgt ggcgtggtgt gcactgtgtt tgctgacgca
      241 acccccactg gttggggcat tgccaccacc tgtcagctcc tttccgggac tttcgctttc
      301 cccctcccta ttgccacggc ggaactcatc gccgcctgcc ttgcccgctg ctggacaggg
      361 gctcggctgt tgggcactga caattccgtg gtgttgtcgg ggaaatcatc gtcctttcct
      421 tggctgctcg cctgtgttgc cacctggatt ctgcgcggga cgtccttctg ctacgtccct
      481 tcggccctca atccagcgga ccttccttcc cgcggcctgc tgccggctct gcggcctctt
      541 ccgcgtcttc gccttcgccc tcagacgagt cggatctccc tttgggccgc ctccccgcgt
      601 cgactttaag accaatgact tacaaggcag ctgtagatct tagccacttt ttaaaagaaa
      661 aggggggact ggaagggcta attcactccc aacgaagaca agatctgctt tttgcttgta
      721 ctgggtctct ctggttagac cagatctgag cctgggagct ctctggctaa ctagggaacc
      781 cactgcttaa gcctcaataa agcttgcctt gagtgcttca agtagtgtgt gcccgtctgt
      841 tgtgtgactc tggtaactag agatccctca gaccctttta gtcagtgtgg aaaatctcta
      901 gcagtacgta tagtagttca tgtcatctta ttattcagta tttataactt gcaaagaaat
      961 gaatatcaga gagtgagagg aacttgttta ttgcagctta taatggttac aaataaagca
     1021 atagcatcac aaatttcaca aataaagcat ttttttcact gcattctagt tgtggtttgt
     1081 ccaaactcat caatgtatct tatcatgtct ggctctagct atcccgcccc taactccgcc
     1141 catcccgccc ctaactccgc ccagttccgc ccattctccg ccccatggct gactaatttt
     1201 ttttatttat gcagaggccg aggccgcctc ggcctctgag ctattccaga agtagtgagg
     1261 aggctttttt ggaggcctag ggacgtaccc aattcgccct atagtgagtc gtattacgcg
     1321 cgctcactgg ccgtcgtttt acaacgtcgt gactgggaaa accctggcgt tacccaactt
     1381 aatcgccttg cagcacatcc ccctttcgcc agctggcgta atagcgaaga ggcccgcacc
     1441 gatcgccctt cccaacagtt gcgcagcctg aatggcgaat gggacgcgcc ctgtagcggc
     1501 gcattaagcg cggcgggtgt ggtggttacg cgcagcgtga ccgctacact tgccagcgcc
     1561 ctagcgcccg ctcctttcgc tttcttccct tcctttctcg ccacgttcgc cggctttccc
     1621 cgtcaagctc taaatcgggg gctcccttta gggttccgat ttagtgcttt acggcacctc
     1681 gaccccaaaa aacttgatta gggtgatggt tcacgtagtg ggccatcgcc ctgatagacg
     1741 gtttttcgcc ctttgacgtt ggagtccacg ttctttaata gtggactctt gttccaaact
     1801 ggaacaacac tcaaccctat ctcggtctat tcttttgatt tataagggat tttgccgatt
     1861 tcggcctatt ggttaaaaaa tgagctgatt taacaaaaat ttaacgcgaa ttttaacaaa
     1921 atattaacgc ttacaattta ggtggcactt ttcggggaaa tgtgcgcgga acccctattt
     1981 gtttattttt ctaaatacat tcaaatatgt atccgctcat gagacaataa ccctgataaa
     2041 tgcttcaata atattgaaaa aggaagagta tgagtattca acatttccgt gtcgccctta
     2101 ttcccttttt tgcggcattt tgccttcctg tttttgctca cccagaaacg ctggtgaaag
     2161 taaaagatgc tgaagatcag ttgggtgcac gagtgggtta catcgaactg gatctcaaca
     2221 gcggtaagat ccttgagagt tttcgccccg aagaacgttt tccaatgatg agcactttta
     2281 aagttctgct atgtggcgcg gtattatccc gtattgacgc cgggcaagag caactcggtc
     2341 gccgcataca ctattctcag aatgacttgg ttgagtactc accagtcaca gaaaagcatc
     2401 ttacggatgg catgacagta agagaattat gcagtgctgc cataaccatg agtgataaca
     2461 ctgcggccaa cttacttctg acaacgatcg gaggaccgaa ggagctaacc gcttttttgc
     2521 acaacatggg ggatcatgta actcgccttg atcgttggga accggagctg aatgaagcca
     2581 taccaaacga cgagcgtgac accacgatgc ctgtagcaat ggcaacaacg ttgcgcaaac
     2641 tattaactgg cgaactactt actctagctt cccggcaaca attaatagac tggatggagg
     2701 cggataaagt tgcaggacca cttctgcgct cggcccttcc ggctggctgg tttattgctg
     2761 ataaatctgg agccggtgag cgtgggtctc gcggtatcat tgcagcactg gggccagatg
     2821 gtaagccctc ccgtatcgta gttatctaca cgacggggag tcaggcaact atggatgaac
     2881 gaaatagaca gatcgctgag ataggtgcct cactgattaa gcattggtaa ctgtcagacc
     2941 aagtttactc atatatactt tagattgatt taaaacttca tttttaattt aaaaggatct
     3001 aggtgaagat cctttttgat aatctcatga ccaaaatccc ttaacgtgag ttttcgttcc
     3061 actgagcgtc agaccccgta gaaaagatca aaggatcttc ttgagatcct ttttttctgc
     3121 gcgtaatctg ctgcttgcaa acaaaaaaac caccgctacc agcggtggtt tgtttgccgg
     3181 atcaagagct accaactctt tttccgaagg taactggctt cagcagagcg cagataccaa
     3241 atactgttct tctagtgtag ccgtagttag gccaccactt caagaactct gtagcaccgc
     3301 ctacatacct cgctctgcta atcctgttac cagtggctgc tgccagtggc gataagtcgt
     3361 gtcttaccgg gttggactca agacgatagt taccggataa ggcgcagcgg tcgggctgaa
     3421 cggggggttc gtgcacacag cccagcttgg agcgaacgac ctacaccgaa ctgagatacc
     3481 tacagcgtga gctatgagaa agcgccacgc ttcccgaagg gagaaaggcg gacaggtatc
     3541 cggtaagcgg cagggtcgga acaggagagc gcacgaggga gcttccaggg ggaaacgcct
     3601 ggtatcttta tagtcctgtc gggtttcgcc acctctgact tgagcgtcga tttttgtgat
     3661 gctcgtcagg ggggcggagc ctatggaaaa acgccagcaa cgcggccttt ttacggttcc
     3721 tggccttttg ctggcctttt gctcacatgt tctttcctgc gttatcccct gattctgtgg
     3781 ataaccgtat taccgccttt gagtgagctg ataccgctcg ccgcagccga acgaccgagc
     3841 gcagcgagtc agtgagcgag gaagcggaag agcgcccaat acgcaaaccg cctctccccg
     3901 cgcgttggcc gattcattaa tgcagctggc acgacaggtt tcccgactgg aaagcgggca
     3961 gtgagcgcaa cgcaattaat gtgagttagc tcactcatta ggcaccccag gctttacact
     4021 ttatgcttcc ggctcgtatg ttgtgtggaa ttgtgagcgg ataacaattt cacacaggaa
     4081 acagctatga ccatgattac gccaagcgcg caattaaccc tcactaaagg gaacaaaagc
     4141 tggagctgca agcttaatgt agtcttatgc aatactcttg tagtcttgca acatggtaac
     4201 gatgagttag caacatgcct tacaaggaga gaaaaagcac cgtgcatgcc gattggtgga
     4261 agtaaggtgg tacgatcgtg ccttattagg aaggcaacag acgggtctga catggattgg
     4321 acgaaccact gaattgccgc attgcagaga tattgtattt aagtgcctag ctcgatacat
     4381 aaacgggtct ctctggttag accagatctg agcctgggag ctctctggct aactagggaa
     4441 cccactgctt aagcctcaat aaagcttgcc ttgagtgctt caagtagtgt gtgcccgtct
     4501 gttgtgtgac tctggtaact agagatccct cagacccttt tagtcagtgt ggaaaatctc
     4561 tagcagtggc gcccgaacag ggacttgaaa gcgaaaggga aaccagagga gctctctcga
     4621 cgcaggactc ggcttgctga agcgcgcacg gcaagaggcg aggggcggcg actggtgagt
     4681 acgccaaaaa ttttgactag cggaggctag aaggagagag atgggtgcga gagcgtcagt
     4741 attaagcggg ggagaattag atcgcgatgg gaaaaaattc ggttaaggcc agggggaaag
     4801 aaaaaatata aattaaaaca tatagtatgg gcaagcaggg agctagaacg attcgcagtt
     4861 aatcctggcc tgttagaaac atcagaaggc tgtagacaaa tactgggaca gctacaacca
     4921 tcccttcaga caggatcaga agaacttaga tcattatata atacagtagc aaccctctat
     4981 tgtgtgcatc aaaggataga gataaaagac accaaggaag ctttagacaa gatagaggaa
     5041 gagcaaaaca aaagtaagac caccgcacag caagcggccg ctgatcttca gacctggagg
     5101 aggagatatg agggacaatt ggagaagtga attatataaa tataaagtag taaaaattga
     5161 accattagga gtagcaccca ccaaggcaaa gagaagagtg gtgcagagag aaaaaagagc
     5221 agtgggaata ggagctttgt tccttgggtt cttgggagca gcaggaagca ctatgggcgc
     5281 agcgtcaatg acgctgacgg tacaggccag acaattattg tctggtatag tgcagcagca
     5341 gaacaatttg ctgagggcta ttgaggcgca acagcatctg ttgcaactca cagtctgggg
     5401 catcaagcag ctccaggcaa gaatcctggc tgtggaaaga tacctaaagg atcaacagct
     5461 cctggggatt tggggttgct ctggaaaact catttgcacc actgctgtgc cttggaatgc
     5521 tagttggagt aataaatctc tggaacagat ttggaatcac acgacctgga tggagtggga
     5581 cagagaaatt aacaattaca caagcttaat acactcctta attgaagaat cgcaaaacca
     5641 gcaagaaaag aatgaacaag aattattgga attagataaa tgggcaagtt tgtggaattg
     5701 gtttaacata acaaattggc tgtggtatat aaaattattc ataatgatag taggaggctt
     5761 ggtaggttta agaatagttt ttgctgtact ttctatagtg aatagagtta ggcagggata
     5821 ttcaccatta tcgtttcaga cccacctccc aaccccgagg ggacccgaca ggcccgaagg
     5881 aatagaagaa gaaggtggag agagagacag agacagatcc attcgattag tgaacggatc
     5941 tcgacggtat cggttaactt ttaaaagaaa aggggggatt ggggggtaca gtgcagggga
     6001 aagaatagta gacataatag caacagacat acaaactaaa gaattacaaa aacaaattac
     6061 aaaaattcaa aattttatcg atgagtaatt catacaaaag gactcgcccc tgccttgggg
     6121 aatcccaggg accgtcgtta aactcccact aacgtagaac ccagagatcg ctgcgttccc
     6181 gccccctcac ccgcccgctc tcgtcatcac tgaggtggag aagagcatgc gtgaggctcc
     6241 ggtgcccgtc agtgggcaga gcgcacatcg cccacagtcc ccgagaagtt ggggggaggg
     6301 gtcggcaatt gaaccggtgc ctagagaagg tggcgcgggg taaactggga aagtgatgtc
     6361 gtgtactggc tccgcctttt tcccgagggt gggggagaac cgtatataag tgcagtagtc
     6421 gccgtgaacg ttctttttcg caacgggttt gccgccagaa cacaggtaag tgccgtgtgt
     6481 ggttcccgcg ggcctggcct ctttacgggt tatggccctt gcgtgccttg aattacttcc
     6541 acgcccctgg ctgcagtacg tgattcttga tcccgagctt cgggttggaa gtgggtggga
     6601 gagttcgagg ccttgcgctt aaggagcccc ttcgcctcgt gcttgagttg aggcctggct
     6661 tgggcgctgg ggccgccgcg tgcgaatctg gtggcacctt cgcgcctgtc tcgctgcttt
     6721 cgataagtct ctagccattt aaaatttttg atgacctgct gcgacgcttt ttttctggca
     6781 agatagtctt gtaaatgcgg gccaagatct gcacactggt atttcggttt ttggggccgc
     6841 gggcggcgac ggggcccgtg cgtcccagcg cacatgttcg gcgaggcggg gcctgcgagc
     6901 gcggccaccg agaatcggac gggggtagtc tcaagctggc cggcctgctc tggtgcctgg
     6961 cctcgcgccg ccgtgtatcg ccccgccctg ggcggcaagg ctggcccggt cggcaccagt
     7021 tgcgtgagcg gaaagatggc cgcttcccgg ccctgctgca gggagctcaa aatggaggac
     7081 gcggcgctcg ggagagcggg cgggtgagtc acccacacaa aggaaaaggg cctttccgtc
     7141 ctcagccgtc gcttcatgtg actccacgga gtaccgggcg ccgtccaggc acctcgatta
     7201 gttctcaagc ttttggagta cgtcgtcttt aggttggggg gaggggtttt atgcgatgga
     7261 gtttccccac actgagtggg tggagactga agttaggcca gcttggcact tgatgtaatt
     7321 ctccttggaa tttgcccttt ttgagtttgg atcttggttc attctcaagc ctcagacagt
     7381 ggttcaaagt ttttttcttc catttcaggt gtcgtgaccc tagcgctacc tctagagcca
     7441 ccatggccgt gatggcccct agaaccctgc tgctgctgct gagcggcgcc ctggccctga
     7501 cccagacatg ggccggcagc cactccatga gatatttctt tacctctgtg agcaggccag
     7561 gaaggggaga gccaaggttc atcgccgtgg gctacgtgga cgatacacag ttcgtgagat
     7621 ttgacagcga tgccgcctcc cagaagatgg agcctagggc accatggatc gagcaggagg
     7681 gaccagagta ttgggatcag gagacacgga acatgaaggc ccactctcag accgacaggg
     7741 caaacctggg cacactgagg ggctactata atcagtctga ggatggcagc cacacaatcc
     7801 agatcatgta cggatgcgat gtgggaccag acggccggtt tctgagagga tacaggcagg
     7861 acgcctatga tggcaaggac tacatcgccc tgaacgagga tctgcgctcc tggaccgcag
     7921 cagacatggc agcccagatc acaaagcgga agtgggaggc agtgcacgca gcagagcagc
     7981 ggagagtgta tctggagggc cgctgcgtgg acggcctgag gcgctacctg gagaatggca
     8041 aggagacact gcagcggaca gatcccccta agacccacat gacacaccac ccaatcagcg
     8101 accacgaggc caccctgagg tgttgggcac tgggcttcta tcctgccgag atcaccctga
     8161 catggcagag ggatggagag gaccagaccc aggatacaga gctggtggag acacggcccg
     8221 caggcgacgg cacatttcag aagtgggcag cagtggtggt gccatccgga gaggagcaga
     8281 gatacacctg ccacgtgcag cacgagggcc tgccaaagcc cctgaccctg agatgggagc
     8341 tgagctccca gcccacaatc cctatcgtgg gcatcatcgc aggcctggtg ctgctgggag
     8401 ccgtgatcac aggagcagtg gtggcagccg tgatgtggag gagaaagtct agcgatagaa
     8461 agggaggatc ctatacccag gcagcatcct ctgattctgc ccagggctcc gacgtgtctc
     8521 tgacagcctg taaggtgtga ggcgcgcctc ccagagccac cgttacacct tggatgactg
     8581 tcaccgggca ccgactcggt gccacttttt caagttgata acggactagc cttattttaa
     8641 cttgctattt ctagctctaa aacttttttt gactagatct tgagacaaat ggcagtattc
     8701 atccacaagg tacgctaaac gcctgcaacc aggacgcgt
//

Lenti plKO_EF1A_ORF_HLA_A*02 01

LOCUS plKO_EF1A_ORF_HLA_A*02 01               8739 bp DNA     circular SYN 24-OCT-2022
DEFINITION  synthetic circular DNA
ACCESSION   .
VERSION     .
KEYWORDS    .
SOURCE      synthetic DNA construct
  ORGANISM  synthetic DNA construct
REFERENCE   1  (bases 1 to 8739)
  AUTHORS   .
  TITLE     Direct Submission
  JOURNAL   Exported Oct 25, 2022 from SnapGene 6.0.5
            https://www.snapgene.com
COMMENT     U515NEI230-114 DNA889096
FEATURES             Location/Qualifiers
     source          1..8739
                     /mol_type="other DNA"
                     /organism="synthetic DNA construct"
     misc_feature    10..598
                     /label=WPRE
     CDS             complement(481..492)
                     /label=Factor Xa site
     LTR             670..903
                     /label=3' LTR (Delta-U3)
     polyA_signal    981..1102
                     /label=SV40 poly(A) signal
     rep_origin      1142..1277
                     /label=SV40 ori
     promoter        complement(1298..1316)
                     /label=T7 promoter
     primer_bind     complement(1326..1342)
                     /label=M13 fwd
     rep_origin      1484..1939
                     /label=f1 ori
     promoter        1965..2069
                     /label=AmpR promoter
     CDS             2070..2138
                     /label=AmpR
     CDS             2139..2930
                     /label=AmpR
     rep_origin      3101..3689
                     /label=ori
     promoter        4019..4036
                     /label=lac promoter
     protein_bind    4051..4067
                     /label=lac operator
     primer_bind     4075..4091
                     /label=M13 rev
     promoter        4112..4130
                     /label=T3 promoter
     promoter        4158..4384
                     /label=RSV promoter
     LTR             4385..4565
                     /label=5' LTR (truncated)
     misc_feature    4612..4737
                     /label=HIV-1 Psi
     misc_feature    5230..5463
                     /label=RRE
     misc_feature    5959..6076
                     /label=cPPT/CTS
     promoter        6236..7417
                     /label=EF-1-alpha promoter
     intron          6466..7408
                     /label=EF-1-alpha intron A
     regulatory      7437..7446
                     /label=Kozak sequence
     CDS             7443..8540
                     /codon_start=1
                     /label=HLA-A*02:01
                     /translation="MAVMAPRTLVLLLSGALALTQTWAGSHSMRYFFTSVSRPGRGEPR
                     FIAVGYVDDTQFVRFDSDAASQRMEPRAPWIEQEGPEYWDGETRKVKAHSQTHRVDLGT
                     LRGYYNQSEAGSHTVQRMYGCDVGSDWRFLRGYHQYAYDGKDYIALKEDLRSWTAADMA
                     AQTTKHKWEAAHVAEQLRAYLEGTCVEWLRRYLENGKETLQRTDAPKTHMTHHAVSDHE
                     ATLRCWALSFYPAEITLTWQRDGEDQTQDTELVETRPAGDGTFQKWAAVVVPSGQEQRY
                     TCHVQHEGLPKPLTLRWEPSSQPTIPIVGIIAGLVLFGAVITGAVVAAVMWRRKSSDRK
                     GGSYSQAASSDSAQGSDVSLTACKV"
     misc_RNA        complement(8588..8663)
                     /label=gRNA scaffold/Barcode
ORIGIN
        1 taagtcgaca atcaacctct ggattacaaa atttgtgaaa gattgactgg tattcttaac
       61 tatgttgctc cttttacgct atgtggatac gctgctttaa tgcctttgta tcatgctatt
      121 gcttcccgta tggctttcat tttctcctcc ttgtataaat cctggttgct gtctctttat
      181 gaggagttgt ggcccgttgt caggcaacgt ggcgtggtgt gcactgtgtt tgctgacgca
      241 acccccactg gttggggcat tgccaccacc tgtcagctcc tttccgggac tttcgctttc
      301 cccctcccta ttgccacggc ggaactcatc gccgcctgcc ttgcccgctg ctggacaggg
      361 gctcggctgt tgggcactga caattccgtg gtgttgtcgg ggaaatcatc gtcctttcct
      421 tggctgctcg cctgtgttgc cacctggatt ctgcgcggga cgtccttctg ctacgtccct
      481 tcggccctca atccagcgga ccttccttcc cgcggcctgc tgccggctct gcggcctctt
      541 ccgcgtcttc gccttcgccc tcagacgagt cggatctccc tttgggccgc ctccccgcgt
      601 cgactttaag accaatgact tacaaggcag ctgtagatct tagccacttt ttaaaagaaa
      661 aggggggact ggaagggcta attcactccc aacgaagaca agatctgctt tttgcttgta
      721 ctgggtctct ctggttagac cagatctgag cctgggagct ctctggctaa ctagggaacc
      781 cactgcttaa gcctcaataa agcttgcctt gagtgcttca agtagtgtgt gcccgtctgt
      841 tgtgtgactc tggtaactag agatccctca gaccctttta gtcagtgtgg aaaatctcta
      901 gcagtacgta tagtagttca tgtcatctta ttattcagta tttataactt gcaaagaaat
      961 gaatatcaga gagtgagagg aacttgttta ttgcagctta taatggttac aaataaagca
     1021 atagcatcac aaatttcaca aataaagcat ttttttcact gcattctagt tgtggtttgt
     1081 ccaaactcat caatgtatct tatcatgtct ggctctagct atcccgcccc taactccgcc
     1141 catcccgccc ctaactccgc ccagttccgc ccattctccg ccccatggct gactaatttt
     1201 ttttatttat gcagaggccg aggccgcctc ggcctctgag ctattccaga agtagtgagg
     1261 aggctttttt ggaggcctag ggacgtaccc aattcgccct atagtgagtc gtattacgcg
     1321 cgctcactgg ccgtcgtttt acaacgtcgt gactgggaaa accctggcgt tacccaactt
     1381 aatcgccttg cagcacatcc ccctttcgcc agctggcgta atagcgaaga ggcccgcacc
     1441 gatcgccctt cccaacagtt gcgcagcctg aatggcgaat gggacgcgcc ctgtagcggc
     1501 gcattaagcg cggcgggtgt ggtggttacg cgcagcgtga ccgctacact tgccagcgcc
     1561 ctagcgcccg ctcctttcgc tttcttccct tcctttctcg ccacgttcgc cggctttccc
     1621 cgtcaagctc taaatcgggg gctcccttta gggttccgat ttagtgcttt acggcacctc
     1681 gaccccaaaa aacttgatta gggtgatggt tcacgtagtg ggccatcgcc ctgatagacg
     1741 gtttttcgcc ctttgacgtt ggagtccacg ttctttaata gtggactctt gttccaaact
     1801 ggaacaacac tcaaccctat ctcggtctat tcttttgatt tataagggat tttgccgatt
     1861 tcggcctatt ggttaaaaaa tgagctgatt taacaaaaat ttaacgcgaa ttttaacaaa
     1921 atattaacgc ttacaattta ggtggcactt ttcggggaaa tgtgcgcgga acccctattt
     1981 gtttattttt ctaaatacat tcaaatatgt atccgctcat gagacaataa ccctgataaa
     2041 tgcttcaata atattgaaaa aggaagagta tgagtattca acatttccgt gtcgccctta
     2101 ttcccttttt tgcggcattt tgccttcctg tttttgctca cccagaaacg ctggtgaaag
     2161 taaaagatgc tgaagatcag ttgggtgcac gagtgggtta catcgaactg gatctcaaca
     2221 gcggtaagat ccttgagagt tttcgccccg aagaacgttt tccaatgatg agcactttta
     2281 aagttctgct atgtggcgcg gtattatccc gtattgacgc cgggcaagag caactcggtc
     2341 gccgcataca ctattctcag aatgacttgg ttgagtactc accagtcaca gaaaagcatc
     2401 ttacggatgg catgacagta agagaattat gcagtgctgc cataaccatg agtgataaca
     2461 ctgcggccaa cttacttctg acaacgatcg gaggaccgaa ggagctaacc gcttttttgc
     2521 acaacatggg ggatcatgta actcgccttg atcgttggga accggagctg aatgaagcca
     2581 taccaaacga cgagcgtgac accacgatgc ctgtagcaat ggcaacaacg ttgcgcaaac
     2641 tattaactgg cgaactactt actctagctt cccggcaaca attaatagac tggatggagg
     2701 cggataaagt tgcaggacca cttctgcgct cggcccttcc ggctggctgg tttattgctg
     2761 ataaatctgg agccggtgag cgtgggtctc gcggtatcat tgcagcactg gggccagatg
     2821 gtaagccctc ccgtatcgta gttatctaca cgacggggag tcaggcaact atggatgaac
     2881 gaaatagaca gatcgctgag ataggtgcct cactgattaa gcattggtaa ctgtcagacc
     2941 aagtttactc atatatactt tagattgatt taaaacttca tttttaattt aaaaggatct
     3001 aggtgaagat cctttttgat aatctcatga ccaaaatccc ttaacgtgag ttttcgttcc
     3061 actgagcgtc agaccccgta gaaaagatca aaggatcttc ttgagatcct ttttttctgc
     3121 gcgtaatctg ctgcttgcaa acaaaaaaac caccgctacc agcggtggtt tgtttgccgg
     3181 atcaagagct accaactctt tttccgaagg taactggctt cagcagagcg cagataccaa
     3241 atactgttct tctagtgtag ccgtagttag gccaccactt caagaactct gtagcaccgc
     3301 ctacatacct cgctctgcta atcctgttac cagtggctgc tgccagtggc gataagtcgt
     3361 gtcttaccgg gttggactca agacgatagt taccggataa ggcgcagcgg tcgggctgaa
     3421 cggggggttc gtgcacacag cccagcttgg agcgaacgac ctacaccgaa ctgagatacc
     3481 tacagcgtga gctatgagaa agcgccacgc ttcccgaagg gagaaaggcg gacaggtatc
     3541 cggtaagcgg cagggtcgga acaggagagc gcacgaggga gcttccaggg ggaaacgcct
     3601 ggtatcttta tagtcctgtc gggtttcgcc acctctgact tgagcgtcga tttttgtgat
     3661 gctcgtcagg ggggcggagc ctatggaaaa acgccagcaa cgcggccttt ttacggttcc
     3721 tggccttttg ctggcctttt gctcacatgt tctttcctgc gttatcccct gattctgtgg
     3781 ataaccgtat taccgccttt gagtgagctg ataccgctcg ccgcagccga acgaccgagc
     3841 gcagcgagtc agtgagcgag gaagcggaag agcgcccaat acgcaaaccg cctctccccg
     3901 cgcgttggcc gattcattaa tgcagctggc acgacaggtt tcccgactgg aaagcgggca
     3961 gtgagcgcaa cgcaattaat gtgagttagc tcactcatta ggcaccccag gctttacact
     4021 ttatgcttcc ggctcgtatg ttgtgtggaa ttgtgagcgg ataacaattt cacacaggaa
     4081 acagctatga ccatgattac gccaagcgcg caattaaccc tcactaaagg gaacaaaagc
     4141 tggagctgca agcttaatgt agtcttatgc aatactcttg tagtcttgca acatggtaac
     4201 gatgagttag caacatgcct tacaaggaga gaaaaagcac cgtgcatgcc gattggtgga
     4261 agtaaggtgg tacgatcgtg ccttattagg aaggcaacag acgggtctga catggattgg
     4321 acgaaccact gaattgccgc attgcagaga tattgtattt aagtgcctag ctcgatacat
     4381 aaacgggtct ctctggttag accagatctg agcctgggag ctctctggct aactagggaa
     4441 cccactgctt aagcctcaat aaagcttgcc ttgagtgctt caagtagtgt gtgcccgtct
     4501 gttgtgtgac tctggtaact agagatccct cagacccttt tagtcagtgt ggaaaatctc
     4561 tagcagtggc gcccgaacag ggacttgaaa gcgaaaggga aaccagagga gctctctcga
     4621 cgcaggactc ggcttgctga agcgcgcacg gcaagaggcg aggggcggcg actggtgagt
     4681 acgccaaaaa ttttgactag cggaggctag aaggagagag atgggtgcga gagcgtcagt
     4741 attaagcggg ggagaattag atcgcgatgg gaaaaaattc ggttaaggcc agggggaaag
     4801 aaaaaatata aattaaaaca tatagtatgg gcaagcaggg agctagaacg attcgcagtt
     4861 aatcctggcc tgttagaaac atcagaaggc tgtagacaaa tactgggaca gctacaacca
     4921 tcccttcaga caggatcaga agaacttaga tcattatata atacagtagc aaccctctat
     4981 tgtgtgcatc aaaggataga gataaaagac accaaggaag ctttagacaa gatagaggaa
     5041 gagcaaaaca aaagtaagac caccgcacag caagcggccg ctgatcttca gacctggagg
     5101 aggagatatg agggacaatt ggagaagtga attatataaa tataaagtag taaaaattga
     5161 accattagga gtagcaccca ccaaggcaaa gagaagagtg gtgcagagag aaaaaagagc
     5221 agtgggaata ggagctttgt tccttgggtt cttgggagca gcaggaagca ctatgggcgc
     5281 agcgtcaatg acgctgacgg tacaggccag acaattattg tctggtatag tgcagcagca
     5341 gaacaatttg ctgagggcta ttgaggcgca acagcatctg ttgcaactca cagtctgggg
     5401 catcaagcag ctccaggcaa gaatcctggc tgtggaaaga tacctaaagg atcaacagct
     5461 cctggggatt tggggttgct ctggaaaact catttgcacc actgctgtgc cttggaatgc
     5521 tagttggagt aataaatctc tggaacagat ttggaatcac acgacctgga tggagtggga
     5581 cagagaaatt aacaattaca caagcttaat acactcctta attgaagaat cgcaaaacca
     5641 gcaagaaaag aatgaacaag aattattgga attagataaa tgggcaagtt tgtggaattg
     5701 gtttaacata acaaattggc tgtggtatat aaaattattc ataatgatag taggaggctt
     5761 ggtaggttta agaatagttt ttgctgtact ttctatagtg aatagagtta ggcagggata
     5821 ttcaccatta tcgtttcaga cccacctccc aaccccgagg ggacccgaca ggcccgaagg
     5881 aatagaagaa gaaggtggag agagagacag agacagatcc attcgattag tgaacggatc
     5941 tcgacggtat cggttaactt ttaaaagaaa aggggggatt ggggggtaca gtgcagggga
     6001 aagaatagta gacataatag caacagacat acaaactaaa gaattacaaa aacaaattac
     6061 aaaaattcaa aattttatcg atgagtaatt catacaaaag gactcgcccc tgccttgggg
     6121 aatcccaggg accgtcgtta aactcccact aacgtagaac ccagagatcg ctgcgttccc
     6181 gccccctcac ccgcccgctc tcgtcatcac tgaggtggag aagagcatgc gtgaggctcc
     6241 ggtgcccgtc agtgggcaga gcgcacatcg cccacagtcc ccgagaagtt ggggggaggg
     6301 gtcggcaatt gaaccggtgc ctagagaagg tggcgcgggg taaactggga aagtgatgtc
     6361 gtgtactggc tccgcctttt tcccgagggt gggggagaac cgtatataag tgcagtagtc
     6421 gccgtgaacg ttctttttcg caacgggttt gccgccagaa cacaggtaag tgccgtgtgt
     6481 ggttcccgcg ggcctggcct ctttacgggt tatggccctt gcgtgccttg aattacttcc
     6541 acgcccctgg ctgcagtacg tgattcttga tcccgagctt cgggttggaa gtgggtggga
     6601 gagttcgagg ccttgcgctt aaggagcccc ttcgcctcgt gcttgagttg aggcctggct
     6661 tgggcgctgg ggccgccgcg tgcgaatctg gtggcacctt cgcgcctgtc tcgctgcttt
     6721 cgataagtct ctagccattt aaaatttttg atgacctgct gcgacgcttt ttttctggca
     6781 agatagtctt gtaaatgcgg gccaagatct gcacactggt atttcggttt ttggggccgc
     6841 gggcggcgac ggggcccgtg cgtcccagcg cacatgttcg gcgaggcggg gcctgcgagc
     6901 gcggccaccg agaatcggac gggggtagtc tcaagctggc cggcctgctc tggtgcctgg
     6961 cctcgcgccg ccgtgtatcg ccccgccctg ggcggcaagg ctggcccggt cggcaccagt
     7021 tgcgtgagcg gaaagatggc cgcttcccgg ccctgctgca gggagctcaa aatggaggac
     7081 gcggcgctcg ggagagcggg cgggtgagtc acccacacaa aggaaaaggg cctttccgtc
     7141 ctcagccgtc gcttcatgtg actccacgga gtaccgggcg ccgtccaggc acctcgatta
     7201 gttctcaagc ttttggagta cgtcgtcttt aggttggggg gaggggtttt atgcgatgga
     7261 gtttccccac actgagtggg tggagactga agttaggcca gcttggcact tgatgtaatt
     7321 ctccttggaa tttgcccttt ttgagtttgg atcttggttc attctcaagc ctcagacagt
     7381 ggttcaaagt ttttttcttc catttcaggt gtcgtgaccc tagcgctacc tctagagcca
     7441 ccatggccgt gatggcacca cggaccctgg tgctgctgct gagcggcgcc ctggccctga
     7501 cccagacatg ggccggcagc cactccatga gatacttctt tacctctgtg agccggcccg
     7561 gaagaggaga gcctcggttc atcgccgtgg gctatgtgga cgatacacag ttcgtgagat
     7621 ttgactctga tgcagcaagc cagaggatgg agccaagagc accatggatc gagcaggagg
     7681 gacctgagta ctgggacggc gagacacgga aggtgaaggc ccactcccag acccacaggg
     7741 tggatctggg cacactgcgc ggctactata accagtccga ggccggctct cacacagtgc
     7801 agagaatgta tggctgcgac gtgggctctg attggaggtt tctgcgcggc taccaccagt
     7861 acgcctatga cggcaaggat tatatcgccc tgaaggagga cctgaggagc tggaccgcag
     7921 cagatatggc agcacagacc acaaagcaca agtgggaggc agcacacgtg gcagagcagc
     7981 tgagggccta cctggagggc acatgcgtgg agtggctgcg gagatatctg gagaatggca
     8041 aggagacact gcagaggaca gacgccccta agacccacat gacacaccac gccgtgagcg
     8101 atcacgaggc caccctgagg tgttgggcac tgtccttcta cccagccgag atcaccctga
     8161 catggcagag ggacggcgag gatcagaccc aggacacaga gctggtggag acacggcccg
     8221 caggcgatgg cacatttcag aagtgggcag cagtggtggt gccatccgga caggagcagc
     8281 ggtatacctg ccacgtgcag cacgagggcc tgccaaagcc tctgaccctg agatgggagc
     8341 ccagctccca gcctacaatc ccaatcgtgg gcatcatcgc aggcctggtg ctgttcggag
     8401 ccgtgatcac cggagcagtg gtggccgccg tgatgtggag gcgcaagtct agcgacagga
     8461 agggaggatc ctactctcag gcagcatcct ctgactctgc ccagggaagc gacgtgagcc
     8521 tgacagcctg taaggtgtga ggcgcgcctc ccagagccac cgttacacgt gggcacagct
     8581 atacctcgca ccgactcggt gccacttttt caagttgata acggactagc cttattttaa
     8641 cttgctattt ctagctctaa aacttttttt gactagatct tgagacaaat ggcagtattc
     8701 atccacaagg tacgctaaac gcctgcaacc aggacgcgt
//

Lenti plKO_EF1A_ORF_HLA_A*03 01

LOCUS plKO_EF1A_ORF_HLA_A*03 01                8739 bp DNA     circular SYN 24-OCT-2022
DEFINITION  synthetic circular DNA
ACCESSION   .
VERSION     .
KEYWORDS    .
SOURCE      synthetic DNA construct
  ORGANISM  synthetic DNA construct
REFERENCE   1  (bases 1 to 8739)
  AUTHORS   .
  TITLE     Direct Submission
  JOURNAL   Exported Oct 25, 2022 from SnapGene 6.0.5
            https://www.snapgene.com
COMMENT     U515NEI230-122 DNA889097
FEATURES             Location/Qualifiers
     source          1..8739
                     /mol_type="other DNA"
                     /organism="synthetic DNA construct"
     misc_feature    10..598
                     /label=WPRE
     CDS             complement(481..492)
                     /label=Factor Xa site
     LTR             670..903
                     /label=3' LTR (Delta-U3)
     polyA_signal    981..1102
                     /label=SV40 poly(A) signal
     rep_origin      1142..1277
                     /label=SV40 ori
     promoter        complement(1298..1316)
                     /label=T7 promoter
     primer_bind     complement(1326..1342)
                     /label=M13 fwd
     rep_origin      1484..1939
                     /label=f1 ori
     promoter        1965..2069
                     /label=AmpR promoter
     CDS             2070..2138
                     /label=AmpR
     CDS             2139..2930
                     /label=AmpR
     rep_origin      3101..3689
                     /label=ori
     promoter        4019..4036
                     /label=lac promoter
     protein_bind    4051..4067
                     /label=lac operator
     primer_bind     4075..4091
                     /label=M13 rev
     promoter        4112..4130
                     /label=T3 promoter
     promoter        4158..4384
                     /label=RSV promoter
     LTR             4385..4565
                     /label=5' LTR (truncated)
     misc_feature    4612..4737
                     /label=HIV-1 Psi
     misc_feature    5230..5463
                     /label=RRE
     misc_feature    5959..6076
                     /label=cPPT/CTS
     promoter        6236..7417
                     /label=EF-1-alpha promoter
     intron          6466..7408
                     /label=EF-1-alpha intron A
     regulatory      7437..7446
                     /label=Kozak sequence
     CDS             7443..8540
                     /codon_start=1
                     /label=HLA-A*03:01
                     /translation="MAVMAPRTLLLLLSGALALTQTWAGSHSMRYFFTSVSRPGRGEPR
                     FIAVGYVDDTQFVRFDSDAASQRMEPRAPWIEQEGPEYWDQETRNVKAQSQTDRVDLGT
                     LRGYYNQSEAGSHTIQIMYGCDVGSDGRFLRGYRQDAYDGKDYIALNEDLRSWTAADMA
                     AQITKRKWEAAHEAEQLRAYLDGTCVEWLRRYLENGKETLQRTDPPKTHMTHHPISDHE
                     ATLRCWALGFYPAEITLTWQRDGEDQTQDTELVETRPAGDGTFQKWAAVVVPSGEEQRY
                     TCHVQHEGLPKPLTLRWELSSQPTIPIVGIIAGLVLLGAVITGAVVAAVMWRRKSSDRK
                     GGSYTQAASSDSAQGSDVSLTACKV"
     misc_RNA        complement(8588..8663)
                     /label=gRNA scaffold/Barcode
ORIGIN
        1 taagtcgaca atcaacctct ggattacaaa atttgtgaaa gattgactgg tattcttaac
       61 tatgttgctc cttttacgct atgtggatac gctgctttaa tgcctttgta tcatgctatt
      121 gcttcccgta tggctttcat tttctcctcc ttgtataaat cctggttgct gtctctttat
      181 gaggagttgt ggcccgttgt caggcaacgt ggcgtggtgt gcactgtgtt tgctgacgca
      241 acccccactg gttggggcat tgccaccacc tgtcagctcc tttccgggac tttcgctttc
      301 cccctcccta ttgccacggc ggaactcatc gccgcctgcc ttgcccgctg ctggacaggg
      361 gctcggctgt tgggcactga caattccgtg gtgttgtcgg ggaaatcatc gtcctttcct
      421 tggctgctcg cctgtgttgc cacctggatt ctgcgcggga cgtccttctg ctacgtccct
      481 tcggccctca atccagcgga ccttccttcc cgcggcctgc tgccggctct gcggcctctt
      541 ccgcgtcttc gccttcgccc tcagacgagt cggatctccc tttgggccgc ctccccgcgt
      601 cgactttaag accaatgact tacaaggcag ctgtagatct tagccacttt ttaaaagaaa
      661 aggggggact ggaagggcta attcactccc aacgaagaca agatctgctt tttgcttgta
      721 ctgggtctct ctggttagac cagatctgag cctgggagct ctctggctaa ctagggaacc
      781 cactgcttaa gcctcaataa agcttgcctt gagtgcttca agtagtgtgt gcccgtctgt
      841 tgtgtgactc tggtaactag agatccctca gaccctttta gtcagtgtgg aaaatctcta
      901 gcagtacgta tagtagttca tgtcatctta ttattcagta tttataactt gcaaagaaat
      961 gaatatcaga gagtgagagg aacttgttta ttgcagctta taatggttac aaataaagca
     1021 atagcatcac aaatttcaca aataaagcat ttttttcact gcattctagt tgtggtttgt
     1081 ccaaactcat caatgtatct tatcatgtct ggctctagct atcccgcccc taactccgcc
     1141 catcccgccc ctaactccgc ccagttccgc ccattctccg ccccatggct gactaatttt
     1201 ttttatttat gcagaggccg aggccgcctc ggcctctgag ctattccaga agtagtgagg
     1261 aggctttttt ggaggcctag ggacgtaccc aattcgccct atagtgagtc gtattacgcg
     1321 cgctcactgg ccgtcgtttt acaacgtcgt gactgggaaa accctggcgt tacccaactt
     1381 aatcgccttg cagcacatcc ccctttcgcc agctggcgta atagcgaaga ggcccgcacc
     1441 gatcgccctt cccaacagtt gcgcagcctg aatggcgaat gggacgcgcc ctgtagcggc
     1501 gcattaagcg cggcgggtgt ggtggttacg cgcagcgtga ccgctacact tgccagcgcc
     1561 ctagcgcccg ctcctttcgc tttcttccct tcctttctcg ccacgttcgc cggctttccc
     1621 cgtcaagctc taaatcgggg gctcccttta gggttccgat ttagtgcttt acggcacctc
     1681 gaccccaaaa aacttgatta gggtgatggt tcacgtagtg ggccatcgcc ctgatagacg
     1741 gtttttcgcc ctttgacgtt ggagtccacg ttctttaata gtggactctt gttccaaact
     1801 ggaacaacac tcaaccctat ctcggtctat tcttttgatt tataagggat tttgccgatt
     1861 tcggcctatt ggttaaaaaa tgagctgatt taacaaaaat ttaacgcgaa ttttaacaaa
     1921 atattaacgc ttacaattta ggtggcactt ttcggggaaa tgtgcgcgga acccctattt
     1981 gtttattttt ctaaatacat tcaaatatgt atccgctcat gagacaataa ccctgataaa
     2041 tgcttcaata atattgaaaa aggaagagta tgagtattca acatttccgt gtcgccctta
     2101 ttcccttttt tgcggcattt tgccttcctg tttttgctca cccagaaacg ctggtgaaag
     2161 taaaagatgc tgaagatcag ttgggtgcac gagtgggtta catcgaactg gatctcaaca
     2221 gcggtaagat ccttgagagt tttcgccccg aagaacgttt tccaatgatg agcactttta
     2281 aagttctgct atgtggcgcg gtattatccc gtattgacgc cgggcaagag caactcggtc
     2341 gccgcataca ctattctcag aatgacttgg ttgagtactc accagtcaca gaaaagcatc
     2401 ttacggatgg catgacagta agagaattat gcagtgctgc cataaccatg agtgataaca
     2461 ctgcggccaa cttacttctg acaacgatcg gaggaccgaa ggagctaacc gcttttttgc
     2521 acaacatggg ggatcatgta actcgccttg atcgttggga accggagctg aatgaagcca
     2581 taccaaacga cgagcgtgac accacgatgc ctgtagcaat ggcaacaacg ttgcgcaaac
     2641 tattaactgg cgaactactt actctagctt cccggcaaca attaatagac tggatggagg
     2701 cggataaagt tgcaggacca cttctgcgct cggcccttcc ggctggctgg tttattgctg
     2761 ataaatctgg agccggtgag cgtgggtctc gcggtatcat tgcagcactg gggccagatg
     2821 gtaagccctc ccgtatcgta gttatctaca cgacggggag tcaggcaact atggatgaac
     2881 gaaatagaca gatcgctgag ataggtgcct cactgattaa gcattggtaa ctgtcagacc
     2941 aagtttactc atatatactt tagattgatt taaaacttca tttttaattt aaaaggatct
     3001 aggtgaagat cctttttgat aatctcatga ccaaaatccc ttaacgtgag ttttcgttcc
     3061 actgagcgtc agaccccgta gaaaagatca aaggatcttc ttgagatcct ttttttctgc
     3121 gcgtaatctg ctgcttgcaa acaaaaaaac caccgctacc agcggtggtt tgtttgccgg
     3181 atcaagagct accaactctt tttccgaagg taactggctt cagcagagcg cagataccaa
     3241 atactgttct tctagtgtag ccgtagttag gccaccactt caagaactct gtagcaccgc
     3301 ctacatacct cgctctgcta atcctgttac cagtggctgc tgccagtggc gataagtcgt
     3361 gtcttaccgg gttggactca agacgatagt taccggataa ggcgcagcgg tcgggctgaa
     3421 cggggggttc gtgcacacag cccagcttgg agcgaacgac ctacaccgaa ctgagatacc
     3481 tacagcgtga gctatgagaa agcgccacgc ttcccgaagg gagaaaggcg gacaggtatc
     3541 cggtaagcgg cagggtcgga acaggagagc gcacgaggga gcttccaggg ggaaacgcct
     3601 ggtatcttta tagtcctgtc gggtttcgcc acctctgact tgagcgtcga tttttgtgat
     3661 gctcgtcagg ggggcggagc ctatggaaaa acgccagcaa cgcggccttt ttacggttcc
     3721 tggccttttg ctggcctttt gctcacatgt tctttcctgc gttatcccct gattctgtgg
     3781 ataaccgtat taccgccttt gagtgagctg ataccgctcg ccgcagccga acgaccgagc
     3841 gcagcgagtc agtgagcgag gaagcggaag agcgcccaat acgcaaaccg cctctccccg
     3901 cgcgttggcc gattcattaa tgcagctggc acgacaggtt tcccgactgg aaagcgggca
     3961 gtgagcgcaa cgcaattaat gtgagttagc tcactcatta ggcaccccag gctttacact
     4021 ttatgcttcc ggctcgtatg ttgtgtggaa ttgtgagcgg ataacaattt cacacaggaa
     4081 acagctatga ccatgattac gccaagcgcg caattaaccc tcactaaagg gaacaaaagc
     4141 tggagctgca agcttaatgt agtcttatgc aatactcttg tagtcttgca acatggtaac
     4201 gatgagttag caacatgcct tacaaggaga gaaaaagcac cgtgcatgcc gattggtgga
     4261 agtaaggtgg tacgatcgtg ccttattagg aaggcaacag acgggtctga catggattgg
     4321 acgaaccact gaattgccgc attgcagaga tattgtattt aagtgcctag ctcgatacat
     4381 aaacgggtct ctctggttag accagatctg agcctgggag ctctctggct aactagggaa
     4441 cccactgctt aagcctcaat aaagcttgcc ttgagtgctt caagtagtgt gtgcccgtct
     4501 gttgtgtgac tctggtaact agagatccct cagacccttt tagtcagtgt ggaaaatctc
     4561 tagcagtggc gcccgaacag ggacttgaaa gcgaaaggga aaccagagga gctctctcga
     4621 cgcaggactc ggcttgctga agcgcgcacg gcaagaggcg aggggcggcg actggtgagt
     4681 acgccaaaaa ttttgactag cggaggctag aaggagagag atgggtgcga gagcgtcagt
     4741 attaagcggg ggagaattag atcgcgatgg gaaaaaattc ggttaaggcc agggggaaag
     4801 aaaaaatata aattaaaaca tatagtatgg gcaagcaggg agctagaacg attcgcagtt
     4861 aatcctggcc tgttagaaac atcagaaggc tgtagacaaa tactgggaca gctacaacca
     4921 tcccttcaga caggatcaga agaacttaga tcattatata atacagtagc aaccctctat
     4981 tgtgtgcatc aaaggataga gataaaagac accaaggaag ctttagacaa gatagaggaa
     5041 gagcaaaaca aaagtaagac caccgcacag caagcggccg ctgatcttca gacctggagg
     5101 aggagatatg agggacaatt ggagaagtga attatataaa tataaagtag taaaaattga
     5161 accattagga gtagcaccca ccaaggcaaa gagaagagtg gtgcagagag aaaaaagagc
     5221 agtgggaata ggagctttgt tccttgggtt cttgggagca gcaggaagca ctatgggcgc
     5281 agcgtcaatg acgctgacgg tacaggccag acaattattg tctggtatag tgcagcagca
     5341 gaacaatttg ctgagggcta ttgaggcgca acagcatctg ttgcaactca cagtctgggg
     5401 catcaagcag ctccaggcaa gaatcctggc tgtggaaaga tacctaaagg atcaacagct
     5461 cctggggatt tggggttgct ctggaaaact catttgcacc actgctgtgc cttggaatgc
     5521 tagttggagt aataaatctc tggaacagat ttggaatcac acgacctgga tggagtggga
     5581 cagagaaatt aacaattaca caagcttaat acactcctta attgaagaat cgcaaaacca
     5641 gcaagaaaag aatgaacaag aattattgga attagataaa tgggcaagtt tgtggaattg
     5701 gtttaacata acaaattggc tgtggtatat aaaattattc ataatgatag taggaggctt
     5761 ggtaggttta agaatagttt ttgctgtact ttctatagtg aatagagtta ggcagggata
     5821 ttcaccatta tcgtttcaga cccacctccc aaccccgagg ggacccgaca ggcccgaagg
     5881 aatagaagaa gaaggtggag agagagacag agacagatcc attcgattag tgaacggatc
     5941 tcgacggtat cggttaactt ttaaaagaaa aggggggatt ggggggtaca gtgcagggga
     6001 aagaatagta gacataatag caacagacat acaaactaaa gaattacaaa aacaaattac
     6061 aaaaattcaa aattttatcg atgagtaatt catacaaaag gactcgcccc tgccttgggg
     6121 aatcccaggg accgtcgtta aactcccact aacgtagaac ccagagatcg ctgcgttccc
     6181 gccccctcac ccgcccgctc tcgtcatcac tgaggtggag aagagcatgc gtgaggctcc
     6241 ggtgcccgtc agtgggcaga gcgcacatcg cccacagtcc ccgagaagtt ggggggaggg
     6301 gtcggcaatt gaaccggtgc ctagagaagg tggcgcgggg taaactggga aagtgatgtc
     6361 gtgtactggc tccgcctttt tcccgagggt gggggagaac cgtatataag tgcagtagtc
     6421 gccgtgaacg ttctttttcg caacgggttt gccgccagaa cacaggtaag tgccgtgtgt
     6481 ggttcccgcg ggcctggcct ctttacgggt tatggccctt gcgtgccttg aattacttcc
     6541 acgcccctgg ctgcagtacg tgattcttga tcccgagctt cgggttggaa gtgggtggga
     6601 gagttcgagg ccttgcgctt aaggagcccc ttcgcctcgt gcttgagttg aggcctggct
     6661 tgggcgctgg ggccgccgcg tgcgaatctg gtggcacctt cgcgcctgtc tcgctgcttt
     6721 cgataagtct ctagccattt aaaatttttg atgacctgct gcgacgcttt ttttctggca
     6781 agatagtctt gtaaatgcgg gccaagatct gcacactggt atttcggttt ttggggccgc
     6841 gggcggcgac ggggcccgtg cgtcccagcg cacatgttcg gcgaggcggg gcctgcgagc
     6901 gcggccaccg agaatcggac gggggtagtc tcaagctggc cggcctgctc tggtgcctgg
     6961 cctcgcgccg ccgtgtatcg ccccgccctg ggcggcaagg ctggcccggt cggcaccagt
     7021 tgcgtgagcg gaaagatggc cgcttcccgg ccctgctgca gggagctcaa aatggaggac
     7081 gcggcgctcg ggagagcggg cgggtgagtc acccacacaa aggaaaaggg cctttccgtc
     7141 ctcagccgtc gcttcatgtg actccacgga gtaccgggcg ccgtccaggc acctcgatta
     7201 gttctcaagc ttttggagta cgtcgtcttt aggttggggg gaggggtttt atgcgatgga
     7261 gtttccccac actgagtggg tggagactga agttaggcca gcttggcact tgatgtaatt
     7321 ctccttggaa tttgcccttt ttgagtttgg atcttggttc attctcaagc ctcagacagt
     7381 ggttcaaagt ttttttcttc catttcaggt gtcgtgaccc tagcgctacc tctagagcca
     7441 ccatggccgt gatggcacca cgcaccctgc tgctgctgct gagcggcgcc ctggccctga
     7501 cccagacatg ggccggcagc cactccatgc ggtatttctt tacctctgtg agccggccag
     7561 gaagaggaga gccacgcttc atcgcagtgg gatacgtgga cgatacacag ttcgtgcggt
     7621 ttgacagcga tgccgcctcc cagagaatgg agcctagggc accatggatc gagcaggagg
     7681 gacctgagta ttgggatcag gagacacgga acgtgaaggc ccagtcccag accgacaggg
     7741 tggatctggg cacactgagg ggctactata atcagtctga ggccggcagc cacacaatcc
     7801 agatcatgta cggctgcgat gtgggaagcg acggcaggtt tctgagggga tacaggcagg
     7861 acgcctatga tggcaaggac tacatcgccc tgaacgagga tctgaggtcc tggaccgcag
     7921 cagacatggc agcacagatc acaaagagaa agtgggaggc agcacacgag gcagagcagc
     7981 tgagggccta tctggacgga acctgcgtgg agtggctgcg gagatacctg gagaatggca
     8041 aggagacact gcagagaaca gatcccccta agacccacat gacacaccac cccatctctg
     8101 accacgaggc cacactgagg tgttgggccc tgggcttcta tcctgccgag atcaccctga
     8161 catggcagag agatggcgag gaccagaccc aggatacaga gctggtggag acacggcccg
     8221 caggcgacgg cacatttcag aagtgggcag cagtggtggt gccatccgga gaggagcagc
     8281 ggtacacctg ccacgtgcag cacgagggcc tgccaaagcc actgaccctg agatgggagc
     8341 tgagctccca gcccacaatc cctatcgtgg gcatcatcgc cggcctggtg ctgctgggag
     8401 ccgtgatcac cggagcagtg gtggccgccg tgatgtggag gcgcaagtct agcgatagga
     8461 agggaggatc ctatacccag gcagcatcct ctgattctgc ccagggctcc gacgtgtctc
     8521 tgacagcctg taaggtgtga ggcgcgcctc ccagagccac cgttacacgc ctcctcctgc
     8581 cgccctcgca ccgactcggt gccacttttt caagttgata acggactagc cttattttaa
     8641 cttgctattt ctagctctaa aacttttttt gactagatct tgagacaaat ggcagtattc
     8701 atccacaagg tacgctaaac gcctgcaacc aggacgcgt
//

Lenti plKO_EF1A_ORF_HLA_A*11 01

LOCUS plKO_EF1A_ORF_HLA_A*11 01                8739 bp DNA     circular SYN 24-OCT-2022
DEFINITION  synthetic circular DNA
ACCESSION   .
VERSION     .
KEYWORDS    .
SOURCE      synthetic DNA construct
  ORGANISM  synthetic DNA construct
REFERENCE   1  (bases 1 to 8739)
  AUTHORS   .
  TITLE     Direct Submission
  JOURNAL   Exported Oct 25, 2022 from SnapGene 6.0.5
            https://www.snapgene.com
COMMENT     U515NEI230-124 DNA889098
FEATURES             Location/Qualifiers
     source          1..8739
                     /mol_type="other DNA"
                     /organism="synthetic DNA construct"
     misc_feature    10..598
                     /label=WPRE
     CDS             complement(481..492)
                     /label=Factor Xa site
     LTR             670..903
                     /label=3' LTR (Delta-U3)
     polyA_signal    981..1102
                     /label=SV40 poly(A) signal
     rep_origin      1142..1277
                     /label=SV40 ori
     promoter        complement(1298..1316)
                     /label=T7 promoter
     primer_bind     complement(1326..1342)
                     /label=M13 fwd
     rep_origin      1484..1939
                     /label=f1 ori
     promoter        1965..2069
                     /label=AmpR promoter
     CDS             2070..2138
                     /label=AmpR
     CDS             2139..2930
                     /label=AmpR
     rep_origin      3101..3689
                     /label=ori
     promoter        4019..4036
                     /label=lac promoter
     protein_bind    4051..4067
                     /label=lac operator
     primer_bind     4075..4091
                     /label=M13 rev
     promoter        4112..4130
                     /label=T3 promoter
     promoter        4158..4384
                     /label=RSV promoter
     LTR             4385..4565
                     /label=5' LTR (truncated)
     misc_feature    4612..4737
                     /label=HIV-1 Psi
     misc_feature    5230..5463
                     /label=RRE
     misc_feature    5959..6076
                     /label=cPPT/CTS
     promoter        6236..7417
                     /label=EF-1-alpha promoter
     intron          6466..7408
                     /label=EF-1-alpha intron A
     regulatory      7437..7446
                     /label=Kozak sequence
     CDS             7443..8540
                     /codon_start=1
                     /label=HLA-A*11:01
                     /translation="MAVMAPRTLLLLLSGALALTQTWAGSHSMRYFYTSVSRPGRGEPR
                     FIAVGYVDDTQFVRFDSDAASQRMEPRAPWIEQEGPEYWDQETRNVKAQSQTDRVDLGT
                     LRGYYNQSEDGSHTIQIMYGCDVGPDGRFLRGYRQDAYDGKDYIALNEDLRSWTAADMA
                     AQITKRKWEAAHAAEQQRAYLEGRCVEWLRRYLENGKETLQRTDPPKTHMTHHPISDHE
                     ATLRCWALGFYPAEITLTWQRDGEDQTQDTELVETRPAGDGTFQKWAAVVVPSGEEQRY
                     TCHVQHEGLPKPLTLRWELSSQPTIPIVGIIAGLVLLGAVITGAVVAAVMWRRKSSDRK
                     GGSYTQAASSDSAQGSDVSLTACKV"
     misc_RNA        complement(8588..8663)
                     /label=gRNA scaffold/Barcode
ORIGIN
        1 taagtcgaca atcaacctct ggattacaaa atttgtgaaa gattgactgg tattcttaac
       61 tatgttgctc cttttacgct atgtggatac gctgctttaa tgcctttgta tcatgctatt
      121 gcttcccgta tggctttcat tttctcctcc ttgtataaat cctggttgct gtctctttat
      181 gaggagttgt ggcccgttgt caggcaacgt ggcgtggtgt gcactgtgtt tgctgacgca
      241 acccccactg gttggggcat tgccaccacc tgtcagctcc tttccgggac tttcgctttc
      301 cccctcccta ttgccacggc ggaactcatc gccgcctgcc ttgcccgctg ctggacaggg
      361 gctcggctgt tgggcactga caattccgtg gtgttgtcgg ggaaatcatc gtcctttcct
      421 tggctgctcg cctgtgttgc cacctggatt ctgcgcggga cgtccttctg ctacgtccct
      481 tcggccctca atccagcgga ccttccttcc cgcggcctgc tgccggctct gcggcctctt
      541 ccgcgtcttc gccttcgccc tcagacgagt cggatctccc tttgggccgc ctccccgcgt
      601 cgactttaag accaatgact tacaaggcag ctgtagatct tagccacttt ttaaaagaaa
      661 aggggggact ggaagggcta attcactccc aacgaagaca agatctgctt tttgcttgta
      721 ctgggtctct ctggttagac cagatctgag cctgggagct ctctggctaa ctagggaacc
      781 cactgcttaa gcctcaataa agcttgcctt gagtgcttca agtagtgtgt gcccgtctgt
      841 tgtgtgactc tggtaactag agatccctca gaccctttta gtcagtgtgg aaaatctcta
      901 gcagtacgta tagtagttca tgtcatctta ttattcagta tttataactt gcaaagaaat
      961 gaatatcaga gagtgagagg aacttgttta ttgcagctta taatggttac aaataaagca
     1021 atagcatcac aaatttcaca aataaagcat ttttttcact gcattctagt tgtggtttgt
     1081 ccaaactcat caatgtatct tatcatgtct ggctctagct atcccgcccc taactccgcc
     1141 catcccgccc ctaactccgc ccagttccgc ccattctccg ccccatggct gactaatttt
     1201 ttttatttat gcagaggccg aggccgcctc ggcctctgag ctattccaga agtagtgagg
     1261 aggctttttt ggaggcctag ggacgtaccc aattcgccct atagtgagtc gtattacgcg
     1321 cgctcactgg ccgtcgtttt acaacgtcgt gactgggaaa accctggcgt tacccaactt
     1381 aatcgccttg cagcacatcc ccctttcgcc agctggcgta atagcgaaga ggcccgcacc
     1441 gatcgccctt cccaacagtt gcgcagcctg aatggcgaat gggacgcgcc ctgtagcggc
     1501 gcattaagcg cggcgggtgt ggtggttacg cgcagcgtga ccgctacact tgccagcgcc
     1561 ctagcgcccg ctcctttcgc tttcttccct tcctttctcg ccacgttcgc cggctttccc
     1621 cgtcaagctc taaatcgggg gctcccttta gggttccgat ttagtgcttt acggcacctc
     1681 gaccccaaaa aacttgatta gggtgatggt tcacgtagtg ggccatcgcc ctgatagacg
     1741 gtttttcgcc ctttgacgtt ggagtccacg ttctttaata gtggactctt gttccaaact
     1801 ggaacaacac tcaaccctat ctcggtctat tcttttgatt tataagggat tttgccgatt
     1861 tcggcctatt ggttaaaaaa tgagctgatt taacaaaaat ttaacgcgaa ttttaacaaa
     1921 atattaacgc ttacaattta ggtggcactt ttcggggaaa tgtgcgcgga acccctattt
     1981 gtttattttt ctaaatacat tcaaatatgt atccgctcat gagacaataa ccctgataaa
     2041 tgcttcaata atattgaaaa aggaagagta tgagtattca acatttccgt gtcgccctta
     2101 ttcccttttt tgcggcattt tgccttcctg tttttgctca cccagaaacg ctggtgaaag
     2161 taaaagatgc tgaagatcag ttgggtgcac gagtgggtta catcgaactg gatctcaaca
     2221 gcggtaagat ccttgagagt tttcgccccg aagaacgttt tccaatgatg agcactttta
     2281 aagttctgct atgtggcgcg gtattatccc gtattgacgc cgggcaagag caactcggtc
     2341 gccgcataca ctattctcag aatgacttgg ttgagtactc accagtcaca gaaaagcatc
     2401 ttacggatgg catgacagta agagaattat gcagtgctgc cataaccatg agtgataaca
     2461 ctgcggccaa cttacttctg acaacgatcg gaggaccgaa ggagctaacc gcttttttgc
     2521 acaacatggg ggatcatgta actcgccttg atcgttggga accggagctg aatgaagcca
     2581 taccaaacga cgagcgtgac accacgatgc ctgtagcaat ggcaacaacg ttgcgcaaac
     2641 tattaactgg cgaactactt actctagctt cccggcaaca attaatagac tggatggagg
     2701 cggataaagt tgcaggacca cttctgcgct cggcccttcc ggctggctgg tttattgctg
     2761 ataaatctgg agccggtgag cgtgggtctc gcggtatcat tgcagcactg gggccagatg
     2821 gtaagccctc ccgtatcgta gttatctaca cgacggggag tcaggcaact atggatgaac
     2881 gaaatagaca gatcgctgag ataggtgcct cactgattaa gcattggtaa ctgtcagacc
     2941 aagtttactc atatatactt tagattgatt taaaacttca tttttaattt aaaaggatct
     3001 aggtgaagat cctttttgat aatctcatga ccaaaatccc ttaacgtgag ttttcgttcc
     3061 actgagcgtc agaccccgta gaaaagatca aaggatcttc ttgagatcct ttttttctgc
     3121 gcgtaatctg ctgcttgcaa acaaaaaaac caccgctacc agcggtggtt tgtttgccgg
     3181 atcaagagct accaactctt tttccgaagg taactggctt cagcagagcg cagataccaa
     3241 atactgttct tctagtgtag ccgtagttag gccaccactt caagaactct gtagcaccgc
     3301 ctacatacct cgctctgcta atcctgttac cagtggctgc tgccagtggc gataagtcgt
     3361 gtcttaccgg gttggactca agacgatagt taccggataa ggcgcagcgg tcgggctgaa
     3421 cggggggttc gtgcacacag cccagcttgg agcgaacgac ctacaccgaa ctgagatacc
     3481 tacagcgtga gctatgagaa agcgccacgc ttcccgaagg gagaaaggcg gacaggtatc
     3541 cggtaagcgg cagggtcgga acaggagagc gcacgaggga gcttccaggg ggaaacgcct
     3601 ggtatcttta tagtcctgtc gggtttcgcc acctctgact tgagcgtcga tttttgtgat
     3661 gctcgtcagg ggggcggagc ctatggaaaa acgccagcaa cgcggccttt ttacggttcc
     3721 tggccttttg ctggcctttt gctcacatgt tctttcctgc gttatcccct gattctgtgg
     3781 ataaccgtat taccgccttt gagtgagctg ataccgctcg ccgcagccga acgaccgagc
     3841 gcagcgagtc agtgagcgag gaagcggaag agcgcccaat acgcaaaccg cctctccccg
     3901 cgcgttggcc gattcattaa tgcagctggc acgacaggtt tcccgactgg aaagcgggca
     3961 gtgagcgcaa cgcaattaat gtgagttagc tcactcatta ggcaccccag gctttacact
     4021 ttatgcttcc ggctcgtatg ttgtgtggaa ttgtgagcgg ataacaattt cacacaggaa
     4081 acagctatga ccatgattac gccaagcgcg caattaaccc tcactaaagg gaacaaaagc
     4141 tggagctgca agcttaatgt agtcttatgc aatactcttg tagtcttgca acatggtaac
     4201 gatgagttag caacatgcct tacaaggaga gaaaaagcac cgtgcatgcc gattggtgga
     4261 agtaaggtgg tacgatcgtg ccttattagg aaggcaacag acgggtctga catggattgg
     4321 acgaaccact gaattgccgc attgcagaga tattgtattt aagtgcctag ctcgatacat
     4381 aaacgggtct ctctggttag accagatctg agcctgggag ctctctggct aactagggaa
     4441 cccactgctt aagcctcaat aaagcttgcc ttgagtgctt caagtagtgt gtgcccgtct
     4501 gttgtgtgac tctggtaact agagatccct cagacccttt tagtcagtgt ggaaaatctc
     4561 tagcagtggc gcccgaacag ggacttgaaa gcgaaaggga aaccagagga gctctctcga
     4621 cgcaggactc ggcttgctga agcgcgcacg gcaagaggcg aggggcggcg actggtgagt
     4681 acgccaaaaa ttttgactag cggaggctag aaggagagag atgggtgcga gagcgtcagt
     4741 attaagcggg ggagaattag atcgcgatgg gaaaaaattc ggttaaggcc agggggaaag
     4801 aaaaaatata aattaaaaca tatagtatgg gcaagcaggg agctagaacg attcgcagtt
     4861 aatcctggcc tgttagaaac atcagaaggc tgtagacaaa tactgggaca gctacaacca
     4921 tcccttcaga caggatcaga agaacttaga tcattatata atacagtagc aaccctctat
     4981 tgtgtgcatc aaaggataga gataaaagac accaaggaag ctttagacaa gatagaggaa
     5041 gagcaaaaca aaagtaagac caccgcacag caagcggccg ctgatcttca gacctggagg
     5101 aggagatatg agggacaatt ggagaagtga attatataaa tataaagtag taaaaattga
     5161 accattagga gtagcaccca ccaaggcaaa gagaagagtg gtgcagagag aaaaaagagc
     5221 agtgggaata ggagctttgt tccttgggtt cttgggagca gcaggaagca ctatgggcgc
     5281 agcgtcaatg acgctgacgg tacaggccag acaattattg tctggtatag tgcagcagca
     5341 gaacaatttg ctgagggcta ttgaggcgca acagcatctg ttgcaactca cagtctgggg
     5401 catcaagcag ctccaggcaa gaatcctggc tgtggaaaga tacctaaagg atcaacagct
     5461 cctggggatt tggggttgct ctggaaaact catttgcacc actgctgtgc cttggaatgc
     5521 tagttggagt aataaatctc tggaacagat ttggaatcac acgacctgga tggagtggga
     5581 cagagaaatt aacaattaca caagcttaat acactcctta attgaagaat cgcaaaacca
     5641 gcaagaaaag aatgaacaag aattattgga attagataaa tgggcaagtt tgtggaattg
     5701 gtttaacata acaaattggc tgtggtatat aaaattattc ataatgatag taggaggctt
     5761 ggtaggttta agaatagttt ttgctgtact ttctatagtg aatagagtta ggcagggata
     5821 ttcaccatta tcgtttcaga cccacctccc aaccccgagg ggacccgaca ggcccgaagg
     5881 aatagaagaa gaaggtggag agagagacag agacagatcc attcgattag tgaacggatc
     5941 tcgacggtat cggttaactt ttaaaagaaa aggggggatt ggggggtaca gtgcagggga
     6001 aagaatagta gacataatag caacagacat acaaactaaa gaattacaaa aacaaattac
     6061 aaaaattcaa aattttatcg atgagtaatt catacaaaag gactcgcccc tgccttgggg
     6121 aatcccaggg accgtcgtta aactcccact aacgtagaac ccagagatcg ctgcgttccc
     6181 gccccctcac ccgcccgctc tcgtcatcac tgaggtggag aagagcatgc gtgaggctcc
     6241 ggtgcccgtc agtgggcaga gcgcacatcg cccacagtcc ccgagaagtt ggggggaggg
     6301 gtcggcaatt gaaccggtgc ctagagaagg tggcgcgggg taaactggga aagtgatgtc
     6361 gtgtactggc tccgcctttt tcccgagggt gggggagaac cgtatataag tgcagtagtc
     6421 gccgtgaacg ttctttttcg caacgggttt gccgccagaa cacaggtaag tgccgtgtgt
     6481 ggttcccgcg ggcctggcct ctttacgggt tatggccctt gcgtgccttg aattacttcc
     6541 acgcccctgg ctgcagtacg tgattcttga tcccgagctt cgggttggaa gtgggtggga
     6601 gagttcgagg ccttgcgctt aaggagcccc ttcgcctcgt gcttgagttg aggcctggct
     6661 tgggcgctgg ggccgccgcg tgcgaatctg gtggcacctt cgcgcctgtc tcgctgcttt
     6721 cgataagtct ctagccattt aaaatttttg atgacctgct gcgacgcttt ttttctggca
     6781 agatagtctt gtaaatgcgg gccaagatct gcacactggt atttcggttt ttggggccgc
     6841 gggcggcgac ggggcccgtg cgtcccagcg cacatgttcg gcgaggcggg gcctgcgagc
     6901 gcggccaccg agaatcggac gggggtagtc tcaagctggc cggcctgctc tggtgcctgg
     6961 cctcgcgccg ccgtgtatcg ccccgccctg ggcggcaagg ctggcccggt cggcaccagt
     7021 tgcgtgagcg gaaagatggc cgcttcccgg ccctgctgca gggagctcaa aatggaggac
     7081 gcggcgctcg ggagagcggg cgggtgagtc acccacacaa aggaaaaggg cctttccgtc
     7141 ctcagccgtc gcttcatgtg actccacgga gtaccgggcg ccgtccaggc acctcgatta
     7201 gttctcaagc ttttggagta cgtcgtcttt aggttggggg gaggggtttt atgcgatgga
     7261 gtttccccac actgagtggg tggagactga agttaggcca gcttggcact tgatgtaatt
     7321 ctccttggaa tttgcccttt ttgagtttgg atcttggttc attctcaagc ctcagacagt
     7381 ggttcaaagt ttttttcttc catttcaggt gtcgtgaccc tagcgctacc tctagagcca
     7441 ccatggccgt gatggcccct agaaccctgc tgctgctgct gagcggcgcc ctggccctga
     7501 cccagacatg ggccggcagc cactccatga ggtacttcta tacctctgtg agccggccag
     7561 gaagaggaga gccacgcttt atcgccgtgg gctatgtgga cgatacacag ttcgtgcggt
     7621 ttgacagcga tgccgcctcc cagagaatgg agcctagggc accatggatc gagcaggagg
     7681 gaccagagta ctgggatcag gagacacgga acgtgaaggc ccagtctcag accgacagag
     7741 tggatctggg cacactgagg ggctactata atcagtctga ggacggcagc cacacaatcc
     7801 agatcatgta tggatgcgat gtgggaccag acggcaggtt cctgagggga taccggcagg
     7861 acgcctatga tggcaaggac tacatcgccc tgaacgagga tctgagatcc tggaccgcag
     7921 cagacatggc agcacagatc acaaagagga agtgggaggc agcacacgca gcagagcagc
     7981 agagggcata tctggagggc cggtgcgtgg agtggctgcg gagatacctg gagaatggca
     8041 aggagacact gcagcgcaca gatcccccta agacccacat gacacaccac ccaatcagcg
     8101 accacgaggc caccctgagg tgttgggcac tgggcttcta tcctgccgag atcaccctga
     8161 catggcagag ggatggagag gaccagaccc aggatacaga gctggtggag acacggcccg
     8221 caggcgacgg cacatttcag aagtgggcag cagtggtggt gccatccgga gaggagcaga
     8281 gatacacctg ccacgtgcag cacgagggcc tgccaaagcc cctgaccctg agatgggagc
     8341 tgagctccca gcccacaatc cctatcgtgg gcatcatcgc aggcctggtg ctgctgggag
     8401 ccgtgatcac aggagcagtg gtggcagccg tgatgtggag gcgcaagtct agcgatagaa
     8461 agggaggatc ctacacccag gcagcatcct ctgattctgc ccagggctcc gacgtgtctc
     8521 tgacagcctg taaggtgtga ggcgcgcctc ccagagccac cgttacactc ctgccgccct
     8581 ctgggctgca ccgactcggt gccacttttt caagttgata acggactagc cttattttaa
     8641 cttgctattt ctagctctaa aacttttttt gactagatct tgagacaaat ggcagtattc
     8701 atccacaagg tacgctaaac gcctgcaacc aggacgcgt
//

Lenti plKO_EF1A_ORF_HLA_A*24 02

LOCUS plKO_EF1A_ORF_HLA_A*24 02                8739 bp DNA     circular SYN 24-OCT-2022
DEFINITION  synthetic circular DNA
ACCESSION   .
VERSION     .
KEYWORDS    .
SOURCE      synthetic DNA construct
  ORGANISM  synthetic DNA construct
REFERENCE   1  (bases 1 to 8739)
  AUTHORS   .
  TITLE     Direct Submission
  JOURNAL   Exported Oct 25, 2022 from SnapGene 6.0.5
            https://www.snapgene.com
COMMENT     U515NEI230-128 DNA889099
FEATURES             Location/Qualifiers
     source          1..8739
                     /mol_type="other DNA"
                     /organism="synthetic DNA construct"
     misc_feature    10..598
                     /label=WPRE
     CDS             complement(481..492)
                     /label=Factor Xa site
     LTR             670..903
                     /label=3' LTR (Delta-U3)
     polyA_signal    981..1102
                     /label=SV40 poly(A) signal
     rep_origin      1142..1277
                     /label=SV40 ori
     promoter        complement(1298..1316)
                     /label=T7 promoter
     primer_bind     complement(1326..1342)
                     /label=M13 fwd
     rep_origin      1484..1939
                     /label=f1 ori
     promoter        1965..2069
                     /label=AmpR promoter
     CDS             2070..2138
                     /label=AmpR
     CDS             2139..2930
                     /label=AmpR
     rep_origin      3101..3689
                     /label=ori
     promoter        4019..4036
                     /label=lac promoter
     protein_bind    4051..4067
                     /label=lac operator
     primer_bind     4075..4091
                     /label=M13 rev
     promoter        4112..4130
                     /label=T3 promoter
     promoter        4158..4384
                     /label=RSV promoter
     LTR             4385..4565
                     /label=5' LTR (truncated)
     misc_feature    4612..4737
                     /label=HIV-1 Psi
     misc_feature    5230..5463
                     /label=RRE
     misc_feature    5959..6076
                     /label=cPPT/CTS
     promoter        6236..7417
                     /label=EF-1-alpha promoter
     intron          6466..7408
                     /label=EF-1-alpha intron A
     regulatory      7437..7446
                     /label=Kozak sequence
     CDS             7443..8540
                     /codon_start=1
                     /label=HLA-A*24:02
                     /translation="MAVMAPRTLVLLLSGALALTQTWAGSHSMRYFSTSVSRPGRGEPR
                     FIAVGYVDDTQFVRFDSDAASQRMEPRAPWIEQEGPEYWDEETGKVKAHSQTDRENLRI
                     ALRYYNQSEAGSHTLQMMFGCDVGSDGRFLRGYHQYAYDGKDYIALKEDLRSWTAADMA
                     AQITKRKWEAAHVAEQQRAYLEGTCVDGLRRYLENGKETLQRTDPPKTHMTHHPISDHE
                     ATLRCWALGFYPAEITLTWQRDGEDQTQDTELVETRPAGDGTFQKWAAVVVPSGEEQRY
                     TCHVQHEGLPKPLTLRWEPSSQPTVPIVGIIAGLVLLGAVITGAVVAAVMWRRNSSDRK
                     GGSYSQAASSDSAQGSDVSLTACKV"
     misc_RNA        complement(8588..8663)
                     /label=gRNA scaffold/Barcode
ORIGIN
        1 taagtcgaca atcaacctct ggattacaaa atttgtgaaa gattgactgg tattcttaac
       61 tatgttgctc cttttacgct atgtggatac gctgctttaa tgcctttgta tcatgctatt
      121 gcttcccgta tggctttcat tttctcctcc ttgtataaat cctggttgct gtctctttat
      181 gaggagttgt ggcccgttgt caggcaacgt ggcgtggtgt gcactgtgtt tgctgacgca
      241 acccccactg gttggggcat tgccaccacc tgtcagctcc tttccgggac tttcgctttc
      301 cccctcccta ttgccacggc ggaactcatc gccgcctgcc ttgcccgctg ctggacaggg
      361 gctcggctgt tgggcactga caattccgtg gtgttgtcgg ggaaatcatc gtcctttcct
      421 tggctgctcg cctgtgttgc cacctggatt ctgcgcggga cgtccttctg ctacgtccct
      481 tcggccctca atccagcgga ccttccttcc cgcggcctgc tgccggctct gcggcctctt
      541 ccgcgtcttc gccttcgccc tcagacgagt cggatctccc tttgggccgc ctccccgcgt
      601 cgactttaag accaatgact tacaaggcag ctgtagatct tagccacttt ttaaaagaaa
      661 aggggggact ggaagggcta attcactccc aacgaagaca agatctgctt tttgcttgta
      721 ctgggtctct ctggttagac cagatctgag cctgggagct ctctggctaa ctagggaacc
      781 cactgcttaa gcctcaataa agcttgcctt gagtgcttca agtagtgtgt gcccgtctgt
      841 tgtgtgactc tggtaactag agatccctca gaccctttta gtcagtgtgg aaaatctcta
      901 gcagtacgta tagtagttca tgtcatctta ttattcagta tttataactt gcaaagaaat
      961 gaatatcaga gagtgagagg aacttgttta ttgcagctta taatggttac aaataaagca
     1021 atagcatcac aaatttcaca aataaagcat ttttttcact gcattctagt tgtggtttgt
     1081 ccaaactcat caatgtatct tatcatgtct ggctctagct atcccgcccc taactccgcc
     1141 catcccgccc ctaactccgc ccagttccgc ccattctccg ccccatggct gactaatttt
     1201 ttttatttat gcagaggccg aggccgcctc ggcctctgag ctattccaga agtagtgagg
     1261 aggctttttt ggaggcctag ggacgtaccc aattcgccct atagtgagtc gtattacgcg
     1321 cgctcactgg ccgtcgtttt acaacgtcgt gactgggaaa accctggcgt tacccaactt
     1381 aatcgccttg cagcacatcc ccctttcgcc agctggcgta atagcgaaga ggcccgcacc
     1441 gatcgccctt cccaacagtt gcgcagcctg aatggcgaat gggacgcgcc ctgtagcggc
     1501 gcattaagcg cggcgggtgt ggtggttacg cgcagcgtga ccgctacact tgccagcgcc
     1561 ctagcgcccg ctcctttcgc tttcttccct tcctttctcg ccacgttcgc cggctttccc
     1621 cgtcaagctc taaatcgggg gctcccttta gggttccgat ttagtgcttt acggcacctc
     1681 gaccccaaaa aacttgatta gggtgatggt tcacgtagtg ggccatcgcc ctgatagacg
     1741 gtttttcgcc ctttgacgtt ggagtccacg ttctttaata gtggactctt gttccaaact
     1801 ggaacaacac tcaaccctat ctcggtctat tcttttgatt tataagggat tttgccgatt
     1861 tcggcctatt ggttaaaaaa tgagctgatt taacaaaaat ttaacgcgaa ttttaacaaa
     1921 atattaacgc ttacaattta ggtggcactt ttcggggaaa tgtgcgcgga acccctattt
     1981 gtttattttt ctaaatacat tcaaatatgt atccgctcat gagacaataa ccctgataaa
     2041 tgcttcaata atattgaaaa aggaagagta tgagtattca acatttccgt gtcgccctta
     2101 ttcccttttt tgcggcattt tgccttcctg tttttgctca cccagaaacg ctggtgaaag
     2161 taaaagatgc tgaagatcag ttgggtgcac gagtgggtta catcgaactg gatctcaaca
     2221 gcggtaagat ccttgagagt tttcgccccg aagaacgttt tccaatgatg agcactttta
     2281 aagttctgct atgtggcgcg gtattatccc gtattgacgc cgggcaagag caactcggtc
     2341 gccgcataca ctattctcag aatgacttgg ttgagtactc accagtcaca gaaaagcatc
     2401 ttacggatgg catgacagta agagaattat gcagtgctgc cataaccatg agtgataaca
     2461 ctgcggccaa cttacttctg acaacgatcg gaggaccgaa ggagctaacc gcttttttgc
     2521 acaacatggg ggatcatgta actcgccttg atcgttggga accggagctg aatgaagcca
     2581 taccaaacga cgagcgtgac accacgatgc ctgtagcaat ggcaacaacg ttgcgcaaac
     2641 tattaactgg cgaactactt actctagctt cccggcaaca attaatagac tggatggagg
     2701 cggataaagt tgcaggacca cttctgcgct cggcccttcc ggctggctgg tttattgctg
     2761 ataaatctgg agccggtgag cgtgggtctc gcggtatcat tgcagcactg gggccagatg
     2821 gtaagccctc ccgtatcgta gttatctaca cgacggggag tcaggcaact atggatgaac
     2881 gaaatagaca gatcgctgag ataggtgcct cactgattaa gcattggtaa ctgtcagacc
     2941 aagtttactc atatatactt tagattgatt taaaacttca tttttaattt aaaaggatct
     3001 aggtgaagat cctttttgat aatctcatga ccaaaatccc ttaacgtgag ttttcgttcc
     3061 actgagcgtc agaccccgta gaaaagatca aaggatcttc ttgagatcct ttttttctgc
     3121 gcgtaatctg ctgcttgcaa acaaaaaaac caccgctacc agcggtggtt tgtttgccgg
     3181 atcaagagct accaactctt tttccgaagg taactggctt cagcagagcg cagataccaa
     3241 atactgttct tctagtgtag ccgtagttag gccaccactt caagaactct gtagcaccgc
     3301 ctacatacct cgctctgcta atcctgttac cagtggctgc tgccagtggc gataagtcgt
     3361 gtcttaccgg gttggactca agacgatagt taccggataa ggcgcagcgg tcgggctgaa
     3421 cggggggttc gtgcacacag cccagcttgg agcgaacgac ctacaccgaa ctgagatacc
     3481 tacagcgtga gctatgagaa agcgccacgc ttcccgaagg gagaaaggcg gacaggtatc
     3541 cggtaagcgg cagggtcgga acaggagagc gcacgaggga gcttccaggg ggaaacgcct
     3601 ggtatcttta tagtcctgtc gggtttcgcc acctctgact tgagcgtcga tttttgtgat
     3661 gctcgtcagg ggggcggagc ctatggaaaa acgccagcaa cgcggccttt ttacggttcc
     3721 tggccttttg ctggcctttt gctcacatgt tctttcctgc gttatcccct gattctgtgg
     3781 ataaccgtat taccgccttt gagtgagctg ataccgctcg ccgcagccga acgaccgagc
     3841 gcagcgagtc agtgagcgag gaagcggaag agcgcccaat acgcaaaccg cctctccccg
     3901 cgcgttggcc gattcattaa tgcagctggc acgacaggtt tcccgactgg aaagcgggca
     3961 gtgagcgcaa cgcaattaat gtgagttagc tcactcatta ggcaccccag gctttacact
     4021 ttatgcttcc ggctcgtatg ttgtgtggaa ttgtgagcgg ataacaattt cacacaggaa
     4081 acagctatga ccatgattac gccaagcgcg caattaaccc tcactaaagg gaacaaaagc
     4141 tggagctgca agcttaatgt agtcttatgc aatactcttg tagtcttgca acatggtaac
     4201 gatgagttag caacatgcct tacaaggaga gaaaaagcac cgtgcatgcc gattggtgga
     4261 agtaaggtgg tacgatcgtg ccttattagg aaggcaacag acgggtctga catggattgg
     4321 acgaaccact gaattgccgc attgcagaga tattgtattt aagtgcctag ctcgatacat
     4381 aaacgggtct ctctggttag accagatctg agcctgggag ctctctggct aactagggaa
     4441 cccactgctt aagcctcaat aaagcttgcc ttgagtgctt caagtagtgt gtgcccgtct
     4501 gttgtgtgac tctggtaact agagatccct cagacccttt tagtcagtgt ggaaaatctc
     4561 tagcagtggc gcccgaacag ggacttgaaa gcgaaaggga aaccagagga gctctctcga
     4621 cgcaggactc ggcttgctga agcgcgcacg gcaagaggcg aggggcggcg actggtgagt
     4681 acgccaaaaa ttttgactag cggaggctag aaggagagag atgggtgcga gagcgtcagt
     4741 attaagcggg ggagaattag atcgcgatgg gaaaaaattc ggttaaggcc agggggaaag
     4801 aaaaaatata aattaaaaca tatagtatgg gcaagcaggg agctagaacg attcgcagtt
     4861 aatcctggcc tgttagaaac atcagaaggc tgtagacaaa tactgggaca gctacaacca
     4921 tcccttcaga caggatcaga agaacttaga tcattatata atacagtagc aaccctctat
     4981 tgtgtgcatc aaaggataga gataaaagac accaaggaag ctttagacaa gatagaggaa
     5041 gagcaaaaca aaagtaagac caccgcacag caagcggccg ctgatcttca gacctggagg
     5101 aggagatatg agggacaatt ggagaagtga attatataaa tataaagtag taaaaattga
     5161 accattagga gtagcaccca ccaaggcaaa gagaagagtg gtgcagagag aaaaaagagc
     5221 agtgggaata ggagctttgt tccttgggtt cttgggagca gcaggaagca ctatgggcgc
     5281 agcgtcaatg acgctgacgg tacaggccag acaattattg tctggtatag tgcagcagca
     5341 gaacaatttg ctgagggcta ttgaggcgca acagcatctg ttgcaactca cagtctgggg
     5401 catcaagcag ctccaggcaa gaatcctggc tgtggaaaga tacctaaagg atcaacagct
     5461 cctggggatt tggggttgct ctggaaaact catttgcacc actgctgtgc cttggaatgc
     5521 tagttggagt aataaatctc tggaacagat ttggaatcac acgacctgga tggagtggga
     5581 cagagaaatt aacaattaca caagcttaat acactcctta attgaagaat cgcaaaacca
     5641 gcaagaaaag aatgaacaag aattattgga attagataaa tgggcaagtt tgtggaattg
     5701 gtttaacata acaaattggc tgtggtatat aaaattattc ataatgatag taggaggctt
     5761 ggtaggttta agaatagttt ttgctgtact ttctatagtg aatagagtta ggcagggata
     5821 ttcaccatta tcgtttcaga cccacctccc aaccccgagg ggacccgaca ggcccgaagg
     5881 aatagaagaa gaaggtggag agagagacag agacagatcc attcgattag tgaacggatc
     5941 tcgacggtat cggttaactt ttaaaagaaa aggggggatt ggggggtaca gtgcagggga
     6001 aagaatagta gacataatag caacagacat acaaactaaa gaattacaaa aacaaattac
     6061 aaaaattcaa aattttatcg atgagtaatt catacaaaag gactcgcccc tgccttgggg
     6121 aatcccaggg accgtcgtta aactcccact aacgtagaac ccagagatcg ctgcgttccc
     6181 gccccctcac ccgcccgctc tcgtcatcac tgaggtggag aagagcatgc gtgaggctcc
     6241 ggtgcccgtc agtgggcaga gcgcacatcg cccacagtcc ccgagaagtt ggggggaggg
     6301 gtcggcaatt gaaccggtgc ctagagaagg tggcgcgggg taaactggga aagtgatgtc
     6361 gtgtactggc tccgcctttt tcccgagggt gggggagaac cgtatataag tgcagtagtc
     6421 gccgtgaacg ttctttttcg caacgggttt gccgccagaa cacaggtaag tgccgtgtgt
     6481 ggttcccgcg ggcctggcct ctttacgggt tatggccctt gcgtgccttg aattacttcc
     6541 acgcccctgg ctgcagtacg tgattcttga tcccgagctt cgggttggaa gtgggtggga
     6601 gagttcgagg ccttgcgctt aaggagcccc ttcgcctcgt gcttgagttg aggcctggct
     6661 tgggcgctgg ggccgccgcg tgcgaatctg gtggcacctt cgcgcctgtc tcgctgcttt
     6721 cgataagtct ctagccattt aaaatttttg atgacctgct gcgacgcttt ttttctggca
     6781 agatagtctt gtaaatgcgg gccaagatct gcacactggt atttcggttt ttggggccgc
     6841 gggcggcgac ggggcccgtg cgtcccagcg cacatgttcg gcgaggcggg gcctgcgagc
     6901 gcggccaccg agaatcggac gggggtagtc tcaagctggc cggcctgctc tggtgcctgg
     6961 cctcgcgccg ccgtgtatcg ccccgccctg ggcggcaagg ctggcccggt cggcaccagt
     7021 tgcgtgagcg gaaagatggc cgcttcccgg ccctgctgca gggagctcaa aatggaggac
     7081 gcggcgctcg ggagagcggg cgggtgagtc acccacacaa aggaaaaggg cctttccgtc
     7141 ctcagccgtc gcttcatgtg actccacgga gtaccgggcg ccgtccaggc acctcgatta
     7201 gttctcaagc ttttggagta cgtcgtcttt aggttggggg gaggggtttt atgcgatgga
     7261 gtttccccac actgagtggg tggagactga agttaggcca gcttggcact tgatgtaatt
     7321 ctccttggaa tttgcccttt ttgagtttgg atcttggttc attctcaagc ctcagacagt
     7381 ggttcaaagt ttttttcttc catttcaggt gtcgtgaccc tagcgctacc tctagagcca
     7441 ccatggccgt gatggcacca cgcaccctgg tgctgctgct gtctggcgcc ctggccctga
     7501 cccagacatg ggcaggcagc cactccatgc ggtacttctc taccagcgtg tcccggccag
     7561 gaagaggaga gccacgcttt atcgccgtgg gctatgtgga cgatacacag ttcgtgcggt
     7621 ttgacagcga tgcagcatcc cagaggatgg agcctagagc accatggatc gagcaggagg
     7681 gaccagagta ctgggacgag gagacaggca aggtgaaggc ccacagccag acagataggg
     7741 agaacctgcg catcgccctg cggtactata atcagtctga ggccggcagc cacaccctgc
     7801 agatgatgtt cggctgcgac gtgggctccg atggcaggtt tctgcgcggc taccaccagt
     7861 acgcctatga cggcaaggat tatatcgccc tgaaggagga cctgagatct tggaccgccg
     7921 ccgatatggc cgcccagatc acaaagagaa agtgggaggc agcacacgtg gcagagcagc
     7981 agagggcata cctggaggga acctgcgtgg acggcctgcg gagatatctg gagaacggca
     8041 aggagacact gcagagaaca gaccccccta agacccacat gacacaccac cccatcagcg
     8101 atcacgaggc cacactgagg tgttgggccc tgggcttcta ccctgccgag atcaccctga
     8161 catggcagag agacggcgag gatcagaccc aggacacaga gctggtggag acacggcccg
     8221 caggcgatgg cacatttcag aagtgggcag cagtggtggt gccatccgga gaggagcagc
     8281 ggtacacctg ccacgtgcag cacgagggcc tgccaaagcc actgaccctg agatgggagc
     8341 ctagctccca gcccacagtg cctatcgtgg gcatcatcgc cggcctggtg ctgctgggag
     8401 ccgtgatcac cggagcagtg gtggccgccg tgatgtggag gcgcaattct agcgatagga
     8461 agggcggctc ttatagccag gcagcatcct ctgactccgc ccagggatcc gacgtgagcc
     8521 tgacagcctg taaggtgtga ggcgcgcctc ccagagccac cgttacaccc tgccgccctc
     8581 tgggctcgca ccgactcggt gccacttttt caagttgata acggactagc cttattttaa
     8641 cttgctattt ctagctctaa aacttttttt gactagatct tgagacaaat ggcagtattc
     8701 atccacaagg tacgctaaac gcctgcaacc aggacgcgt
//

Lenti plKO_EF1A_ORF_HLA_B*07 02

LOCUS plKO_EF1A_ORF_HLA_B*07 02                8730 bp DNA     circular SYN 24-OCT-2022
DEFINITION  synthetic circular DNA
ACCESSION   .
VERSION     .
KEYWORDS    .
SOURCE      synthetic DNA construct
  ORGANISM  synthetic DNA construct
REFERENCE   1  (bases 1 to 8730)
  AUTHORS   .
  TITLE     Direct Submission
  JOURNAL   Exported Oct 25, 2022 from SnapGene 6.0.5
            https://www.snapgene.com
COMMENT     U515NEI230-150 DNA889100
FEATURES             Location/Qualifiers
     source          1..8730
                     /mol_type="other DNA"
                     /organism="synthetic DNA construct"
     misc_feature    10..598
                     /label=WPRE
     CDS             complement(481..492)
                     /label=Factor Xa site
     LTR             670..903
                     /label=3' LTR (Delta-U3)
     polyA_signal    981..1102
                     /label=SV40 poly(A) signal
     rep_origin      1142..1277
                     /label=SV40 ori
     promoter        complement(1298..1316)
                     /label=T7 promoter
     primer_bind     complement(1326..1342)
                     /label=M13 fwd
     rep_origin      1484..1939
                     /label=f1 ori
     promoter        1965..2069
                     /label=AmpR promoter
     CDS             2070..2138
                     /label=AmpR
     CDS             2139..2930
                     /label=AmpR
     rep_origin      3101..3689
                     /label=ori
     promoter        4019..4036
                     /label=lac promoter
     protein_bind    4051..4067
                     /label=lac operator
     primer_bind     4075..4091
                     /label=M13 rev
     promoter        4112..4130
                     /label=T3 promoter
     promoter        4158..4384
                     /label=RSV promoter
     LTR             4385..4565
                     /label=5' LTR (truncated)
     misc_feature    4612..4737
                     /label=HIV-1 Psi
     misc_feature    5230..5463
                     /label=RRE
     misc_feature    5959..6076
                     /label=cPPT/CTS
     promoter        6236..7417
                     /label=EF-1-alpha promoter
     intron          6466..7408
                     /label=EF-1-alpha intron A
     CDS             7443..8531
                     /codon_start=1
                     /label=HLA*B:07:02
                     /translation="MLVMAPRTVLLLLSAALALTETWAGSHSMRYFYTSVSRPGRGEPR
                     FISVGYVDDTQFVRFDSDAASPREEPRAPWIEQEGPEYWDRNTQIYKAQAQTDRESLRN
                     LRGYYNQSEAGSHTLQSMYGCDVGPDGRLLRGHDQYAYDGKDYIALNEDLRSWTAADTA
                     AQITQRKWEAAREAEQRRAYLEGECVEWLRRYLENGKDKLERADPPKTHVTHHPISDHE
                     ATLRCWALGFYPAEITLTWQRDGEDQTQDTELVETRPAGDRTFQKWAAVVVPSGEEQRY
                     TCHVQHEGLPKPLTLRWEPSSQSTVPIVGIVAGLAVLAVVVIGAVVAAVMCRRKSSGGK
                     GGSYSQAACSDSAQGSDVSLTA"
     misc_RNA        complement(8579..8654)
                     /label=gRNA scaffold/Barcode
ORIGIN
        1 taagtcgaca atcaacctct ggattacaaa atttgtgaaa gattgactgg tattcttaac
       61 tatgttgctc cttttacgct atgtggatac gctgctttaa tgcctttgta tcatgctatt
      121 gcttcccgta tggctttcat tttctcctcc ttgtataaat cctggttgct gtctctttat
      181 gaggagttgt ggcccgttgt caggcaacgt ggcgtggtgt gcactgtgtt tgctgacgca
      241 acccccactg gttggggcat tgccaccacc tgtcagctcc tttccgggac tttcgctttc
      301 cccctcccta ttgccacggc ggaactcatc gccgcctgcc ttgcccgctg ctggacaggg
      361 gctcggctgt tgggcactga caattccgtg gtgttgtcgg ggaaatcatc gtcctttcct
      421 tggctgctcg cctgtgttgc cacctggatt ctgcgcggga cgtccttctg ctacgtccct
      481 tcggccctca atccagcgga ccttccttcc cgcggcctgc tgccggctct gcggcctctt
      541 ccgcgtcttc gccttcgccc tcagacgagt cggatctccc tttgggccgc ctccccgcgt
      601 cgactttaag accaatgact tacaaggcag ctgtagatct tagccacttt ttaaaagaaa
      661 aggggggact ggaagggcta attcactccc aacgaagaca agatctgctt tttgcttgta
      721 ctgggtctct ctggttagac cagatctgag cctgggagct ctctggctaa ctagggaacc
      781 cactgcttaa gcctcaataa agcttgcctt gagtgcttca agtagtgtgt gcccgtctgt
      841 tgtgtgactc tggtaactag agatccctca gaccctttta gtcagtgtgg aaaatctcta
      901 gcagtacgta tagtagttca tgtcatctta ttattcagta tttataactt gcaaagaaat
      961 gaatatcaga gagtgagagg aacttgttta ttgcagctta taatggttac aaataaagca
     1021 atagcatcac aaatttcaca aataaagcat ttttttcact gcattctagt tgtggtttgt
     1081 ccaaactcat caatgtatct tatcatgtct ggctctagct atcccgcccc taactccgcc
     1141 catcccgccc ctaactccgc ccagttccgc ccattctccg ccccatggct gactaatttt
     1201 ttttatttat gcagaggccg aggccgcctc ggcctctgag ctattccaga agtagtgagg
     1261 aggctttttt ggaggcctag ggacgtaccc aattcgccct atagtgagtc gtattacgcg
     1321 cgctcactgg ccgtcgtttt acaacgtcgt gactgggaaa accctggcgt tacccaactt
     1381 aatcgccttg cagcacatcc ccctttcgcc agctggcgta atagcgaaga ggcccgcacc
     1441 gatcgccctt cccaacagtt gcgcagcctg aatggcgaat gggacgcgcc ctgtagcggc
     1501 gcattaagcg cggcgggtgt ggtggttacg cgcagcgtga ccgctacact tgccagcgcc
     1561 ctagcgcccg ctcctttcgc tttcttccct tcctttctcg ccacgttcgc cggctttccc
     1621 cgtcaagctc taaatcgggg gctcccttta gggttccgat ttagtgcttt acggcacctc
     1681 gaccccaaaa aacttgatta gggtgatggt tcacgtagtg ggccatcgcc ctgatagacg
     1741 gtttttcgcc ctttgacgtt ggagtccacg ttctttaata gtggactctt gttccaaact
     1801 ggaacaacac tcaaccctat ctcggtctat tcttttgatt tataagggat tttgccgatt
     1861 tcggcctatt ggttaaaaaa tgagctgatt taacaaaaat ttaacgcgaa ttttaacaaa
     1921 atattaacgc ttacaattta ggtggcactt ttcggggaaa tgtgcgcgga acccctattt
     1981 gtttattttt ctaaatacat tcaaatatgt atccgctcat gagacaataa ccctgataaa
     2041 tgcttcaata atattgaaaa aggaagagta tgagtattca acatttccgt gtcgccctta
     2101 ttcccttttt tgcggcattt tgccttcctg tttttgctca cccagaaacg ctggtgaaag
     2161 taaaagatgc tgaagatcag ttgggtgcac gagtgggtta catcgaactg gatctcaaca
     2221 gcggtaagat ccttgagagt tttcgccccg aagaacgttt tccaatgatg agcactttta
     2281 aagttctgct atgtggcgcg gtattatccc gtattgacgc cgggcaagag caactcggtc
     2341 gccgcataca ctattctcag aatgacttgg ttgagtactc accagtcaca gaaaagcatc
     2401 ttacggatgg catgacagta agagaattat gcagtgctgc cataaccatg agtgataaca
     2461 ctgcggccaa cttacttctg acaacgatcg gaggaccgaa ggagctaacc gcttttttgc
     2521 acaacatggg ggatcatgta actcgccttg atcgttggga accggagctg aatgaagcca
     2581 taccaaacga cgagcgtgac accacgatgc ctgtagcaat ggcaacaacg ttgcgcaaac
     2641 tattaactgg cgaactactt actctagctt cccggcaaca attaatagac tggatggagg
     2701 cggataaagt tgcaggacca cttctgcgct cggcccttcc ggctggctgg tttattgctg
     2761 ataaatctgg agccggtgag cgtgggtctc gcggtatcat tgcagcactg gggccagatg
     2821 gtaagccctc ccgtatcgta gttatctaca cgacggggag tcaggcaact atggatgaac
     2881 gaaatagaca gatcgctgag ataggtgcct cactgattaa gcattggtaa ctgtcagacc
     2941 aagtttactc atatatactt tagattgatt taaaacttca tttttaattt aaaaggatct
     3001 aggtgaagat cctttttgat aatctcatga ccaaaatccc ttaacgtgag ttttcgttcc
     3061 actgagcgtc agaccccgta gaaaagatca aaggatcttc ttgagatcct ttttttctgc
     3121 gcgtaatctg ctgcttgcaa acaaaaaaac caccgctacc agcggtggtt tgtttgccgg
     3181 atcaagagct accaactctt tttccgaagg taactggctt cagcagagcg cagataccaa
     3241 atactgttct tctagtgtag ccgtagttag gccaccactt caagaactct gtagcaccgc
     3301 ctacatacct cgctctgcta atcctgttac cagtggctgc tgccagtggc gataagtcgt
     3361 gtcttaccgg gttggactca agacgatagt taccggataa ggcgcagcgg tcgggctgaa
     3421 cggggggttc gtgcacacag cccagcttgg agcgaacgac ctacaccgaa ctgagatacc
     3481 tacagcgtga gctatgagaa agcgccacgc ttcccgaagg gagaaaggcg gacaggtatc
     3541 cggtaagcgg cagggtcgga acaggagagc gcacgaggga gcttccaggg ggaaacgcct
     3601 ggtatcttta tagtcctgtc gggtttcgcc acctctgact tgagcgtcga tttttgtgat
     3661 gctcgtcagg ggggcggagc ctatggaaaa acgccagcaa cgcggccttt ttacggttcc
     3721 tggccttttg ctggcctttt gctcacatgt tctttcctgc gttatcccct gattctgtgg
     3781 ataaccgtat taccgccttt gagtgagctg ataccgctcg ccgcagccga acgaccgagc
     3841 gcagcgagtc agtgagcgag gaagcggaag agcgcccaat acgcaaaccg cctctccccg
     3901 cgcgttggcc gattcattaa tgcagctggc acgacaggtt tcccgactgg aaagcgggca
     3961 gtgagcgcaa cgcaattaat gtgagttagc tcactcatta ggcaccccag gctttacact
     4021 ttatgcttcc ggctcgtatg ttgtgtggaa ttgtgagcgg ataacaattt cacacaggaa
     4081 acagctatga ccatgattac gccaagcgcg caattaaccc tcactaaagg gaacaaaagc
     4141 tggagctgca agcttaatgt agtcttatgc aatactcttg tagtcttgca acatggtaac
     4201 gatgagttag caacatgcct tacaaggaga gaaaaagcac cgtgcatgcc gattggtgga
     4261 agtaaggtgg tacgatcgtg ccttattagg aaggcaacag acgggtctga catggattgg
     4321 acgaaccact gaattgccgc attgcagaga tattgtattt aagtgcctag ctcgatacat
     4381 aaacgggtct ctctggttag accagatctg agcctgggag ctctctggct aactagggaa
     4441 cccactgctt aagcctcaat aaagcttgcc ttgagtgctt caagtagtgt gtgcccgtct
     4501 gttgtgtgac tctggtaact agagatccct cagacccttt tagtcagtgt ggaaaatctc
     4561 tagcagtggc gcccgaacag ggacttgaaa gcgaaaggga aaccagagga gctctctcga
     4621 cgcaggactc ggcttgctga agcgcgcacg gcaagaggcg aggggcggcg actggtgagt
     4681 acgccaaaaa ttttgactag cggaggctag aaggagagag atgggtgcga gagcgtcagt
     4741 attaagcggg ggagaattag atcgcgatgg gaaaaaattc ggttaaggcc agggggaaag
     4801 aaaaaatata aattaaaaca tatagtatgg gcaagcaggg agctagaacg attcgcagtt
     4861 aatcctggcc tgttagaaac atcagaaggc tgtagacaaa tactgggaca gctacaacca
     4921 tcccttcaga caggatcaga agaacttaga tcattatata atacagtagc aaccctctat
     4981 tgtgtgcatc aaaggataga gataaaagac accaaggaag ctttagacaa gatagaggaa
     5041 gagcaaaaca aaagtaagac caccgcacag caagcggccg ctgatcttca gacctggagg
     5101 aggagatatg agggacaatt ggagaagtga attatataaa tataaagtag taaaaattga
     5161 accattagga gtagcaccca ccaaggcaaa gagaagagtg gtgcagagag aaaaaagagc
     5221 agtgggaata ggagctttgt tccttgggtt cttgggagca gcaggaagca ctatgggcgc
     5281 agcgtcaatg acgctgacgg tacaggccag acaattattg tctggtatag tgcagcagca
     5341 gaacaatttg ctgagggcta ttgaggcgca acagcatctg ttgcaactca cagtctgggg
     5401 catcaagcag ctccaggcaa gaatcctggc tgtggaaaga tacctaaagg atcaacagct
     5461 cctggggatt tggggttgct ctggaaaact catttgcacc actgctgtgc cttggaatgc
     5521 tagttggagt aataaatctc tggaacagat ttggaatcac acgacctgga tggagtggga
     5581 cagagaaatt aacaattaca caagcttaat acactcctta attgaagaat cgcaaaacca
     5641 gcaagaaaag aatgaacaag aattattgga attagataaa tgggcaagtt tgtggaattg
     5701 gtttaacata acaaattggc tgtggtatat aaaattattc ataatgatag taggaggctt
     5761 ggtaggttta agaatagttt ttgctgtact ttctatagtg aatagagtta ggcagggata
     5821 ttcaccatta tcgtttcaga cccacctccc aaccccgagg ggacccgaca ggcccgaagg
     5881 aatagaagaa gaaggtggag agagagacag agacagatcc attcgattag tgaacggatc
     5941 tcgacggtat cggttaactt ttaaaagaaa aggggggatt ggggggtaca gtgcagggga
     6001 aagaatagta gacataatag caacagacat acaaactaaa gaattacaaa aacaaattac
     6061 aaaaattcaa aattttatcg atgagtaatt catacaaaag gactcgcccc tgccttgggg
     6121 aatcccaggg accgtcgtta aactcccact aacgtagaac ccagagatcg ctgcgttccc
     6181 gccccctcac ccgcccgctc tcgtcatcac tgaggtggag aagagcatgc gtgaggctcc
     6241 ggtgcccgtc agtgggcaga gcgcacatcg cccacagtcc ccgagaagtt ggggggaggg
     6301 gtcggcaatt gaaccggtgc ctagagaagg tggcgcgggg taaactggga aagtgatgtc
     6361 gtgtactggc tccgcctttt tcccgagggt gggggagaac cgtatataag tgcagtagtc
     6421 gccgtgaacg ttctttttcg caacgggttt gccgccagaa cacaggtaag tgccgtgtgt
     6481 ggttcccgcg ggcctggcct ctttacgggt tatggccctt gcgtgccttg aattacttcc
     6541 acgcccctgg ctgcagtacg tgattcttga tcccgagctt cgggttggaa gtgggtggga
     6601 gagttcgagg ccttgcgctt aaggagcccc ttcgcctcgt gcttgagttg aggcctggct
     6661 tgggcgctgg ggccgccgcg tgcgaatctg gtggcacctt cgcgcctgtc tcgctgcttt
     6721 cgataagtct ctagccattt aaaatttttg atgacctgct gcgacgcttt ttttctggca
     6781 agatagtctt gtaaatgcgg gccaagatct gcacactggt atttcggttt ttggggccgc
     6841 gggcggcgac ggggcccgtg cgtcccagcg cacatgttcg gcgaggcggg gcctgcgagc
     6901 gcggccaccg agaatcggac gggggtagtc tcaagctggc cggcctgctc tggtgcctgg
     6961 cctcgcgccg ccgtgtatcg ccccgccctg ggcggcaagg ctggcccggt cggcaccagt
     7021 tgcgtgagcg gaaagatggc cgcttcccgg ccctgctgca gggagctcaa aatggaggac
     7081 gcggcgctcg ggagagcggg cgggtgagtc acccacacaa aggaaaaggg cctttccgtc
     7141 ctcagccgtc gcttcatgtg actccacgga gtaccgggcg ccgtccaggc acctcgatta
     7201 gttctcaagc ttttggagta cgtcgtcttt aggttggggg gaggggtttt atgcgatgga
     7261 gtttccccac actgagtggg tggagactga agttaggcca gcttggcact tgatgtaatt
     7321 ctccttggaa tttgcccttt ttgagtttgg atcttggttc attctcaagc ctcagacagt
     7381 ggttcaaagt ttttttcttc catttcaggt gtcgtgaccc tagcgctacc tctagagcca
     7441 ccatgctggt catggcccct aggaccgtgc tgctgctgct gtccgccgcc ctggccctga
     7501 ccgagacatg ggccggctcc cactctatgc gctacttcta taccagcgtg tccaggcccg
     7561 gaaggggaga gcctcggttt atctctgtgg gctacgtgga cgatacacag ttcgtgagat
     7621 ttgactctga tgcagcaagc cctagggagg agcctagagc accatggatc gagcaggagg
     7681 gcccagagta ctgggacagg aacacccaga tctataaggc ccaggcccag acagatcgcg
     7741 agagcctgcg gaacctgaga ggctactata atcagtccga ggccggctct cacaccctgc
     7801 agtccatgta cggctgcgac gtgggaccag atggcaggct gctgagggga cacgaccagt
     7861 acgcctatga cggcaaggat tatatcgccc tgaacgagga cctgcggagc tggaccgcag
     7921 cagatacagc agcccagatc acacagagga agtgggaggc agcaagagag gcagagcaga
     7981 ggagagcata cctggaggga gagtgcgtgg agtggctgag gcgctatctg gagaatggca
     8041 aggacaagct ggagagggcc gatcccccta agacccacgt gacacaccac ccaatctccg
     8101 atcacgaggc caccctgagg tgttgggcac tgggcttcta cccagcagag atcaccctga
     8161 catggcagag ggacggagag gatcagaccc aggacacaga gctggtggag acacggcccg
     8221 caggcgatcg cacatttcag aagtgggcag cagtggtggt gccttctgga gaggagcagc
     8281 ggtatacatg tcacgtgcag cacgagggcc tgccaaagcc cctgaccctg agatgggagc
     8341 caagctccca gagcacagtg ccaatcgtgg gaatcgtggc aggcctggcc gtgctggccg
     8401 tggtggtcat cggagcagtg gtggcagccg tgatgtgccg gagaaagtct agcggaggca
     8461 agggaggatc ttacagccag gcagcatgtt ccgactctgc ccagggaagc gacgtgagcc
     8521 tgaccgcatg aggcgcgcct cccagagcca ccgttacacg gcacccgagt gtgcgctcgc
     8581 accgactcgg tgccactttt tcaagttgat aacggactag ccttatttta acttgctatt
     8641 tctagctcta aaactttttt tgactagatc ttgagacaaa tggcagtatt catccacaag
     8701 gtacgctaaa cgcctgcaac caggacgcgt
//

Lenti plKO_EF1A_ORF_HLA_B*08 01

LOCUS plKO_EF1A_ORF_HLA_B*08 01       Exported                8730 bp DNA     circular SYN 24-OCT-2022
DEFINITION  synthetic circular DNA
ACCESSION   .
VERSION     .
KEYWORDS    .
SOURCE      synthetic DNA construct
  ORGANISM  synthetic DNA construct
REFERENCE   1  (bases 1 to 8730)
  AUTHORS   .
  TITLE     Direct Submission
  JOURNAL   Exported Oct 25, 2022 from SnapGene 6.0.5
            https://www.snapgene.com
COMMENT     U515NEI230-152 DNA889101
FEATURES             Location/Qualifiers
     source          1..8730
                     /mol_type="other DNA"
                     /organism="synthetic DNA construct"
     misc_feature    10..598
                     /label=WPRE
     CDS             complement(481..492)
                     /label=Factor Xa site
     LTR             670..903
                     /label=3' LTR (Delta-U3)
     polyA_signal    981..1102
                     /label=SV40 poly(A) signal
     rep_origin      1142..1277
                     /label=SV40 ori
     promoter        complement(1298..1316)
                     /label=T7 promoter
     primer_bind     complement(1326..1342)
                     /label=M13 fwd
     rep_origin      1484..1939
                     /label=f1 ori
     promoter        1965..2069
                     /label=AmpR promoter
     CDS             2070..2138
                     /label=AmpR
     CDS             2139..2930
                     /label=AmpR
     rep_origin      3101..3689
                     /label=ori
     promoter        4019..4036
                     /label=lac promoter
     protein_bind    4051..4067
                     /label=lac operator
     primer_bind     4075..4091
                     /label=M13 rev
     promoter        4112..4130
                     /label=T3 promoter
     promoter        4158..4384
                     /label=RSV promoter
     LTR             4385..4565
                     /label=5' LTR (truncated)
     misc_feature    4612..4737
                     /label=HIV-1 Psi
     misc_feature    5230..5463
                     /label=RRE
     misc_feature    5959..6076
                     /label=cPPT/CTS
     promoter        6236..7417
                     /label=EF-1-alpha promoter
     intron          6466..7408
                     /label=EF-1-alpha intron A
     CDS             7443..8531
                     /codon_start=1
                     /label=HLA-B*08:01
                     /translation="MLVMAPRTVLLLLSAALALTETWAGSHSMRYFDTAMSRPGRGEPR
                     FISVGYVDDTQFVRFDSDAASPREEPRAPWIEQEGPEYWDRNTQIFKTNTQTDRESLRN
                     LRGYYNQSEAGSHTLQSMYGCDVGPDGRLLRGHNQYAYDGKDYIALNEDLRSWTAADTA
                     AQITQRKWEAARVAEQDRAYLEGTCVEWLRRYLENGKDTLERADPPKTHVTHHPISDHE
                     ATLRCWALGFYPAEITLTWQRDGEDQTQDTELVETRPAGDRTFQKWAAVVVPSGEEQRY
                     TCHVQHEGLPKPLTLRWEPSSQSTVPIVGIVAGLAVLAVVVIGAVVAAVMCRRKSSGGK
                     GGSYSQAACSDSAQGSDVSLTA"
     misc_RNA        complement(8579..8654)
                     /label=gRNA scaffold/Barcode
ORIGIN
        1 taagtcgaca atcaacctct ggattacaaa atttgtgaaa gattgactgg tattcttaac
       61 tatgttgctc cttttacgct atgtggatac gctgctttaa tgcctttgta tcatgctatt
      121 gcttcccgta tggctttcat tttctcctcc ttgtataaat cctggttgct gtctctttat
      181 gaggagttgt ggcccgttgt caggcaacgt ggcgtggtgt gcactgtgtt tgctgacgca
      241 acccccactg gttggggcat tgccaccacc tgtcagctcc tttccgggac tttcgctttc
      301 cccctcccta ttgccacggc ggaactcatc gccgcctgcc ttgcccgctg ctggacaggg
      361 gctcggctgt tgggcactga caattccgtg gtgttgtcgg ggaaatcatc gtcctttcct
      421 tggctgctcg cctgtgttgc cacctggatt ctgcgcggga cgtccttctg ctacgtccct
      481 tcggccctca atccagcgga ccttccttcc cgcggcctgc tgccggctct gcggcctctt
      541 ccgcgtcttc gccttcgccc tcagacgagt cggatctccc tttgggccgc ctccccgcgt
      601 cgactttaag accaatgact tacaaggcag ctgtagatct tagccacttt ttaaaagaaa
      661 aggggggact ggaagggcta attcactccc aacgaagaca agatctgctt tttgcttgta
      721 ctgggtctct ctggttagac cagatctgag cctgggagct ctctggctaa ctagggaacc
      781 cactgcttaa gcctcaataa agcttgcctt gagtgcttca agtagtgtgt gcccgtctgt
      841 tgtgtgactc tggtaactag agatccctca gaccctttta gtcagtgtgg aaaatctcta
      901 gcagtacgta tagtagttca tgtcatctta ttattcagta tttataactt gcaaagaaat
      961 gaatatcaga gagtgagagg aacttgttta ttgcagctta taatggttac aaataaagca
     1021 atagcatcac aaatttcaca aataaagcat ttttttcact gcattctagt tgtggtttgt
     1081 ccaaactcat caatgtatct tatcatgtct ggctctagct atcccgcccc taactccgcc
     1141 catcccgccc ctaactccgc ccagttccgc ccattctccg ccccatggct gactaatttt
     1201 ttttatttat gcagaggccg aggccgcctc ggcctctgag ctattccaga agtagtgagg
     1261 aggctttttt ggaggcctag ggacgtaccc aattcgccct atagtgagtc gtattacgcg
     1321 cgctcactgg ccgtcgtttt acaacgtcgt gactgggaaa accctggcgt tacccaactt
     1381 aatcgccttg cagcacatcc ccctttcgcc agctggcgta atagcgaaga ggcccgcacc
     1441 gatcgccctt cccaacagtt gcgcagcctg aatggcgaat gggacgcgcc ctgtagcggc
     1501 gcattaagcg cggcgggtgt ggtggttacg cgcagcgtga ccgctacact tgccagcgcc
     1561 ctagcgcccg ctcctttcgc tttcttccct tcctttctcg ccacgttcgc cggctttccc
     1621 cgtcaagctc taaatcgggg gctcccttta gggttccgat ttagtgcttt acggcacctc
     1681 gaccccaaaa aacttgatta gggtgatggt tcacgtagtg ggccatcgcc ctgatagacg
     1741 gtttttcgcc ctttgacgtt ggagtccacg ttctttaata gtggactctt gttccaaact
     1801 ggaacaacac tcaaccctat ctcggtctat tcttttgatt tataagggat tttgccgatt
     1861 tcggcctatt ggttaaaaaa tgagctgatt taacaaaaat ttaacgcgaa ttttaacaaa
     1921 atattaacgc ttacaattta ggtggcactt ttcggggaaa tgtgcgcgga acccctattt
     1981 gtttattttt ctaaatacat tcaaatatgt atccgctcat gagacaataa ccctgataaa
     2041 tgcttcaata atattgaaaa aggaagagta tgagtattca acatttccgt gtcgccctta
     2101 ttcccttttt tgcggcattt tgccttcctg tttttgctca cccagaaacg ctggtgaaag
     2161 taaaagatgc tgaagatcag ttgggtgcac gagtgggtta catcgaactg gatctcaaca
     2221 gcggtaagat ccttgagagt tttcgccccg aagaacgttt tccaatgatg agcactttta
     2281 aagttctgct atgtggcgcg gtattatccc gtattgacgc cgggcaagag caactcggtc
     2341 gccgcataca ctattctcag aatgacttgg ttgagtactc accagtcaca gaaaagcatc
     2401 ttacggatgg catgacagta agagaattat gcagtgctgc cataaccatg agtgataaca
     2461 ctgcggccaa cttacttctg acaacgatcg gaggaccgaa ggagctaacc gcttttttgc
     2521 acaacatggg ggatcatgta actcgccttg atcgttggga accggagctg aatgaagcca
     2581 taccaaacga cgagcgtgac accacgatgc ctgtagcaat ggcaacaacg ttgcgcaaac
     2641 tattaactgg cgaactactt actctagctt cccggcaaca attaatagac tggatggagg
     2701 cggataaagt tgcaggacca cttctgcgct cggcccttcc ggctggctgg tttattgctg
     2761 ataaatctgg agccggtgag cgtgggtctc gcggtatcat tgcagcactg gggccagatg
     2821 gtaagccctc ccgtatcgta gttatctaca cgacggggag tcaggcaact atggatgaac
     2881 gaaatagaca gatcgctgag ataggtgcct cactgattaa gcattggtaa ctgtcagacc
     2941 aagtttactc atatatactt tagattgatt taaaacttca tttttaattt aaaaggatct
     3001 aggtgaagat cctttttgat aatctcatga ccaaaatccc ttaacgtgag ttttcgttcc
     3061 actgagcgtc agaccccgta gaaaagatca aaggatcttc ttgagatcct ttttttctgc
     3121 gcgtaatctg ctgcttgcaa acaaaaaaac caccgctacc agcggtggtt tgtttgccgg
     3181 atcaagagct accaactctt tttccgaagg taactggctt cagcagagcg cagataccaa
     3241 atactgttct tctagtgtag ccgtagttag gccaccactt caagaactct gtagcaccgc
     3301 ctacatacct cgctctgcta atcctgttac cagtggctgc tgccagtggc gataagtcgt
     3361 gtcttaccgg gttggactca agacgatagt taccggataa ggcgcagcgg tcgggctgaa
     3421 cggggggttc gtgcacacag cccagcttgg agcgaacgac ctacaccgaa ctgagatacc
     3481 tacagcgtga gctatgagaa agcgccacgc ttcccgaagg gagaaaggcg gacaggtatc
     3541 cggtaagcgg cagggtcgga acaggagagc gcacgaggga gcttccaggg ggaaacgcct
     3601 ggtatcttta tagtcctgtc gggtttcgcc acctctgact tgagcgtcga tttttgtgat
     3661 gctcgtcagg ggggcggagc ctatggaaaa acgccagcaa cgcggccttt ttacggttcc
     3721 tggccttttg ctggcctttt gctcacatgt tctttcctgc gttatcccct gattctgtgg
     3781 ataaccgtat taccgccttt gagtgagctg ataccgctcg ccgcagccga acgaccgagc
     3841 gcagcgagtc agtgagcgag gaagcggaag agcgcccaat acgcaaaccg cctctccccg
     3901 cgcgttggcc gattcattaa tgcagctggc acgacaggtt tcccgactgg aaagcgggca
     3961 gtgagcgcaa cgcaattaat gtgagttagc tcactcatta ggcaccccag gctttacact
     4021 ttatgcttcc ggctcgtatg ttgtgtggaa ttgtgagcgg ataacaattt cacacaggaa
     4081 acagctatga ccatgattac gccaagcgcg caattaaccc tcactaaagg gaacaaaagc
     4141 tggagctgca agcttaatgt agtcttatgc aatactcttg tagtcttgca acatggtaac
     4201 gatgagttag caacatgcct tacaaggaga gaaaaagcac cgtgcatgcc gattggtgga
     4261 agtaaggtgg tacgatcgtg ccttattagg aaggcaacag acgggtctga catggattgg
     4321 acgaaccact gaattgccgc attgcagaga tattgtattt aagtgcctag ctcgatacat
     4381 aaacgggtct ctctggttag accagatctg agcctgggag ctctctggct aactagggaa
     4441 cccactgctt aagcctcaat aaagcttgcc ttgagtgctt caagtagtgt gtgcccgtct
     4501 gttgtgtgac tctggtaact agagatccct cagacccttt tagtcagtgt ggaaaatctc
     4561 tagcagtggc gcccgaacag ggacttgaaa gcgaaaggga aaccagagga gctctctcga
     4621 cgcaggactc ggcttgctga agcgcgcacg gcaagaggcg aggggcggcg actggtgagt
     4681 acgccaaaaa ttttgactag cggaggctag aaggagagag atgggtgcga gagcgtcagt
     4741 attaagcggg ggagaattag atcgcgatgg gaaaaaattc ggttaaggcc agggggaaag
     4801 aaaaaatata aattaaaaca tatagtatgg gcaagcaggg agctagaacg attcgcagtt
     4861 aatcctggcc tgttagaaac atcagaaggc tgtagacaaa tactgggaca gctacaacca
     4921 tcccttcaga caggatcaga agaacttaga tcattatata atacagtagc aaccctctat
     4981 tgtgtgcatc aaaggataga gataaaagac accaaggaag ctttagacaa gatagaggaa
     5041 gagcaaaaca aaagtaagac caccgcacag caagcggccg ctgatcttca gacctggagg
     5101 aggagatatg agggacaatt ggagaagtga attatataaa tataaagtag taaaaattga
     5161 accattagga gtagcaccca ccaaggcaaa gagaagagtg gtgcagagag aaaaaagagc
     5221 agtgggaata ggagctttgt tccttgggtt cttgggagca gcaggaagca ctatgggcgc
     5281 agcgtcaatg acgctgacgg tacaggccag acaattattg tctggtatag tgcagcagca
     5341 gaacaatttg ctgagggcta ttgaggcgca acagcatctg ttgcaactca cagtctgggg
     5401 catcaagcag ctccaggcaa gaatcctggc tgtggaaaga tacctaaagg atcaacagct
     5461 cctggggatt tggggttgct ctggaaaact catttgcacc actgctgtgc cttggaatgc
     5521 tagttggagt aataaatctc tggaacagat ttggaatcac acgacctgga tggagtggga
     5581 cagagaaatt aacaattaca caagcttaat acactcctta attgaagaat cgcaaaacca
     5641 gcaagaaaag aatgaacaag aattattgga attagataaa tgggcaagtt tgtggaattg
     5701 gtttaacata acaaattggc tgtggtatat aaaattattc ataatgatag taggaggctt
     5761 ggtaggttta agaatagttt ttgctgtact ttctatagtg aatagagtta ggcagggata
     5821 ttcaccatta tcgtttcaga cccacctccc aaccccgagg ggacccgaca ggcccgaagg
     5881 aatagaagaa gaaggtggag agagagacag agacagatcc attcgattag tgaacggatc
     5941 tcgacggtat cggttaactt ttaaaagaaa aggggggatt ggggggtaca gtgcagggga
     6001 aagaatagta gacataatag caacagacat acaaactaaa gaattacaaa aacaaattac
     6061 aaaaattcaa aattttatcg atgagtaatt catacaaaag gactcgcccc tgccttgggg
     6121 aatcccaggg accgtcgtta aactcccact aacgtagaac ccagagatcg ctgcgttccc
     6181 gccccctcac ccgcccgctc tcgtcatcac tgaggtggag aagagcatgc gtgaggctcc
     6241 ggtgcccgtc agtgggcaga gcgcacatcg cccacagtcc ccgagaagtt ggggggaggg
     6301 gtcggcaatt gaaccggtgc ctagagaagg tggcgcgggg taaactggga aagtgatgtc
     6361 gtgtactggc tccgcctttt tcccgagggt gggggagaac cgtatataag tgcagtagtc
     6421 gccgtgaacg ttctttttcg caacgggttt gccgccagaa cacaggtaag tgccgtgtgt
     6481 ggttcccgcg ggcctggcct ctttacgggt tatggccctt gcgtgccttg aattacttcc
     6541 acgcccctgg ctgcagtacg tgattcttga tcccgagctt cgggttggaa gtgggtggga
     6601 gagttcgagg ccttgcgctt aaggagcccc ttcgcctcgt gcttgagttg aggcctggct
     6661 tgggcgctgg ggccgccgcg tgcgaatctg gtggcacctt cgcgcctgtc tcgctgcttt
     6721 cgataagtct ctagccattt aaaatttttg atgacctgct gcgacgcttt ttttctggca
     6781 agatagtctt gtaaatgcgg gccaagatct gcacactggt atttcggttt ttggggccgc
     6841 gggcggcgac ggggcccgtg cgtcccagcg cacatgttcg gcgaggcggg gcctgcgagc
     6901 gcggccaccg agaatcggac gggggtagtc tcaagctggc cggcctgctc tggtgcctgg
     6961 cctcgcgccg ccgtgtatcg ccccgccctg ggcggcaagg ctggcccggt cggcaccagt
     7021 tgcgtgagcg gaaagatggc cgcttcccgg ccctgctgca gggagctcaa aatggaggac
     7081 gcggcgctcg ggagagcggg cgggtgagtc acccacacaa aggaaaaggg cctttccgtc
     7141 ctcagccgtc gcttcatgtg actccacgga gtaccgggcg ccgtccaggc acctcgatta
     7201 gttctcaagc ttttggagta cgtcgtcttt aggttggggg gaggggtttt atgcgatgga
     7261 gtttccccac actgagtggg tggagactga agttaggcca gcttggcact tgatgtaatt
     7321 ctccttggaa tttgcccttt ttgagtttgg atcttggttc attctcaagc ctcagacagt
     7381 ggttcaaagt ttttttcttc catttcaggt gtcgtgaccc tagcgctacc tctagagcca
     7441 ccatgctggt catggcccct agaaccgtgc tgctgctgct gagcgccgcc ctggccctga
     7501 ccgagacatg ggccggctcc cactctatga ggtacttcga cacagcaatg tcccggcccg
     7561 gaagaggaga gcctcgcttt atctctgtgg gctatgtgga cgatacccag ttcgtgcggt
     7621 ttgacagcga tgcagcatcc cctagggagg agcctagggc accatggatc gagcaggagg
     7681 gcccagagta ctgggacaga aacacacaga tcttcaagac caatacacag accgataggg
     7741 agagcctgag gaacctgcgc ggctactata atcagtctga ggccggcagc cacaccctgc
     7801 agtccatgta cggatgcgac gtgggaccag atggccggct gctgagaggc cacaaccagt
     7861 acgcctatga cggcaaggat tatatcgccc tgaatgagga cctgcgctcc tggacagcag
     7921 cagataccgc agcccagatc acacagagga agtgggaggc agcccgcgtg gcagagcagg
     7981 acagggccta cctggagggc acatgcgtgg agtggctgcg gagatatctg gagaacggca
     8041 aggacaccct ggagagagcc gatcccccta agacacacgt gacccaccac ccaatctctg
     8101 atcacgaggc caccctgaga tgttgggccc tgggcttcta ccccgccgag atcacactga
     8161 cctggcagag ggacggcgag gatcagacac aggacaccga gctggtggag acacggcccg
     8221 caggcgatag aacctttcag aagtgggcag cagtggtggt gccttctgga gaggagcagc
     8281 ggtacacatg tcacgtgcag cacgagggcc tgccaaagcc cctgacactg cggtgggagc
     8341 caagctccca gagcaccgtg cccatcgtgg gaatcgtggc aggcctggcc gtgctggccg
     8401 tggtggtcat cggagcagtg gtggcagccg tgatgtgcag gaggaagtct agcggaggca
     8461 agggaggaag ctattcccag gcagcatgtt ctgacagcgc ccagggatcc gacgtgagcc
     8521 tgaccgcctg aggcgcgcct cccagagcca ccgttacacc acggccagca ttcgggaggc
     8581 accgactcgg tgccactttt tcaagttgat aacggactag ccttatttta acttgctatt
     8641 tctagctcta aaactttttt tgactagatc ttgagacaaa tggcagtatt catccacaag
     8701 gtacgctaaa cgcctgcaac caggacgcgt
//

Lenti plKO_EF1A_ORF_HLA_B*35 01

LOCUS plKO_EF1A_ORF_HLA_B*35 01                8730 bp DNA     circular SYN 24-OCT-2022
DEFINITION  synthetic circular DNA
ACCESSION   .
VERSION     .
KEYWORDS    .
SOURCE      synthetic DNA construct
  ORGANISM  synthetic DNA construct
REFERENCE   1  (bases 1 to 8730)
  AUTHORS   .
  TITLE     Direct Submission
  JOURNAL   Exported Oct 25, 2022 from SnapGene 6.0.5
            https://www.snapgene.com
COMMENT     U515NEI230-158 DNA889102
FEATURES             Location/Qualifiers
     source          1..8730
                     /mol_type="other DNA"
                     /organism="synthetic DNA construct"
     misc_feature    10..598
                     /label=WPRE
     CDS             complement(481..492)
                     /label=Factor Xa site
     LTR             670..903
                     /label=3' LTR (Delta-U3)
     polyA_signal    981..1102
                     /label=SV40 poly(A) signal
     rep_origin      1142..1277
                     /label=SV40 ori
     promoter        complement(1298..1316)
                     /label=T7 promoter
     primer_bind     complement(1326..1342)
                     /label=M13 fwd
     rep_origin      1484..1939
                     /label=f1 ori
     promoter        1965..2069
                     /label=AmpR promoter
     CDS             2070..2138
                     /label=AmpR
     CDS             2139..2930
                     /label=AmpR
     rep_origin      3101..3689
                     /label=ori
     promoter        4019..4036
                     /label=lac promoter
     protein_bind    4051..4067
                     /label=lac operator
     primer_bind     4075..4091
                     /label=M13 rev
     promoter        4112..4130
                     /label=T3 promoter
     promoter        4158..4384
                     /label=RSV promoter
     LTR             4385..4565
                     /label=5' LTR (truncated)
     misc_feature    4612..4737
                     /label=HIV-1 Psi
     misc_feature    5230..5463
                     /label=RRE
     misc_feature    5959..6076
                     /label=cPPT/CTS
     promoter        6236..7417
                     /label=EF-1-alpha promoter
     intron          6466..7408
                     /label=EF-1-alpha intron A
     CDS             7443..8531
                     /codon_start=1
                     /label=HLA-B*35:01
                     /translation="MRVTAPRTVLLLLWGAVALTETWAGSHSMRYFYTAMSRPGRGEPR
                     FIAVGYVDDTQFVRFDSDAASPRTEPRAPWIEQEGPEYWDRNTQIFKTNTQTYRESLRN
                     LRGYYNQSEAGSHIIQRMYGCDLGPDGRLLRGHDQSAYDGKDYIALNEDLSSWTAADTA
                     AQITQRKWEAARVAEQLRAYLEGLCVEWLRRYLENGKETLQRADPPKTHVTHHPVSDHE
                     ATLRCWALGFYPAEITLTWQRDGEDQTQDTELVETRPAGDRTFQKWAAVVVPSGEEQRY
                     TCHVQHEGLPKPLTLRWEPSSQSTIPIVGIVAGLAVLAVVVIGAVVATVMCRRKSSGGK
                     GGSYSQAASSDSAQGSDVSLTA"
     misc_RNA        complement(8579..8654)
                     /label=gRNA scaffold/Barcode
ORIGIN
        1 taagtcgaca atcaacctct ggattacaaa atttgtgaaa gattgactgg tattcttaac
       61 tatgttgctc cttttacgct atgtggatac gctgctttaa tgcctttgta tcatgctatt
      121 gcttcccgta tggctttcat tttctcctcc ttgtataaat cctggttgct gtctctttat
      181 gaggagttgt ggcccgttgt caggcaacgt ggcgtggtgt gcactgtgtt tgctgacgca
      241 acccccactg gttggggcat tgccaccacc tgtcagctcc tttccgggac tttcgctttc
      301 cccctcccta ttgccacggc ggaactcatc gccgcctgcc ttgcccgctg ctggacaggg
      361 gctcggctgt tgggcactga caattccgtg gtgttgtcgg ggaaatcatc gtcctttcct
      421 tggctgctcg cctgtgttgc cacctggatt ctgcgcggga cgtccttctg ctacgtccct
      481 tcggccctca atccagcgga ccttccttcc cgcggcctgc tgccggctct gcggcctctt
      541 ccgcgtcttc gccttcgccc tcagacgagt cggatctccc tttgggccgc ctccccgcgt
      601 cgactttaag accaatgact tacaaggcag ctgtagatct tagccacttt ttaaaagaaa
      661 aggggggact ggaagggcta attcactccc aacgaagaca agatctgctt tttgcttgta
      721 ctgggtctct ctggttagac cagatctgag cctgggagct ctctggctaa ctagggaacc
      781 cactgcttaa gcctcaataa agcttgcctt gagtgcttca agtagtgtgt gcccgtctgt
      841 tgtgtgactc tggtaactag agatccctca gaccctttta gtcagtgtgg aaaatctcta
      901 gcagtacgta tagtagttca tgtcatctta ttattcagta tttataactt gcaaagaaat
      961 gaatatcaga gagtgagagg aacttgttta ttgcagctta taatggttac aaataaagca
     1021 atagcatcac aaatttcaca aataaagcat ttttttcact gcattctagt tgtggtttgt
     1081 ccaaactcat caatgtatct tatcatgtct ggctctagct atcccgcccc taactccgcc
     1141 catcccgccc ctaactccgc ccagttccgc ccattctccg ccccatggct gactaatttt
     1201 ttttatttat gcagaggccg aggccgcctc ggcctctgag ctattccaga agtagtgagg
     1261 aggctttttt ggaggcctag ggacgtaccc aattcgccct atagtgagtc gtattacgcg
     1321 cgctcactgg ccgtcgtttt acaacgtcgt gactgggaaa accctggcgt tacccaactt
     1381 aatcgccttg cagcacatcc ccctttcgcc agctggcgta atagcgaaga ggcccgcacc
     1441 gatcgccctt cccaacagtt gcgcagcctg aatggcgaat gggacgcgcc ctgtagcggc
     1501 gcattaagcg cggcgggtgt ggtggttacg cgcagcgtga ccgctacact tgccagcgcc
     1561 ctagcgcccg ctcctttcgc tttcttccct tcctttctcg ccacgttcgc cggctttccc
     1621 cgtcaagctc taaatcgggg gctcccttta gggttccgat ttagtgcttt acggcacctc
     1681 gaccccaaaa aacttgatta gggtgatggt tcacgtagtg ggccatcgcc ctgatagacg
     1741 gtttttcgcc ctttgacgtt ggagtccacg ttctttaata gtggactctt gttccaaact
     1801 ggaacaacac tcaaccctat ctcggtctat tcttttgatt tataagggat tttgccgatt
     1861 tcggcctatt ggttaaaaaa tgagctgatt taacaaaaat ttaacgcgaa ttttaacaaa
     1921 atattaacgc ttacaattta ggtggcactt ttcggggaaa tgtgcgcgga acccctattt
     1981 gtttattttt ctaaatacat tcaaatatgt atccgctcat gagacaataa ccctgataaa
     2041 tgcttcaata atattgaaaa aggaagagta tgagtattca acatttccgt gtcgccctta
     2101 ttcccttttt tgcggcattt tgccttcctg tttttgctca cccagaaacg ctggtgaaag
     2161 taaaagatgc tgaagatcag ttgggtgcac gagtgggtta catcgaactg gatctcaaca
     2221 gcggtaagat ccttgagagt tttcgccccg aagaacgttt tccaatgatg agcactttta
     2281 aagttctgct atgtggcgcg gtattatccc gtattgacgc cgggcaagag caactcggtc
     2341 gccgcataca ctattctcag aatgacttgg ttgagtactc accagtcaca gaaaagcatc
     2401 ttacggatgg catgacagta agagaattat gcagtgctgc cataaccatg agtgataaca
     2461 ctgcggccaa cttacttctg acaacgatcg gaggaccgaa ggagctaacc gcttttttgc
     2521 acaacatggg ggatcatgta actcgccttg atcgttggga accggagctg aatgaagcca
     2581 taccaaacga cgagcgtgac accacgatgc ctgtagcaat ggcaacaacg ttgcgcaaac
     2641 tattaactgg cgaactactt actctagctt cccggcaaca attaatagac tggatggagg
     2701 cggataaagt tgcaggacca cttctgcgct cggcccttcc ggctggctgg tttattgctg
     2761 ataaatctgg agccggtgag cgtgggtctc gcggtatcat tgcagcactg gggccagatg
     2821 gtaagccctc ccgtatcgta gttatctaca cgacggggag tcaggcaact atggatgaac
     2881 gaaatagaca gatcgctgag ataggtgcct cactgattaa gcattggtaa ctgtcagacc
     2941 aagtttactc atatatactt tagattgatt taaaacttca tttttaattt aaaaggatct
     3001 aggtgaagat cctttttgat aatctcatga ccaaaatccc ttaacgtgag ttttcgttcc
     3061 actgagcgtc agaccccgta gaaaagatca aaggatcttc ttgagatcct ttttttctgc
     3121 gcgtaatctg ctgcttgcaa acaaaaaaac caccgctacc agcggtggtt tgtttgccgg
     3181 atcaagagct accaactctt tttccgaagg taactggctt cagcagagcg cagataccaa
     3241 atactgttct tctagtgtag ccgtagttag gccaccactt caagaactct gtagcaccgc
     3301 ctacatacct cgctctgcta atcctgttac cagtggctgc tgccagtggc gataagtcgt
     3361 gtcttaccgg gttggactca agacgatagt taccggataa ggcgcagcgg tcgggctgaa
     3421 cggggggttc gtgcacacag cccagcttgg agcgaacgac ctacaccgaa ctgagatacc
     3481 tacagcgtga gctatgagaa agcgccacgc ttcccgaagg gagaaaggcg gacaggtatc
     3541 cggtaagcgg cagggtcgga acaggagagc gcacgaggga gcttccaggg ggaaacgcct
     3601 ggtatcttta tagtcctgtc gggtttcgcc acctctgact tgagcgtcga tttttgtgat
     3661 gctcgtcagg ggggcggagc ctatggaaaa acgccagcaa cgcggccttt ttacggttcc
     3721 tggccttttg ctggcctttt gctcacatgt tctttcctgc gttatcccct gattctgtgg
     3781 ataaccgtat taccgccttt gagtgagctg ataccgctcg ccgcagccga acgaccgagc
     3841 gcagcgagtc agtgagcgag gaagcggaag agcgcccaat acgcaaaccg cctctccccg
     3901 cgcgttggcc gattcattaa tgcagctggc acgacaggtt tcccgactgg aaagcgggca
     3961 gtgagcgcaa cgcaattaat gtgagttagc tcactcatta ggcaccccag gctttacact
     4021 ttatgcttcc ggctcgtatg ttgtgtggaa ttgtgagcgg ataacaattt cacacaggaa
     4081 acagctatga ccatgattac gccaagcgcg caattaaccc tcactaaagg gaacaaaagc
     4141 tggagctgca agcttaatgt agtcttatgc aatactcttg tagtcttgca acatggtaac
     4201 gatgagttag caacatgcct tacaaggaga gaaaaagcac cgtgcatgcc gattggtgga
     4261 agtaaggtgg tacgatcgtg ccttattagg aaggcaacag acgggtctga catggattgg
     4321 acgaaccact gaattgccgc attgcagaga tattgtattt aagtgcctag ctcgatacat
     4381 aaacgggtct ctctggttag accagatctg agcctgggag ctctctggct aactagggaa
     4441 cccactgctt aagcctcaat aaagcttgcc ttgagtgctt caagtagtgt gtgcccgtct
     4501 gttgtgtgac tctggtaact agagatccct cagacccttt tagtcagtgt ggaaaatctc
     4561 tagcagtggc gcccgaacag ggacttgaaa gcgaaaggga aaccagagga gctctctcga
     4621 cgcaggactc ggcttgctga agcgcgcacg gcaagaggcg aggggcggcg actggtgagt
     4681 acgccaaaaa ttttgactag cggaggctag aaggagagag atgggtgcga gagcgtcagt
     4741 attaagcggg ggagaattag atcgcgatgg gaaaaaattc ggttaaggcc agggggaaag
     4801 aaaaaatata aattaaaaca tatagtatgg gcaagcaggg agctagaacg attcgcagtt
     4861 aatcctggcc tgttagaaac atcagaaggc tgtagacaaa tactgggaca gctacaacca
     4921 tcccttcaga caggatcaga agaacttaga tcattatata atacagtagc aaccctctat
     4981 tgtgtgcatc aaaggataga gataaaagac accaaggaag ctttagacaa gatagaggaa
     5041 gagcaaaaca aaagtaagac caccgcacag caagcggccg ctgatcttca gacctggagg
     5101 aggagatatg agggacaatt ggagaagtga attatataaa tataaagtag taaaaattga
     5161 accattagga gtagcaccca ccaaggcaaa gagaagagtg gtgcagagag aaaaaagagc
     5221 agtgggaata ggagctttgt tccttgggtt cttgggagca gcaggaagca ctatgggcgc
     5281 agcgtcaatg acgctgacgg tacaggccag acaattattg tctggtatag tgcagcagca
     5341 gaacaatttg ctgagggcta ttgaggcgca acagcatctg ttgcaactca cagtctgggg
     5401 catcaagcag ctccaggcaa gaatcctggc tgtggaaaga tacctaaagg atcaacagct
     5461 cctggggatt tggggttgct ctggaaaact catttgcacc actgctgtgc cttggaatgc
     5521 tagttggagt aataaatctc tggaacagat ttggaatcac acgacctgga tggagtggga
     5581 cagagaaatt aacaattaca caagcttaat acactcctta attgaagaat cgcaaaacca
     5641 gcaagaaaag aatgaacaag aattattgga attagataaa tgggcaagtt tgtggaattg
     5701 gtttaacata acaaattggc tgtggtatat aaaattattc ataatgatag taggaggctt
     5761 ggtaggttta agaatagttt ttgctgtact ttctatagtg aatagagtta ggcagggata
     5821 ttcaccatta tcgtttcaga cccacctccc aaccccgagg ggacccgaca ggcccgaagg
     5881 aatagaagaa gaaggtggag agagagacag agacagatcc attcgattag tgaacggatc
     5941 tcgacggtat cggttaactt ttaaaagaaa aggggggatt ggggggtaca gtgcagggga
     6001 aagaatagta gacataatag caacagacat acaaactaaa gaattacaaa aacaaattac
     6061 aaaaattcaa aattttatcg atgagtaatt catacaaaag gactcgcccc tgccttgggg
     6121 aatcccaggg accgtcgtta aactcccact aacgtagaac ccagagatcg ctgcgttccc
     6181 gccccctcac ccgcccgctc tcgtcatcac tgaggtggag aagagcatgc gtgaggctcc
     6241 ggtgcccgtc agtgggcaga gcgcacatcg cccacagtcc ccgagaagtt ggggggaggg
     6301 gtcggcaatt gaaccggtgc ctagagaagg tggcgcgggg taaactggga aagtgatgtc
     6361 gtgtactggc tccgcctttt tcccgagggt gggggagaac cgtatataag tgcagtagtc
     6421 gccgtgaacg ttctttttcg caacgggttt gccgccagaa cacaggtaag tgccgtgtgt
     6481 ggttcccgcg ggcctggcct ctttacgggt tatggccctt gcgtgccttg aattacttcc
     6541 acgcccctgg ctgcagtacg tgattcttga tcccgagctt cgggttggaa gtgggtggga
     6601 gagttcgagg ccttgcgctt aaggagcccc ttcgcctcgt gcttgagttg aggcctggct
     6661 tgggcgctgg ggccgccgcg tgcgaatctg gtggcacctt cgcgcctgtc tcgctgcttt
     6721 cgataagtct ctagccattt aaaatttttg atgacctgct gcgacgcttt ttttctggca
     6781 agatagtctt gtaaatgcgg gccaagatct gcacactggt atttcggttt ttggggccgc
     6841 gggcggcgac ggggcccgtg cgtcccagcg cacatgttcg gcgaggcggg gcctgcgagc
     6901 gcggccaccg agaatcggac gggggtagtc tcaagctggc cggcctgctc tggtgcctgg
     6961 cctcgcgccg ccgtgtatcg ccccgccctg ggcggcaagg ctggcccggt cggcaccagt
     7021 tgcgtgagcg gaaagatggc cgcttcccgg ccctgctgca gggagctcaa aatggaggac
     7081 gcggcgctcg ggagagcggg cgggtgagtc acccacacaa aggaaaaggg cctttccgtc
     7141 ctcagccgtc gcttcatgtg actccacgga gtaccgggcg ccgtccaggc acctcgatta
     7201 gttctcaagc ttttggagta cgtcgtcttt aggttggggg gaggggtttt atgcgatgga
     7261 gtttccccac actgagtggg tggagactga agttaggcca gcttggcact tgatgtaatt
     7321 ctccttggaa tttgcccttt ttgagtttgg atcttggttc attctcaagc ctcagacagt
     7381 ggttcaaagt ttttttcttc catttcaggt gtcgtgaccc tagcgctacc tctagagcca
     7441 ccatgcgggt gacagcccct agaaccgtgc tgctgctgct gtggggagca gtggccctga
     7501 ccgagacatg ggccggctct cacagcatga ggtacttcta tacagcaatg agccggcccg
     7561 gaagaggaga gcctcggttc atcgccgtgg gctacgtgga cgatacacag ttcgtgaggt
     7621 ttgactctga tgcagcaagc cctaggaccg agcctagggc accatggatc gagcaggagg
     7681 gcccagagta ctgggaccgc aacacccaga tcttcaagac caatacacag acctatcggg
     7741 agtctctgag gaacctgcgc ggctactata atcagtccga ggccggctct cacatcatcc
     7801 agcgcatgta tggatgcgac ctgggaccag atggccggct gctgagaggc cacgatcaga
     7861 gcgcctacga cggcaaggat tatatcgccc tgaacgagga cctgagctcc tggacagcag
     7921 cagataccgc agcacagatc acccagagga agtgggaggc agccagagtg gcagagcagc
     7981 tgagggcata cctggagggc ctgtgcgtgg agtggctgcg gagatatctg gagaatggca
     8041 aggagacact gcagagggca gacccaccta agacacacgt gacccaccac ccagtgtccg
     8101 atcacgaggc caccctgagg tgttgggcac tgggcttcta cccagccgag atcacactga
     8161 cctggcagcg ggacggcgag gatcagacac aggacaccga gctggtggag acaaggcccg
     8221 caggcgatcg cacctttcag aagtgggcag cagtggtggt gcctagcgga gaggagcaga
     8281 gatatacctg ccacgtgcag cacgagggcc tgccaaagcc cctgacactg agatgggagc
     8341 catctagcca gtccaccatc ccaatcgtgg gaatcgtggc aggcctggcc gtgctggccg
     8401 tggtggtcat cggagcagtg gtggcaaccg tgatgtgcag gaggaagtcc tctggaggca
     8461 agggaggatc ctactctcag gcagcaagct ccgactctgc ccagggaagc gacgtgagcc
     8521 tgaccgcatg aggcgcgcct cccagagcca ccgttacacc agtactggag aaccagtagc
     8581 accgactcgg tgccactttt tcaagttgat aacggactag ccttatttta acttgctatt
     8641 tctagctcta aaactttttt tgactagatc ttgagacaaa tggcagtatt catccacaag
     8701 gtacgctaaa cgcctgcaac caggacgcgt
//

Lenti plKO_EF1A_ORF_HLA_B*51 01

LOCUS plKO_EF1A_ORF_HLA_B*51 01                8730 bp DNA     circular SYN 24-OCT-2022
DEFINITION  synthetic circular DNA
ACCESSION   .
VERSION     .
KEYWORDS    .
SOURCE      synthetic DNA construct
  ORGANISM  synthetic DNA construct
REFERENCE   1  (bases 1 to 8730)
  AUTHORS   .
  TITLE     Direct Submission
  JOURNAL   Exported Oct 25, 2022 from SnapGene 6.0.5
            https://www.snapgene.com
COMMENT     U515NEI230-174 DNA889104
FEATURES             Location/Qualifiers
     source          1..8730
                     /mol_type="other DNA"
                     /organism="synthetic DNA construct"
     misc_feature    10..598
                     /label=WPRE
     CDS             complement(481..492)
                     /label=Factor Xa site
     LTR             670..903
                     /label=3' LTR (Delta-U3)
     polyA_signal    981..1102
                     /label=SV40 poly(A) signal
     rep_origin      1142..1277
                     /label=SV40 ori
     promoter        complement(1298..1316)
                     /label=T7 promoter
     primer_bind     complement(1326..1342)
                     /label=M13 fwd
     rep_origin      1484..1939
                     /label=f1 ori
     promoter        1965..2069
                     /label=AmpR promoter
     CDS             2070..2138
                     /label=AmpR
     CDS             2139..2930
                     /label=AmpR
     rep_origin      3101..3689
                     /label=ori
     promoter        4019..4036
                     /label=lac promoter
     protein_bind    4051..4067
                     /label=lac operator
     primer_bind     4075..4091
                     /label=M13 rev
     promoter        4112..4130
                     /label=T3 promoter
     promoter        4158..4384
                     /label=RSV promoter
     LTR             4385..4565
                     /label=5' LTR (truncated)
     misc_feature    4612..4737
                     /label=HIV-1 Psi
     misc_feature    5230..5463
                     /label=RRE
     misc_feature    5959..6076
                     /label=cPPT/CTS
     promoter        6236..7417
                     /label=EF-1-alpha promoter
     intron          6466..7408
                     /label=EF-1-alpha intron A
     CDS             7443..8531
                     /codon_start=1
                     /label=HLA-B*51:01
                     /translation="MRVTAPRTVLLLLWGAVALTETWAGSHSMRYFYTAMSRPGRGEPR
                     FIAVGYVDDTQFVRFDSDAASPRTEPRAPWIEQEGPEYWDRNTQIFKTNTQTYRENLRI
                     ALRYYNQSEAGSHTWQTMYGCDVGPDGRLLRGHNQYAYDGKDYIALNEDLSSWTAADTA
                     AQITQRKWEAAREAEQLRAYLEGLCVEWLRRHLENGKETLQRADPPKTHVTHHPVSDHE
                     ATLRCWALGFYPAEITLTWQRDGEDQTQDTELVETRPAGDRTFQKWAAVVVPSGEEQRY
                     TCHVQHEGLPKPLTLRWEPSSQSTIPIVGIVAGLAVLAVVVIGAVVATVMCRRKSSGGK
                     GGSYSQAASSDSAQGSDVSLTA"
     misc_RNA        complement(8579..8654)
                     /label=gRNA scaffold/Barcode
ORIGIN
        1 taagtcgaca atcaacctct ggattacaaa atttgtgaaa gattgactgg tattcttaac
       61 tatgttgctc cttttacgct atgtggatac gctgctttaa tgcctttgta tcatgctatt
      121 gcttcccgta tggctttcat tttctcctcc ttgtataaat cctggttgct gtctctttat
      181 gaggagttgt ggcccgttgt caggcaacgt ggcgtggtgt gcactgtgtt tgctgacgca
      241 acccccactg gttggggcat tgccaccacc tgtcagctcc tttccgggac tttcgctttc
      301 cccctcccta ttgccacggc ggaactcatc gccgcctgcc ttgcccgctg ctggacaggg
      361 gctcggctgt tgggcactga caattccgtg gtgttgtcgg ggaaatcatc gtcctttcct
      421 tggctgctcg cctgtgttgc cacctggatt ctgcgcggga cgtccttctg ctacgtccct
      481 tcggccctca atccagcgga ccttccttcc cgcggcctgc tgccggctct gcggcctctt
      541 ccgcgtcttc gccttcgccc tcagacgagt cggatctccc tttgggccgc ctccccgcgt
      601 cgactttaag accaatgact tacaaggcag ctgtagatct tagccacttt ttaaaagaaa
      661 aggggggact ggaagggcta attcactccc aacgaagaca agatctgctt tttgcttgta
      721 ctgggtctct ctggttagac cagatctgag cctgggagct ctctggctaa ctagggaacc
      781 cactgcttaa gcctcaataa agcttgcctt gagtgcttca agtagtgtgt gcccgtctgt
      841 tgtgtgactc tggtaactag agatccctca gaccctttta gtcagtgtgg aaaatctcta
      901 gcagtacgta tagtagttca tgtcatctta ttattcagta tttataactt gcaaagaaat
      961 gaatatcaga gagtgagagg aacttgttta ttgcagctta taatggttac aaataaagca
     1021 atagcatcac aaatttcaca aataaagcat ttttttcact gcattctagt tgtggtttgt
     1081 ccaaactcat caatgtatct tatcatgtct ggctctagct atcccgcccc taactccgcc
     1141 catcccgccc ctaactccgc ccagttccgc ccattctccg ccccatggct gactaatttt
     1201 ttttatttat gcagaggccg aggccgcctc ggcctctgag ctattccaga agtagtgagg
     1261 aggctttttt ggaggcctag ggacgtaccc aattcgccct atagtgagtc gtattacgcg
     1321 cgctcactgg ccgtcgtttt acaacgtcgt gactgggaaa accctggcgt tacccaactt
     1381 aatcgccttg cagcacatcc ccctttcgcc agctggcgta atagcgaaga ggcccgcacc
     1441 gatcgccctt cccaacagtt gcgcagcctg aatggcgaat gggacgcgcc ctgtagcggc
     1501 gcattaagcg cggcgggtgt ggtggttacg cgcagcgtga ccgctacact tgccagcgcc
     1561 ctagcgcccg ctcctttcgc tttcttccct tcctttctcg ccacgttcgc cggctttccc
     1621 cgtcaagctc taaatcgggg gctcccttta gggttccgat ttagtgcttt acggcacctc
     1681 gaccccaaaa aacttgatta gggtgatggt tcacgtagtg ggccatcgcc ctgatagacg
     1741 gtttttcgcc ctttgacgtt ggagtccacg ttctttaata gtggactctt gttccaaact
     1801 ggaacaacac tcaaccctat ctcggtctat tcttttgatt tataagggat tttgccgatt
     1861 tcggcctatt ggttaaaaaa tgagctgatt taacaaaaat ttaacgcgaa ttttaacaaa
     1921 atattaacgc ttacaattta ggtggcactt ttcggggaaa tgtgcgcgga acccctattt
     1981 gtttattttt ctaaatacat tcaaatatgt atccgctcat gagacaataa ccctgataaa
     2041 tgcttcaata atattgaaaa aggaagagta tgagtattca acatttccgt gtcgccctta
     2101 ttcccttttt tgcggcattt tgccttcctg tttttgctca cccagaaacg ctggtgaaag
     2161 taaaagatgc tgaagatcag ttgggtgcac gagtgggtta catcgaactg gatctcaaca
     2221 gcggtaagat ccttgagagt tttcgccccg aagaacgttt tccaatgatg agcactttta
     2281 aagttctgct atgtggcgcg gtattatccc gtattgacgc cgggcaagag caactcggtc
     2341 gccgcataca ctattctcag aatgacttgg ttgagtactc accagtcaca gaaaagcatc
     2401 ttacggatgg catgacagta agagaattat gcagtgctgc cataaccatg agtgataaca
     2461 ctgcggccaa cttacttctg acaacgatcg gaggaccgaa ggagctaacc gcttttttgc
     2521 acaacatggg ggatcatgta actcgccttg atcgttggga accggagctg aatgaagcca
     2581 taccaaacga cgagcgtgac accacgatgc ctgtagcaat ggcaacaacg ttgcgcaaac
     2641 tattaactgg cgaactactt actctagctt cccggcaaca attaatagac tggatggagg
     2701 cggataaagt tgcaggacca cttctgcgct cggcccttcc ggctggctgg tttattgctg
     2761 ataaatctgg agccggtgag cgtgggtctc gcggtatcat tgcagcactg gggccagatg
     2821 gtaagccctc ccgtatcgta gttatctaca cgacggggag tcaggcaact atggatgaac
     2881 gaaatagaca gatcgctgag ataggtgcct cactgattaa gcattggtaa ctgtcagacc
     2941 aagtttactc atatatactt tagattgatt taaaacttca tttttaattt aaaaggatct
     3001 aggtgaagat cctttttgat aatctcatga ccaaaatccc ttaacgtgag ttttcgttcc
     3061 actgagcgtc agaccccgta gaaaagatca aaggatcttc ttgagatcct ttttttctgc
     3121 gcgtaatctg ctgcttgcaa acaaaaaaac caccgctacc agcggtggtt tgtttgccgg
     3181 atcaagagct accaactctt tttccgaagg taactggctt cagcagagcg cagataccaa
     3241 atactgttct tctagtgtag ccgtagttag gccaccactt caagaactct gtagcaccgc
     3301 ctacatacct cgctctgcta atcctgttac cagtggctgc tgccagtggc gataagtcgt
     3361 gtcttaccgg gttggactca agacgatagt taccggataa ggcgcagcgg tcgggctgaa
     3421 cggggggttc gtgcacacag cccagcttgg agcgaacgac ctacaccgaa ctgagatacc
     3481 tacagcgtga gctatgagaa agcgccacgc ttcccgaagg gagaaaggcg gacaggtatc
     3541 cggtaagcgg cagggtcgga acaggagagc gcacgaggga gcttccaggg ggaaacgcct
     3601 ggtatcttta tagtcctgtc gggtttcgcc acctctgact tgagcgtcga tttttgtgat
     3661 gctcgtcagg ggggcggagc ctatggaaaa acgccagcaa cgcggccttt ttacggttcc
     3721 tggccttttg ctggcctttt gctcacatgt tctttcctgc gttatcccct gattctgtgg
     3781 ataaccgtat taccgccttt gagtgagctg ataccgctcg ccgcagccga acgaccgagc
     3841 gcagcgagtc agtgagcgag gaagcggaag agcgcccaat acgcaaaccg cctctccccg
     3901 cgcgttggcc gattcattaa tgcagctggc acgacaggtt tcccgactgg aaagcgggca
     3961 gtgagcgcaa cgcaattaat gtgagttagc tcactcatta ggcaccccag gctttacact
     4021 ttatgcttcc ggctcgtatg ttgtgtggaa ttgtgagcgg ataacaattt cacacaggaa
     4081 acagctatga ccatgattac gccaagcgcg caattaaccc tcactaaagg gaacaaaagc
     4141 tggagctgca agcttaatgt agtcttatgc aatactcttg tagtcttgca acatggtaac
     4201 gatgagttag caacatgcct tacaaggaga gaaaaagcac cgtgcatgcc gattggtgga
     4261 agtaaggtgg tacgatcgtg ccttattagg aaggcaacag acgggtctga catggattgg
     4321 acgaaccact gaattgccgc attgcagaga tattgtattt aagtgcctag ctcgatacat
     4381 aaacgggtct ctctggttag accagatctg agcctgggag ctctctggct aactagggaa
     4441 cccactgctt aagcctcaat aaagcttgcc ttgagtgctt caagtagtgt gtgcccgtct
     4501 gttgtgtgac tctggtaact agagatccct cagacccttt tagtcagtgt ggaaaatctc
     4561 tagcagtggc gcccgaacag ggacttgaaa gcgaaaggga aaccagagga gctctctcga
     4621 cgcaggactc ggcttgctga agcgcgcacg gcaagaggcg aggggcggcg actggtgagt
     4681 acgccaaaaa ttttgactag cggaggctag aaggagagag atgggtgcga gagcgtcagt
     4741 attaagcggg ggagaattag atcgcgatgg gaaaaaattc ggttaaggcc agggggaaag
     4801 aaaaaatata aattaaaaca tatagtatgg gcaagcaggg agctagaacg attcgcagtt
     4861 aatcctggcc tgttagaaac atcagaaggc tgtagacaaa tactgggaca gctacaacca
     4921 tcccttcaga caggatcaga agaacttaga tcattatata atacagtagc aaccctctat
     4981 tgtgtgcatc aaaggataga gataaaagac accaaggaag ctttagacaa gatagaggaa
     5041 gagcaaaaca aaagtaagac caccgcacag caagcggccg ctgatcttca gacctggagg
     5101 aggagatatg agggacaatt ggagaagtga attatataaa tataaagtag taaaaattga
     5161 accattagga gtagcaccca ccaaggcaaa gagaagagtg gtgcagagag aaaaaagagc
     5221 agtgggaata ggagctttgt tccttgggtt cttgggagca gcaggaagca ctatgggcgc
     5281 agcgtcaatg acgctgacgg tacaggccag acaattattg tctggtatag tgcagcagca
     5341 gaacaatttg ctgagggcta ttgaggcgca acagcatctg ttgcaactca cagtctgggg
     5401 catcaagcag ctccaggcaa gaatcctggc tgtggaaaga tacctaaagg atcaacagct
     5461 cctggggatt tggggttgct ctggaaaact catttgcacc actgctgtgc cttggaatgc
     5521 tagttggagt aataaatctc tggaacagat ttggaatcac acgacctgga tggagtggga
     5581 cagagaaatt aacaattaca caagcttaat acactcctta attgaagaat cgcaaaacca
     5641 gcaagaaaag aatgaacaag aattattgga attagataaa tgggcaagtt tgtggaattg
     5701 gtttaacata acaaattggc tgtggtatat aaaattattc ataatgatag taggaggctt
     5761 ggtaggttta agaatagttt ttgctgtact ttctatagtg aatagagtta ggcagggata
     5821 ttcaccatta tcgtttcaga cccacctccc aaccccgagg ggacccgaca ggcccgaagg
     5881 aatagaagaa gaaggtggag agagagacag agacagatcc attcgattag tgaacggatc
     5941 tcgacggtat cggttaactt ttaaaagaaa aggggggatt ggggggtaca gtgcagggga
     6001 aagaatagta gacataatag caacagacat acaaactaaa gaattacaaa aacaaattac
     6061 aaaaattcaa aattttatcg atgagtaatt catacaaaag gactcgcccc tgccttgggg
     6121 aatcccaggg accgtcgtta aactcccact aacgtagaac ccagagatcg ctgcgttccc
     6181 gccccctcac ccgcccgctc tcgtcatcac tgaggtggag aagagcatgc gtgaggctcc
     6241 ggtgcccgtc agtgggcaga gcgcacatcg cccacagtcc ccgagaagtt ggggggaggg
     6301 gtcggcaatt gaaccggtgc ctagagaagg tggcgcgggg taaactggga aagtgatgtc
     6361 gtgtactggc tccgcctttt tcccgagggt gggggagaac cgtatataag tgcagtagtc
     6421 gccgtgaacg ttctttttcg caacgggttt gccgccagaa cacaggtaag tgccgtgtgt
     6481 ggttcccgcg ggcctggcct ctttacgggt tatggccctt gcgtgccttg aattacttcc
     6541 acgcccctgg ctgcagtacg tgattcttga tcccgagctt cgggttggaa gtgggtggga
     6601 gagttcgagg ccttgcgctt aaggagcccc ttcgcctcgt gcttgagttg aggcctggct
     6661 tgggcgctgg ggccgccgcg tgcgaatctg gtggcacctt cgcgcctgtc tcgctgcttt
     6721 cgataagtct ctagccattt aaaatttttg atgacctgct gcgacgcttt ttttctggca
     6781 agatagtctt gtaaatgcgg gccaagatct gcacactggt atttcggttt ttggggccgc
     6841 gggcggcgac ggggcccgtg cgtcccagcg cacatgttcg gcgaggcggg gcctgcgagc
     6901 gcggccaccg agaatcggac gggggtagtc tcaagctggc cggcctgctc tggtgcctgg
     6961 cctcgcgccg ccgtgtatcg ccccgccctg ggcggcaagg ctggcccggt cggcaccagt
     7021 tgcgtgagcg gaaagatggc cgcttcccgg ccctgctgca gggagctcaa aatggaggac
     7081 gcggcgctcg ggagagcggg cgggtgagtc acccacacaa aggaaaaggg cctttccgtc
     7141 ctcagccgtc gcttcatgtg actccacgga gtaccgggcg ccgtccaggc acctcgatta
     7201 gttctcaagc ttttggagta cgtcgtcttt aggttggggg gaggggtttt atgcgatgga
     7261 gtttccccac actgagtggg tggagactga agttaggcca gcttggcact tgatgtaatt
     7321 ctccttggaa tttgcccttt ttgagtttgg atcttggttc attctcaagc ctcagacagt
     7381 ggttcaaagt ttttttcttc catttcaggt gtcgtgaccc tagcgctacc tctagagcca
     7441 ccatgagagt gacagcccct aggaccgtgc tgctgctgct gtggggagca gtggccctga
     7501 ccgagacatg ggccggctct cacagcatga ggtacttcta tacagcaatg agccggcccg
     7561 gaagaggaga gcctcggttt atcgccgtgg gctacgtgga cgatacacag ttcgtgagat
     7621 ttgactctga tgcagcaagc cctaggaccg agcctagagc accatggatc gagcaggagg
     7681 gcccagagta ctgggaccgc aacacccaga tcttcaagac caatacacag acctataggg
     7741 agaacctgcg catcgccctg cggtactata atcagtccga ggccggctct cacacatggc
     7801 agaccatgta cggctgcgac gtgggaccag atggcaggct gctgaggggc cacaatcagt
     7861 acgcctatga cggcaaggat tatatcgccc tgaacgagga cctgagctcc tggacagcag
     7921 cagataccgc agcacagatc acccagagaa agtgggaggc agcaagggag gcagagcagc
     7981 tgagggcata tctggagggc ctgtgcgtgg agtggctgcg gagacacctg gagaacggca
     8041 aggagacact gcagagagcc gaccccccta agacacacgt gacccaccac ccagtgtctg
     8101 atcacgaggc caccctgagg tgttgggcac tgggcttcta cccagcagag atcacactga
     8161 cctggcagcg cgacggcgag gatcagacac aggacaccga gctggtggag acacgccccg
     8221 caggcgatcg gacctttcag aagtgggcag cagtggtggt gcctagcgga gaggagcagc
     8281 ggtatacctg ccacgtgcag cacgagggcc tgccaaagcc cctgacactg agatgggagc
     8341 catctagcca gtccaccatc ccaatcgtgg gaatcgtggc aggcctggcc gtgctggccg
     8401 tggtggtcat cggagcagtg gtggcaaccg tgatgtgcag gaggaagtcc tctggaggca
     8461 agggaggatc ctactctcag gcagcaagct ccgactccgc ccagggaagc gacgtgagcc
     8521 tgaccgcctg aggcgcgcct cccagagcca ccgttacaca acattctccc ggtatcccgc
     8581 accgactcgg tgccactttt tcaagttgat aacggactag ccttatttta acttgctatt
     8641 tctagctcta aaactttttt tgactagatc ttgagacaaa tggcagtatt catccacaag
     8701 gtacgctaaa cgcctgcaac caggacgcgt
//

Lenti plKO_EF1A_ORF_HLA_C*03 04

LOCUS plKO_EF1A_ORF_HLA_C*03 04                8742 bp DNA     circular SYN 24-OCT-2022
DEFINITION  synthetic circular DNA
ACCESSION   .
VERSION     .
KEYWORDS    .
SOURCE      synthetic DNA construct
  ORGANISM  synthetic DNA construct
REFERENCE   1  (bases 1 to 8742)
  AUTHORS   .
  TITLE     Direct Submission
  JOURNAL   Exported Oct 25, 2022 from SnapGene 6.0.5
            https://www.snapgene.com
COMMENT     U515NEI230-192 DNA889107
FEATURES             Location/Qualifiers
     source          1..8742
                     /mol_type="other DNA"
                     /organism="synthetic DNA construct"
     misc_feature    10..598
                     /label=WPRE
     CDS             complement(481..492)
                     /label=Factor Xa site
     LTR             670..903
                     /label=3' LTR (Delta-U3)
     polyA_signal    981..1102
                     /label=SV40 poly(A) signal
     rep_origin      1142..1277
                     /label=SV40 ori
     promoter        complement(1298..1316)
                     /label=T7 promoter
     primer_bind     complement(1326..1342)
                     /label=M13 fwd
     rep_origin      1484..1939
                     /label=f1 ori
     promoter        1965..2069
                     /label=AmpR promoter
     CDS             2070..2138
                     /label=AmpR
     CDS             2139..2930
                     /label=AmpR
     rep_origin      3101..3689
                     /label=ori
     promoter        4019..4036
                     /label=lac promoter
     protein_bind    4051..4067
                     /label=lac operator
     primer_bind     4075..4091
                     /label=M13 rev
     promoter        4112..4130
                     /label=T3 promoter
     promoter        4158..4384
                     /label=RSV promoter
     LTR             4385..4565
                     /label=5' LTR (truncated)
     misc_feature    4612..4737
                     /label=HIV-1 Psi
     misc_feature    5230..5463
                     /label=RRE
     misc_feature    5959..6076
                     /label=cPPT/CTS
     promoter        6236..7417
                     /label=EF-1-alpha promoter
     intron          6466..7408
                     /label=EF-1-alpha intron A
     CDS             7443..8543
                     /codon_start=1
                     /label=HLA-C*03:04
                     /translation="MRVMAPRTLILLLSGALALTETWAGSHSMRYFYTAVSRPGRGEPH
                     FIAVGYVDDTQFVRFDSDAASPRGEPRAPWVEQEGPEYWDRETQKYKRQAQTDRVSLRN
                     LRGYYNQSEAGSHIIQRMYGCDVGPDGRLLRGYDQYAYDGKDYIALNEDLRSWTAADTA
                     AQITQRKWEAAREAEQLRAYLEGLCVEWLRRYLKNGKETLQRAEHPKTHVTHHPVSDHE
                     ATLRCWALGFYPAEITLTWQWDGEDQTQDTELVETRPAGDGTFQKWAAVVVPSGEEQRY
                     TCHVQHEGLPEPLTLRWEPSSQPTIPIVGIVAGLAVLAVLAVLGAVVAVVMCRRKSSGG
                     KGGSCSQAASSNSAQGSDESLIACKA"
     misc_RNA        complement(8591..8666)
                     /label=gRNA scaffold/Barcode
ORIGIN
        1 taagtcgaca atcaacctct ggattacaaa atttgtgaaa gattgactgg tattcttaac
       61 tatgttgctc cttttacgct atgtggatac gctgctttaa tgcctttgta tcatgctatt
      121 gcttcccgta tggctttcat tttctcctcc ttgtataaat cctggttgct gtctctttat
      181 gaggagttgt ggcccgttgt caggcaacgt ggcgtggtgt gcactgtgtt tgctgacgca
      241 acccccactg gttggggcat tgccaccacc tgtcagctcc tttccgggac tttcgctttc
      301 cccctcccta ttgccacggc ggaactcatc gccgcctgcc ttgcccgctg ctggacaggg
      361 gctcggctgt tgggcactga caattccgtg gtgttgtcgg ggaaatcatc gtcctttcct
      421 tggctgctcg cctgtgttgc cacctggatt ctgcgcggga cgtccttctg ctacgtccct
      481 tcggccctca atccagcgga ccttccttcc cgcggcctgc tgccggctct gcggcctctt
      541 ccgcgtcttc gccttcgccc tcagacgagt cggatctccc tttgggccgc ctccccgcgt
      601 cgactttaag accaatgact tacaaggcag ctgtagatct tagccacttt ttaaaagaaa
      661 aggggggact ggaagggcta attcactccc aacgaagaca agatctgctt tttgcttgta
      721 ctgggtctct ctggttagac cagatctgag cctgggagct ctctggctaa ctagggaacc
      781 cactgcttaa gcctcaataa agcttgcctt gagtgcttca agtagtgtgt gcccgtctgt
      841 tgtgtgactc tggtaactag agatccctca gaccctttta gtcagtgtgg aaaatctcta
      901 gcagtacgta tagtagttca tgtcatctta ttattcagta tttataactt gcaaagaaat
      961 gaatatcaga gagtgagagg aacttgttta ttgcagctta taatggttac aaataaagca
     1021 atagcatcac aaatttcaca aataaagcat ttttttcact gcattctagt tgtggtttgt
     1081 ccaaactcat caatgtatct tatcatgtct ggctctagct atcccgcccc taactccgcc
     1141 catcccgccc ctaactccgc ccagttccgc ccattctccg ccccatggct gactaatttt
     1201 ttttatttat gcagaggccg aggccgcctc ggcctctgag ctattccaga agtagtgagg
     1261 aggctttttt ggaggcctag ggacgtaccc aattcgccct atagtgagtc gtattacgcg
     1321 cgctcactgg ccgtcgtttt acaacgtcgt gactgggaaa accctggcgt tacccaactt
     1381 aatcgccttg cagcacatcc ccctttcgcc agctggcgta atagcgaaga ggcccgcacc
     1441 gatcgccctt cccaacagtt gcgcagcctg aatggcgaat gggacgcgcc ctgtagcggc
     1501 gcattaagcg cggcgggtgt ggtggttacg cgcagcgtga ccgctacact tgccagcgcc
     1561 ctagcgcccg ctcctttcgc tttcttccct tcctttctcg ccacgttcgc cggctttccc
     1621 cgtcaagctc taaatcgggg gctcccttta gggttccgat ttagtgcttt acggcacctc
     1681 gaccccaaaa aacttgatta gggtgatggt tcacgtagtg ggccatcgcc ctgatagacg
     1741 gtttttcgcc ctttgacgtt ggagtccacg ttctttaata gtggactctt gttccaaact
     1801 ggaacaacac tcaaccctat ctcggtctat tcttttgatt tataagggat tttgccgatt
     1861 tcggcctatt ggttaaaaaa tgagctgatt taacaaaaat ttaacgcgaa ttttaacaaa
     1921 atattaacgc ttacaattta ggtggcactt ttcggggaaa tgtgcgcgga acccctattt
     1981 gtttattttt ctaaatacat tcaaatatgt atccgctcat gagacaataa ccctgataaa
     2041 tgcttcaata atattgaaaa aggaagagta tgagtattca acatttccgt gtcgccctta
     2101 ttcccttttt tgcggcattt tgccttcctg tttttgctca cccagaaacg ctggtgaaag
     2161 taaaagatgc tgaagatcag ttgggtgcac gagtgggtta catcgaactg gatctcaaca
     2221 gcggtaagat ccttgagagt tttcgccccg aagaacgttt tccaatgatg agcactttta
     2281 aagttctgct atgtggcgcg gtattatccc gtattgacgc cgggcaagag caactcggtc
     2341 gccgcataca ctattctcag aatgacttgg ttgagtactc accagtcaca gaaaagcatc
     2401 ttacggatgg catgacagta agagaattat gcagtgctgc cataaccatg agtgataaca
     2461 ctgcggccaa cttacttctg acaacgatcg gaggaccgaa ggagctaacc gcttttttgc
     2521 acaacatggg ggatcatgta actcgccttg atcgttggga accggagctg aatgaagcca
     2581 taccaaacga cgagcgtgac accacgatgc ctgtagcaat ggcaacaacg ttgcgcaaac
     2641 tattaactgg cgaactactt actctagctt cccggcaaca attaatagac tggatggagg
     2701 cggataaagt tgcaggacca cttctgcgct cggcccttcc ggctggctgg tttattgctg
     2761 ataaatctgg agccggtgag cgtgggtctc gcggtatcat tgcagcactg gggccagatg
     2821 gtaagccctc ccgtatcgta gttatctaca cgacggggag tcaggcaact atggatgaac
     2881 gaaatagaca gatcgctgag ataggtgcct cactgattaa gcattggtaa ctgtcagacc
     2941 aagtttactc atatatactt tagattgatt taaaacttca tttttaattt aaaaggatct
     3001 aggtgaagat cctttttgat aatctcatga ccaaaatccc ttaacgtgag ttttcgttcc
     3061 actgagcgtc agaccccgta gaaaagatca aaggatcttc ttgagatcct ttttttctgc
     3121 gcgtaatctg ctgcttgcaa acaaaaaaac caccgctacc agcggtggtt tgtttgccgg
     3181 atcaagagct accaactctt tttccgaagg taactggctt cagcagagcg cagataccaa
     3241 atactgttct tctagtgtag ccgtagttag gccaccactt caagaactct gtagcaccgc
     3301 ctacatacct cgctctgcta atcctgttac cagtggctgc tgccagtggc gataagtcgt
     3361 gtcttaccgg gttggactca agacgatagt taccggataa ggcgcagcgg tcgggctgaa
     3421 cggggggttc gtgcacacag cccagcttgg agcgaacgac ctacaccgaa ctgagatacc
     3481 tacagcgtga gctatgagaa agcgccacgc ttcccgaagg gagaaaggcg gacaggtatc
     3541 cggtaagcgg cagggtcgga acaggagagc gcacgaggga gcttccaggg ggaaacgcct
     3601 ggtatcttta tagtcctgtc gggtttcgcc acctctgact tgagcgtcga tttttgtgat
     3661 gctcgtcagg ggggcggagc ctatggaaaa acgccagcaa cgcggccttt ttacggttcc
     3721 tggccttttg ctggcctttt gctcacatgt tctttcctgc gttatcccct gattctgtgg
     3781 ataaccgtat taccgccttt gagtgagctg ataccgctcg ccgcagccga acgaccgagc
     3841 gcagcgagtc agtgagcgag gaagcggaag agcgcccaat acgcaaaccg cctctccccg
     3901 cgcgttggcc gattcattaa tgcagctggc acgacaggtt tcccgactgg aaagcgggca
     3961 gtgagcgcaa cgcaattaat gtgagttagc tcactcatta ggcaccccag gctttacact
     4021 ttatgcttcc ggctcgtatg ttgtgtggaa ttgtgagcgg ataacaattt cacacaggaa
     4081 acagctatga ccatgattac gccaagcgcg caattaaccc tcactaaagg gaacaaaagc
     4141 tggagctgca agcttaatgt agtcttatgc aatactcttg tagtcttgca acatggtaac
     4201 gatgagttag caacatgcct tacaaggaga gaaaaagcac cgtgcatgcc gattggtgga
     4261 agtaaggtgg tacgatcgtg ccttattagg aaggcaacag acgggtctga catggattgg
     4321 acgaaccact gaattgccgc attgcagaga tattgtattt aagtgcctag ctcgatacat
     4381 aaacgggtct ctctggttag accagatctg agcctgggag ctctctggct aactagggaa
     4441 cccactgctt aagcctcaat aaagcttgcc ttgagtgctt caagtagtgt gtgcccgtct
     4501 gttgtgtgac tctggtaact agagatccct cagacccttt tagtcagtgt ggaaaatctc
     4561 tagcagtggc gcccgaacag ggacttgaaa gcgaaaggga aaccagagga gctctctcga
     4621 cgcaggactc ggcttgctga agcgcgcacg gcaagaggcg aggggcggcg actggtgagt
     4681 acgccaaaaa ttttgactag cggaggctag aaggagagag atgggtgcga gagcgtcagt
     4741 attaagcggg ggagaattag atcgcgatgg gaaaaaattc ggttaaggcc agggggaaag
     4801 aaaaaatata aattaaaaca tatagtatgg gcaagcaggg agctagaacg attcgcagtt
     4861 aatcctggcc tgttagaaac atcagaaggc tgtagacaaa tactgggaca gctacaacca
     4921 tcccttcaga caggatcaga agaacttaga tcattatata atacagtagc aaccctctat
     4981 tgtgtgcatc aaaggataga gataaaagac accaaggaag ctttagacaa gatagaggaa
     5041 gagcaaaaca aaagtaagac caccgcacag caagcggccg ctgatcttca gacctggagg
     5101 aggagatatg agggacaatt ggagaagtga attatataaa tataaagtag taaaaattga
     5161 accattagga gtagcaccca ccaaggcaaa gagaagagtg gtgcagagag aaaaaagagc
     5221 agtgggaata ggagctttgt tccttgggtt cttgggagca gcaggaagca ctatgggcgc
     5281 agcgtcaatg acgctgacgg tacaggccag acaattattg tctggtatag tgcagcagca
     5341 gaacaatttg ctgagggcta ttgaggcgca acagcatctg ttgcaactca cagtctgggg
     5401 catcaagcag ctccaggcaa gaatcctggc tgtggaaaga tacctaaagg atcaacagct
     5461 cctggggatt tggggttgct ctggaaaact catttgcacc actgctgtgc cttggaatgc
     5521 tagttggagt aataaatctc tggaacagat ttggaatcac acgacctgga tggagtggga
     5581 cagagaaatt aacaattaca caagcttaat acactcctta attgaagaat cgcaaaacca
     5641 gcaagaaaag aatgaacaag aattattgga attagataaa tgggcaagtt tgtggaattg
     5701 gtttaacata acaaattggc tgtggtatat aaaattattc ataatgatag taggaggctt
     5761 ggtaggttta agaatagttt ttgctgtact ttctatagtg aatagagtta ggcagggata
     5821 ttcaccatta tcgtttcaga cccacctccc aaccccgagg ggacccgaca ggcccgaagg
     5881 aatagaagaa gaaggtggag agagagacag agacagatcc attcgattag tgaacggatc
     5941 tcgacggtat cggttaactt ttaaaagaaa aggggggatt ggggggtaca gtgcagggga
     6001 aagaatagta gacataatag caacagacat acaaactaaa gaattacaaa aacaaattac
     6061 aaaaattcaa aattttatcg atgagtaatt catacaaaag gactcgcccc tgccttgggg
     6121 aatcccaggg accgtcgtta aactcccact aacgtagaac ccagagatcg ctgcgttccc
     6181 gccccctcac ccgcccgctc tcgtcatcac tgaggtggag aagagcatgc gtgaggctcc
     6241 ggtgcccgtc agtgggcaga gcgcacatcg cccacagtcc ccgagaagtt ggggggaggg
     6301 gtcggcaatt gaaccggtgc ctagagaagg tggcgcgggg taaactggga aagtgatgtc
     6361 gtgtactggc tccgcctttt tcccgagggt gggggagaac cgtatataag tgcagtagtc
     6421 gccgtgaacg ttctttttcg caacgggttt gccgccagaa cacaggtaag tgccgtgtgt
     6481 ggttcccgcg ggcctggcct ctttacgggt tatggccctt gcgtgccttg aattacttcc
     6541 acgcccctgg ctgcagtacg tgattcttga tcccgagctt cgggttggaa gtgggtggga
     6601 gagttcgagg ccttgcgctt aaggagcccc ttcgcctcgt gcttgagttg aggcctggct
     6661 tgggcgctgg ggccgccgcg tgcgaatctg gtggcacctt cgcgcctgtc tcgctgcttt
     6721 cgataagtct ctagccattt aaaatttttg atgacctgct gcgacgcttt ttttctggca
     6781 agatagtctt gtaaatgcgg gccaagatct gcacactggt atttcggttt ttggggccgc
     6841 gggcggcgac ggggcccgtg cgtcccagcg cacatgttcg gcgaggcggg gcctgcgagc
     6901 gcggccaccg agaatcggac gggggtagtc tcaagctggc cggcctgctc tggtgcctgg
     6961 cctcgcgccg ccgtgtatcg ccccgccctg ggcggcaagg ctggcccggt cggcaccagt
     7021 tgcgtgagcg gaaagatggc cgcttcccgg ccctgctgca gggagctcaa aatggaggac
     7081 gcggcgctcg ggagagcggg cgggtgagtc acccacacaa aggaaaaggg cctttccgtc
     7141 ctcagccgtc gcttcatgtg actccacgga gtaccgggcg ccgtccaggc acctcgatta
     7201 gttctcaagc ttttggagta cgtcgtcttt aggttggggg gaggggtttt atgcgatgga
     7261 gtttccccac actgagtggg tggagactga agttaggcca gcttggcact tgatgtaatt
     7321 ctccttggaa tttgcccttt ttgagtttgg atcttggttc attctcaagc ctcagacagt
     7381 ggttcaaagt ttttttcttc catttcaggt gtcgtgaccc tagcgctacc tctagagcca
     7441 ccatgagagt gatggcccct agaaccctga tcctgctgct gtccggcgcc ctggccctga
     7501 ccgagacatg ggccggcagc cactccatgc gctacttcta taccgccgtg tctcggcccg
     7561 gaagaggaga gcctcacttt atcgccgtgg gctacgtgga cgatacacag ttcgtgcgct
     7621 ttgacagcga tgcagcatcc cctaggggag agccaagggc accctgggtg gagcaggagg
     7681 gcccagagta ctgggaccgg gagacacaga agtataagag gcaggcccag acagatcgcg
     7741 tgagcctgag gaacctgcgc ggctactata atcagtctga ggccggcagc cacatcatcc
     7801 agagaatgta cggatgcgac gtgggaccag atggccggct gctgagaggc tacgaccagt
     7861 acgcctatga cggcaaggat tatatcgccc tgaacgagga cctgaggtcc tggaccgcag
     7921 cagatacagc agcacagatc acccagagga agtgggaggc agcaagagag gcagagcagc
     7981 tgagggcata cctggagggc ctgtgcgtgg agtggctgcg gagatatctg aagaatggca
     8041 aggagacact gcagagagcc gagcacccaa agacccacgt gacacaccac cccgtgtctg
     8101 atcacgaggc cacactgagg tgttgggccc tgggcttcta cccagccgag atcaccctga
     8161 catggcagtg ggacggcgag gatcagaccc aggacacaga gctggtggag acacggcccg
     8221 caggcgatgg cacatttcag aagtgggcag cagtggtggt gccttccgga gaggagcagc
     8281 ggtacacatg ccacgtgcag cacgagggcc tgccagagcc tctgaccctg agatgggagc
     8341 caagctccca gcctacaatc ccaatcgtgg gaatcgtggc aggcctggcc gtgctggccg
     8401 tgctggccgt gctgggtgca gtggtggcag tggtcatgtg caggaggaag tctagcggag
     8461 gcaagggagg atcttgtagc caggcagcat cctctaacag cgcccaggga tccgacgagt
     8521 ctctgatcgc ctgtaaggcc tgaggcgcgc ctcccagagc caccgttaca cagtgtgttc
     8581 ccgtccgcac gcaccgactc ggtgccactt tttcaagttg ataacggact agccttattt
     8641 taacttgcta tttctagctc taaaactttt tttgactaga tcttgagaca aatggcagta
     8701 ttcatccaca aggtacgcta aacgcctgca accaggacgc gt
//

Lenti plKO_EF1A_ORF_HLA_C*04 01

LOCUS plKO_EF1A_ORF_HLA_C*04 01                8742 bp DNA     circular SYN 24-OCT-2022
DEFINITION  synthetic circular DNA
ACCESSION   .
VERSION     .
KEYWORDS    .
SOURCE      synthetic DNA construct
  ORGANISM  synthetic DNA construct
REFERENCE   1  (bases 1 to 8742)
  AUTHORS   .
  TITLE     Direct Submission
  JOURNAL   Exported Oct 25, 2022 from SnapGene 6.0.5
            https://www.snapgene.com
COMMENT     U515NEI230-194 DNA889108
FEATURES             Location/Qualifiers
     source          1..8742
                     /mol_type="other DNA"
                     /organism="synthetic DNA construct"
     misc_feature    10..598
                     /label=WPRE
     CDS             complement(481..492)
                     /label=Factor Xa site
     LTR             670..903
                     /label=3' LTR (Delta-U3)
     polyA_signal    981..1102
                     /label=SV40 poly(A) signal
     rep_origin      1142..1277
                     /label=SV40 ori
     promoter        complement(1298..1316)
                     /label=T7 promoter
     primer_bind     complement(1326..1342)
                     /label=M13 fwd
     rep_origin      1484..1939
                     /label=f1 ori
     promoter        1965..2069
                     /label=AmpR promoter
     CDS             2070..2138
                     /label=AmpR
     CDS             2139..2930
                     /label=AmpR
     rep_origin      3101..3689
                     /label=ori
     promoter        4019..4036
                     /label=lac promoter
     protein_bind    4051..4067
                     /label=lac operator
     primer_bind     4075..4091
                     /label=M13 rev
     promoter        4112..4130
                     /label=T3 promoter
     promoter        4158..4384
                     /label=RSV promoter
     LTR             4385..4565
                     /label=5' LTR (truncated)
     misc_feature    4612..4737
                     /label=HIV-1 Psi
     misc_feature    5230..5463
                     /label=RRE
     misc_feature    5959..6076
                     /label=cPPT/CTS
     promoter        6236..7417
                     /label=EF-1-alpha promoter
     intron          6466..7408
                     /label=EF-1-alpha intron A
     CDS             7443..8543
                     /codon_start=1
                     /label=HLA-C*04:01
                     /translation="MRVMAPRTLILLLSGALALTETWAGSHSMRYFSTSVSWPGRGEPR
                     FIAVGYVDDTQFVRFDSDAASPRGEPREPWVEQEGPEYWDRETQKYKRQAQADRVNLRK
                     LRGYYNQSEDGSHTLQRMFGCDLGPDGRLLRGYNQFAYDGKDYIALNEDLRSWTAADTA
                     AQITQRKWEAAREAEQRRAYLEGTCVEWLRRYLENGKETLQRAEHPKTHVTHHPVSDHE
                     ATLRCWALGFYPAEITLTWQWDGEDQTQDTELVETRPAGDGTFQKWAAVVVPSGEEQRY
                     TCHVQHEGLPEPLTLRWKPSSQPTIPIVGIVAGLAVLAVLAVLGAMVAVVMCRRKSSGG
                     KGGSCSQAASSNSAQGSDESLIACKA"
     misc_RNA        complement(8591..8666)
                     /label=gRNA scaffold/Barcode
ORIGIN
        1 taagtcgaca atcaacctct ggattacaaa atttgtgaaa gattgactgg tattcttaac
       61 tatgttgctc cttttacgct atgtggatac gctgctttaa tgcctttgta tcatgctatt
      121 gcttcccgta tggctttcat tttctcctcc ttgtataaat cctggttgct gtctctttat
      181 gaggagttgt ggcccgttgt caggcaacgt ggcgtggtgt gcactgtgtt tgctgacgca
      241 acccccactg gttggggcat tgccaccacc tgtcagctcc tttccgggac tttcgctttc
      301 cccctcccta ttgccacggc ggaactcatc gccgcctgcc ttgcccgctg ctggacaggg
      361 gctcggctgt tgggcactga caattccgtg gtgttgtcgg ggaaatcatc gtcctttcct
      421 tggctgctcg cctgtgttgc cacctggatt ctgcgcggga cgtccttctg ctacgtccct
      481 tcggccctca atccagcgga ccttccttcc cgcggcctgc tgccggctct gcggcctctt
      541 ccgcgtcttc gccttcgccc tcagacgagt cggatctccc tttgggccgc ctccccgcgt
      601 cgactttaag accaatgact tacaaggcag ctgtagatct tagccacttt ttaaaagaaa
      661 aggggggact ggaagggcta attcactccc aacgaagaca agatctgctt tttgcttgta
      721 ctgggtctct ctggttagac cagatctgag cctgggagct ctctggctaa ctagggaacc
      781 cactgcttaa gcctcaataa agcttgcctt gagtgcttca agtagtgtgt gcccgtctgt
      841 tgtgtgactc tggtaactag agatccctca gaccctttta gtcagtgtgg aaaatctcta
      901 gcagtacgta tagtagttca tgtcatctta ttattcagta tttataactt gcaaagaaat
      961 gaatatcaga gagtgagagg aacttgttta ttgcagctta taatggttac aaataaagca
     1021 atagcatcac aaatttcaca aataaagcat ttttttcact gcattctagt tgtggtttgt
     1081 ccaaactcat caatgtatct tatcatgtct ggctctagct atcccgcccc taactccgcc
     1141 catcccgccc ctaactccgc ccagttccgc ccattctccg ccccatggct gactaatttt
     1201 ttttatttat gcagaggccg aggccgcctc ggcctctgag ctattccaga agtagtgagg
     1261 aggctttttt ggaggcctag ggacgtaccc aattcgccct atagtgagtc gtattacgcg
     1321 cgctcactgg ccgtcgtttt acaacgtcgt gactgggaaa accctggcgt tacccaactt
     1381 aatcgccttg cagcacatcc ccctttcgcc agctggcgta atagcgaaga ggcccgcacc
     1441 gatcgccctt cccaacagtt gcgcagcctg aatggcgaat gggacgcgcc ctgtagcggc
     1501 gcattaagcg cggcgggtgt ggtggttacg cgcagcgtga ccgctacact tgccagcgcc
     1561 ctagcgcccg ctcctttcgc tttcttccct tcctttctcg ccacgttcgc cggctttccc
     1621 cgtcaagctc taaatcgggg gctcccttta gggttccgat ttagtgcttt acggcacctc
     1681 gaccccaaaa aacttgatta gggtgatggt tcacgtagtg ggccatcgcc ctgatagacg
     1741 gtttttcgcc ctttgacgtt ggagtccacg ttctttaata gtggactctt gttccaaact
     1801 ggaacaacac tcaaccctat ctcggtctat tcttttgatt tataagggat tttgccgatt
     1861 tcggcctatt ggttaaaaaa tgagctgatt taacaaaaat ttaacgcgaa ttttaacaaa
     1921 atattaacgc ttacaattta ggtggcactt ttcggggaaa tgtgcgcgga acccctattt
     1981 gtttattttt ctaaatacat tcaaatatgt atccgctcat gagacaataa ccctgataaa
     2041 tgcttcaata atattgaaaa aggaagagta tgagtattca acatttccgt gtcgccctta
     2101 ttcccttttt tgcggcattt tgccttcctg tttttgctca cccagaaacg ctggtgaaag
     2161 taaaagatgc tgaagatcag ttgggtgcac gagtgggtta catcgaactg gatctcaaca
     2221 gcggtaagat ccttgagagt tttcgccccg aagaacgttt tccaatgatg agcactttta
     2281 aagttctgct atgtggcgcg gtattatccc gtattgacgc cgggcaagag caactcggtc
     2341 gccgcataca ctattctcag aatgacttgg ttgagtactc accagtcaca gaaaagcatc
     2401 ttacggatgg catgacagta agagaattat gcagtgctgc cataaccatg agtgataaca
     2461 ctgcggccaa cttacttctg acaacgatcg gaggaccgaa ggagctaacc gcttttttgc
     2521 acaacatggg ggatcatgta actcgccttg atcgttggga accggagctg aatgaagcca
     2581 taccaaacga cgagcgtgac accacgatgc ctgtagcaat ggcaacaacg ttgcgcaaac
     2641 tattaactgg cgaactactt actctagctt cccggcaaca attaatagac tggatggagg
     2701 cggataaagt tgcaggacca cttctgcgct cggcccttcc ggctggctgg tttattgctg
     2761 ataaatctgg agccggtgag cgtgggtctc gcggtatcat tgcagcactg gggccagatg
     2821 gtaagccctc ccgtatcgta gttatctaca cgacggggag tcaggcaact atggatgaac
     2881 gaaatagaca gatcgctgag ataggtgcct cactgattaa gcattggtaa ctgtcagacc
     2941 aagtttactc atatatactt tagattgatt taaaacttca tttttaattt aaaaggatct
     3001 aggtgaagat cctttttgat aatctcatga ccaaaatccc ttaacgtgag ttttcgttcc
     3061 actgagcgtc agaccccgta gaaaagatca aaggatcttc ttgagatcct ttttttctgc
     3121 gcgtaatctg ctgcttgcaa acaaaaaaac caccgctacc agcggtggtt tgtttgccgg
     3181 atcaagagct accaactctt tttccgaagg taactggctt cagcagagcg cagataccaa
     3241 atactgttct tctagtgtag ccgtagttag gccaccactt caagaactct gtagcaccgc
     3301 ctacatacct cgctctgcta atcctgttac cagtggctgc tgccagtggc gataagtcgt
     3361 gtcttaccgg gttggactca agacgatagt taccggataa ggcgcagcgg tcgggctgaa
     3421 cggggggttc gtgcacacag cccagcttgg agcgaacgac ctacaccgaa ctgagatacc
     3481 tacagcgtga gctatgagaa agcgccacgc ttcccgaagg gagaaaggcg gacaggtatc
     3541 cggtaagcgg cagggtcgga acaggagagc gcacgaggga gcttccaggg ggaaacgcct
     3601 ggtatcttta tagtcctgtc gggtttcgcc acctctgact tgagcgtcga tttttgtgat
     3661 gctcgtcagg ggggcggagc ctatggaaaa acgccagcaa cgcggccttt ttacggttcc
     3721 tggccttttg ctggcctttt gctcacatgt tctttcctgc gttatcccct gattctgtgg
     3781 ataaccgtat taccgccttt gagtgagctg ataccgctcg ccgcagccga acgaccgagc
     3841 gcagcgagtc agtgagcgag gaagcggaag agcgcccaat acgcaaaccg cctctccccg
     3901 cgcgttggcc gattcattaa tgcagctggc acgacaggtt tcccgactgg aaagcgggca
     3961 gtgagcgcaa cgcaattaat gtgagttagc tcactcatta ggcaccccag gctttacact
     4021 ttatgcttcc ggctcgtatg ttgtgtggaa ttgtgagcgg ataacaattt cacacaggaa
     4081 acagctatga ccatgattac gccaagcgcg caattaaccc tcactaaagg gaacaaaagc
     4141 tggagctgca agcttaatgt agtcttatgc aatactcttg tagtcttgca acatggtaac
     4201 gatgagttag caacatgcct tacaaggaga gaaaaagcac cgtgcatgcc gattggtgga
     4261 agtaaggtgg tacgatcgtg ccttattagg aaggcaacag acgggtctga catggattgg
     4321 acgaaccact gaattgccgc attgcagaga tattgtattt aagtgcctag ctcgatacat
     4381 aaacgggtct ctctggttag accagatctg agcctgggag ctctctggct aactagggaa
     4441 cccactgctt aagcctcaat aaagcttgcc ttgagtgctt caagtagtgt gtgcccgtct
     4501 gttgtgtgac tctggtaact agagatccct cagacccttt tagtcagtgt ggaaaatctc
     4561 tagcagtggc gcccgaacag ggacttgaaa gcgaaaggga aaccagagga gctctctcga
     4621 cgcaggactc ggcttgctga agcgcgcacg gcaagaggcg aggggcggcg actggtgagt
     4681 acgccaaaaa ttttgactag cggaggctag aaggagagag atgggtgcga gagcgtcagt
     4741 attaagcggg ggagaattag atcgcgatgg gaaaaaattc ggttaaggcc agggggaaag
     4801 aaaaaatata aattaaaaca tatagtatgg gcaagcaggg agctagaacg attcgcagtt
     4861 aatcctggcc tgttagaaac atcagaaggc tgtagacaaa tactgggaca gctacaacca
     4921 tcccttcaga caggatcaga agaacttaga tcattatata atacagtagc aaccctctat
     4981 tgtgtgcatc aaaggataga gataaaagac accaaggaag ctttagacaa gatagaggaa
     5041 gagcaaaaca aaagtaagac caccgcacag caagcggccg ctgatcttca gacctggagg
     5101 aggagatatg agggacaatt ggagaagtga attatataaa tataaagtag taaaaattga
     5161 accattagga gtagcaccca ccaaggcaaa gagaagagtg gtgcagagag aaaaaagagc
     5221 agtgggaata ggagctttgt tccttgggtt cttgggagca gcaggaagca ctatgggcgc
     5281 agcgtcaatg acgctgacgg tacaggccag acaattattg tctggtatag tgcagcagca
     5341 gaacaatttg ctgagggcta ttgaggcgca acagcatctg ttgcaactca cagtctgggg
     5401 catcaagcag ctccaggcaa gaatcctggc tgtggaaaga tacctaaagg atcaacagct
     5461 cctggggatt tggggttgct ctggaaaact catttgcacc actgctgtgc cttggaatgc
     5521 tagttggagt aataaatctc tggaacagat ttggaatcac acgacctgga tggagtggga
     5581 cagagaaatt aacaattaca caagcttaat acactcctta attgaagaat cgcaaaacca
     5641 gcaagaaaag aatgaacaag aattattgga attagataaa tgggcaagtt tgtggaattg
     5701 gtttaacata acaaattggc tgtggtatat aaaattattc ataatgatag taggaggctt
     5761 ggtaggttta agaatagttt ttgctgtact ttctatagtg aatagagtta ggcagggata
     5821 ttcaccatta tcgtttcaga cccacctccc aaccccgagg ggacccgaca ggcccgaagg
     5881 aatagaagaa gaaggtggag agagagacag agacagatcc attcgattag tgaacggatc
     5941 tcgacggtat cggttaactt ttaaaagaaa aggggggatt ggggggtaca gtgcagggga
     6001 aagaatagta gacataatag caacagacat acaaactaaa gaattacaaa aacaaattac
     6061 aaaaattcaa aattttatcg atgagtaatt catacaaaag gactcgcccc tgccttgggg
     6121 aatcccaggg accgtcgtta aactcccact aacgtagaac ccagagatcg ctgcgttccc
     6181 gccccctcac ccgcccgctc tcgtcatcac tgaggtggag aagagcatgc gtgaggctcc
     6241 ggtgcccgtc agtgggcaga gcgcacatcg cccacagtcc ccgagaagtt ggggggaggg
     6301 gtcggcaatt gaaccggtgc ctagagaagg tggcgcgggg taaactggga aagtgatgtc
     6361 gtgtactggc tccgcctttt tcccgagggt gggggagaac cgtatataag tgcagtagtc
     6421 gccgtgaacg ttctttttcg caacgggttt gccgccagaa cacaggtaag tgccgtgtgt
     6481 ggttcccgcg ggcctggcct ctttacgggt tatggccctt gcgtgccttg aattacttcc
     6541 acgcccctgg ctgcagtacg tgattcttga tcccgagctt cgggttggaa gtgggtggga
     6601 gagttcgagg ccttgcgctt aaggagcccc ttcgcctcgt gcttgagttg aggcctggct
     6661 tgggcgctgg ggccgccgcg tgcgaatctg gtggcacctt cgcgcctgtc tcgctgcttt
     6721 cgataagtct ctagccattt aaaatttttg atgacctgct gcgacgcttt ttttctggca
     6781 agatagtctt gtaaatgcgg gccaagatct gcacactggt atttcggttt ttggggccgc
     6841 gggcggcgac ggggcccgtg cgtcccagcg cacatgttcg gcgaggcggg gcctgcgagc
     6901 gcggccaccg agaatcggac gggggtagtc tcaagctggc cggcctgctc tggtgcctgg
     6961 cctcgcgccg ccgtgtatcg ccccgccctg ggcggcaagg ctggcccggt cggcaccagt
     7021 tgcgtgagcg gaaagatggc cgcttcccgg ccctgctgca gggagctcaa aatggaggac
     7081 gcggcgctcg ggagagcggg cgggtgagtc acccacacaa aggaaaaggg cctttccgtc
     7141 ctcagccgtc gcttcatgtg actccacgga gtaccgggcg ccgtccaggc acctcgatta
     7201 gttctcaagc ttttggagta cgtcgtcttt aggttggggg gaggggtttt atgcgatgga
     7261 gtttccccac actgagtggg tggagactga agttaggcca gcttggcact tgatgtaatt
     7321 ctccttggaa tttgcccttt ttgagtttgg atcttggttc attctcaagc ctcagacagt
     7381 ggttcaaagt ttttttcttc catttcaggt gtcgtgaccc tagcgctacc tctagagcca
     7441 ccatgagagt gatggcccct cgcaccctga tcctgctgct gtccggcgcc ctggccctga
     7501 ccgagacatg ggccggcagc cactccatgc ggtatttctc taccagcgtg tcctggccag
     7561 gaaggggaga gcctcgcttt atcgccgtgg gctacgtgga cgatacacag ttcgtgcggt
     7621 ttgacagcga tgcagcatcc cctaggggag agccaagaga gccatgggtg gagcaggagg
     7681 gaccagagta ttgggacaga gagacacaga agtacaagcg gcaggcccag gccgatagag
     7741 tgaacctgag gaagctgcgc ggctactata atcagtctga ggacggcagc cacacactgc
     7801 agaggatgtt cggatgcgac ctgggaccag atggccggct gctgagaggc tacaaccagt
     7861 ttgcctatga cggcaaggat tacatcgccc tgaatgagga cctgcgcagc tggaccgcag
     7921 cagatacagc agcacagatc acccagagga agtgggaggc agcaagggag gcagagcagc
     7981 ggagagccta tctggaggga acctgcgtgg agtggctgag gcgctacctg gagaacggca
     8041 aggagacact gcagagggcc gagcacccaa agacccacgt gacacaccac cccgtgtccg
     8101 atcacgaggc cacactgagg tgttgggcac tgggcttcta tccagccgag atcaccctga
     8161 catggcagtg ggacggcgag gatcagaccc aggacacaga gctggtggag acacggcccg
     8221 caggcgatgg cacatttcag aagtgggcag cagtggtggt gccttctgga gaggagcagc
     8281 ggtacacctg ccacgtgcag cacgagggcc tgccagagcc tctgaccctg agatggaagc
     8341 ccagctccca gcctacaatc ccaatcgtgg gaatcgtggc aggcctggcc gtgctggccg
     8401 tgctggccgt gctgggtgca atggtggcag tggtcatgtg ccggagaaag tctagcggag
     8461 gcaagggagg atcttgtagc caggcagcat cctctaattc cgcccagggc tccgacgagt
     8521 ctctgatcgc ctgtaaggcc tgaggcgcgc ctcccagagc caccgttaca cgcagtagtc
     8581 ccggtgcgga gcaccgactc ggtgccactt tttcaagttg ataacggact agccttattt
     8641 taacttgcta tttctagctc taaaactttt tttgactaga tcttgagaca aatggcagta
     8701 ttcatccaca aggtacgcta aacgcctgca accaggacgc gt
//

Lenti plKO_EF1A_ORF_HLA_C*05 01

LOCUS plKO_EF1A_ORF_HLA_C*05 01                8742 bp DNA     circular SYN 24-OCT-2022
DEFINITION  synthetic circular DNA
ACCESSION   .
VERSION     .
KEYWORDS    .
SOURCE      synthetic DNA construct
  ORGANISM  synthetic DNA construct
REFERENCE   1  (bases 1 to 8742)
  AUTHORS   .
  TITLE     Direct Submission
  JOURNAL   Exported Oct 25, 2022 from SnapGene 6.0.5
            https://www.snapgene.com
COMMENT     U515NEI230-196 DNA889005
FEATURES             Location/Qualifiers
     source          1..8742
                     /mol_type="other DNA"
                     /organism="synthetic DNA construct"
     misc_feature    10..598
                     /label=WPRE
     CDS             complement(481..492)
                     /label=Factor Xa site
     LTR             670..903
                     /label=3' LTR (Delta-U3)
     polyA_signal    981..1102
                     /label=SV40 poly(A) signal
     rep_origin      1142..1277
                     /label=SV40 ori
     promoter        complement(1298..1316)
                     /label=T7 promoter
     primer_bind     complement(1326..1342)
                     /label=M13 fwd
     rep_origin      1484..1939
                     /label=f1 ori
     promoter        1965..2069
                     /label=AmpR promoter
     CDS             2070..2138
                     /label=AmpR
     CDS             2139..2930
                     /label=AmpR
     rep_origin      3101..3689
                     /label=ori
     promoter        4019..4036
                     /label=lac promoter
     protein_bind    4051..4067
                     /label=lac operator
     primer_bind     4075..4091
                     /label=M13 rev
     promoter        4112..4130
                     /label=T3 promoter
     promoter        4158..4384
                     /label=RSV promoter
     LTR             4385..4565
                     /label=5' LTR (truncated)
     misc_feature    4612..4737
                     /label=HIV-1 Psi
     misc_feature    5230..5463
                     /label=RRE
     misc_feature    5959..6076
                     /label=cPPT/CTS
     promoter        6236..7417
                     /label=EF-1-alpha promoter
     intron          6466..7408
                     /label=EF-1-alpha intron A
     CDS             7443..8543
                     /codon_start=1
                     /label=HLA-C*05:01
                     /translation="MRVMAPRTLILLLSGALALTETWACSHSMRYFYTAVSRPGRGEPR
                     FIAVGYVDDTQFVQFDSDAASPRGEPRAPWVEQEGPEYWDRETQKYKRQAQTDRVNLRK
                     LRGYYNQSEAGSHTLQRMYGCDLGPDGRLLRGYNQFAYDGKDYIALNEDLRSWTAADKA
                     AQITQRKWEAAREAEQRRAYLEGTCVEWLRRYLENGKKTLQRAEHPKTHVTHHPVSDHE
                     ATLRCWALGFYPAEITLTWQRDGEDQTQDTELVETRPAGDGTFQKWAAVVVPSGEEQRY
                     TCHVQHEGLPEPLTLRWGPSSQPTIPIVGIVAGLAVLAVLAVLGAVMAVVMCRRKSSGG
                     KGGSCSQAASSNSAQGSDESLIACKA"
     misc_RNA        complement(8591..8666)
                     /label=gRNA scaffold/Barcode
ORIGIN
        1 taagtcgaca atcaacctct ggattacaaa atttgtgaaa gattgactgg tattcttaac
       61 tatgttgctc cttttacgct atgtggatac gctgctttaa tgcctttgta tcatgctatt
      121 gcttcccgta tggctttcat tttctcctcc ttgtataaat cctggttgct gtctctttat
      181 gaggagttgt ggcccgttgt caggcaacgt ggcgtggtgt gcactgtgtt tgctgacgca
      241 acccccactg gttggggcat tgccaccacc tgtcagctcc tttccgggac tttcgctttc
      301 cccctcccta ttgccacggc ggaactcatc gccgcctgcc ttgcccgctg ctggacaggg
      361 gctcggctgt tgggcactga caattccgtg gtgttgtcgg ggaaatcatc gtcctttcct
      421 tggctgctcg cctgtgttgc cacctggatt ctgcgcggga cgtccttctg ctacgtccct
      481 tcggccctca atccagcgga ccttccttcc cgcggcctgc tgccggctct gcggcctctt
      541 ccgcgtcttc gccttcgccc tcagacgagt cggatctccc tttgggccgc ctccccgcgt
      601 cgactttaag accaatgact tacaaggcag ctgtagatct tagccacttt ttaaaagaaa
      661 aggggggact ggaagggcta attcactccc aacgaagaca agatctgctt tttgcttgta
      721 ctgggtctct ctggttagac cagatctgag cctgggagct ctctggctaa ctagggaacc
      781 cactgcttaa gcctcaataa agcttgcctt gagtgcttca agtagtgtgt gcccgtctgt
      841 tgtgtgactc tggtaactag agatccctca gaccctttta gtcagtgtgg aaaatctcta
      901 gcagtacgta tagtagttca tgtcatctta ttattcagta tttataactt gcaaagaaat
      961 gaatatcaga gagtgagagg aacttgttta ttgcagctta taatggttac aaataaagca
     1021 atagcatcac aaatttcaca aataaagcat ttttttcact gcattctagt tgtggtttgt
     1081 ccaaactcat caatgtatct tatcatgtct ggctctagct atcccgcccc taactccgcc
     1141 catcccgccc ctaactccgc ccagttccgc ccattctccg ccccatggct gactaatttt
     1201 ttttatttat gcagaggccg aggccgcctc ggcctctgag ctattccaga agtagtgagg
     1261 aggctttttt ggaggcctag ggacgtaccc aattcgccct atagtgagtc gtattacgcg
     1321 cgctcactgg ccgtcgtttt acaacgtcgt gactgggaaa accctggcgt tacccaactt
     1381 aatcgccttg cagcacatcc ccctttcgcc agctggcgta atagcgaaga ggcccgcacc
     1441 gatcgccctt cccaacagtt gcgcagcctg aatggcgaat gggacgcgcc ctgtagcggc
     1501 gcattaagcg cggcgggtgt ggtggttacg cgcagcgtga ccgctacact tgccagcgcc
     1561 ctagcgcccg ctcctttcgc tttcttccct tcctttctcg ccacgttcgc cggctttccc
     1621 cgtcaagctc taaatcgggg gctcccttta gggttccgat ttagtgcttt acggcacctc
     1681 gaccccaaaa aacttgatta gggtgatggt tcacgtagtg ggccatcgcc ctgatagacg
     1741 gtttttcgcc ctttgacgtt ggagtccacg ttctttaata gtggactctt gttccaaact
     1801 ggaacaacac tcaaccctat ctcggtctat tcttttgatt tataagggat tttgccgatt
     1861 tcggcctatt ggttaaaaaa tgagctgatt taacaaaaat ttaacgcgaa ttttaacaaa
     1921 atattaacgc ttacaattta ggtggcactt ttcggggaaa tgtgcgcgga acccctattt
     1981 gtttattttt ctaaatacat tcaaatatgt atccgctcat gagacaataa ccctgataaa
     2041 tgcttcaata atattgaaaa aggaagagta tgagtattca acatttccgt gtcgccctta
     2101 ttcccttttt tgcggcattt tgccttcctg tttttgctca cccagaaacg ctggtgaaag
     2161 taaaagatgc tgaagatcag ttgggtgcac gagtgggtta catcgaactg gatctcaaca
     2221 gcggtaagat ccttgagagt tttcgccccg aagaacgttt tccaatgatg agcactttta
     2281 aagttctgct atgtggcgcg gtattatccc gtattgacgc cgggcaagag caactcggtc
     2341 gccgcataca ctattctcag aatgacttgg ttgagtactc accagtcaca gaaaagcatc
     2401 ttacggatgg catgacagta agagaattat gcagtgctgc cataaccatg agtgataaca
     2461 ctgcggccaa cttacttctg acaacgatcg gaggaccgaa ggagctaacc gcttttttgc
     2521 acaacatggg ggatcatgta actcgccttg atcgttggga accggagctg aatgaagcca
     2581 taccaaacga cgagcgtgac accacgatgc ctgtagcaat ggcaacaacg ttgcgcaaac
     2641 tattaactgg cgaactactt actctagctt cccggcaaca attaatagac tggatggagg
     2701 cggataaagt tgcaggacca cttctgcgct cggcccttcc ggctggctgg tttattgctg
     2761 ataaatctgg agccggtgag cgtgggtctc gcggtatcat tgcagcactg gggccagatg
     2821 gtaagccctc ccgtatcgta gttatctaca cgacggggag tcaggcaact atggatgaac
     2881 gaaatagaca gatcgctgag ataggtgcct cactgattaa gcattggtaa ctgtcagacc
     2941 aagtttactc atatatactt tagattgatt taaaacttca tttttaattt aaaaggatct
     3001 aggtgaagat cctttttgat aatctcatga ccaaaatccc ttaacgtgag ttttcgttcc
     3061 actgagcgtc agaccccgta gaaaagatca aaggatcttc ttgagatcct ttttttctgc
     3121 gcgtaatctg ctgcttgcaa acaaaaaaac caccgctacc agcggtggtt tgtttgccgg
     3181 atcaagagct accaactctt tttccgaagg taactggctt cagcagagcg cagataccaa
     3241 atactgttct tctagtgtag ccgtagttag gccaccactt caagaactct gtagcaccgc
     3301 ctacatacct cgctctgcta atcctgttac cagtggctgc tgccagtggc gataagtcgt
     3361 gtcttaccgg gttggactca agacgatagt taccggataa ggcgcagcgg tcgggctgaa
     3421 cggggggttc gtgcacacag cccagcttgg agcgaacgac ctacaccgaa ctgagatacc
     3481 tacagcgtga gctatgagaa agcgccacgc ttcccgaagg gagaaaggcg gacaggtatc
     3541 cggtaagcgg cagggtcgga acaggagagc gcacgaggga gcttccaggg ggaaacgcct
     3601 ggtatcttta tagtcctgtc gggtttcgcc acctctgact tgagcgtcga tttttgtgat
     3661 gctcgtcagg ggggcggagc ctatggaaaa acgccagcaa cgcggccttt ttacggttcc
     3721 tggccttttg ctggcctttt gctcacatgt tctttcctgc gttatcccct gattctgtgg
     3781 ataaccgtat taccgccttt gagtgagctg ataccgctcg ccgcagccga acgaccgagc
     3841 gcagcgagtc agtgagcgag gaagcggaag agcgcccaat acgcaaaccg cctctccccg
     3901 cgcgttggcc gattcattaa tgcagctggc acgacaggtt tcccgactgg aaagcgggca
     3961 gtgagcgcaa cgcaattaat gtgagttagc tcactcatta ggcaccccag gctttacact
     4021 ttatgcttcc ggctcgtatg ttgtgtggaa ttgtgagcgg ataacaattt cacacaggaa
     4081 acagctatga ccatgattac gccaagcgcg caattaaccc tcactaaagg gaacaaaagc
     4141 tggagctgca agcttaatgt agtcttatgc aatactcttg tagtcttgca acatggtaac
     4201 gatgagttag caacatgcct tacaaggaga gaaaaagcac cgtgcatgcc gattggtgga
     4261 agtaaggtgg tacgatcgtg ccttattagg aaggcaacag acgggtctga catggattgg
     4321 acgaaccact gaattgccgc attgcagaga tattgtattt aagtgcctag ctcgatacat
     4381 aaacgggtct ctctggttag accagatctg agcctgggag ctctctggct aactagggaa
     4441 cccactgctt aagcctcaat aaagcttgcc ttgagtgctt caagtagtgt gtgcccgtct
     4501 gttgtgtgac tctggtaact agagatccct cagacccttt tagtcagtgt ggaaaatctc
     4561 tagcagtggc gcccgaacag ggacttgaaa gcgaaaggga aaccagagga gctctctcga
     4621 cgcaggactc ggcttgctga agcgcgcacg gcaagaggcg aggggcggcg actggtgagt
     4681 acgccaaaaa ttttgactag cggaggctag aaggagagag atgggtgcga gagcgtcagt
     4741 attaagcggg ggagaattag atcgcgatgg gaaaaaattc ggttaaggcc agggggaaag
     4801 aaaaaatata aattaaaaca tatagtatgg gcaagcaggg agctagaacg attcgcagtt
     4861 aatcctggcc tgttagaaac atcagaaggc tgtagacaaa tactgggaca gctacaacca
     4921 tcccttcaga caggatcaga agaacttaga tcattatata atacagtagc aaccctctat
     4981 tgtgtgcatc aaaggataga gataaaagac accaaggaag ctttagacaa gatagaggaa
     5041 gagcaaaaca aaagtaagac caccgcacag caagcggccg ctgatcttca gacctggagg
     5101 aggagatatg agggacaatt ggagaagtga attatataaa tataaagtag taaaaattga
     5161 accattagga gtagcaccca ccaaggcaaa gagaagagtg gtgcagagag aaaaaagagc
     5221 agtgggaata ggagctttgt tccttgggtt cttgggagca gcaggaagca ctatgggcgc
     5281 agcgtcaatg acgctgacgg tacaggccag acaattattg tctggtatag tgcagcagca
     5341 gaacaatttg ctgagggcta ttgaggcgca acagcatctg ttgcaactca cagtctgggg
     5401 catcaagcag ctccaggcaa gaatcctggc tgtggaaaga tacctaaagg atcaacagct
     5461 cctggggatt tggggttgct ctggaaaact catttgcacc actgctgtgc cttggaatgc
     5521 tagttggagt aataaatctc tggaacagat ttggaatcac acgacctgga tggagtggga
     5581 cagagaaatt aacaattaca caagcttaat acactcctta attgaagaat cgcaaaacca
     5641 gcaagaaaag aatgaacaag aattattgga attagataaa tgggcaagtt tgtggaattg
     5701 gtttaacata acaaattggc tgtggtatat aaaattattc ataatgatag taggaggctt
     5761 ggtaggttta agaatagttt ttgctgtact ttctatagtg aatagagtta ggcagggata
     5821 ttcaccatta tcgtttcaga cccacctccc aaccccgagg ggacccgaca ggcccgaagg
     5881 aatagaagaa gaaggtggag agagagacag agacagatcc attcgattag tgaacggatc
     5941 tcgacggtat cggttaactt ttaaaagaaa aggggggatt ggggggtaca gtgcagggga
     6001 aagaatagta gacataatag caacagacat acaaactaaa gaattacaaa aacaaattac
     6061 aaaaattcaa aattttatcg atgagtaatt catacaaaag gactcgcccc tgccttgggg
     6121 aatcccaggg accgtcgtta aactcccact aacgtagaac ccagagatcg ctgcgttccc
     6181 gccccctcac ccgcccgctc tcgtcatcac tgaggtggag aagagcatgc gtgaggctcc
     6241 ggtgcccgtc agtgggcaga gcgcacatcg cccacagtcc ccgagaagtt ggggggaggg
     6301 gtcggcaatt gaaccggtgc ctagagaagg tggcgcgggg taaactggga aagtgatgtc
     6361 gtgtactggc tccgcctttt tcccgagggt gggggagaac cgtatataag tgcagtagtc
     6421 gccgtgaacg ttctttttcg caacgggttt gccgccagaa cacaggtaag tgccgtgtgt
     6481 ggttcccgcg ggcctggcct ctttacgggt tatggccctt gcgtgccttg aattacttcc
     6541 acgcccctgg ctgcagtacg tgattcttga tcccgagctt cgggttggaa gtgggtggga
     6601 gagttcgagg ccttgcgctt aaggagcccc ttcgcctcgt gcttgagttg aggcctggct
     6661 tgggcgctgg ggccgccgcg tgcgaatctg gtggcacctt cgcgcctgtc tcgctgcttt
     6721 cgataagtct ctagccattt aaaatttttg atgacctgct gcgacgcttt ttttctggca
     6781 agatagtctt gtaaatgcgg gccaagatct gcacactggt atttcggttt ttggggccgc
     6841 gggcggcgac ggggcccgtg cgtcccagcg cacatgttcg gcgaggcggg gcctgcgagc
     6901 gcggccaccg agaatcggac gggggtagtc tcaagctggc cggcctgctc tggtgcctgg
     6961 cctcgcgccg ccgtgtatcg ccccgccctg ggcggcaagg ctggcccggt cggcaccagt
     7021 tgcgtgagcg gaaagatggc cgcttcccgg ccctgctgca gggagctcaa aatggaggac
     7081 gcggcgctcg ggagagcggg cgggtgagtc acccacacaa aggaaaaggg cctttccgtc
     7141 ctcagccgtc gcttcatgtg actccacgga gtaccgggcg ccgtccaggc acctcgatta
     7201 gttctcaagc ttttggagta cgtcgtcttt aggttggggg gaggggtttt atgcgatgga
     7261 gtttccccac actgagtggg tggagactga agttaggcca gcttggcact tgatgtaatt
     7321 ctccttggaa tttgcccttt ttgagtttgg atcttggttc attctcaagc ctcagacagt
     7381 ggttcaaagt ttttttcttc catttcaggt gtcgtgaccc tagcgctacc tctagagcca
     7441 ccatgagagt gatggcccct cgcaccctga tcctgctgct gtccggcgcc ctggccctga
     7501 ccgagacatg ggcctgcagc cactccatgc ggtacttcta taccgccgtg tctaggcccg
     7561 gaaggggaga gcctcggttc atcgccgtgg gctatgtgga cgatacacag ttcgtgcagt
     7621 ttgacagcga tgcagcatcc cctaggggag agccaagagc accatgggtg gagcaggagg
     7681 gaccagagta ttgggaccgg gagacacaga agtacaagcg gcaggcccag acagatagag
     7741 tgaacctgcg gaagctgaga ggctactata atcagtctga ggcaggcagc cacaccctgc
     7801 agaggatgta cggatgtgac ctgggaccag atggcaggct gctgaggggc tacaaccagt
     7861 tcgcctatga cggcaaggat tacatcgccc tgaatgagga cctgagatct tggaccgccg
     7921 ccgataaggc cgcccagatc acacagagga agtgggaggc agcaagggag gcagagcagc
     7981 ggagagccta tctggaggga acctgcgtgg agtggctgag gcgctacctg gagaacggca
     8041 agaagacact gcagagggcc gagcacccaa agacccacgt gacacaccac cccgtgagcg
     8101 accacgaggc caccctgagg tgttgggcac tgggcttcta tccagccgag atcaccctga
     8161 catggcagag ggacggcgag gatcagaccc aggacacaga gctggtggag acacggcccg
     8221 caggcgatgg cacatttcag aagtgggcag cagtggtggt gccttccgga gaggagcagc
     8281 ggtacacatg ccacgtgcag cacgagggcc tgccagagcc tctgaccctg agatggggac
     8341 caagctccca gcctacaatc ccaatcgtgg gaatcgtggc aggcctggcc gtgctggccg
     8401 tgctggccgt gctgggtgcc gtgatggcag tggtcatgtg ccggagaaag tctagcggag
     8461 gcaagggagg atcttgtagc caggcagcat cctctaattc cgcccagggc tccgatgagt
     8521 ctctgatcgc ctgtaaggcc tgaggcgcgc ctcccagagc caccgttaca ccaccgggac
     8581 tactgcgtcc gcaccgactc ggtgccactt tttcaagttg ataacggact agccttattt
     8641 taacttgcta tttctagctc taaaactttt tttgactaga tcttgagaca aatggcagta
     8701 ttcatccaca aggtacgcta aacgcctgca accaggacgc gt
//

Lenti plKO_EF1A_ORF_HLA_C*06 02

LOCUS plKO_EF1A_ORF_HLA_C*06 02               8742 bp DNA     circular SYN 24-OCT-2022
DEFINITION  synthetic circular DNA
ACCESSION   .
VERSION     .
KEYWORDS    .
SOURCE      synthetic DNA construct
  ORGANISM  synthetic DNA construct
REFERENCE   1  (bases 1 to 8742)
  AUTHORS   .
  TITLE     Direct Submission
  JOURNAL   Exported Oct 25, 2022 from SnapGene 6.0.5
            https://www.snapgene.com
COMMENT     U515NEI230-198 DNA889109
FEATURES             Location/Qualifiers
     source          1..8742
                     /mol_type="other DNA"
                     /organism="synthetic DNA construct"
     misc_feature    10..598
                     /label=WPRE
     CDS             complement(481..492)
                     /label=Factor Xa site
     LTR             670..903
                     /label=3' LTR (Delta-U3)
     polyA_signal    981..1102
                     /label=SV40 poly(A) signal
     rep_origin      1142..1277
                     /label=SV40 ori
     promoter        complement(1298..1316)
                     /label=T7 promoter
     primer_bind     complement(1326..1342)
                     /label=M13 fwd
     rep_origin      1484..1939
                     /label=f1 ori
     promoter        1965..2069
                     /label=AmpR promoter
     CDS             2070..2138
                     /label=AmpR
     CDS             2139..2930
                     /label=AmpR
     rep_origin      3101..3689
                     /label=ori
     promoter        4019..4036
                     /label=lac promoter
     protein_bind    4051..4067
                     /label=lac operator
     primer_bind     4075..4091
                     /label=M13 rev
     promoter        4112..4130
                     /label=T3 promoter
     promoter        4158..4384
                     /label=RSV promoter
     LTR             4385..4565
                     /label=5' LTR (truncated)
     misc_feature    4612..4737
                     /label=HIV-1 Psi
     misc_feature    5230..5463
                     /label=RRE
     misc_feature    5959..6076
                     /label=cPPT/CTS
     promoter        6236..7417
                     /label=EF-1-alpha promoter
     intron          6466..7408
                     /label=EF-1-alpha intron A
     CDS             7443..8543
                     /codon_start=1
                     /label=HLA-C*06:02
                     /translation="MRVMAPRTLILLLSGALALTETWACSHSMRYFDTAVSRPGRGEPR
                     FISVGYVDDTQFVRFDSDAASPRGEPRAPWVEQEGPEYWDRETQKYKRQAQADRVNLRK
                     LRGYYNQSEDGSHTLQWMYGCDLGPDGRLLRGYDQSAYDGKDYIALNEDLRSWTAADTA
                     AQITQRKWEAAREAEQWRAYLEGTCVEWLRRYLENGKETLQRAEHPKTHVTHHPVSDHE
                     ATLRCWALGFYPAEITLTWQRDGEDQTQDTELVETRPAGDGTFQKWAAVVVPSGEEQRY
                     TCHVQHEGLPEPLTLRWEPSSQPTIPIVGIVAGLAVLAVLAVLGAVMAVVMCRRKSSGG
                     KGGSCSQAASSNSAQGSDESLIACKA"
     misc_RNA        complement(8591..8666)
                     /label=gRNA scaffold/Barcode
ORIGIN
        1 taagtcgaca atcaacctct ggattacaaa atttgtgaaa gattgactgg tattcttaac
       61 tatgttgctc cttttacgct atgtggatac gctgctttaa tgcctttgta tcatgctatt
      121 gcttcccgta tggctttcat tttctcctcc ttgtataaat cctggttgct gtctctttat
      181 gaggagttgt ggcccgttgt caggcaacgt ggcgtggtgt gcactgtgtt tgctgacgca
      241 acccccactg gttggggcat tgccaccacc tgtcagctcc tttccgggac tttcgctttc
      301 cccctcccta ttgccacggc ggaactcatc gccgcctgcc ttgcccgctg ctggacaggg
      361 gctcggctgt tgggcactga caattccgtg gtgttgtcgg ggaaatcatc gtcctttcct
      421 tggctgctcg cctgtgttgc cacctggatt ctgcgcggga cgtccttctg ctacgtccct
      481 tcggccctca atccagcgga ccttccttcc cgcggcctgc tgccggctct gcggcctctt
      541 ccgcgtcttc gccttcgccc tcagacgagt cggatctccc tttgggccgc ctccccgcgt
      601 cgactttaag accaatgact tacaaggcag ctgtagatct tagccacttt ttaaaagaaa
      661 aggggggact ggaagggcta attcactccc aacgaagaca agatctgctt tttgcttgta
      721 ctgggtctct ctggttagac cagatctgag cctgggagct ctctggctaa ctagggaacc
      781 cactgcttaa gcctcaataa agcttgcctt gagtgcttca agtagtgtgt gcccgtctgt
      841 tgtgtgactc tggtaactag agatccctca gaccctttta gtcagtgtgg aaaatctcta
      901 gcagtacgta tagtagttca tgtcatctta ttattcagta tttataactt gcaaagaaat
      961 gaatatcaga gagtgagagg aacttgttta ttgcagctta taatggttac aaataaagca
     1021 atagcatcac aaatttcaca aataaagcat ttttttcact gcattctagt tgtggtttgt
     1081 ccaaactcat caatgtatct tatcatgtct ggctctagct atcccgcccc taactccgcc
     1141 catcccgccc ctaactccgc ccagttccgc ccattctccg ccccatggct gactaatttt
     1201 ttttatttat gcagaggccg aggccgcctc ggcctctgag ctattccaga agtagtgagg
     1261 aggctttttt ggaggcctag ggacgtaccc aattcgccct atagtgagtc gtattacgcg
     1321 cgctcactgg ccgtcgtttt acaacgtcgt gactgggaaa accctggcgt tacccaactt
     1381 aatcgccttg cagcacatcc ccctttcgcc agctggcgta atagcgaaga ggcccgcacc
     1441 gatcgccctt cccaacagtt gcgcagcctg aatggcgaat gggacgcgcc ctgtagcggc
     1501 gcattaagcg cggcgggtgt ggtggttacg cgcagcgtga ccgctacact tgccagcgcc
     1561 ctagcgcccg ctcctttcgc tttcttccct tcctttctcg ccacgttcgc cggctttccc
     1621 cgtcaagctc taaatcgggg gctcccttta gggttccgat ttagtgcttt acggcacctc
     1681 gaccccaaaa aacttgatta gggtgatggt tcacgtagtg ggccatcgcc ctgatagacg
     1741 gtttttcgcc ctttgacgtt ggagtccacg ttctttaata gtggactctt gttccaaact
     1801 ggaacaacac tcaaccctat ctcggtctat tcttttgatt tataagggat tttgccgatt
     1861 tcggcctatt ggttaaaaaa tgagctgatt taacaaaaat ttaacgcgaa ttttaacaaa
     1921 atattaacgc ttacaattta ggtggcactt ttcggggaaa tgtgcgcgga acccctattt
     1981 gtttattttt ctaaatacat tcaaatatgt atccgctcat gagacaataa ccctgataaa
     2041 tgcttcaata atattgaaaa aggaagagta tgagtattca acatttccgt gtcgccctta
     2101 ttcccttttt tgcggcattt tgccttcctg tttttgctca cccagaaacg ctggtgaaag
     2161 taaaagatgc tgaagatcag ttgggtgcac gagtgggtta catcgaactg gatctcaaca
     2221 gcggtaagat ccttgagagt tttcgccccg aagaacgttt tccaatgatg agcactttta
     2281 aagttctgct atgtggcgcg gtattatccc gtattgacgc cgggcaagag caactcggtc
     2341 gccgcataca ctattctcag aatgacttgg ttgagtactc accagtcaca gaaaagcatc
     2401 ttacggatgg catgacagta agagaattat gcagtgctgc cataaccatg agtgataaca
     2461 ctgcggccaa cttacttctg acaacgatcg gaggaccgaa ggagctaacc gcttttttgc
     2521 acaacatggg ggatcatgta actcgccttg atcgttggga accggagctg aatgaagcca
     2581 taccaaacga cgagcgtgac accacgatgc ctgtagcaat ggcaacaacg ttgcgcaaac
     2641 tattaactgg cgaactactt actctagctt cccggcaaca attaatagac tggatggagg
     2701 cggataaagt tgcaggacca cttctgcgct cggcccttcc ggctggctgg tttattgctg
     2761 ataaatctgg agccggtgag cgtgggtctc gcggtatcat tgcagcactg gggccagatg
     2821 gtaagccctc ccgtatcgta gttatctaca cgacggggag tcaggcaact atggatgaac
     2881 gaaatagaca gatcgctgag ataggtgcct cactgattaa gcattggtaa ctgtcagacc
     2941 aagtttactc atatatactt tagattgatt taaaacttca tttttaattt aaaaggatct
     3001 aggtgaagat cctttttgat aatctcatga ccaaaatccc ttaacgtgag ttttcgttcc
     3061 actgagcgtc agaccccgta gaaaagatca aaggatcttc ttgagatcct ttttttctgc
     3121 gcgtaatctg ctgcttgcaa acaaaaaaac caccgctacc agcggtggtt tgtttgccgg
     3181 atcaagagct accaactctt tttccgaagg taactggctt cagcagagcg cagataccaa
     3241 atactgttct tctagtgtag ccgtagttag gccaccactt caagaactct gtagcaccgc
     3301 ctacatacct cgctctgcta atcctgttac cagtggctgc tgccagtggc gataagtcgt
     3361 gtcttaccgg gttggactca agacgatagt taccggataa ggcgcagcgg tcgggctgaa
     3421 cggggggttc gtgcacacag cccagcttgg agcgaacgac ctacaccgaa ctgagatacc
     3481 tacagcgtga gctatgagaa agcgccacgc ttcccgaagg gagaaaggcg gacaggtatc
     3541 cggtaagcgg cagggtcgga acaggagagc gcacgaggga gcttccaggg ggaaacgcct
     3601 ggtatcttta tagtcctgtc gggtttcgcc acctctgact tgagcgtcga tttttgtgat
     3661 gctcgtcagg ggggcggagc ctatggaaaa acgccagcaa cgcggccttt ttacggttcc
     3721 tggccttttg ctggcctttt gctcacatgt tctttcctgc gttatcccct gattctgtgg
     3781 ataaccgtat taccgccttt gagtgagctg ataccgctcg ccgcagccga acgaccgagc
     3841 gcagcgagtc agtgagcgag gaagcggaag agcgcccaat acgcaaaccg cctctccccg
     3901 cgcgttggcc gattcattaa tgcagctggc acgacaggtt tcccgactgg aaagcgggca
     3961 gtgagcgcaa cgcaattaat gtgagttagc tcactcatta ggcaccccag gctttacact
     4021 ttatgcttcc ggctcgtatg ttgtgtggaa ttgtgagcgg ataacaattt cacacaggaa
     4081 acagctatga ccatgattac gccaagcgcg caattaaccc tcactaaagg gaacaaaagc
     4141 tggagctgca agcttaatgt agtcttatgc aatactcttg tagtcttgca acatggtaac
     4201 gatgagttag caacatgcct tacaaggaga gaaaaagcac cgtgcatgcc gattggtgga
     4261 agtaaggtgg tacgatcgtg ccttattagg aaggcaacag acgggtctga catggattgg
     4321 acgaaccact gaattgccgc attgcagaga tattgtattt aagtgcctag ctcgatacat
     4381 aaacgggtct ctctggttag accagatctg agcctgggag ctctctggct aactagggaa
     4441 cccactgctt aagcctcaat aaagcttgcc ttgagtgctt caagtagtgt gtgcccgtct
     4501 gttgtgtgac tctggtaact agagatccct cagacccttt tagtcagtgt ggaaaatctc
     4561 tagcagtggc gcccgaacag ggacttgaaa gcgaaaggga aaccagagga gctctctcga
     4621 cgcaggactc ggcttgctga agcgcgcacg gcaagaggcg aggggcggcg actggtgagt
     4681 acgccaaaaa ttttgactag cggaggctag aaggagagag atgggtgcga gagcgtcagt
     4741 attaagcggg ggagaattag atcgcgatgg gaaaaaattc ggttaaggcc agggggaaag
     4801 aaaaaatata aattaaaaca tatagtatgg gcaagcaggg agctagaacg attcgcagtt
     4861 aatcctggcc tgttagaaac atcagaaggc tgtagacaaa tactgggaca gctacaacca
     4921 tcccttcaga caggatcaga agaacttaga tcattatata atacagtagc aaccctctat
     4981 tgtgtgcatc aaaggataga gataaaagac accaaggaag ctttagacaa gatagaggaa
     5041 gagcaaaaca aaagtaagac caccgcacag caagcggccg ctgatcttca gacctggagg
     5101 aggagatatg agggacaatt ggagaagtga attatataaa tataaagtag taaaaattga
     5161 accattagga gtagcaccca ccaaggcaaa gagaagagtg gtgcagagag aaaaaagagc
     5221 agtgggaata ggagctttgt tccttgggtt cttgggagca gcaggaagca ctatgggcgc
     5281 agcgtcaatg acgctgacgg tacaggccag acaattattg tctggtatag tgcagcagca
     5341 gaacaatttg ctgagggcta ttgaggcgca acagcatctg ttgcaactca cagtctgggg
     5401 catcaagcag ctccaggcaa gaatcctggc tgtggaaaga tacctaaagg atcaacagct
     5461 cctggggatt tggggttgct ctggaaaact catttgcacc actgctgtgc cttggaatgc
     5521 tagttggagt aataaatctc tggaacagat ttggaatcac acgacctgga tggagtggga
     5581 cagagaaatt aacaattaca caagcttaat acactcctta attgaagaat cgcaaaacca
     5641 gcaagaaaag aatgaacaag aattattgga attagataaa tgggcaagtt tgtggaattg
     5701 gtttaacata acaaattggc tgtggtatat aaaattattc ataatgatag taggaggctt
     5761 ggtaggttta agaatagttt ttgctgtact ttctatagtg aatagagtta ggcagggata
     5821 ttcaccatta tcgtttcaga cccacctccc aaccccgagg ggacccgaca ggcccgaagg
     5881 aatagaagaa gaaggtggag agagagacag agacagatcc attcgattag tgaacggatc
     5941 tcgacggtat cggttaactt ttaaaagaaa aggggggatt ggggggtaca gtgcagggga
     6001 aagaatagta gacataatag caacagacat acaaactaaa gaattacaaa aacaaattac
     6061 aaaaattcaa aattttatcg atgagtaatt catacaaaag gactcgcccc tgccttgggg
     6121 aatcccaggg accgtcgtta aactcccact aacgtagaac ccagagatcg ctgcgttccc
     6181 gccccctcac ccgcccgctc tcgtcatcac tgaggtggag aagagcatgc gtgaggctcc
     6241 ggtgcccgtc agtgggcaga gcgcacatcg cccacagtcc ccgagaagtt ggggggaggg
     6301 gtcggcaatt gaaccggtgc ctagagaagg tggcgcgggg taaactggga aagtgatgtc
     6361 gtgtactggc tccgcctttt tcccgagggt gggggagaac cgtatataag tgcagtagtc
     6421 gccgtgaacg ttctttttcg caacgggttt gccgccagaa cacaggtaag tgccgtgtgt
     6481 ggttcccgcg ggcctggcct ctttacgggt tatggccctt gcgtgccttg aattacttcc
     6541 acgcccctgg ctgcagtacg tgattcttga tcccgagctt cgggttggaa gtgggtggga
     6601 gagttcgagg ccttgcgctt aaggagcccc ttcgcctcgt gcttgagttg aggcctggct
     6661 tgggcgctgg ggccgccgcg tgcgaatctg gtggcacctt cgcgcctgtc tcgctgcttt
     6721 cgataagtct ctagccattt aaaatttttg atgacctgct gcgacgcttt ttttctggca
     6781 agatagtctt gtaaatgcgg gccaagatct gcacactggt atttcggttt ttggggccgc
     6841 gggcggcgac ggggcccgtg cgtcccagcg cacatgttcg gcgaggcggg gcctgcgagc
     6901 gcggccaccg agaatcggac gggggtagtc tcaagctggc cggcctgctc tggtgcctgg
     6961 cctcgcgccg ccgtgtatcg ccccgccctg ggcggcaagg ctggcccggt cggcaccagt
     7021 tgcgtgagcg gaaagatggc cgcttcccgg ccctgctgca gggagctcaa aatggaggac
     7081 gcggcgctcg ggagagcggg cgggtgagtc acccacacaa aggaaaaggg cctttccgtc
     7141 ctcagccgtc gcttcatgtg actccacgga gtaccgggcg ccgtccaggc acctcgatta
     7201 gttctcaagc ttttggagta cgtcgtcttt aggttggggg gaggggtttt atgcgatgga
     7261 gtttccccac actgagtggg tggagactga agttaggcca gcttggcact tgatgtaatt
     7321 ctccttggaa tttgcccttt ttgagtttgg atcttggttc attctcaagc ctcagacagt
     7381 ggttcaaagt ttttttcttc catttcaggt gtcgtgaccc tagcgctacc tctagagcca
     7441 ccatgagagt gatggcccct agaaccctga tcctgctgct gagcggcgcc ctggccctga
     7501 ccgagacatg ggcctgcagc cactccatgc gctatttcga caccgccgtg tcccggcccg
     7561 gaagaggaga gcctcgcttt atctctgtgg gctacgtgga cgatacacag ttcgtgcggt
     7621 ttgacagcga tgcagcatcc cctaggggag agccaagggc accctgggtg gagcaggagg
     7681 gcccagagta ttgggaccgg gagacacaga agtacaagag gcaggcccag gccgatcgcg
     7741 tgaacctgag gaagctgcgc ggctactata atcagtctga ggacggcagc cacacactgc
     7801 agtggatgta tggatgtgac ctgggaccag atggccggct gctgagaggc tacgatcaga
     7861 gcgcctatga cggcaaggat tacatcgccc tgaacgagga cctgaggtcc tggaccgcag
     7921 cagatacagc agcacagatc acccagagga agtgggaggc agcaagagag gcagagcagt
     7981 ggagggccta tctggaggga acctgcgtgg agtggctgcg gagatacctg gagaatggca
     8041 aggagacact gcagagagcc gagcacccaa agacccacgt gacacaccac cccgtgtctg
     8101 accacgaggc cacactgagg tgttgggccc tgggcttcta tccagccgag atcaccctga
     8161 catggcagag agacggcgag gatcagaccc aggacacaga gctggtggag acacggcccg
     8221 caggcgatgg cacatttcag aagtgggcag cagtggtggt gccttccgga gaggagcagc
     8281 ggtacacctg ccacgtgcag cacgagggcc tgccagagcc tctgaccctg agatgggagc
     8341 caagctccca gcctacaatc ccaatcgtgg gaatcgtggc aggcctggcc gtgctggccg
     8401 tgctggccgt gctgggtgcc gtgatggccg tggtcatgtg caggaggaag tctagcggag
     8461 gcaagggagg atcttgtagc caggcagcat cctctaactc cgcccaggga tccgatgagt
     8521 ctctgatcgc ctgtaaggcc tgaggcgcgc ctcccagagc caccgttaca cggacgcagt
     8581 agtcccggtg gcaccgactc ggtgccactt tttcaagttg ataacggact agccttattt
     8641 taacttgcta tttctagctc taaaactttt tttgactaga tcttgagaca aatggcagta
     8701 ttcatccaca aggtacgcta aacgcctgca accaggacgc gt
//

Lenti plKO_EF1A_ORF_HLA_C*07 01

LOCUS plKO_EF1A_ORF_HLA_C*07 01                 8742 bp DNA     circular SYN 24-OCT-2022
DEFINITION  synthetic circular DNA
ACCESSION   .
VERSION     .
KEYWORDS    .
SOURCE      synthetic DNA construct
  ORGANISM  synthetic DNA construct
REFERENCE   1  (bases 1 to 8742)
  AUTHORS   .
  TITLE     Direct Submission
  JOURNAL   Exported Oct 25, 2022 from SnapGene 6.0.5
            https://www.snapgene.com
COMMENT     U515NEI230-200 DNA889110
FEATURES             Location/Qualifiers
     source          1..8742
                     /mol_type="other DNA"
                     /organism="synthetic DNA construct"
     misc_feature    10..598
                     /label=WPRE
     CDS             complement(481..492)
                     /label=Factor Xa site
     LTR             670..903
                     /label=3' LTR (Delta-U3)
     polyA_signal    981..1102
                     /label=SV40 poly(A) signal
     rep_origin      1142..1277
                     /label=SV40 ori
     promoter        complement(1298..1316)
                     /label=T7 promoter
     primer_bind     complement(1326..1342)
                     /label=M13 fwd
     rep_origin      1484..1939
                     /label=f1 ori
     promoter        1965..2069
                     /label=AmpR promoter
     CDS             2070..2138
                     /label=AmpR
     CDS             2139..2930
                     /label=AmpR
     rep_origin      3101..3689
                     /label=ori
     promoter        4019..4036
                     /label=lac promoter
     protein_bind    4051..4067
                     /label=lac operator
     primer_bind     4075..4091
                     /label=M13 rev
     promoter        4112..4130
                     /label=T3 promoter
     promoter        4158..4384
                     /label=RSV promoter
     LTR             4385..4565
                     /label=5' LTR (truncated)
     misc_feature    4612..4737
                     /label=HIV-1 Psi
     misc_feature    5230..5463
                     /label=RRE
     misc_feature    5959..6076
                     /label=cPPT/CTS
     promoter        6236..7417
                     /label=EF-1-alpha promoter
     intron          6466..7408
                     /label=EF-1-alpha intron A
     CDS             7443..8543
                     /codon_start=1
                     /label=HLA-C*07:01
                     /translation="MRVMAPRALLLLLSGGLALTETWACSHSMRYFDTAVSRPGRGEPR
                     FISVGYVDDTQFVRFDSDAASPRGEPRAPWVEQEGPEYWDRETQNYKRQAQADRVSLRN
                     LRGYYNQSEDGSHTLQRMYGCDLGPDGRLLRGYDQSAYDGKDYIALNEDLRSWTAADTA
                     AQITQRKLEAARAAEQLRAYLEGTCVEWLRRYLENGKETLQRAEPPKTHVTHHPLSDHE
                     ATLRCWALGFYPAEITLTWQRDGEDQTQDTELVETRPAGDGTFQKWAAVVVPSGQEQRY
                     TCHMQHEGLQEPLTLSWEPSSQPTIPIMGIVAGLAVLVVLAVLGAVVTAMMCRRKSSGG
                     KGGSCSQAACSNSAQGSDESLITCKA"
     misc_RNA        complement(8591..8666)
                     /label=gRNA scaffold/Barcode
ORIGIN
        1 taagtcgaca atcaacctct ggattacaaa atttgtgaaa gattgactgg tattcttaac
       61 tatgttgctc cttttacgct atgtggatac gctgctttaa tgcctttgta tcatgctatt
      121 gcttcccgta tggctttcat tttctcctcc ttgtataaat cctggttgct gtctctttat
      181 gaggagttgt ggcccgttgt caggcaacgt ggcgtggtgt gcactgtgtt tgctgacgca
      241 acccccactg gttggggcat tgccaccacc tgtcagctcc tttccgggac tttcgctttc
      301 cccctcccta ttgccacggc ggaactcatc gccgcctgcc ttgcccgctg ctggacaggg
      361 gctcggctgt tgggcactga caattccgtg gtgttgtcgg ggaaatcatc gtcctttcct
      421 tggctgctcg cctgtgttgc cacctggatt ctgcgcggga cgtccttctg ctacgtccct
      481 tcggccctca atccagcgga ccttccttcc cgcggcctgc tgccggctct gcggcctctt
      541 ccgcgtcttc gccttcgccc tcagacgagt cggatctccc tttgggccgc ctccccgcgt
      601 cgactttaag accaatgact tacaaggcag ctgtagatct tagccacttt ttaaaagaaa
      661 aggggggact ggaagggcta attcactccc aacgaagaca agatctgctt tttgcttgta
      721 ctgggtctct ctggttagac cagatctgag cctgggagct ctctggctaa ctagggaacc
      781 cactgcttaa gcctcaataa agcttgcctt gagtgcttca agtagtgtgt gcccgtctgt
      841 tgtgtgactc tggtaactag agatccctca gaccctttta gtcagtgtgg aaaatctcta
      901 gcagtacgta tagtagttca tgtcatctta ttattcagta tttataactt gcaaagaaat
      961 gaatatcaga gagtgagagg aacttgttta ttgcagctta taatggttac aaataaagca
     1021 atagcatcac aaatttcaca aataaagcat ttttttcact gcattctagt tgtggtttgt
     1081 ccaaactcat caatgtatct tatcatgtct ggctctagct atcccgcccc taactccgcc
     1141 catcccgccc ctaactccgc ccagttccgc ccattctccg ccccatggct gactaatttt
     1201 ttttatttat gcagaggccg aggccgcctc ggcctctgag ctattccaga agtagtgagg
     1261 aggctttttt ggaggcctag ggacgtaccc aattcgccct atagtgagtc gtattacgcg
     1321 cgctcactgg ccgtcgtttt acaacgtcgt gactgggaaa accctggcgt tacccaactt
     1381 aatcgccttg cagcacatcc ccctttcgcc agctggcgta atagcgaaga ggcccgcacc
     1441 gatcgccctt cccaacagtt gcgcagcctg aatggcgaat gggacgcgcc ctgtagcggc
     1501 gcattaagcg cggcgggtgt ggtggttacg cgcagcgtga ccgctacact tgccagcgcc
     1561 ctagcgcccg ctcctttcgc tttcttccct tcctttctcg ccacgttcgc cggctttccc
     1621 cgtcaagctc taaatcgggg gctcccttta gggttccgat ttagtgcttt acggcacctc
     1681 gaccccaaaa aacttgatta gggtgatggt tcacgtagtg ggccatcgcc ctgatagacg
     1741 gtttttcgcc ctttgacgtt ggagtccacg ttctttaata gtggactctt gttccaaact
     1801 ggaacaacac tcaaccctat ctcggtctat tcttttgatt tataagggat tttgccgatt
     1861 tcggcctatt ggttaaaaaa tgagctgatt taacaaaaat ttaacgcgaa ttttaacaaa
     1921 atattaacgc ttacaattta ggtggcactt ttcggggaaa tgtgcgcgga acccctattt
     1981 gtttattttt ctaaatacat tcaaatatgt atccgctcat gagacaataa ccctgataaa
     2041 tgcttcaata atattgaaaa aggaagagta tgagtattca acatttccgt gtcgccctta
     2101 ttcccttttt tgcggcattt tgccttcctg tttttgctca cccagaaacg ctggtgaaag
     2161 taaaagatgc tgaagatcag ttgggtgcac gagtgggtta catcgaactg gatctcaaca
     2221 gcggtaagat ccttgagagt tttcgccccg aagaacgttt tccaatgatg agcactttta
     2281 aagttctgct atgtggcgcg gtattatccc gtattgacgc cgggcaagag caactcggtc
     2341 gccgcataca ctattctcag aatgacttgg ttgagtactc accagtcaca gaaaagcatc
     2401 ttacggatgg catgacagta agagaattat gcagtgctgc cataaccatg agtgataaca
     2461 ctgcggccaa cttacttctg acaacgatcg gaggaccgaa ggagctaacc gcttttttgc
     2521 acaacatggg ggatcatgta actcgccttg atcgttggga accggagctg aatgaagcca
     2581 taccaaacga cgagcgtgac accacgatgc ctgtagcaat ggcaacaacg ttgcgcaaac
     2641 tattaactgg cgaactactt actctagctt cccggcaaca attaatagac tggatggagg
     2701 cggataaagt tgcaggacca cttctgcgct cggcccttcc ggctggctgg tttattgctg
     2761 ataaatctgg agccggtgag cgtgggtctc gcggtatcat tgcagcactg gggccagatg
     2821 gtaagccctc ccgtatcgta gttatctaca cgacggggag tcaggcaact atggatgaac
     2881 gaaatagaca gatcgctgag ataggtgcct cactgattaa gcattggtaa ctgtcagacc
     2941 aagtttactc atatatactt tagattgatt taaaacttca tttttaattt aaaaggatct
     3001 aggtgaagat cctttttgat aatctcatga ccaaaatccc ttaacgtgag ttttcgttcc
     3061 actgagcgtc agaccccgta gaaaagatca aaggatcttc ttgagatcct ttttttctgc
     3121 gcgtaatctg ctgcttgcaa acaaaaaaac caccgctacc agcggtggtt tgtttgccgg
     3181 atcaagagct accaactctt tttccgaagg taactggctt cagcagagcg cagataccaa
     3241 atactgttct tctagtgtag ccgtagttag gccaccactt caagaactct gtagcaccgc
     3301 ctacatacct cgctctgcta atcctgttac cagtggctgc tgccagtggc gataagtcgt
     3361 gtcttaccgg gttggactca agacgatagt taccggataa ggcgcagcgg tcgggctgaa
     3421 cggggggttc gtgcacacag cccagcttgg agcgaacgac ctacaccgaa ctgagatacc
     3481 tacagcgtga gctatgagaa agcgccacgc ttcccgaagg gagaaaggcg gacaggtatc
     3541 cggtaagcgg cagggtcgga acaggagagc gcacgaggga gcttccaggg ggaaacgcct
     3601 ggtatcttta tagtcctgtc gggtttcgcc acctctgact tgagcgtcga tttttgtgat
     3661 gctcgtcagg ggggcggagc ctatggaaaa acgccagcaa cgcggccttt ttacggttcc
     3721 tggccttttg ctggcctttt gctcacatgt tctttcctgc gttatcccct gattctgtgg
     3781 ataaccgtat taccgccttt gagtgagctg ataccgctcg ccgcagccga acgaccgagc
     3841 gcagcgagtc agtgagcgag gaagcggaag agcgcccaat acgcaaaccg cctctccccg
     3901 cgcgttggcc gattcattaa tgcagctggc acgacaggtt tcccgactgg aaagcgggca
     3961 gtgagcgcaa cgcaattaat gtgagttagc tcactcatta ggcaccccag gctttacact
     4021 ttatgcttcc ggctcgtatg ttgtgtggaa ttgtgagcgg ataacaattt cacacaggaa
     4081 acagctatga ccatgattac gccaagcgcg caattaaccc tcactaaagg gaacaaaagc
     4141 tggagctgca agcttaatgt agtcttatgc aatactcttg tagtcttgca acatggtaac
     4201 gatgagttag caacatgcct tacaaggaga gaaaaagcac cgtgcatgcc gattggtgga
     4261 agtaaggtgg tacgatcgtg ccttattagg aaggcaacag acgggtctga catggattgg
     4321 acgaaccact gaattgccgc attgcagaga tattgtattt aagtgcctag ctcgatacat
     4381 aaacgggtct ctctggttag accagatctg agcctgggag ctctctggct aactagggaa
     4441 cccactgctt aagcctcaat aaagcttgcc ttgagtgctt caagtagtgt gtgcccgtct
     4501 gttgtgtgac tctggtaact agagatccct cagacccttt tagtcagtgt ggaaaatctc
     4561 tagcagtggc gcccgaacag ggacttgaaa gcgaaaggga aaccagagga gctctctcga
     4621 cgcaggactc ggcttgctga agcgcgcacg gcaagaggcg aggggcggcg actggtgagt
     4681 acgccaaaaa ttttgactag cggaggctag aaggagagag atgggtgcga gagcgtcagt
     4741 attaagcggg ggagaattag atcgcgatgg gaaaaaattc ggttaaggcc agggggaaag
     4801 aaaaaatata aattaaaaca tatagtatgg gcaagcaggg agctagaacg attcgcagtt
     4861 aatcctggcc tgttagaaac atcagaaggc tgtagacaaa tactgggaca gctacaacca
     4921 tcccttcaga caggatcaga agaacttaga tcattatata atacagtagc aaccctctat
     4981 tgtgtgcatc aaaggataga gataaaagac accaaggaag ctttagacaa gatagaggaa
     5041 gagcaaaaca aaagtaagac caccgcacag caagcggccg ctgatcttca gacctggagg
     5101 aggagatatg agggacaatt ggagaagtga attatataaa tataaagtag taaaaattga
     5161 accattagga gtagcaccca ccaaggcaaa gagaagagtg gtgcagagag aaaaaagagc
     5221 agtgggaata ggagctttgt tccttgggtt cttgggagca gcaggaagca ctatgggcgc
     5281 agcgtcaatg acgctgacgg tacaggccag acaattattg tctggtatag tgcagcagca
     5341 gaacaatttg ctgagggcta ttgaggcgca acagcatctg ttgcaactca cagtctgggg
     5401 catcaagcag ctccaggcaa gaatcctggc tgtggaaaga tacctaaagg atcaacagct
     5461 cctggggatt tggggttgct ctggaaaact catttgcacc actgctgtgc cttggaatgc
     5521 tagttggagt aataaatctc tggaacagat ttggaatcac acgacctgga tggagtggga
     5581 cagagaaatt aacaattaca caagcttaat acactcctta attgaagaat cgcaaaacca
     5641 gcaagaaaag aatgaacaag aattattgga attagataaa tgggcaagtt tgtggaattg
     5701 gtttaacata acaaattggc tgtggtatat aaaattattc ataatgatag taggaggctt
     5761 ggtaggttta agaatagttt ttgctgtact ttctatagtg aatagagtta ggcagggata
     5821 ttcaccatta tcgtttcaga cccacctccc aaccccgagg ggacccgaca ggcccgaagg
     5881 aatagaagaa gaaggtggag agagagacag agacagatcc attcgattag tgaacggatc
     5941 tcgacggtat cggttaactt ttaaaagaaa aggggggatt ggggggtaca gtgcagggga
     6001 aagaatagta gacataatag caacagacat acaaactaaa gaattacaaa aacaaattac
     6061 aaaaattcaa aattttatcg atgagtaatt catacaaaag gactcgcccc tgccttgggg
     6121 aatcccaggg accgtcgtta aactcccact aacgtagaac ccagagatcg ctgcgttccc
     6181 gccccctcac ccgcccgctc tcgtcatcac tgaggtggag aagagcatgc gtgaggctcc
     6241 ggtgcccgtc agtgggcaga gcgcacatcg cccacagtcc ccgagaagtt ggggggaggg
     6301 gtcggcaatt gaaccggtgc ctagagaagg tggcgcgggg taaactggga aagtgatgtc
     6361 gtgtactggc tccgcctttt tcccgagggt gggggagaac cgtatataag tgcagtagtc
     6421 gccgtgaacg ttctttttcg caacgggttt gccgccagaa cacaggtaag tgccgtgtgt
     6481 ggttcccgcg ggcctggcct ctttacgggt tatggccctt gcgtgccttg aattacttcc
     6541 acgcccctgg ctgcagtacg tgattcttga tcccgagctt cgggttggaa gtgggtggga
     6601 gagttcgagg ccttgcgctt aaggagcccc ttcgcctcgt gcttgagttg aggcctggct
     6661 tgggcgctgg ggccgccgcg tgcgaatctg gtggcacctt cgcgcctgtc tcgctgcttt
     6721 cgataagtct ctagccattt aaaatttttg atgacctgct gcgacgcttt ttttctggca
     6781 agatagtctt gtaaatgcgg gccaagatct gcacactggt atttcggttt ttggggccgc
     6841 gggcggcgac ggggcccgtg cgtcccagcg cacatgttcg gcgaggcggg gcctgcgagc
     6901 gcggccaccg agaatcggac gggggtagtc tcaagctggc cggcctgctc tggtgcctgg
     6961 cctcgcgccg ccgtgtatcg ccccgccctg ggcggcaagg ctggcccggt cggcaccagt
     7021 tgcgtgagcg gaaagatggc cgcttcccgg ccctgctgca gggagctcaa aatggaggac
     7081 gcggcgctcg ggagagcggg cgggtgagtc acccacacaa aggaaaaggg cctttccgtc
     7141 ctcagccgtc gcttcatgtg actccacgga gtaccgggcg ccgtccaggc acctcgatta
     7201 gttctcaagc ttttggagta cgtcgtcttt aggttggggg gaggggtttt atgcgatgga
     7261 gtttccccac actgagtggg tggagactga agttaggcca gcttggcact tgatgtaatt
     7321 ctccttggaa tttgcccttt ttgagtttgg atcttggttc attctcaagc ctcagacagt
     7381 ggttcaaagt ttttttcttc catttcaggt gtcgtgaccc tagcgctacc tctagagcca
     7441 ccatgagagt gatggcccct agagccctgc tgctgctgct gtccggaggc ctggccctga
     7501 ccgagacatg ggcctgctcc cactctatgc gctatttcga caccgccgtg tcccggcccg
     7561 gaagaggaga gcctcgcttt atctctgtgg gctacgtgga cgatacacag ttcgtgcggt
     7621 ttgacagcga tgcagcatcc cctaggggag agccaagggc accctgggtg gagcaggagg
     7681 gcccagagta ttgggaccgg gagacacaga actacaagag gcaggcccag gccgatcgcg
     7741 tgtctctgag gaacctgcgc ggctactata atcagtctga ggacggcagc cacacactgc
     7801 agagaatgta tggatgtgac ctgggaccag atggccggct gctgagaggc tacgatcaga
     7861 gcgcctatga cggcaaggat tacatcgccc tgaacgagga cctgaggtcc tggaccgcag
     7921 cagatacagc agcacagatc acccagcgga agctggaggc agcaagagca gcagagcagc
     7981 tgagggccta tctggaggga acctgcgtgg agtggctgcg gagatacctg gagaatggca
     8041 aggagacact gcagagggca gagccaccta agacccacgt gacacaccac ccactgagcg
     8101 accacgaggc caccctgagg tgttgggcac tgggcttcta tcccgccgag atcaccctga
     8161 catggcagag agacggcgag gatcagaccc aggacacaga gctggtggag acacggcccg
     8221 caggcgatgg cacatttcag aagtgggcag cagtggtggt gccttctgga caggagcaga
     8281 ggtacacctg ccacatgcag cacgagggcc tgcaggagcc actgaccctg agctgggagc
     8341 caagctccca gcctacaatc ccaatcatgg gcatcgtggc aggcctggcc gtgctggtgg
     8401 tgctggccgt gctgggcgcc gtggtgaccg ccatgatgtg caggaggaag tctagcggag
     8461 gcaagggagg aagctgctcc caggccgcct gttctaatag cgcccagggc tccgatgagt
     8521 ctctgatcac ctgtaaggcc tgaggcgcgc ctcccagagc caccgttaca ctggcgcttc
     8581 tgctgtgcgc gcaccgactc ggtgccactt tttcaagttg ataacggact agccttattt
     8641 taacttgcta tttctagctc taaaactttt tttgactaga tcttgagaca aatggcagta
     8701 ttcatccaca aggtacgcta aacgcctgca accaggacgc gt
//

Lenti plKO_EF1A_ORF_HLA_C*07 02

LOCUS  plKO_EF1A_ORF_HLA_C*07 02                8742 bp DNA     circular SYN 24-OCT-2022
DEFINITION  synthetic circular DNA
ACCESSION   .
VERSION     .
KEYWORDS    .
SOURCE      synthetic DNA construct
  ORGANISM  synthetic DNA construct
REFERENCE   1  (bases 1 to 8742)
  AUTHORS   .
  TITLE     Direct Submission
  JOURNAL   Exported Oct 25, 2022 from SnapGene 6.0.5
            https://www.snapgene.com
COMMENT     U515NEI230-202 DNA889111
FEATURES             Location/Qualifiers
     source          1..8742
                     /mol_type="other DNA"
                     /organism="synthetic DNA construct"
     misc_feature    10..598
                     /label=WPRE
     CDS             complement(481..492)
                     /label=Factor Xa site
     LTR             670..903
                     /label=3' LTR (Delta-U3)
     polyA_signal    981..1102
                     /label=SV40 poly(A) signal
     rep_origin      1142..1277
                     /label=SV40 ori
     promoter        complement(1298..1316)
                     /label=T7 promoter
     primer_bind     complement(1326..1342)
                     /label=M13 fwd
     rep_origin      1484..1939
                     /label=f1 ori
     promoter        1965..2069
                     /label=AmpR promoter
     CDS             2070..2138
                     /label=AmpR
     CDS             2139..2930
                     /label=AmpR
     rep_origin      3101..3689
                     /label=ori
     promoter        4019..4036
                     /label=lac promoter
     protein_bind    4051..4067
                     /label=lac operator
     primer_bind     4075..4091
                     /label=M13 rev
     promoter        4112..4130
                     /label=T3 promoter
     promoter        4158..4384
                     /label=RSV promoter
     LTR             4385..4565
                     /label=5' LTR (truncated)
     misc_feature    4612..4737
                     /label=HIV-1 Psi
     misc_feature    5230..5463
                     /label=RRE
     misc_feature    5959..6076
                     /label=cPPT/CTS
     promoter        6236..7417
                     /label=EF-1-alpha promoter
     intron          6466..7408
                     /label=EF-1-alpha intron A
     CDS             7443..8543
                     /codon_start=1
                     /label=HLA-C*07:02
                     /translation="MRVMAPRALLLLLSGGLALTETWACSHSMRYFDTAVSRPGRGEPR
                     FISVGYVDDTQFVRFDSDAASPRGEPRAPWVEQEGPEYWDRETQKYKRQAQADRVSLRN
                     LRGYYNQSEDGSHTLQRMSGCDLGPDGRLLRGYDQSAYDGKDYIALNEDLRSWTAADTA
                     AQITQRKLEAARAAEQLRAYLEGTCVEWLRRYLENGKETLQRAEPPKTHVTHHPLSDHE
                     ATLRCWALGFYPAEITLTWQRDGEDQTQDTELVETRPAGDGTFQKWAAVVVPSGQEQRY
                     TCHMQHEGLQEPLTLSWEPSSQPTIPIMGIVAGLAVLVVLAVLGAVVTAMMCRRKSSGG
                     KGGSCSQAACSNSAQGSDESLITCKA"
     misc_RNA        complement(8591..8666)
                     /label=gRNA scaffold/Barcode
ORIGIN
        1 taagtcgaca atcaacctct ggattacaaa atttgtgaaa gattgactgg tattcttaac
       61 tatgttgctc cttttacgct atgtggatac gctgctttaa tgcctttgta tcatgctatt
      121 gcttcccgta tggctttcat tttctcctcc ttgtataaat cctggttgct gtctctttat
      181 gaggagttgt ggcccgttgt caggcaacgt ggcgtggtgt gcactgtgtt tgctgacgca
      241 acccccactg gttggggcat tgccaccacc tgtcagctcc tttccgggac tttcgctttc
      301 cccctcccta ttgccacggc ggaactcatc gccgcctgcc ttgcccgctg ctggacaggg
      361 gctcggctgt tgggcactga caattccgtg gtgttgtcgg ggaaatcatc gtcctttcct
      421 tggctgctcg cctgtgttgc cacctggatt ctgcgcggga cgtccttctg ctacgtccct
      481 tcggccctca atccagcgga ccttccttcc cgcggcctgc tgccggctct gcggcctctt
      541 ccgcgtcttc gccttcgccc tcagacgagt cggatctccc tttgggccgc ctccccgcgt
      601 cgactttaag accaatgact tacaaggcag ctgtagatct tagccacttt ttaaaagaaa
      661 aggggggact ggaagggcta attcactccc aacgaagaca agatctgctt tttgcttgta
      721 ctgggtctct ctggttagac cagatctgag cctgggagct ctctggctaa ctagggaacc
      781 cactgcttaa gcctcaataa agcttgcctt gagtgcttca agtagtgtgt gcccgtctgt
      841 tgtgtgactc tggtaactag agatccctca gaccctttta gtcagtgtgg aaaatctcta
      901 gcagtacgta tagtagttca tgtcatctta ttattcagta tttataactt gcaaagaaat
      961 gaatatcaga gagtgagagg aacttgttta ttgcagctta taatggttac aaataaagca
     1021 atagcatcac aaatttcaca aataaagcat ttttttcact gcattctagt tgtggtttgt
     1081 ccaaactcat caatgtatct tatcatgtct ggctctagct atcccgcccc taactccgcc
     1141 catcccgccc ctaactccgc ccagttccgc ccattctccg ccccatggct gactaatttt
     1201 ttttatttat gcagaggccg aggccgcctc ggcctctgag ctattccaga agtagtgagg
     1261 aggctttttt ggaggcctag ggacgtaccc aattcgccct atagtgagtc gtattacgcg
     1321 cgctcactgg ccgtcgtttt acaacgtcgt gactgggaaa accctggcgt tacccaactt
     1381 aatcgccttg cagcacatcc ccctttcgcc agctggcgta atagcgaaga ggcccgcacc
     1441 gatcgccctt cccaacagtt gcgcagcctg aatggcgaat gggacgcgcc ctgtagcggc
     1501 gcattaagcg cggcgggtgt ggtggttacg cgcagcgtga ccgctacact tgccagcgcc
     1561 ctagcgcccg ctcctttcgc tttcttccct tcctttctcg ccacgttcgc cggctttccc
     1621 cgtcaagctc taaatcgggg gctcccttta gggttccgat ttagtgcttt acggcacctc
     1681 gaccccaaaa aacttgatta gggtgatggt tcacgtagtg ggccatcgcc ctgatagacg
     1741 gtttttcgcc ctttgacgtt ggagtccacg ttctttaata gtggactctt gttccaaact
     1801 ggaacaacac tcaaccctat ctcggtctat tcttttgatt tataagggat tttgccgatt
     1861 tcggcctatt ggttaaaaaa tgagctgatt taacaaaaat ttaacgcgaa ttttaacaaa
     1921 atattaacgc ttacaattta ggtggcactt ttcggggaaa tgtgcgcgga acccctattt
     1981 gtttattttt ctaaatacat tcaaatatgt atccgctcat gagacaataa ccctgataaa
     2041 tgcttcaata atattgaaaa aggaagagta tgagtattca acatttccgt gtcgccctta
     2101 ttcccttttt tgcggcattt tgccttcctg tttttgctca cccagaaacg ctggtgaaag
     2161 taaaagatgc tgaagatcag ttgggtgcac gagtgggtta catcgaactg gatctcaaca
     2221 gcggtaagat ccttgagagt tttcgccccg aagaacgttt tccaatgatg agcactttta
     2281 aagttctgct atgtggcgcg gtattatccc gtattgacgc cgggcaagag caactcggtc
     2341 gccgcataca ctattctcag aatgacttgg ttgagtactc accagtcaca gaaaagcatc
     2401 ttacggatgg catgacagta agagaattat gcagtgctgc cataaccatg agtgataaca
     2461 ctgcggccaa cttacttctg acaacgatcg gaggaccgaa ggagctaacc gcttttttgc
     2521 acaacatggg ggatcatgta actcgccttg atcgttggga accggagctg aatgaagcca
     2581 taccaaacga cgagcgtgac accacgatgc ctgtagcaat ggcaacaacg ttgcgcaaac
     2641 tattaactgg cgaactactt actctagctt cccggcaaca attaatagac tggatggagg
     2701 cggataaagt tgcaggacca cttctgcgct cggcccttcc ggctggctgg tttattgctg
     2761 ataaatctgg agccggtgag cgtgggtctc gcggtatcat tgcagcactg gggccagatg
     2821 gtaagccctc ccgtatcgta gttatctaca cgacggggag tcaggcaact atggatgaac
     2881 gaaatagaca gatcgctgag ataggtgcct cactgattaa gcattggtaa ctgtcagacc
     2941 aagtttactc atatatactt tagattgatt taaaacttca tttttaattt aaaaggatct
     3001 aggtgaagat cctttttgat aatctcatga ccaaaatccc ttaacgtgag ttttcgttcc
     3061 actgagcgtc agaccccgta gaaaagatca aaggatcttc ttgagatcct ttttttctgc
     3121 gcgtaatctg ctgcttgcaa acaaaaaaac caccgctacc agcggtggtt tgtttgccgg
     3181 atcaagagct accaactctt tttccgaagg taactggctt cagcagagcg cagataccaa
     3241 atactgttct tctagtgtag ccgtagttag gccaccactt caagaactct gtagcaccgc
     3301 ctacatacct cgctctgcta atcctgttac cagtggctgc tgccagtggc gataagtcgt
     3361 gtcttaccgg gttggactca agacgatagt taccggataa ggcgcagcgg tcgggctgaa
     3421 cggggggttc gtgcacacag cccagcttgg agcgaacgac ctacaccgaa ctgagatacc
     3481 tacagcgtga gctatgagaa agcgccacgc ttcccgaagg gagaaaggcg gacaggtatc
     3541 cggtaagcgg cagggtcgga acaggagagc gcacgaggga gcttccaggg ggaaacgcct
     3601 ggtatcttta tagtcctgtc gggtttcgcc acctctgact tgagcgtcga tttttgtgat
     3661 gctcgtcagg ggggcggagc ctatggaaaa acgccagcaa cgcggccttt ttacggttcc
     3721 tggccttttg ctggcctttt gctcacatgt tctttcctgc gttatcccct gattctgtgg
     3781 ataaccgtat taccgccttt gagtgagctg ataccgctcg ccgcagccga acgaccgagc
     3841 gcagcgagtc agtgagcgag gaagcggaag agcgcccaat acgcaaaccg cctctccccg
     3901 cgcgttggcc gattcattaa tgcagctggc acgacaggtt tcccgactgg aaagcgggca
     3961 gtgagcgcaa cgcaattaat gtgagttagc tcactcatta ggcaccccag gctttacact
     4021 ttatgcttcc ggctcgtatg ttgtgtggaa ttgtgagcgg ataacaattt cacacaggaa
     4081 acagctatga ccatgattac gccaagcgcg caattaaccc tcactaaagg gaacaaaagc
     4141 tggagctgca agcttaatgt agtcttatgc aatactcttg tagtcttgca acatggtaac
     4201 gatgagttag caacatgcct tacaaggaga gaaaaagcac cgtgcatgcc gattggtgga
     4261 agtaaggtgg tacgatcgtg ccttattagg aaggcaacag acgggtctga catggattgg
     4321 acgaaccact gaattgccgc attgcagaga tattgtattt aagtgcctag ctcgatacat
     4381 aaacgggtct ctctggttag accagatctg agcctgggag ctctctggct aactagggaa
     4441 cccactgctt aagcctcaat aaagcttgcc ttgagtgctt caagtagtgt gtgcccgtct
     4501 gttgtgtgac tctggtaact agagatccct cagacccttt tagtcagtgt ggaaaatctc
     4561 tagcagtggc gcccgaacag ggacttgaaa gcgaaaggga aaccagagga gctctctcga
     4621 cgcaggactc ggcttgctga agcgcgcacg gcaagaggcg aggggcggcg actggtgagt
     4681 acgccaaaaa ttttgactag cggaggctag aaggagagag atgggtgcga gagcgtcagt
     4741 attaagcggg ggagaattag atcgcgatgg gaaaaaattc ggttaaggcc agggggaaag
     4801 aaaaaatata aattaaaaca tatagtatgg gcaagcaggg agctagaacg attcgcagtt
     4861 aatcctggcc tgttagaaac atcagaaggc tgtagacaaa tactgggaca gctacaacca
     4921 tcccttcaga caggatcaga agaacttaga tcattatata atacagtagc aaccctctat
     4981 tgtgtgcatc aaaggataga gataaaagac accaaggaag ctttagacaa gatagaggaa
     5041 gagcaaaaca aaagtaagac caccgcacag caagcggccg ctgatcttca gacctggagg
     5101 aggagatatg agggacaatt ggagaagtga attatataaa tataaagtag taaaaattga
     5161 accattagga gtagcaccca ccaaggcaaa gagaagagtg gtgcagagag aaaaaagagc
     5221 agtgggaata ggagctttgt tccttgggtt cttgggagca gcaggaagca ctatgggcgc
     5281 agcgtcaatg acgctgacgg tacaggccag acaattattg tctggtatag tgcagcagca
     5341 gaacaatttg ctgagggcta ttgaggcgca acagcatctg ttgcaactca cagtctgggg
     5401 catcaagcag ctccaggcaa gaatcctggc tgtggaaaga tacctaaagg atcaacagct
     5461 cctggggatt tggggttgct ctggaaaact catttgcacc actgctgtgc cttggaatgc
     5521 tagttggagt aataaatctc tggaacagat ttggaatcac acgacctgga tggagtggga
     5581 cagagaaatt aacaattaca caagcttaat acactcctta attgaagaat cgcaaaacca
     5641 gcaagaaaag aatgaacaag aattattgga attagataaa tgggcaagtt tgtggaattg
     5701 gtttaacata acaaattggc tgtggtatat aaaattattc ataatgatag taggaggctt
     5761 ggtaggttta agaatagttt ttgctgtact ttctatagtg aatagagtta ggcagggata
     5821 ttcaccatta tcgtttcaga cccacctccc aaccccgagg ggacccgaca ggcccgaagg
     5881 aatagaagaa gaaggtggag agagagacag agacagatcc attcgattag tgaacggatc
     5941 tcgacggtat cggttaactt ttaaaagaaa aggggggatt ggggggtaca gtgcagggga
     6001 aagaatagta gacataatag caacagacat acaaactaaa gaattacaaa aacaaattac
     6061 aaaaattcaa aattttatcg atgagtaatt catacaaaag gactcgcccc tgccttgggg
     6121 aatcccaggg accgtcgtta aactcccact aacgtagaac ccagagatcg ctgcgttccc
     6181 gccccctcac ccgcccgctc tcgtcatcac tgaggtggag aagagcatgc gtgaggctcc
     6241 ggtgcccgtc agtgggcaga gcgcacatcg cccacagtcc ccgagaagtt ggggggaggg
     6301 gtcggcaatt gaaccggtgc ctagagaagg tggcgcgggg taaactggga aagtgatgtc
     6361 gtgtactggc tccgcctttt tcccgagggt gggggagaac cgtatataag tgcagtagtc
     6421 gccgtgaacg ttctttttcg caacgggttt gccgccagaa cacaggtaag tgccgtgtgt
     6481 ggttcccgcg ggcctggcct ctttacgggt tatggccctt gcgtgccttg aattacttcc
     6541 acgcccctgg ctgcagtacg tgattcttga tcccgagctt cgggttggaa gtgggtggga
     6601 gagttcgagg ccttgcgctt aaggagcccc ttcgcctcgt gcttgagttg aggcctggct
     6661 tgggcgctgg ggccgccgcg tgcgaatctg gtggcacctt cgcgcctgtc tcgctgcttt
     6721 cgataagtct ctagccattt aaaatttttg atgacctgct gcgacgcttt ttttctggca
     6781 agatagtctt gtaaatgcgg gccaagatct gcacactggt atttcggttt ttggggccgc
     6841 gggcggcgac ggggcccgtg cgtcccagcg cacatgttcg gcgaggcggg gcctgcgagc
     6901 gcggccaccg agaatcggac gggggtagtc tcaagctggc cggcctgctc tggtgcctgg
     6961 cctcgcgccg ccgtgtatcg ccccgccctg ggcggcaagg ctggcccggt cggcaccagt
     7021 tgcgtgagcg gaaagatggc cgcttcccgg ccctgctgca gggagctcaa aatggaggac
     7081 gcggcgctcg ggagagcggg cgggtgagtc acccacacaa aggaaaaggg cctttccgtc
     7141 ctcagccgtc gcttcatgtg actccacgga gtaccgggcg ccgtccaggc acctcgatta
     7201 gttctcaagc ttttggagta cgtcgtcttt aggttggggg gaggggtttt atgcgatgga
     7261 gtttccccac actgagtggg tggagactga agttaggcca gcttggcact tgatgtaatt
     7321 ctccttggaa tttgcccttt ttgagtttgg atcttggttc attctcaagc ctcagacagt
     7381 ggttcaaagt ttttttcttc catttcaggt gtcgtgaccc tagcgctacc tctagagcca
     7441 ccatgagagt gatggcccct agagccctgc tgctgctgct gagcggaggc ctggccctga
     7501 ccgagacatg ggcctgctcc cactctatgc gctatttcga caccgccgtg tcccggcccg
     7561 gaagaggaga gcctcgcttt atctctgtgg gctacgtgga cgatacacag ttcgtgcggt
     7621 ttgacagcga tgcagcatcc cctaggggag agccaagggc accctgggtg gagcaggagg
     7681 gcccagagta ttgggaccgg gagacacaga agtacaagag gcaggcccag gccgatcgcg
     7741 tgtctctgag gaacctgcgc ggctactata atcagtctga ggacggcagc cacacactgc
     7801 agagaatgag cggatgtgac ctgggaccag atggccggct gctgagaggc tacgatcaga
     7861 gcgcctatga cggcaaggat tacatcgccc tgaacgagga cctgaggtcc tggaccgcag
     7921 cagatacagc agcacagatc acccagcgga agctggaggc agcaagagca gcagagcagc
     7981 tgagggccta tctggaggga acctgcgtgg agtggctgcg gagatacctg gagaatggca
     8041 aggagacact gcagagggca gagccaccta agacccacgt gacacaccac ccactgtccg
     8101 accacgaggc caccctgagg tgttgggcac tgggcttcta tcccgccgag atcaccctga
     8161 catggcagag agacggcgag gatcagaccc aggacacaga gctggtggag acacggcccg
     8221 caggcgatgg cacatttcag aagtgggcag cagtggtggt gccttccgga caggagcaga
     8281 ggtacacctg ccacatgcag cacgagggcc tgcaggagcc actgaccctg tcttgggagc
     8341 ccagctccca gcctacaatc ccaatcatgg gcatcgtggc aggcctggcc gtgctggtgg
     8401 tgctggccgt gctgggcgcc gtggtgaccg ccatgatgtg caggaggaag tctagcggag
     8461 gcaagggagg aagctgctcc caggccgcct gttctaacag cgcccagggc tccgatgagt
     8521 ctctgatcac ctgtaaggcc tgaggcgcgc ctcccagagc caccgttaca ccgagtcgga
     8581 gatgacttcc gcaccgactc ggtgccactt tttcaagttg ataacggact agccttattt
     8641 taacttgcta tttctagctc taaaactttt tttgactaga tcttgagaca aatggcagta
     8701 ttcatccaca aggtacgcta aacgcctgca accaggacgc gt
//
```
